# Molecular dynamics simulations provide insights into ULK-101 potency and selectivity toward autophagic kinases ULK1/2

**DOI:** 10.1101/2023.12.01.569261

**Authors:** Robert M. Vaughan, Bradley M. Dickson, Katie R. Martin, Jeffrey P. MacKeigan

## Abstract

Kinase domains are highly conserved within protein kinases in both sequence and structure. Many factors, including phosphorylation, amino acid substitutions or mutations, and small molecule inhibitor binding, influence conformations of the kinase domain and enzymatic activity. The serine/threonine kinases ULK1 and ULK2 are highly conserved with N- and C-terminal domains, phosphate-binding P-loops, -αC-helix, regulatory and catalytic spines, and activation loop DFG and APE motifs. Here, we performed molecular dynamics (MD) simulations to understand better the potency and selectivity of the ULK1/2 small molecule inhibitor, ULK-101. We observed stable bound states for ULK-101 to the adenosine triphosphate (ATP)-binding site of ULK2, coordinated by hydrogen bonding with the hinge backbone and the catalytic lysine sidechain. Notably, ULK-101 occupies a hydrophobic pocket associated with the N-terminus of the -αC-helix. Large movements in the P-loop are also associated with ULK-101 inhibitor binding and exit from ULK2. Our data further suggests that ULK-101 could induce a folded P-loop conformation and hydrophobic pocket reflected in its nanomolar potency and kinome selectivity.

## Introduction

Macroautophagy, commonly referred to as autophagy, is an essential cellular recycling process^1^. It begins with the formation of phagophores, which evolve into double-membraned autophagosomes, engulfing cytoplasmic components. These autophagosomes then merge with lysosomes, where their contents are broken down into amino acids, lipids, and carbohydrates, available for reuse by the cell^2^. This efficient system helps cells survive during stressful conditions, such as nutrient scarcity. Autophagy plays a complex role in cancer, particularly in KRAS-driven tumors^3,4^. Consequently, inhibiting autophagy is now being explored to enhance current cancer treatments^5^.

Autophagy is initiated by the serine/threonine kinases ULK1/2 (unc-51 like autophagy initiating kinase 1/2), mammalian equivalents of yeast ATG1^6^. ULK1/2 forms a complex with ATG13, RB1CC1 (also known as FIP200), and ATG101. This complex allows ULK1/2 to integrate signals from nutrient-sensing (mTORC1) and energy-sensing (AMPK) pathways^7^, triggering the early stages of autophagic membrane production. The critical role of ULK1/2 in autophagy has been demonstrated in studies showing that depleting or inhibiting ULK1/2 impairs autophagy, making them targets for autophagy inhibition. Accordingly, several small molecule inhibitors of ULK1/2 have been developed, offering insights into its structure and the effectiveness of ULK1/2 inhibition in suppressing autophagy^8–11^. However, these inhibitors vary in potency, specificity, and cellular activity. ULK-101 shows improved potency and selectivity compared to existing inhibitors, and effectively blocks autophagy induction and reduces autophagic flux in various conditions^8^. Here, we focus on the small molecule inhibitor, ULK-101, in molecular dynamics (MD) simulations and report the first MD simulations of ULK2 binding with ULK-101.

## Results

In order to understand molecular details of the high-affinity interactions between ULK-101 (Fig. 1A) and the serine/threonine kinases ULK1 (Fig. 1C) and ULK2 (Fig. 1D), we obtained the existing X-ray crystal structures solved with small molecule inhibitors (ULK1, PDB# 4WNO; ULK2, PDB# 6YID). Active states were captured for each ULK kinase, highlighted by DFG-in conformation and phosphorylated Thr-180 (p-T180) in ULK1 and the phosphomimetic Asp-173 substitution in ULK2. Important structural features of the kinase domains are highlighted (Fig. 1C,D). Overall, the ULK1 and ULK2 kinases share a high degree of structural similarity, with ULK-101 inhibiting ULK1 and ULK2 at nanomolar biochemical activity^8^. ULK-101 inhibits ULK1 with an IC_50_ of 8.3 nM (95% CI: 7.2–9.6 nM) and ULK2 with an IC_50_ of 30 nM (95% CI: 26–35 nM) for ULK2 (Fig. 1B, data from ref^8^).

**Figure 1.**
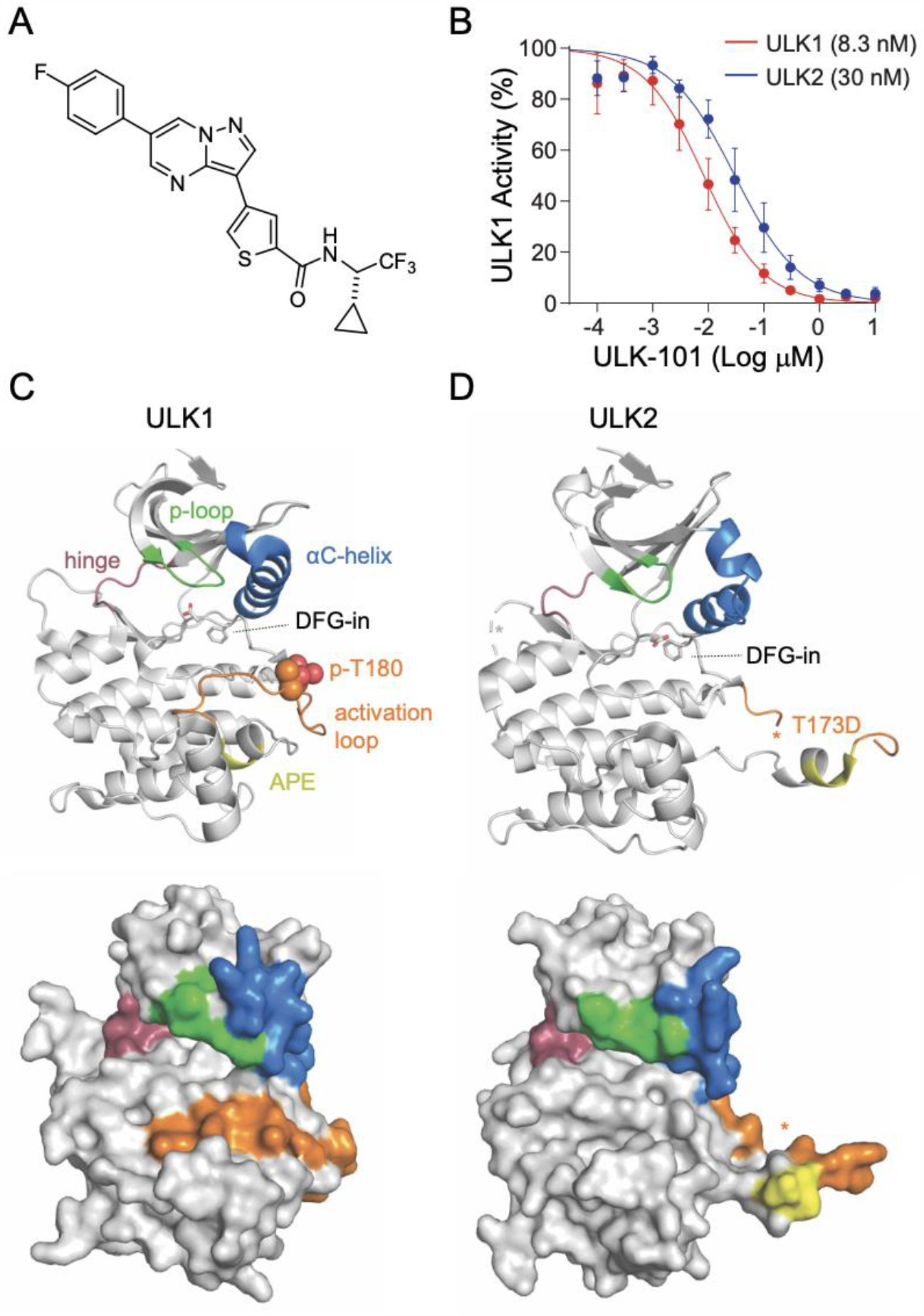
ULK-101 is a potent inhibitor of ULK1 and ULK2 kinases. (**A**) Chemical structure of ULK-101. (**B**) ULK1 (red) and ULK2 (blue) kinase activity was measured in the presence of ULK-101 at half-log dilutions from top concentrations of 10 μM, as we have previously reported^8^. Data are represented as mean activity normalized to control (0 μM inhibitor) ± SD. Solid lines represent IC_50_ curves fit by non-linear regression. (C,D) Cartoon (top) or surface representations (bottom) of (**C**) ULK1 (PDB# 4WNO) or (**D**) ULK2 (lPDB# 6YID) with ligands removed, and * denotes unresolved regions.

To understand how ULK-101 interacts with ULK1/2, we performed all-atom MD simulations with a single molecule of both ULK-101 and ULK2 kinase domain (PDB# 6YID) corresponding to amino acids 1-276 of ULK2 (Fig. 2A). Noteworthy, is that ULK-101 molecular dynamics simulation was not started in the ATP-binding site nor directed toward the active site. In order to avoid sampling unreliable or weakly bound states, we utilized adaptive biasing with overfill protection as implemented by fABMACS^12,13^. The free energy of the system is shown with respect to two different collective variables (Fig. 2B). Each collective variable consisted of three atoms on either end of ULK-101, and the root mean square deviation (RMSD) was calculated relative to the predicted binding pocket. In other words, the closer each collective variable (CV_1_, CV_2_) was to 0, the closer ULK-101 was bound to the active site of ULK2. In one MD simulation, totaling 790 ns, we achieved stable poses for ULK-101 near the ATP-binding pocket of the ULK2 kinase domain (Fig. 2C), occurring roughly between 400 and 600 ns of simulation time. Indeed, the most stable poses (lowest free energy) occurred with ULK-101 near the known ATP-binding site (Fig. 2D).

**Figure 2.**
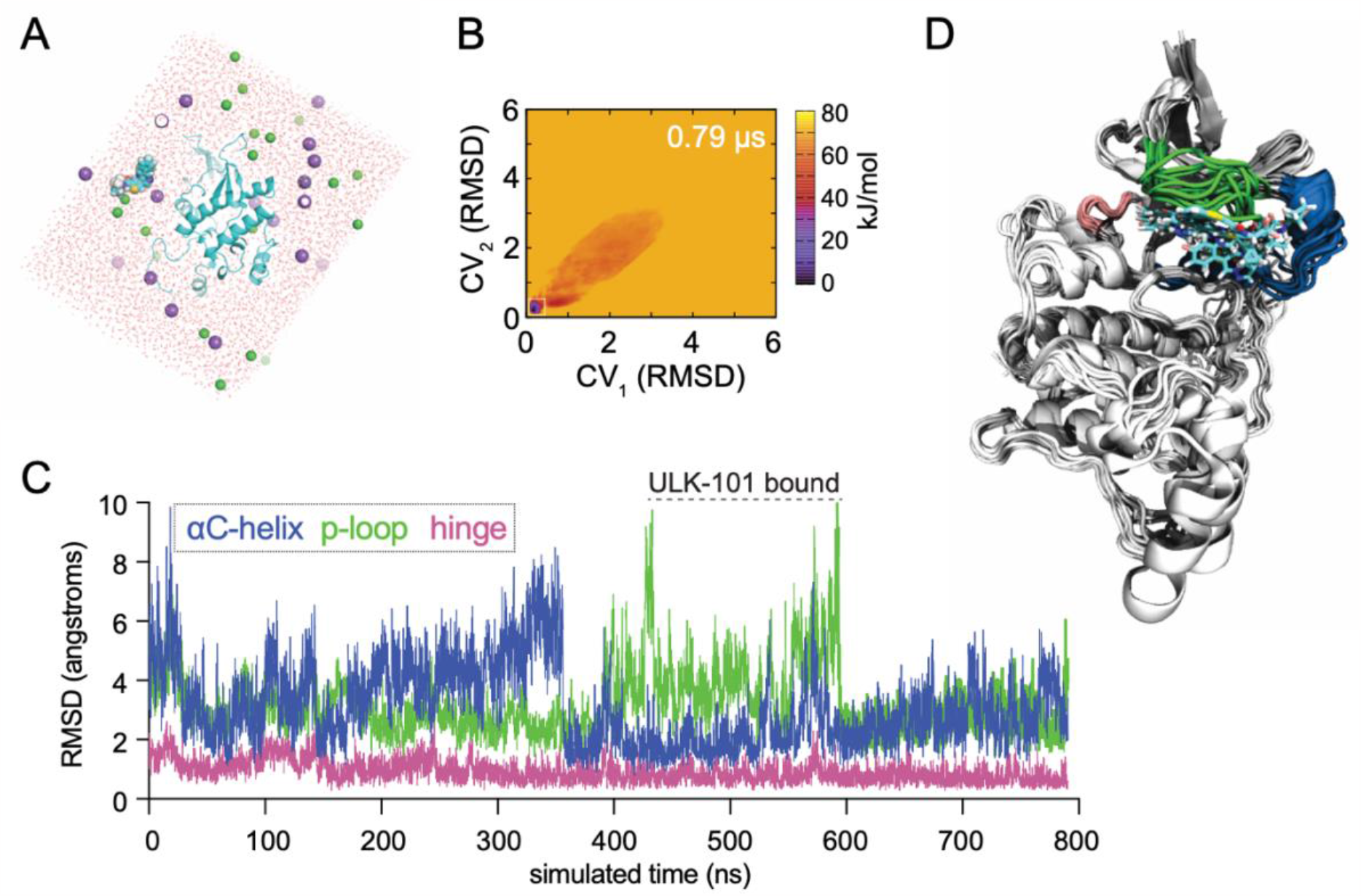
Molecular dynamics simulations show stable bound states for ULK-101 with ULK2. (**A**) Beginning state of ULK-101 with ULK2 kinase for all-atom molecular dynamics simulation; ULK2 is shown in cyan cartoon, ULK-101 in cyan spheres, water molecules in red, and Na^+^ and Cl^-^ in purple and green spheres, respectively. (**B**) Free energy calculation of the system shown in 2A as a function of ULK-101 location. RMSD is for each CV relative to the predicted three-atom collective variables in ULK-101, relative to the predicted binding pocket. (**C**) RMSD of selected kinase features as a function of simulated time; see methods for exact residues. Stable bound poses of ULK-101 are indicated between 400-600 ns. (**D**) Cartoon representation of 10 ns steps from the poses with ULK-101 bound near the canonical inhibitor binding pocket in ULK2, aligned via C-terminal helices.

To further elucidate critical factors for ULK-101 binding to ULK2, we first identified hydrogen bonds between the two molecules as a function of simulated time (Fig. 3A). The two most frequently occurring hydrogen bonds were between the backbone of Cys-88 (C88) and the fluorobenzene on ULK-101, and between the sidechain of the catalytic Lys-39 (K39) and the carbonyl oxygen on ULK-101 (Fig. 3A), occurring in 2.4% and 1.9% of all frames, respectively. It’s noteworthy that the initial states with two or three hydrogen bonds (near time = 0 ns) are not more frequently sampled, demonstrating the utility of adaptive biasing and avoidance of unreliable states. To confirm the states where the ligand ULK-101 was in the binding pocket of ULK2, we measured the distance between the gatekeeper residue Met-85 (M85) and the carbon attached to the fluorine on the fluorobenzene ring, as well as from the beta carbon of Ile-42 (I42) to a carbon on the small cyclopropane ring (Fig. 3B). These two measurements indicate the location of ULK-101 to the left side (ATP binding) or right side (hydrophobic region) of the pocket in ULK2, respectively. ULK-101 made contact with the hydrophobic pocket residues before entry and after exit from the ULK2 binding pocket, indicating that the hydrophobic pocket contributes to the stability of the interaction between ULK2 and ULK-101 in this simulation. In addition to insertion into the hydrophobic pocket, a P-loop rearrangement is required for ULK-101 entry and binding of ULK2 (Fig. 3C).

**Figure 3.**
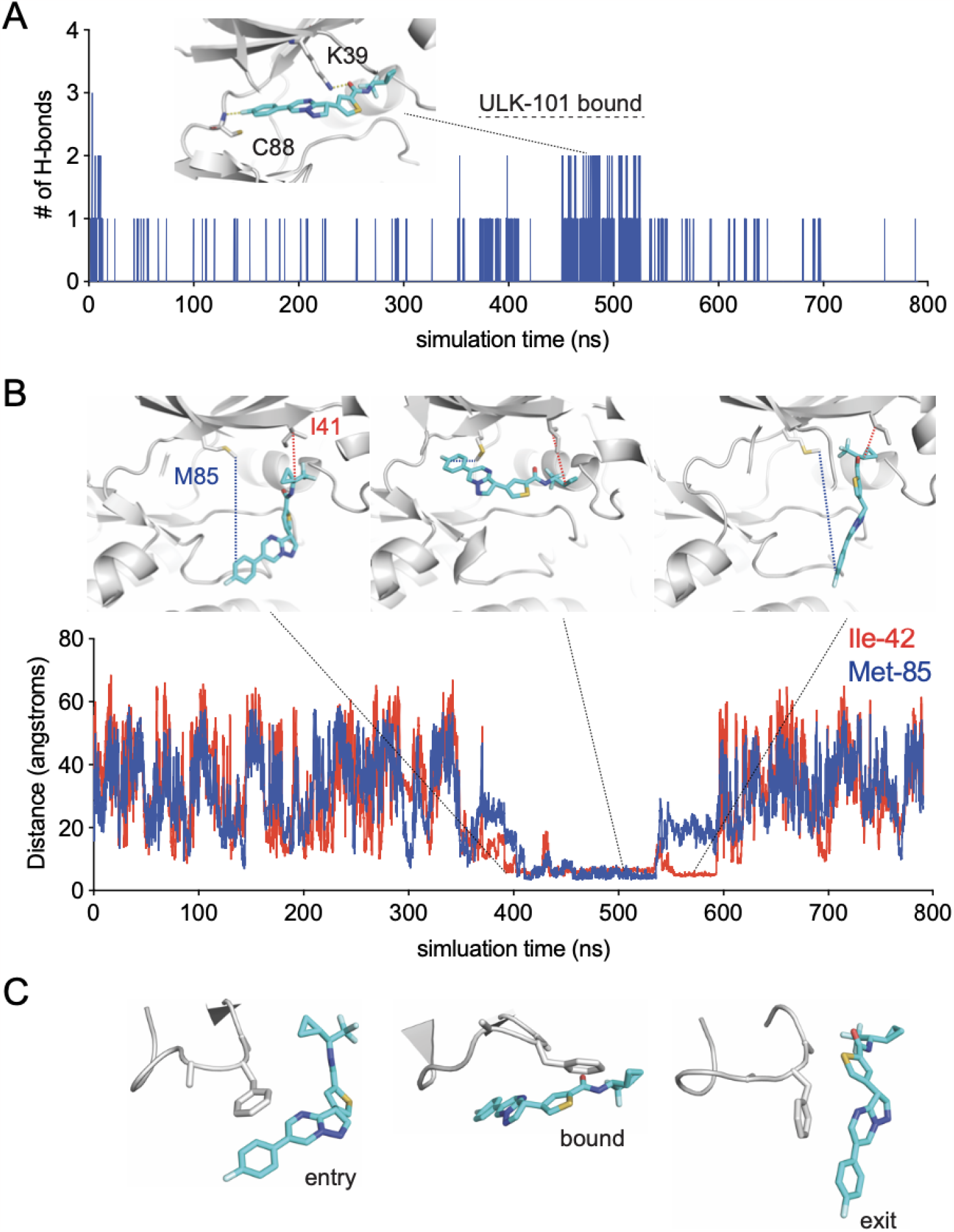
Molecular details of ULK-101 recognition by ULK2 kinase. (**A**) Number of hydrogen bonds formed between ULK-101 and ULK2 as a function of simulated time; inset shows the two major hydrogen bonds formed between the indicated ULK-101 atoms and the backbone of C88 or the sidechain of K39. (**B**) Distance between indicated carbon atoms of ULK-101 and either Met-85 (blue) or Ile-42 (red) as a function of simulated time. (**C**) Examples of P-loop conformations from this simulation for indicated events, same frames shown as in 3B.

The P-loop rearrangement is visible when comparing the starting state of the simulation (relaxed and equilibrated) with a stable bound state (Fig. 4A). In the unbound, or t = 0 ns frame, ULK2 has an extended P-loop conformation. When a stable pose is achieved, the P-loop is more collapsed or folded into the ATP-binding area. Comparison of a stable ULK2 pose with bound ULK-101 pose to the crystal structures of ULK2 with two structurally distinct small molecule inhibitors, SBI-0206965 or MRT68921, illustrates the alternate P-loop conformation, highlighted by large shifts in residues between His-17 (H17) and Phe-20 (F20) (Fig. 4B). A comparison of ULK-101 and MRT68921 in ULK2 revealed differences in the space occupied by each ligand (Fig. 4C). MRT68921 occupies space toward the ATP binding site and may stabilize the P-loop in an extended conformation. The contacts required to stabilize the extended P-loop conformation were missing from our ULK-101 poses. Rather, the P-loop was free to collapse and remain flexible. As part of ULK-101 sits in a hydrophobic back pocket (Fig. 4D), a feature that is unique among resolved ULK1/2 kinase inhibitors, we posit that a combination of the occupancy of the hydrophobic pocket and a folded P-loop contributes to ULK1/2 kinase selectivity for ULK-101^8^.

**Figure 4.**
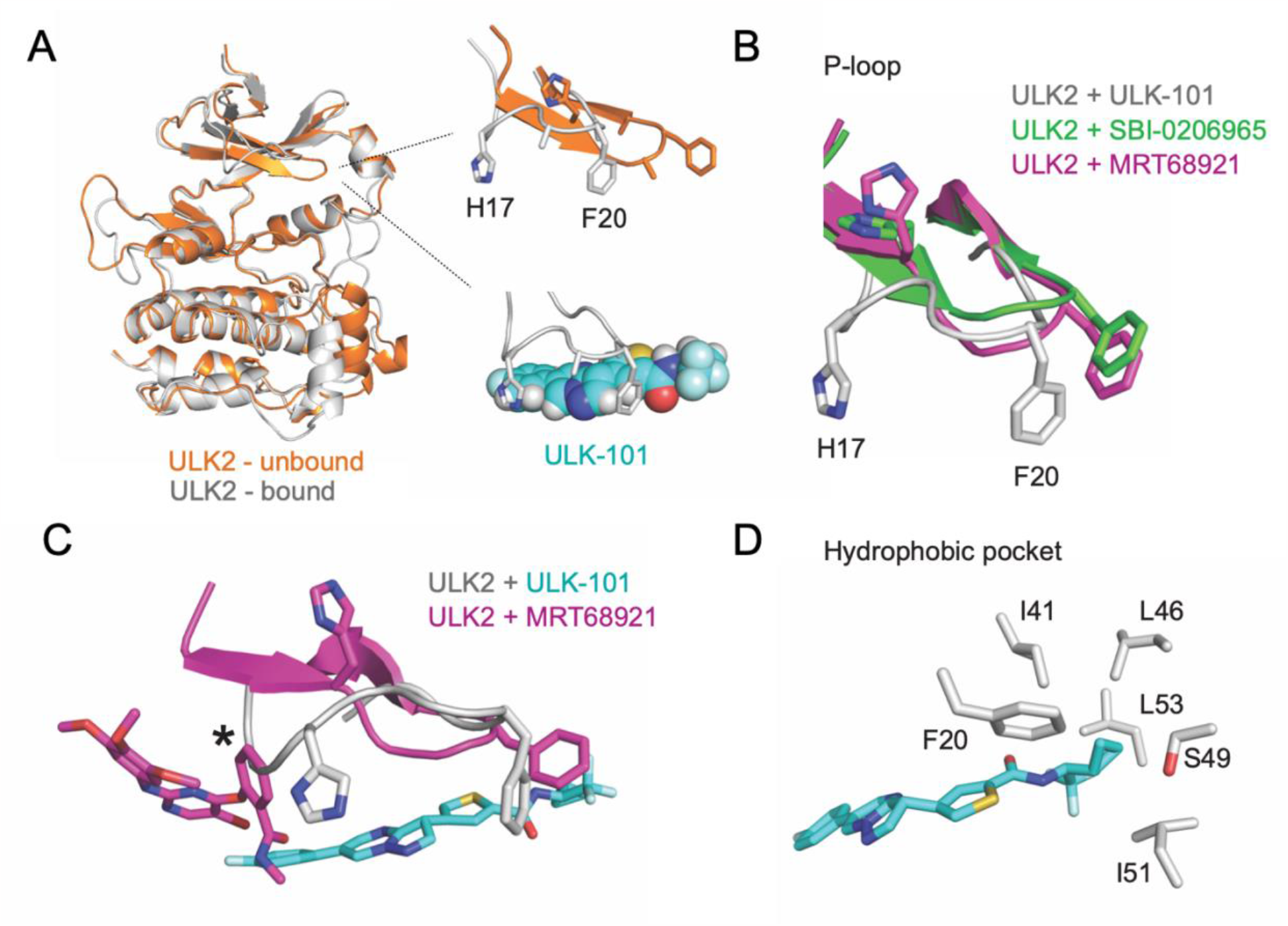
ULK-101 bound with a folded P-loop conformation and hydrophobic pocket. (**A**) Comparison of ULK2 structures at the start of simulation (orange, unbound) and bound to ULK-101 (gray), highlighting a shift from an extended to a folded P-loop conformation. (**B**) Overlay of ULK2 P-loops from ULK2 bound to ULK-101 (gray), ULK2 bound to SBI-0206965 (green), or ULK2 bound to MRT68921 (pink) highlighting two bulky P-loop sidechains (H17, F20) with the ligands masked. (**C**) Same as in 4B, but with ligands shown for selected models, after alignment with residues 5 to 45; * denotes clash with MRT68921. (**D**) Sidechains of residues within 4 angstroms of ULK-101 and involved in a hydrophobic pocket are shown.

## Discussion

To understand the relative selectivity of ULK-101 as an inhibitor of ULK1/2, we set up a host of simulations between ULK1, ULK2, and other known kinase inhibitors. The various experiments were between 500 to 2500 ns in total of adaptively biased simulation time. Of those efforts, we achieved a stable bound pose for ULK-101 with ULK2 that lasted roughly 120 ns from a single 790 ns simulation. Notably, the entry and exit of ULK-101 interaction with ULK2 appears driven largely by a hydrophobic pocket in the -αC-helix, which carries a bend at the N-terminus in both ULK1 (Fig. 1C) and ULK2 (Fig. 1D). We speculate that our findings in ULK2 may hold true in ULK1 based on homology to ULK2 and shared high-affinity for ULK-101.

However, this non-exhaustive list of references demonstrates remaining challenges in dissecting precise contributions of ULK kinases to various processes in various systems^14–17^. What remains, however, is that high affinity ligands for ULK1 maintain high affinity for ULK2^8–10^.

In line with other kinase inhibitors, ULK-101 would be predicted to be ATP competitive as it appears coordinated by hydrogen bonded in the hinge region and catalytic lysine (Fig. 3A). We hypothesize that the hydrophobic pocket that was bound by ULK-101 in ULK2, explains some selectivity for the ULK kinases. The pocket is formed by residues of the N-terminus of the -αC-helix and is responsible for binding the cyclopropyl or the trifluoromethyl groups of ULK-101. This position is supported by interactions with the P-loop, a behavior that is perhaps anticipated, as kinase inhibitors often emulate and compete with nucleotides^18^, and the P-loop is involved in nucleotide recognition^19^. Further, the P-loop undergoes a conformational change to accommodate ULK-101 binding (Fig. 2C, 3C, 4A, 4B). Our simulations begin with ULK2 P-loop in an extended conformation (Fig. 4A, orange), but quickly folds or collapses into the ATP pocket; this is, of course, in the absence of the small molecule, SBI-0206965^11^, as it was removed from the crystal structure model prior to relaxation and equilibration of the simulation system.

Mutation of the gatekeeper residue is a mechanism of acquired resistance to kinase inhibitors and clinically problematic^20^. Accordingly, inhibitors are being developed in light of specific and prevalent gatekeeper mutations^21^. Suppose the positioning of ULK-101 in our MD simulations is correct or partially correct. In that case, it positions our ligand, on average, further from the ULK2 gatekeeper (Met-85) than other solved structures for ULK2 (PDB# 6QAT, 6QAU, 6QAV, 6YID, only one ligand shown in Fig. 4C). The increased distance between gatekeeper and ligand, potentially as a result of insertion in the hydrophobic back pocket, may reduce ULK-101 sensitivity to gatekeeper mutations. Further, this work provides the structural framework for the identification of additional novel small molecule inhibitors capable of binding the hydrophobic back pocket with a folded P-loop, which could be a novel route to targeting ULK1/2.

## Methods

### Visualization of atomic structures

Visual representation of existing structural models (ULK1, PDB# 4WNO; ULK2, PDB# 6YID) was performed with PyMOL^22^. Visuals related to trajectories were generated in VMD (https://www.ks.uiuc.edu/Research/vmd/).

### Biochemical ULK-101 inhibition of ULK1/2 kinases

Eurofins Pharma Discovery Services UK Limited performed all kinase assays, as we have previously reported^8^. Briefly, IC_50_ data for ULK1 and ULK2 was generated using 10-point IC_50_ Profiler assays with half-log dilutions from top concentrations of 10 μM (4 replicates) and 1 μM (4 replicates), giving 8 data points for most concentrations in the curve. The percent activity remaining and percent inhibition were calculated from negative control wells. GraphPad Prism was used for IC_50_ determinations by fitting curves with variable slope (four-parameter) non-linear regression models using top and bottom constraints of 100% and 0%, respectively.

### Preparation of ULK2 kinase domain and ULK-101 for simulation

All naming and numbering are according to UniProt. The X-ray crystal of ULK2 in complex with inhibitor SBI-0206965 (PDB# 6YID, ref^11^) was the structural basis for our simulation. We began with the coordinates of a monomeric ULK2 kinase domain (a.a. 1-276) that was engineered with a phosphomimetic T173D in the activation loop. The inhibitor and water molecules were removed from the crystal structure, and unresolved loop residues were modeled with Chimera v1.15^23^. The refined ULK2 kinase was brought into GROMACS v5.0.5^24^ with the CHARMM 36 force field^25^. To prepare ULK-101 for simulation, the inhibitor was first converted from SMILES to .mol2 and then parameterized using SwissParam^26,27^. The coordinates for ULK-101 and ULK2 were combined (ULK-101 out of the kinase pocket) and solvated in TIP3P water with 150 nM NaCl, followed by steepest descent for 5,000 steps (converged to Fmax < 1000 in 1425 steps), a 0.1 ns relaxation in the canonical ensemble, and then 5 ns pressure equilibration in the isothermal-isobaric ensemble. Free energy of ULK-101 binding to ULK2 was computed with adaptive biasing potential and overfill protection^12,13^ in the canonical ensemble with the adaptive biasing potential parameters c = 0.01, b = 0.9, α = 8 and an overfill limit of 70 kJ/mol. See Supplemental Data for necessary files and parameters to set up the simulation and run fABMACS.

### Representation of molecular dynamics simulations

To represent free energy calculation as a function of ULK-101 location, two groups of three atoms were selected on each end of ULK-101 as collective variables (CV_1_, CV_2_,). RMSD were calculated for each CV relative to an approximation of the ATP binding pocket (x=0, y=0, z=0). The RMSD Trajectory Tool in VMD was used to measure the RMSD of the following kinase features: P-loop (ULK1, a.a. 23-29; ULK2, a.a. 17-21), -αC-helix (ULK1, a.a. 50-68; ULK2, a.a. 43-62), hinge region (ULK1, a.a. 93-98; ULK2, a.a. 86-91), after alignment of all frames to the top frame (based on a.a. 200-260, the relatively stable C-terminus of ULK2 kinase domain). The Hydrogen Bond tool in VMD was used to identify and count the frequency of hydrogen bonds made between ULK2 and ULK-101 with a distance and angle cutoff of 3 angstroms and 20 degrees, respectively. The distance between ULK-101 and indicated atoms was generated using the labels function in VMD.

## Author contributions

Designing research studies: R.M.V., K.R.M., J.P.M.; conducting experiments: R.M.V., B.M.D.; analyzing data: R.M.V., B.M.D., K.R.M., J.P.M.; writing the manuscript: R.M.V., J.P.M.; acquisition of funding: R.M.V., J.P.M.; final approval of this manuscript: R.M.V., B.M.D., K.R.M., J.P.M.

## Acknowledgments

R.M.V. (K00CA245821) and J.P.M. (R21CA270588, R21CA252430, and R21CA263133) have research support from the National Cancer Institute. This work was partly supported through computational resources and services provided by the Institute for Cyber-Enabled Research at Michigan State University.

