## Supplemental Methods for "Molecular dynamics simulations provide insights into ULK-101 potency and selectivity toward autophagic kinases ULK1/2"

- Example code for how these simulations were run on a cluster
- Required input files to fun fABMACS. https://github.com/BradleyDickson/fABMACS
- proc.pdb – coordinate file for refined ULK2 and ULK-101 for simulation

***Code for equilibration with GROMACS and running fABMACS.***

#build system and relax/equilibrate with GROMACS

```
pdb2gmx_mpi -f FILE -o proc.pdb -water TIP3P -ignh -ter
#where FILE is just your clean protein
#add ligand pdb ATOM after TER in proc.pdb followed by ENDMDL
#add to topol.top  #include "ligand.itp" after charm in forcefield parameters and add ligand.itp
to working dir and add ligand to [ molecules ] section

editconf_mpi -f proc.pdb -o boxed.gro -c -d 1 -bt triclinic

gmx_mpi solvate -cp boxed.gro -cs spc216.gro -o solv.gro -p topol.top

grompp_mpi -f ~/GROrep/minim.mdp -c solv.gro -p topol.top -o ions.tpr

genion_mpi -s ions.tpr -o solv_ions.gro -p topol.top -pname NA -nname CL -neutral -conc 0.15

grompp_mpi -f ~/GROrep/minim.mdp -c solv_ions.gro -p topol.top -o em.tpr

mpirun -np 120 /opt/software/GROMACS/5.0.5-intel-2015a-hybrid/bin/mdrun_mpi -deffnm em

grompp_mpi -f ~/GROrep/nvt.mdp -c em.gro -p topol.top -o canon.tpr

mpirun -np 120 /opt/software/GROMACS/5.0.5-intel-2015a-hybrid/bin/mdrun_mpi -deffnm canon

grompp_mpi -f ~/GROrep/npt.mdp -c canon.gro -p topol.top -o isob.tpr

mpirun -np 120 /opt/software/GROMACS/5.0.5-intel-2015a-hybrid/bin/mdrun_mpi -deffnm isob
```

#build fABMACS

```
#git clone https://github.com/BradleyDickson/fABMACS.git
awk '{print $2/10,$3/10,$4/10}' listIndex > Reference
#the +0 or +1 is for if you use the atom # after selecting
awk '{print $1+0}' listIndex > fABMACS/list
tail -1 isob.gro |awk '{print $1/2"\n"$2/2"\n"$3/2}' > sphpoints

module load ifort/2015.1.133-GCC-4.9.2  impi/5.0.2.044
module load GROMACS/5.0.5-hybrid
module load cmake

cd fABMACS
./PATCHscript.sh 480 6  3  3 6 5.5 OVERFILL
cd ..
mkdir fBuild
mkdir fBin
cd fBuild
cmake ../fABMACS -DGMX_BUILD_OWN_FFTW=ON -DGMX_SIMD=AVX2_256 -DGMX_OPENMP=OFF -DGMX_MPI=ON -
DCMAKE_INSTALL_PREFIX=../fBin
make -j24
make install
cd ..
```

#run fABMACS

```
grompp_mpi -f ~/GROrep/md.mdp -c isob.gro -p topol.top -o run1.tpr
```

```
mpirun -np 120 PATHtoFAB/mdrun_mpi -deffnm run1
```

#params

```
300.0
0.9
0.05
10.0
7.17029   9.22826   7.87935
4.6
20.0
0
70.0
```

#sphpoints

```
3.58514
4.61413
3.93967
```

#listIndex (atoms from ULK-101 for collective variables)

```
4432 27.857 38.636 37.254
4433 27.977 39.566 36.254
4434 28.807 39.156 35.224
4447 27.847 47.316 34.714
4448 27.237 46.086 34.904
4450 27.937 48.206 36.184
```

#Reference

```
2.7857 3.8636 3.7254
2.7977 3.9566 3.6254
2.8807 3.9156 3.5224
2.7847 4.7316 3.4714
2.7237 4.6086 3.4904
2.7937 4.8206 3.6184
```

#backs.itp non-mobile atoms to restrain protein in space

```
; position restraints for C-alpha of Protein in water

[ position_restraints ]
;  i funct       fcx        fcy        fcz
1686 1 1000 1000 1000
1687 1 1000 1000 1000
1688 1 1000 1000 1000
1689 1 1000 1000 1000
1690 1 1000 1000 1000
1691 1 1000 1000 1000
1692 1 1000 1000 1000
1693 1 1000 1000 1000
1694 1 1000 1000 1000
1695 1 1000 1000 1000
```

| | | | | |
|---|---|---|---|---|
| 1696 | 1 | 1000 | 1000 | 1000 |
| 1697 | 1 | 1000 | 1000 | 1000 |
| 1698 | 1 | 1000 | 1000 | 1000 |
| 1699 | 1 | 1000 | 1000 | 1000 |
| 1700 | 1 | 1000 | 1000 | 1000 |
| 1701 | 1 | 1000 | 1000 | 1000 |
| 1702 | 1 | 1000 | 1000 | 1000 |
| 1703 | 1 | 1000 | 1000 | 1000 |
| 1704 | 1 | 1000 | 1000 | 1000 |
| 1705 | 1 | 1000 | 1000 | 1000 |
| 1706 | 1 | 1000 | 1000 | 1000 |
| 1707 | 1 | 1000 | 1000 | 1000 |
| 1708 | 1 | 1000 | 1000 | 1000 |
| 1709 | 1 | 1000 | 1000 | 1000 |
| 1710 | 1 | 1000 | 1000 | 1000 |
| 1711 | 1 | 1000 | 1000 | 1000 |
| 1712 | 1 | 1000 | 1000 | 1000 |
| 1713 | 1 | 1000 | 1000 | 1000 |
| 1714 | 1 | 1000 | 1000 | 1000 |
| 1715 | 1 | 1000 | 1000 | 1000 |
| 1716 | 1 | 1000 | 1000 | 1000 |
| 1717 | 1 | 1000 | 1000 | 1000 |
| 1718 | 1 | 1000 | 1000 | 1000 |
| 1719 | 1 | 1000 | 1000 | 1000 |
| 1720 | 1 | 1000 | 1000 | 1000 |
| 1721 | 1 | 1000 | 1000 | 1000 |
| 1722 | 1 | 1000 | 1000 | 1000 |
| 1723 | 1 | 1000 | 1000 | 1000 |
| 1724 | 1 | 1000 | 1000 | 1000 |
| 1725 | 1 | 1000 | 1000 | 1000 |
| 1726 | 1 | 1000 | 1000 | 1000 |
| 1727 | 1 | 1000 | 1000 | 1000 |
| 1728 | 1 | 1000 | 1000 | 1000 |
| 1729 | 1 | 1000 | 1000 | 1000 |
| 1730 | 1 | 1000 | 1000 | 1000 |
| 1731 | 1 | 1000 | 1000 | 1000 |
| 1732 | 1 | 1000 | 1000 | 1000 |
| 1733 | 1 | 1000 | 1000 | 1000 |
| 1734 | 1 | 1000 | 1000 | 1000 |
| 1735 | 1 | 1000 | 1000 | 1000 |
| 1736 | 1 | 1000 | 1000 | 1000 |
| 1737 | 1 | 1000 | 1000 | 1000 |
| 1738 | 1 | 1000 | 1000 | 1000 |
| 1739 | 1 | 1000 | 1000 | 1000 |
| 1740 | 1 | 1000 | 1000 | 1000 |
| 1741 | 1 | 1000 | 1000 | 1000 |
| 1742 | 1 | 1000 | 1000 | 1000 |
| 1743 | 1 | 1000 | 1000 | 1000 |
| 1744 | 1 | 1000 | 1000 | 1000 |
| 1745 | 1 | 1000 | 1000 | 1000 |
| 1746 | 1 | 1000 | 1000 | 1000 |
| 1747 | 1 | 1000 | 1000 | 1000 |
| 1748 | 1 | 1000 | 1000 | 1000 |
| 1749 | 1 | 1000 | 1000 | 1000 |
| 1750 | 1 | 1000 | 1000 | 1000 |
| 1751 | 1 | 1000 | 1000 | 1000 |
| 1752 | 1 | 1000 | 1000 | 1000 |
| 1753 | 1 | 1000 | 1000 | 1000 |
| 1754 | 1 | 1000 | 1000 | 1000 |
| 1755 | 1 | 1000 | 1000 | 1000 |
| 1756 | 1 | 1000 | 1000 | 1000 |
| 1757 | 1 | 1000 | 1000 | 1000 |
| 1758 | 1 | 1000 | 1000 | 1000 |
| 1759 | 1 | 1000 | 1000 | 1000 |
| 1760 | 1 | 1000 | 1000 | 1000 |
| 1761 | 1 | 1000 | 1000 | 1000 |
| 1762 | 1 | 1000 | 1000 | 1000 |
| 1763 | 1 | 1000 | 1000 | 1000 |
| 1764 | 1 | 1000 | 1000 | 1000 |
| 1765 | 1 | 1000 | 1000 | 1000 |
| 1766 | 1 | 1000 | 1000 | 1000 |

| | | | | |
|---|---|---|---|---|
| 1767 | 1 | 1000 | 1000 | 1000 |
| 1768 | 1 | 1000 | 1000 | 1000 |
| 1769 | 1 | 1000 | 1000 | 1000 |
| 1770 | 1 | 1000 | 1000 | 1000 |
| 1771 | 1 | 1000 | 1000 | 1000 |
| 1772 | 1 | 1000 | 1000 | 1000 |
| 1773 | 1 | 1000 | 1000 | 1000 |
| 1774 | 1 | 1000 | 1000 | 1000 |
| 1775 | 1 | 1000 | 1000 | 1000 |
| 1776 | 1 | 1000 | 1000 | 1000 |
| 1777 | 1 | 1000 | 1000 | 1000 |
| 1778 | 1 | 1000 | 1000 | 1000 |
| 1779 | 1 | 1000 | 1000 | 1000 |
| 1780 | 1 | 1000 | 1000 | 1000 |
| 1781 | 1 | 1000 | 1000 | 1000 |
| 1782 | 1 | 1000 | 1000 | 1000 |
| 1783 | 1 | 1000 | 1000 | 1000 |
| 1784 | 1 | 1000 | 1000 | 1000 |
| 1785 | 1 | 1000 | 1000 | 1000 |
| 1786 | 1 | 1000 | 1000 | 1000 |
| 1787 | 1 | 1000 | 1000 | 1000 |
| 1788 | 1 | 1000 | 1000 | 1000 |
| 1789 | 1 | 1000 | 1000 | 1000 |
| 1790 | 1 | 1000 | 1000 | 1000 |
| 1791 | 1 | 1000 | 1000 | 1000 |
| 1792 | 1 | 1000 | 1000 | 1000 |
| 1793 | 1 | 1000 | 1000 | 1000 |
| 1794 | 1 | 1000 | 1000 | 1000 |
| 1795 | 1 | 1000 | 1000 | 1000 |
| 1796 | 1 | 1000 | 1000 | 1000 |
| 1797 | 1 | 1000 | 1000 | 1000 |
| 1798 | 1 | 1000 | 1000 | 1000 |
| 1799 | 1 | 1000 | 1000 | 1000 |
| 1800 | 1 | 1000 | 1000 | 1000 |
| 1801 | 1 | 1000 | 1000 | 1000 |
| 1802 | 1 | 1000 | 1000 | 1000 |
| 1803 | 1 | 1000 | 1000 | 1000 |
| 1804 | 1 | 1000 | 1000 | 1000 |
| 1805 | 1 | 1000 | 1000 | 1000 |
| 1806 | 1 | 1000 | 1000 | 1000 |
| 1807 | 1 | 1000 | 1000 | 1000 |
| 1808 | 1 | 1000 | 1000 | 1000 |
| 1809 | 1 | 1000 | 1000 | 1000 |
| 1810 | 1 | 1000 | 1000 | 1000 |
| 1811 | 1 | 1000 | 1000 | 1000 |
| 1812 | 1 | 1000 | 1000 | 1000 |
| 1813 | 1 | 1000 | 1000 | 1000 |
| 1814 | 1 | 1000 | 1000 | 1000 |
| 1815 | 1 | 1000 | 1000 | 1000 |
| 1816 | 1 | 1000 | 1000 | 1000 |
| 1817 | 1 | 1000 | 1000 | 1000 |
| 1818 | 1 | 1000 | 1000 | 1000 |
| 1819 | 1 | 1000 | 1000 | 1000 |
| 1820 | 1 | 1000 | 1000 | 1000 |
| 1821 | 1 | 1000 | 1000 | 1000 |
| 1822 | 1 | 1000 | 1000 | 1000 |
| 1823 | 1 | 1000 | 1000 | 1000 |
| 1824 | 1 | 1000 | 1000 | 1000 |
| 1825 | 1 | 1000 | 1000 | 1000 |
| 1826 | 1 | 1000 | 1000 | 1000 |
| 1827 | 1 | 1000 | 1000 | 1000 |
| 1828 | 1 | 1000 | 1000 | 1000 |
| 1829 | 1 | 1000 | 1000 | 1000 |
| 1830 | 1 | 1000 | 1000 | 1000 |
| 1831 | 1 | 1000 | 1000 | 1000 |
| 1832 | 1 | 1000 | 1000 | 1000 |
| 1833 | 1 | 1000 | 1000 | 1000 |
| 1834 | 1 | 1000 | 1000 | 1000 |
| 1835 | 1 | 1000 | 1000 | 1000 |
| 1836 | 1 | 1000 | 1000 | 1000 |
| 1837 | 1 | 1000 | 1000 | 1000 |

```
1838 1 1000 1000 1000
1839 1 1000 1000 1000
1840 1 1000 1000 1000
1841 1 1000 1000 1000
1842 1 1000 1000 1000
1843 1 1000 1000 1000
1844 1 1000 1000 1000
1845 1 1000 1000 1000
1846 1 1000 1000 1000
1847 1 1000 1000 1000
1848 1 1000 1000 1000
1849 1 1000 1000 1000
1850 1 1000 1000 1000
1851 1 1000 1000 1000
1852 1 1000 1000 1000
1853 1 1000 1000 1000
1854 1 1000 1000 1000
1855 1 1000 1000 1000
1856 1 1000 1000 1000
1857 1 1000 1000 1000
1858 1 1000 1000 1000
1859 1 1000 1000 1000
1860 1 1000 1000 1000
1861 1 1000 1000 1000
1862 1 1000 1000 1000
1863 1 1000 1000 1000
1864 1 1000 1000 1000
1865 1 1000 1000 1000
1866 1 1000 1000 1000
1867 1 1000 1000 1000
1868 1 1000 1000 1000
1869 1 1000 1000 1000
1870 1 1000 1000 1000
1871 1 1000 1000 1000
1872 1 1000 1000 1000
1873 1 1000 1000 1000
1874 1 1000 1000 1000
1875 1 1000 1000 1000
1876 1 1000 1000 1000
1877 1 1000 1000 1000
1878 1 1000 1000 1000
1879 1 1000 1000 1000
1880 1 1000 1000 1000
1881 1 1000 1000 1000
1882 1 1000 1000 1000
1883 1 1000 1000 1000
1884 1 1000 1000 1000
1885 1 1000 1000 1000
1886 1 1000 1000 1000
1887 1 1000 1000 1000
1888 1 1000 1000 1000
1889 1 1000 1000 1000
1890 1 1000 1000 1000
1891 1 1000 1000 1000
1892 1 1000 1000 1000
1893 1 1000 1000 1000
1894 1 1000 1000 1000
1895 1 1000 1000 1000
1896 1 1000 1000 1000
1897 1 1000 1000 1000
1898 1 1000 1000 1000
1899 1 1000 1000 1000
1900 1 1000 1000 1000
1901 1 1000 1000 1000
1902 1 1000 1000 1000
1903 1 1000 1000 1000
1904 1 1000 1000 1000
1905 1 1000 1000 1000
1906 1 1000 1000 1000
1907 1 1000 1000 1000
1908 1 1000 1000 1000
```

```
1909 1 1000 1000 1000
1910 1 1000 1000 1000
1911 1 1000 1000 1000
1912 1 1000 1000 1000
1913 1 1000 1000 1000
1914 1 1000 1000 1000
1915 1 1000 1000 1000
1916 1 1000 1000 1000
1917 1 1000 1000 1000
1918 1 1000 1000 1000
1919 1 1000 1000 1000
1920 1 1000 1000 1000
1921 1 1000 1000 1000
1922 1 1000 1000 1000
1923 1 1000 1000 1000
1924 1 1000 1000 1000
4000 1 1000 1000 1000
4001 1 1000 1000 1000
4002 1 1000 1000 1000
4003 1 1000 1000 1000
4004 1 1000 1000 1000
4005 1 1000 1000 1000
4006 1 1000 1000 1000
4007 1 1000 1000 1000
4008 1 1000 1000 1000
4009 1 1000 1000 1000
4010 1 1000 1000 1000
4011 1 1000 1000 1000
4012 1 1000 1000 1000
4013 1 1000 1000 1000
4014 1 1000 1000 1000
4015 1 1000 1000 1000
4016 1 1000 1000 1000
4017 1 1000 1000 1000
4018 1 1000 1000 1000
4019 1 1000 1000 1000
4020 1 1000 1000 1000
4021 1 1000 1000 1000
4022 1 1000 1000 1000
4023 1 1000 1000 1000
4024 1 1000 1000 1000
4025 1 1000 1000 1000
4026 1 1000 1000 1000
4027 1 1000 1000 1000
4028 1 1000 1000 1000
4029 1 1000 1000 1000
4030 1 1000 1000 1000
4031 1 1000 1000 1000
4032 1 1000 1000 1000
4033 1 1000 1000 1000
4034 1 1000 1000 1000
4035 1 1000 1000 1000
4036 1 1000 1000 1000
4037 1 1000 1000 1000
4038 1 1000 1000 1000
4039 1 1000 1000 1000
4040 1 1000 1000 1000
4041 1 1000 1000 1000
4042 1 1000 1000 1000
4043 1 1000 1000 1000
4044 1 1000 1000 1000
4045 1 1000 1000 1000
4046 1 1000 1000 1000
4047 1 1000 1000 1000
4048 1 1000 1000 1000
4049 1 1000 1000 1000
4050 1 1000 1000 1000
4051 1 1000 1000 1000
4052 1 1000 1000 1000
4053 1 1000 1000 1000
4054 1 1000 1000 1000
```

| | | | | |
|---|---|---|---|---|
| 4055 | 1 | 1000 | 1000 | 1000 |
| 4056 | 1 | 1000 | 1000 | 1000 |
| 4057 | 1 | 1000 | 1000 | 1000 |
| 4058 | 1 | 1000 | 1000 | 1000 |
| 4059 | 1 | 1000 | 1000 | 1000 |
| 4060 | 1 | 1000 | 1000 | 1000 |
| 4061 | 1 | 1000 | 1000 | 1000 |
| 4062 | 1 | 1000 | 1000 | 1000 |
| 4063 | 1 | 1000 | 1000 | 1000 |
| 4064 | 1 | 1000 | 1000 | 1000 |
| 4065 | 1 | 1000 | 1000 | 1000 |
| 4066 | 1 | 1000 | 1000 | 1000 |
| 4067 | 1 | 1000 | 1000 | 1000 |
| 4068 | 1 | 1000 | 1000 | 1000 |
| 4069 | 1 | 1000 | 1000 | 1000 |
| 4070 | 1 | 1000 | 1000 | 1000 |
| 4071 | 1 | 1000 | 1000 | 1000 |
| 4072 | 1 | 1000 | 1000 | 1000 |
| 4073 | 1 | 1000 | 1000 | 1000 |
| 4074 | 1 | 1000 | 1000 | 1000 |
| 4075 | 1 | 1000 | 1000 | 1000 |
| 4076 | 1 | 1000 | 1000 | 1000 |
| 4077 | 1 | 1000 | 1000 | 1000 |
| 4078 | 1 | 1000 | 1000 | 1000 |
| 4079 | 1 | 1000 | 1000 | 1000 |
| 4080 | 1 | 1000 | 1000 | 1000 |
| 4081 | 1 | 1000 | 1000 | 1000 |
| 4082 | 1 | 1000 | 1000 | 1000 |
| 4083 | 1 | 1000 | 1000 | 1000 |
| 4084 | 1 | 1000 | 1000 | 1000 |
| 4085 | 1 | 1000 | 1000 | 1000 |
| 4086 | 1 | 1000 | 1000 | 1000 |
| 4087 | 1 | 1000 | 1000 | 1000 |
| 4088 | 1 | 1000 | 1000 | 1000 |
| 4089 | 1 | 1000 | 1000 | 1000 |
| 4090 | 1 | 1000 | 1000 | 1000 |
| 4091 | 1 | 1000 | 1000 | 1000 |
| 4092 | 1 | 1000 | 1000 | 1000 |
| 4093 | 1 | 1000 | 1000 | 1000 |
| 4094 | 1 | 1000 | 1000 | 1000 |
| 4095 | 1 | 1000 | 1000 | 1000 |
| 4096 | 1 | 1000 | 1000 | 1000 |
| 4097 | 1 | 1000 | 1000 | 1000 |
| 4098 | 1 | 1000 | 1000 | 1000 |
| 4099 | 1 | 1000 | 1000 | 1000 |
| 4100 | 1 | 1000 | 1000 | 1000 |
| 4101 | 1 | 1000 | 1000 | 1000 |
| 4102 | 1 | 1000 | 1000 | 1000 |
| 4103 | 1 | 1000 | 1000 | 1000 |
| 4104 | 1 | 1000 | 1000 | 1000 |
| 4105 | 1 | 1000 | 1000 | 1000 |
| 4106 | 1 | 1000 | 1000 | 1000 |
| 4107 | 1 | 1000 | 1000 | 1000 |
| 4108 | 1 | 1000 | 1000 | 1000 |
| 4109 | 1 | 1000 | 1000 | 1000 |
| 4110 | 1 | 1000 | 1000 | 1000 |
| 4111 | 1 | 1000 | 1000 | 1000 |
| 4112 | 1 | 1000 | 1000 | 1000 |
| 4113 | 1 | 1000 | 1000 | 1000 |
| 4114 | 1 | 1000 | 1000 | 1000 |
| 4115 | 1 | 1000 | 1000 | 1000 |
| 4116 | 1 | 1000 | 1000 | 1000 |
| 4117 | 1 | 1000 | 1000 | 1000 |
| 4118 | 1 | 1000 | 1000 | 1000 |

#proc.pdb

```
TITLE     Gromacs Runs On Most of All Computer Systems
MODEL        1
ATOM      1  N   SER A   1     -20.284 -17.510   3.277  1.00  0.00            N
ATOM      2  H1  SER A   1     -20.993 -17.913   2.699  1.00  0.00
ATOM      3  H2  SER A   1     -19.385 -17.815   2.963  1.00  0.00
ATOM      4  CA  SER A   1     -20.362 -16.005   3.194  1.00  0.00            C
ATOM      5  HA  SER A   1     -20.213 -15.746   2.240  1.00  0.00
ATOM      6  CB  SER A   1     -21.733 -15.502   3.574  1.00  0.00            C
ATOM      7  HB1 SER A   1     -22.418 -15.884   2.953  1.00  0.00
ATOM      8  HB2 SER A   1     -21.945 -15.774   4.513  1.00  0.00
ATOM      9  OG  SER A   1     -21.832 -14.073   3.503  1.00  0.00            O
ATOM     10  HG1 SER A   1     -22.755 -13.790   3.763  1.00  0.00
ATOM     11  C   SER A   1     -19.279 -15.369   4.077  1.00  0.00            C
ATOM     12  O   SER A   1     -19.298 -15.510   5.318  1.00  0.00            O
ATOM     13  N   MET A   2     -18.330 -14.694   3.425  1.00  0.00            N
ATOM     14  HN  MET A   2     -18.340 -14.708   2.425  1.00  0.00
ATOM     15  CA  MET A   2     -17.276 -13.935   4.099  1.00  0.00            C
ATOM     16  HA  MET A   2     -17.342 -14.150   5.073  1.00  0.00
ATOM     17  CB  MET A   2     -15.885 -14.334   3.597  1.00  0.00            C
ATOM     18  HB1 MET A   2     -15.851 -14.202   2.606  1.00  0.00
ATOM     19  HB2 MET A   2     -15.205 -13.746   4.035  1.00  0.00
ATOM     20  CG  MET A   2     -15.519 -15.765   3.888  1.00  0.00            C
ATOM     21  HG1 MET A   2     -16.395 -16.236   3.995  1.00  0.00
ATOM     22  HG2 MET A   2     -15.067 -16.087   3.056  1.00  0.00
ATOM     23  SD  MET A   2     -14.473 -15.924   5.341  1.00  0.00            S
ATOM     24  CE  MET A   2     -12.900 -16.397   4.617  1.00  0.00            C
ATOM     25  HE1 MET A   2     -12.222 -16.517   5.342  1.00  0.00
ATOM     26  HE2 MET A   2     -13.008 -17.256   4.116  1.00  0.00
ATOM     27  HE3 MET A   2     -12.593 -15.681   3.989  1.00  0.00
ATOM     28  C   MET A   2     -17.514 -12.452   3.804  1.00  0.00            C
ATOM     29  O   MET A   2     -17.953 -12.094   2.718  1.00  0.00            O
ATOM     30  N   GLU A   3     -17.255 -11.606   4.804  1.00  0.00            N
ATOM     31  HN  GLU A   3     -17.027 -11.990   5.699  1.00  0.00
ATOM     32  CA  GLU A   3     -17.285 -10.153   4.667  1.00  0.00            C
ATOM     33  HA  GLU A   3     -18.009  -9.937   4.012  1.00  0.00
ATOM     34  CB  GLU A   3     -17.616  -9.482   6.003  1.00  0.00            C
ATOM     35  HB1 GLU A   3     -17.334 -10.095   6.741  1.00  0.00
ATOM     36  HB2 GLU A   3     -17.099  -8.628   6.063  1.00  0.00
ATOM     37  CG  GLU A   3     -19.092  -9.157   6.194  1.00  0.00            C
ATOM     38  HG1 GLU A   3     -19.619  -9.980   5.979  1.00  0.00
ATOM     39  HG2 GLU A   3     -19.232  -8.913   7.154  1.00  0.00
ATOM     40  CD  GLU A   3     -19.625  -8.013   5.331  1.00  0.00            C
ATOM     41  OE1 GLU A   3     -18.821  -7.265   4.691  1.00  0.00            O
ATOM     42  OE2 GLU A   3     -20.848  -7.841   5.327  1.00  0.00            O
ATOM     43  C   GLU A   3     -15.915  -9.690   4.160  1.00  0.00            C
ATOM     44  O   GLU A   3     -14.887 -10.183   4.575  1.00  0.00            O
ATOM     45  N   VAL A   4     -15.934  -8.747   3.235  1.00  0.00            N
ATOM     46  HN  VAL A   4     -16.809  -8.321   3.004  1.00  0.00
ATOM     47  CA  VAL A   4     -14.767  -8.313   2.555  1.00  0.00            C
ATOM     48  HA  VAL A   4     -14.056  -8.984   2.764  1.00  0.00
ATOM     49  CB  VAL A   4     -15.011  -8.300   1.039  1.00  0.00            C
ATOM     50  HB  VAL A   4     -15.890  -7.856   0.868  1.00  0.00
ATOM     51  CG1 VAL A   4     -13.969  -7.489   0.296  1.00  0.00            C
ATOM     52 1HG1 VAL A   4     -14.168  -7.509  -0.684  1.00  0.00
ATOM     53 2HG1 VAL A   4     -13.989  -6.543   0.621  1.00  0.00
ATOM     54 3HG1 VAL A   4     -13.063  -7.879   0.461  1.00  0.00
ATOM     55  CG2 VAL A   4     -15.075  -9.715   0.504  1.00  0.00            C
ATOM     56 1HG2 VAL A   4     -15.234  -9.691  -0.483  1.00  0.00
ATOM     57 2HG2 VAL A   4     -14.211 -10.182   0.691  1.00  0.00
ATOM     58 3HG2 VAL A   4     -15.823 -10.207   0.950  1.00  0.00
ATOM     59  C   VAL A   4     -14.364  -6.940   3.095  1.00  0.00            C
ATOM     60  O   VAL A   4     -15.195  -6.071   3.270  1.00  0.00            O
ATOM     61  N   VAL A   5     -13.069  -6.790   3.357  1.00  0.00            N
ATOM     62  HN  VAL A   5     -12.472  -7.583   3.237  1.00  0.00
ATOM     63  CA  VAL A   5     -12.471  -5.554   3.804  1.00  0.00            C
ATOM     64  HA  VAL A   5     -13.141  -4.833   3.625  1.00  0.00
ATOM     65  CB  VAL A   5     -12.171  -5.608   5.309  1.00  0.00            C
ATOM     66  HB  VAL A   5     -11.519  -6.350   5.465  1.00  0.00
```

```
ATOM     67  CG1 VAL A   5     -11.521  -4.319   5.797  1.00  0.00           C
ATOM     68 1HG1 VAL A   5     -11.340  -4.388   6.778  1.00  0.00
ATOM     69 2HG1 VAL A   5     -10.661  -4.174   5.308  1.00  0.00
ATOM     70 3HG1 VAL A   5     -12.136  -3.550   5.625  1.00  0.00
ATOM     71  CG2 VAL A   5     -13.412  -5.936   6.112  1.00  0.00           C
ATOM     72 1HG2 VAL A   5     -13.182  -5.963   7.085  1.00  0.00
ATOM     73 2HG2 VAL A   5     -14.107  -5.235   5.953  1.00  0.00
ATOM     74 3HG2 VAL A   5     -13.767  -6.827   5.828  1.00  0.00
ATOM     75  C   VAL A   5     -11.184  -5.329   3.010  1.00  0.00           C
ATOM     76  O   VAL A   5     -10.147  -5.922   3.329  1.00  0.00           O
ATOM     77  N   GLY A   6     -11.257  -4.468   1.985  1.00  0.00           N
ATOM     78  HN  GLY A   6     -12.072  -3.896   1.891  1.00  0.00
ATOM     79  CA  GLY A   6     -10.195  -4.339   1.009  1.00  0.00           C
ATOM     80  HA1 GLY A   6     -10.542  -3.803   0.239  1.00  0.00
ATOM     81  HA2 GLY A   6      -9.434  -3.852   1.438  1.00  0.00
ATOM     82  C   GLY A   6      -9.739  -5.709   0.532  1.00  0.00           C
ATOM     83  O   GLY A   6     -10.550  -6.512   0.085  1.00  0.00           O
ATOM     84  N   ASP A   7      -8.433  -5.989   0.679  1.00  0.00           N
ATOM     85  HN  ASP A   7      -7.850  -5.324   1.145  1.00  0.00
ATOM     86  CA  ASP A   7      -7.827  -7.231   0.184  1.00  0.00           C
ATOM     87  HA  ASP A   7      -8.347  -7.450  -0.642  1.00  0.00
ATOM     88  CB  ASP A   7      -6.362  -7.026  -0.171  1.00  0.00           C
ATOM     89  HB1 ASP A   7      -5.887  -6.684   0.640  1.00  0.00
ATOM     90  HB2 ASP A   7      -5.976  -7.910  -0.436  1.00  0.00
ATOM     91  CG  ASP A   7      -6.130  -6.044  -1.309  1.00  0.00           C
ATOM     92  OD1 ASP A   7      -6.959  -5.992  -2.235  1.00  0.00           O
ATOM     93  OD2 ASP A   7      -5.103  -5.362  -1.276  1.00  0.00           O
ATOM     94  C   ASP A   7      -7.952  -8.369   1.211  1.00  0.00           C
ATOM     95  O   ASP A   7      -7.379  -9.446   1.002  1.00  0.00           O
ATOM     96  N   PHE A   8      -8.701  -8.135   2.301  1.00  0.00           N
ATOM     97  HN  PHE A   8      -9.175  -7.257   2.371  1.00  0.00
ATOM     98  CA  PHE A   8      -8.858  -9.094   3.387  1.00  0.00           C
ATOM     99  HA  PHE A   8      -8.290  -9.884   3.158  1.00  0.00
ATOM    100  CB  PHE A   8      -8.374  -8.495   4.704  1.00  0.00           C
ATOM    101  HB1 PHE A   8      -8.949  -7.708   4.928  1.00  0.00
ATOM    102  HB2 PHE A   8      -8.462  -9.186   5.421  1.00  0.00
ATOM    103  CG  PHE A   8      -6.939  -8.034   4.669  1.00  0.00           C
ATOM    104  CD1 PHE A   8      -6.614  -6.761   4.209  1.00  0.00           C
ATOM    105  HD1 PHE A   8      -7.340  -6.142   3.910  1.00  0.00
ATOM    106  CE1 PHE A   8      -5.294  -6.345   4.162  1.00  0.00           C
ATOM    107  HE1 PHE A   8      -5.074  -5.425   3.836  1.00  0.00
ATOM    108  CZ  PHE A   8      -4.277  -7.194   4.563  1.00  0.00           C
ATOM    109  HZ  PHE A   8      -3.325  -6.891   4.524  1.00  0.00
ATOM    110  CD2 PHE A   8      -5.908  -8.877   5.074  1.00  0.00           C
ATOM    111  HD2 PHE A   8      -6.122  -9.794   5.410  1.00  0.00
ATOM    112  CE2 PHE A   8      -4.583  -8.456   5.016  1.00  0.00           C
ATOM    113  HE2 PHE A   8      -3.851  -9.073   5.305  1.00  0.00
ATOM    114  C   PHE A   8     -10.324  -9.504   3.489  1.00  0.00           C
ATOM    115  O   PHE A   8     -11.188  -8.849   2.922  1.00  0.00           O
ATOM    116  N   GLU A   9     -10.585 -10.589   4.220  1.00  0.00           N
ATOM    117  HN  GLU A   9      -9.821 -11.082   4.636  1.00  0.00
ATOM    118  CA  GLU A   9     -11.932 -11.083   4.437  1.00  0.00           C
ATOM    119  HA  GLU A   9     -12.543 -10.292   4.398  1.00  0.00
ATOM    120  CB  GLU A   9     -12.322 -12.056   3.329  1.00  0.00           C
ATOM    121  HB1 GLU A   9     -13.274 -12.331   3.466  1.00  0.00
ATOM    122  HB2 GLU A   9     -12.233 -11.590   2.449  1.00  0.00
ATOM    123  CG  GLU A   9     -11.470 -13.305   3.291  1.00  0.00           C
ATOM    124  HG1 GLU A   9     -10.516 -13.017   3.214  1.00  0.00
ATOM    125  HG2 GLU A   9     -11.604 -13.788   4.157  1.00  0.00
ATOM    126  CD  GLU A   9     -11.763 -14.283   2.153  1.00  0.00           C
ATOM    127  OE1 GLU A   9     -12.826 -14.149   1.491  1.00  0.00           O
ATOM    128  OE2 GLU A   9     -10.913 -15.193   1.922  1.00  0.00           O
ATOM    129  C   GLU A   9     -12.011 -11.740   5.818  1.00  0.00           C
ATOM    130  O   GLU A   9     -10.995 -12.099   6.390  1.00  0.00           O
ATOM    131  N   TYR A  10     -13.231 -11.876   6.340  1.00  0.00           N
ATOM    132  HN  TYR A  10     -14.013 -11.515   5.832  1.00  0.00
ATOM    133  CA  TYR A  10     -13.470 -12.524   7.611  1.00  0.00           C
ATOM    134  HA  TYR A  10     -12.805 -13.270   7.641  1.00  0.00
ATOM    135  CB  TYR A  10     -13.243 -11.560   8.782  1.00  0.00           C
ATOM    136  HB1 TYR A  10     -13.213 -12.092   9.628  1.00  0.00
ATOM    137  HB2 TYR A  10     -12.366 -11.100   8.647  1.00  0.00
```

```
ATOM    138  CG  TYR A  10     -14.300 -10.494   8.947  1.00  0.00           C
ATOM    139  CD1 TYR A  10     -14.243  -9.353   8.189  1.00  0.00           C
ATOM    140  HD1 TYR A  10     -13.500  -9.245   7.529  1.00  0.00
ATOM    141  CE1 TYR A  10     -15.182  -8.348   8.311  1.00  0.00           C
ATOM    142  HE1 TYR A  10     -15.114  -7.530   7.739  1.00  0.00
ATOM    143  CZ  TYR A  10     -16.206  -8.467   9.212  1.00  0.00           C
ATOM    144  OH  TYR A  10     -17.091  -7.427   9.282  1.00  0.00           O
ATOM    145  HH  TYR A  10     -17.788  -7.628   9.970  1.00  0.00
ATOM    146  CD2 TYR A  10     -15.338 -10.613   9.856  1.00  0.00           C
ATOM    147  HD2 TYR A  10     -15.405 -11.435  10.422  1.00  0.00
ATOM    148  CE2 TYR A  10     -16.287  -9.609   9.993  1.00  0.00           C
ATOM    149  HE2 TYR A  10     -17.029  -9.710  10.655  1.00  0.00
ATOM    150  C   TYR A  10     -14.904 -13.051   7.667  1.00  0.00           C
ATOM    151  O   TYR A  10     -15.774 -12.608   6.950  1.00  0.00           O
ATOM    152  N   SER A  11     -15.102 -14.016   8.557  1.00  0.00           N
ATOM    153  HN  SER A  11     -14.296 -14.388   9.018  1.00  0.00
ATOM    154  CA  SER A  11     -16.375 -14.568   8.911  1.00  0.00           C
ATOM    155  HA  SER A  11     -17.031 -14.298   8.206  1.00  0.00
ATOM    156  CB  SER A  11     -16.273 -16.067   8.925  1.00  0.00           C
ATOM    157  HB1 SER A  11     -16.304 -16.427   7.992  1.00  0.00
ATOM    158  HB2 SER A  11     -15.422 -16.354   9.365  1.00  0.00
ATOM    159  OG  SER A  11     -17.347 -16.639   9.645  1.00  0.00           O
ATOM    160  HG1 SER A  11     -17.258 -17.635   9.641  1.00  0.00
ATOM    161  C   SER A  11     -16.790 -14.017  10.283  1.00  0.00           C
ATOM    162  O   SER A  11     -15.965 -13.919  11.191  1.00  0.00           O
ATOM    163  N   LYS A  12     -18.076 -13.681  10.422  1.00  0.00           N
ATOM    164  HN  LYS A  12     -18.689 -13.821   9.644  1.00  0.00
ATOM    165  CA  LYS A  12     -18.636 -13.118  11.654  1.00  0.00           C
ATOM    166  HA  LYS A  12     -18.038 -12.364  11.926  1.00  0.00
ATOM    167  CB  LYS A  12     -20.031 -12.566  11.385  1.00  0.00           C
ATOM    168  HB1 LYS A  12     -20.582 -13.284  10.959  1.00  0.00
ATOM    169  HB2 LYS A  12     -20.444 -12.302  12.257  1.00  0.00
ATOM    170  CG  LYS A  12     -20.025 -11.354  10.467  1.00  0.00           C
ATOM    171  HG1 LYS A  12     -19.354 -10.699  10.815  1.00  0.00
ATOM    172  HG2 LYS A  12     -19.754 -11.654   9.552  1.00  0.00
ATOM    173  CD  LYS A  12     -21.354 -10.645  10.352  1.00  0.00           C
ATOM    174  HD1 LYS A  12     -21.942 -11.168   9.735  1.00  0.00
ATOM    175  HD2 LYS A  12     -21.774 -10.605  11.259  1.00  0.00
ATOM    176  CE  LYS A  12     -21.214  -9.233   9.815  1.00  0.00           C
ATOM    177  HE1 LYS A  12     -20.979  -8.616  10.566  1.00  0.00
ATOM    178  HE2 LYS A  12     -20.489  -9.212   9.127  1.00  0.00
ATOM    179  NZ  LYS A  12     -22.472  -8.751   9.187  1.00  0.00           N
ATOM    180  HZ1 LYS A  12     -22.339  -7.820   8.847  1.00  0.00
ATOM    181  HZ2 LYS A  12     -22.716  -9.352   8.426  1.00  0.00
ATOM    182  HZ3 LYS A  12     -23.207  -8.756   9.865  1.00  0.00
ATOM    183  C   LYS A  12     -18.639 -14.164  12.773  1.00  0.00           C
ATOM    184  O   LYS A  12     -18.708 -13.800  13.937  1.00  0.00           O
ATOM    185  N   ARG A  13     -18.506 -15.447  12.408  1.00  0.00           N
ATOM    186  HN  ARG A  13     -18.493 -15.670  11.433  1.00  0.00
ATOM    187  CA  ARG A  13     -18.379 -16.530  13.374  1.00  0.00           C
ATOM    188  HA  ARG A  13     -19.189 -16.433  13.953  1.00  0.00
ATOM    189  CB  ARG A  13     -18.348 -17.903  12.685  1.00  0.00           C
ATOM    190  HB1 ARG A  13     -17.696 -17.856  11.929  1.00  0.00
ATOM    191  HB2 ARG A  13     -18.035 -18.578  13.353  1.00  0.00
ATOM    192  CG  ARG A  13     -19.672 -18.395  12.120  1.00  0.00           C
ATOM    193  HG1 ARG A  13     -20.387 -18.270  12.808  1.00  0.00
ATOM    194  HG2 ARG A  13     -19.897 -17.862  11.304  1.00  0.00
ATOM    195  CD  ARG A  13     -19.584 -19.875  11.747  1.00  0.00           C
ATOM    196  HD1 ARG A  13     -20.384 -20.126  11.201  1.00  0.00
ATOM    197  HD2 ARG A  13     -18.753 -20.032  11.213  1.00  0.00
ATOM    198  NE  ARG A  13     -19.540 -20.778  12.902  1.00  0.00           N
ATOM    199  HE  ARG A  13     -20.130 -20.557  13.679  1.00  0.00
ATOM    200  CZ  ARG A  13     -18.777 -21.874  13.005  1.00  0.00           C
ATOM    201  NH1 ARG A  13     -17.872 -22.156  12.081  1.00  0.00           N
ATOM    202 1HH1 ARG A  13     -17.753 -21.548  11.296  1.00  0.00
ATOM    203 2HH1 ARG A  13     -17.308 -22.977  12.168  1.00  0.00
ATOM    204  NH2 ARG A  13     -18.909 -22.670  14.048  1.00  0.00           N
ATOM    205 1HH2 ARG A  13     -19.578 -22.455  14.760  1.00  0.00
ATOM    206 2HH2 ARG A  13     -18.340 -23.489  14.127  1.00  0.00
ATOM    207  C   ARG A  13     -17.075 -16.384  14.166  1.00  0.00           C
ATOM    208  O   ARG A  13     -17.006 -16.795  15.319  1.00  0.00           O
```

| | | | | | | | | | | | |
|---|---|---|---|---|---|---|---|---|---|---|---|
| ATOM | 209 | N | ASP | A | 14 | -16.037 | -15.843 | 13.522 | 1.00 | 0.00 | N |
| ATOM | 210 | HN | ASP | A | 14 | -16.198 | -15.397 | 12.641 | 1.00 | 0.00 | |
| ATOM | 211 | CA | ASP | A | 14 | -14.679 | -15.876 | 14.051 | 1.00 | 0.00 | C |
| ATOM | 212 | HA | ASP | A | 14 | -14.626 | -16.683 | 14.639 | 1.00 | 0.00 | |
| ATOM | 213 | CB | ASP | A | 14 | -13.651 | -16.043 | 12.930 | 1.00 | 0.00 | C |
| ATOM | 214 | HB1 | ASP | A | 14 | -13.810 | -15.332 | 12.245 | 1.00 | 0.00 | |
| ATOM | 215 | HB2 | ASP | A | 14 | -12.736 | -15.932 | 13.319 | 1.00 | 0.00 | |
| ATOM | 216 | CG | ASP | A | 14 | -13.705 | -17.386 | 12.234 | 1.00 | 0.00 | C |
| ATOM | 217 | OD1 | ASP | A | 14 | -13.731 | -18.405 | 12.952 | 1.00 | 0.00 | O |
| ATOM | 218 | OD2 | ASP | A | 14 | -13.739 | -17.392 | 10.961 | 1.00 | 0.00 | O |
| ATOM | 219 | C | ASP | A | 14 | -14.423 | -14.611 | 14.876 | 1.00 | 0.00 | C |
| ATOM | 220 | O | ASP | A | 14 | -13.489 | -13.866 | 14.617 | 1.00 | 0.00 | O |
| ATOM | 221 | N | LEU | A | 15 | -15.262 | -14.414 | 15.897 | 1.00 | 0.00 | N |
| ATOM | 222 | HN | LEU | A | 15 | -15.983 | -15.091 | 16.046 | 1.00 | 0.00 | |
| ATOM | 223 | CA | LEU | A | 15 | -15.197 | -13.283 | 16.804 | 1.00 | 0.00 | C |
| ATOM | 224 | HA | LEU | A | 15 | -15.033 | -12.453 | 16.272 | 1.00 | 0.00 | |
| ATOM | 225 | CB | LEU | A | 15 | -16.541 | -13.202 | 17.532 | 1.00 | 0.00 | C |
| ATOM | 226 | HB1 | LEU | A | 15 | -17.261 | -13.144 | 16.841 | 1.00 | 0.00 | |
| ATOM | 227 | HB2 | LEU | A | 15 | -16.654 | -14.043 | 18.062 | 1.00 | 0.00 | |
| ATOM | 228 | CG | LEU | A | 15 | -16.722 | -12.028 | 18.486 | 1.00 | 0.00 | C |
| ATOM | 229 | HG | LEU | A | 15 | -15.982 | -12.055 | 19.158 | 1.00 | 0.00 | |
| ATOM | 230 | CD1 | LEU | A | 15 | -16.645 | -10.709 | 17.738 | 1.00 | 0.00 | C |
| ATOM | 231 | 1HD1 | LEU | A | 15 | -16.766 | -9.953 | 18.382 | 1.00 | 0.00 | |
| ATOM | 232 | 2HD1 | LEU | A | 15 | -15.752 | -10.629 | 17.294 | 1.00 | 0.00 | |
| ATOM | 233 | 3HD1 | LEU | A | 15 | -17.366 | -10.676 | 17.046 | 1.00 | 0.00 | |
| ATOM | 234 | CD2 | LEU | A | 15 | -18.059 | -12.133 | 19.209 | 1.00 | 0.00 | C |
| ATOM | 235 | 1HD2 | LEU | A | 15 | -18.164 | -11.357 | 19.831 | 1.00 | 0.00 | |
| ATOM | 236 | 2HD2 | LEU | A | 15 | -18.802 | -12.126 | 18.539 | 1.00 | 0.00 | |
| ATOM | 237 | 3HD2 | LEU | A | 15 | -18.089 | -12.985 | 19.732 | 1.00 | 0.00 | |
| ATOM | 238 | C | LEU | A | 15 | -14.035 | -13.484 | 17.785 | 1.00 | 0.00 | C |
| ATOM | 239 | O | LEU | A | 15 | -13.827 | -14.577 | 18.267 | 1.00 | 0.00 | O |
| ATOM | 240 | N | VAL | A | 16 | -13.252 | -12.427 | 18.020 | 1.00 | 0.00 | N |
| ATOM | 241 | HN | VAL | A | 16 | -13.438 | -11.573 | 17.534 | 1.00 | 0.00 | |
| ATOM | 242 | CA | VAL | A | 16 | -12.132 | -12.471 | 18.962 | 1.00 | 0.00 | C |
| ATOM | 243 | HA | VAL | A | 16 | -11.916 | -13.438 | 19.094 | 1.00 | 0.00 | |
| ATOM | 244 | CB | VAL | A | 16 | -10.873 | -11.786 | 18.400 | 1.00 | 0.00 | C |
| ATOM | 245 | HB | VAL | A | 16 | -11.060 | -10.804 | 18.359 | 1.00 | 0.00 | |
| ATOM | 246 | CG1 | VAL | A | 16 | -9.669 | -12.017 | 19.295 | 1.00 | 0.00 | C |
| ATOM | 247 | 1HG1 | VAL | A | 16 | -8.870 | -11.561 | 18.903 | 1.00 | 0.00 | |
| ATOM | 248 | 2HG1 | VAL | A | 16 | -9.853 | -11.644 | 20.204 | 1.00 | 0.00 | |
| ATOM | 249 | 3HG1 | VAL | A | 16 | -9.490 | -12.998 | 19.367 | 1.00 | 0.00 | |
| ATOM | 250 | CG2 | VAL | A | 16 | -10.556 | -12.246 | 17.000 | 1.00 | 0.00 | C |
| ATOM | 251 | 1HG2 | VAL | A | 16 | -9.734 | -11.779 | 16.674 | 1.00 | 0.00 | |
| ATOM | 252 | 2HG2 | VAL | A | 16 | -10.400 | -13.234 | 17.001 | 1.00 | 0.00 | |
| ATOM | 253 | 3HG2 | VAL | A | 16 | -11.324 | -12.031 | 16.397 | 1.00 | 0.00 | |
| ATOM | 254 | C | VAL | A | 16 | -12.578 | -11.826 | 20.278 | 1.00 | 0.00 | C |
| ATOM | 255 | O | VAL | A | 16 | -12.377 | -12.394 | 21.344 | 1.00 | 0.00 | O |
| ATOM | 256 | N | GLY | A | 17 | -13.193 | -10.643 | 20.184 | 1.00 | 0.00 | N |
| ATOM | 257 | HN | GLY | A | 17 | -13.283 | -10.220 | 19.283 | 1.00 | 0.00 | |
| ATOM | 258 | CA | GLY | A | 17 | -13.737 | -9.946 | 21.340 | 1.00 | 0.00 | C |
| ATOM | 259 | HA1 | GLY | A | 17 | -14.499 | -10.486 | 21.697 | 1.00 | 0.00 | |
| ATOM | 260 | HA2 | GLY | A | 17 | -13.018 | -9.886 | 22.032 | 1.00 | 0.00 | |
| ATOM | 261 | C | GLY | A | 17 | -14.218 | -8.553 | 20.973 | 1.00 | 0.00 | C |
| ATOM | 262 | O | GLY | A | 17 | -14.445 | -8.258 | 19.780 | 1.00 | 0.00 | O |
| ATOM | 263 | N | HIS | A | 18 | -14.385 | -7.704 | 21.991 | 1.00 | 0.00 | N |
| ATOM | 264 | HN | HIS | A | 18 | -14.113 | -7.996 | 22.908 | 1.00 | 0.00 | |
| ATOM | 265 | CA | HIS | A | 18 | -14.947 | -6.370 | 21.821 | 1.00 | 0.00 | C |
| ATOM | 266 | HA | HIS | A | 18 | -15.003 | -6.217 | 20.834 | 1.00 | 0.00 | |
| ATOM | 267 | CB | HIS | A | 18 | -16.362 | -6.289 | 22.423 | 1.00 | 0.00 | C |
| ATOM | 268 | HB1 | HIS | A | 18 | -16.291 | -6.324 | 23.420 | 1.00 | 0.00 | |
| ATOM | 269 | HB2 | HIS | A | 18 | -16.780 | -5.423 | 22.149 | 1.00 | 0.00 | |
| ATOM | 270 | ND1 | HIS | A | 18 | -17.449 | -8.548 | 22.751 | 1.00 | 0.00 | N |
| ATOM | 271 | CG | HIS | A | 18 | -17.287 | -7.381 | 22.005 | 1.00 | 0.00 | C |
| ATOM | 272 | CE1 | HIS | A | 18 | -18.330 | -9.327 | 22.155 | 1.00 | 0.00 | C |
| ATOM | 273 | HE1 | HIS | A | 18 | -18.628 | -10.227 | 22.473 | 1.00 | 0.00 | |
| ATOM | 274 | NE2 | HIS | A | 18 | -18.757 | -8.701 | 21.046 | 1.00 | 0.00 | N |
| ATOM | 275 | HE2 | HIS | A | 18 | -19.431 | -9.058 | 20.399 | 1.00 | 0.00 | |
| ATOM | 276 | CD2 | HIS | A | 18 | -18.116 | -7.489 | 20.945 | 1.00 | 0.00 | C |
| ATOM | 277 | HD2 | HIS | A | 18 | -18.241 | -6.812 | 20.220 | 1.00 | 0.00 | |
| ATOM | 278 | C | HIS | A | 18 | -14.014 | -5.337 | 22.467 | 1.00 | 0.00 | C |
| ATOM | 279 | O | HIS | A | 18 | -13.277 | -5.645 | 23.398 | 1.00 | 0.00 | O |

| | | | | | | | | | | | |
|---|---|---|---|---|---|---|---|---|---|---|---|
| ATOM | 280 | N | GLY | A | 19 | -14.052 | -4.106 | 21.942 | 1.00 | 0.00 | N |
| ATOM | 281 | HN | GLY | A | 19 | -14.322 | -4.006 | 20.984 | 1.00 | 0.00 | |
| ATOM | 282 | CA | GLY | A | 19 | -13.721 | -2.907 | 22.694 | 1.00 | 0.00 | C |
| ATOM | 283 | HA1 | GLY | A | 19 | -13.163 | -3.171 | 23.480 | 1.00 | 0.00 | |
| ATOM | 284 | HA2 | GLY | A | 19 | -13.195 | -2.297 | 22.101 | 1.00 | 0.00 | |
| ATOM | 285 | C | GLY | A | 19 | -14.992 | -2.218 | 23.160 | 1.00 | 0.00 | C |
| ATOM | 286 | O | GLY | A | 19 | -16.086 | -2.781 | 23.080 | 1.00 | 0.00 | O |
| ATOM | 287 | N | ALA | A | 20 | -14.863 | -0.974 | 23.602 | 1.00 | 0.00 | N |
| ATOM | 288 | HN | ALA | A | 20 | -13.949 | -0.581 | 23.700 | 1.00 | 0.00 | |
| ATOM | 289 | CA | ALA | A | 20 | -16.024 | -0.173 | 23.946 | 1.00 | 0.00 | C |
| ATOM | 290 | HA | ALA | A | 20 | -16.594 | -0.726 | 24.554 | 1.00 | 0.00 | |
| ATOM | 291 | CB | ALA | A | 20 | -15.602 | 1.060 | 24.712 | 1.00 | 0.00 | C |
| ATOM | 292 | HB1 | ALA | A | 20 | -16.410 | 1.602 | 24.942 | 1.00 | 0.00 | |
| ATOM | 293 | HB2 | ALA | A | 20 | -15.133 | 0.786 | 25.552 | 1.00 | 0.00 | |
| ATOM | 294 | HB3 | ALA | A | 20 | -14.983 | 1.607 | 24.148 | 1.00 | 0.00 | |
| ATOM | 295 | C | ALA | A | 20 | -16.797 | 0.181 | 22.660 | 1.00 | 0.00 | C |
| ATOM | 296 | O | ALA | A | 20 | -18.023 | 0.232 | 22.660 | 1.00 | 0.00 | O |
| ATOM | 297 | N | PHE | A | 21 | -16.060 | 0.406 | 21.562 | 1.00 | 0.00 | N |
| ATOM | 298 | HN | PHE | A | 21 | -15.082 | 0.200 | 21.593 | 1.00 | 0.00 | |
| ATOM | 299 | CA | PHE | A | 21 | -16.627 | 0.942 | 20.314 | 1.00 | 0.00 | C |
| ATOM | 300 | HA | PHE | A | 21 | -17.615 | 1.003 | 20.453 | 1.00 | 0.00 | |
| ATOM | 301 | CB | PHE | A | 21 | -16.045 | 2.327 | 20.056 | 1.00 | 0.00 | C |
| ATOM | 302 | HB1 | PHE | A | 21 | -15.047 | 2.275 | 20.102 | 1.00 | 0.00 | |
| ATOM | 303 | HB2 | PHE | A | 21 | -16.322 | 2.632 | 19.145 | 1.00 | 0.00 | |
| ATOM | 304 | CG | PHE | A | 21 | -16.514 | 3.344 | 21.060 | 1.00 | 0.00 | C |
| ATOM | 305 | CD1 | PHE | A | 21 | -17.762 | 3.924 | 20.936 | 1.00 | 0.00 | C |
| ATOM | 306 | HD1 | PHE | A | 21 | -18.350 | 3.671 | 20.168 | 1.00 | 0.00 | |
| ATOM | 307 | CE1 | PHE | A | 21 | -18.209 | 4.854 | 21.865 | 1.00 | 0.00 | C |
| ATOM | 308 | HE1 | PHE | A | 21 | -19.101 | 5.290 | 21.745 | 1.00 | 0.00 | |
| ATOM | 309 | CZ | PHE | A | 21 | -17.426 | 5.177 | 22.953 | 1.00 | 0.00 | C |
| ATOM | 310 | HZ | PHE | A | 21 | -17.739 | 5.860 | 23.613 | 1.00 | 0.00 | |
| ATOM | 311 | CD2 | PHE | A | 21 | -15.753 | 3.651 | 22.176 | 1.00 | 0.00 | C |
| ATOM | 312 | HD2 | PHE | A | 21 | -14.861 | 3.215 | 22.299 | 1.00 | 0.00 | |
| ATOM | 313 | CE2 | PHE | A | 21 | -16.210 | 4.554 | 23.125 | 1.00 | 0.00 | C |
| ATOM | 314 | HE2 | PHE | A | 21 | -15.656 | 4.752 | 23.934 | 1.00 | 0.00 | |
| ATOM | 315 | C | PHE | A | 21 | -16.370 | 0.006 | 19.126 | 1.00 | 0.00 | C |
| ATOM | 316 | O | PHE | A | 21 | -16.924 | 0.214 | 18.059 | 1.00 | 0.00 | O |
| ATOM | 317 | N | ALA | A | 22 | -15.539 | -1.023 | 19.324 | 1.00 | 0.00 | N |
| ATOM | 318 | HN | ALA | A | 22 | -15.265 | -1.227 | 20.264 | 1.00 | 0.00 | |
| ATOM | 319 | CA | ALA | A | 22 | -15.011 | -1.859 | 18.273 | 1.00 | 0.00 | C |
| ATOM | 320 | HA | ALA | A | 22 | -15.412 | -1.526 | 17.420 | 1.00 | 0.00 | |
| ATOM | 321 | CB | ALA | A | 22 | -13.510 | -1.707 | 18.221 | 1.00 | 0.00 | C |
| ATOM | 322 | HB1 | ALA | A | 22 | -13.141 | -2.285 | 17.493 | 1.00 | 0.00 | |
| ATOM | 323 | HB2 | ALA | A | 22 | -13.278 | -0.752 | 18.037 | 1.00 | 0.00 | |
| ATOM | 324 | HB3 | ALA | A | 22 | -13.116 | -1.983 | 19.098 | 1.00 | 0.00 | |
| ATOM | 325 | C | ALA | A | 22 | -15.409 | -3.325 | 18.498 | 1.00 | 0.00 | C |
| ATOM | 326 | O | ALA | A | 22 | -15.451 | -3.797 | 19.628 | 1.00 | 0.00 | O |
| ATOM | 327 | N | VAL | A | 23 | -15.639 | -4.041 | 17.392 | 1.00 | 0.00 | N |
| ATOM | 328 | HN | VAL | A | 23 | -15.757 | -3.557 | 16.525 | 1.00 | 0.00 | |
| ATOM | 329 | CA | VAL | A | 23 | -15.725 | -5.500 | 17.395 | 1.00 | 0.00 | C |
| ATOM | 330 | HA | VAL | A | 23 | -15.694 | -5.801 | 18.348 | 1.00 | 0.00 | |
| ATOM | 331 | CB | VAL | A | 23 | -17.053 | -5.984 | 16.777 | 1.00 | 0.00 | C |
| ATOM | 332 | HB | VAL | A | 23 | -16.991 | -5.851 | 15.788 | 1.00 | 0.00 | |
| ATOM | 333 | CG1 | VAL | A | 23 | -17.281 | -7.461 | 17.026 | 1.00 | 0.00 | C |
| ATOM | 334 | 1HG1 | VAL | A | 23 | -18.148 | -7.739 | 16.613 | 1.00 | 0.00 | |
| ATOM | 335 | 2HG1 | VAL | A | 23 | -16.534 | -7.987 | 16.619 | 1.00 | 0.00 | |
| ATOM | 336 | 3HG1 | VAL | A | 23 | -17.311 | -7.632 | 18.011 | 1.00 | 0.00 | |
| ATOM | 337 | CG2 | VAL | A | 23 | -18.238 | -5.175 | 17.275 | 1.00 | 0.00 | C |
| ATOM | 338 | 1HG2 | VAL | A | 23 | -19.077 | -5.518 | 16.853 | 1.00 | 0.00 | |
| ATOM | 339 | 2HG2 | VAL | A | 23 | -18.306 | -5.261 | 18.269 | 1.00 | 0.00 | |
| ATOM | 340 | 3HG2 | VAL | A | 23 | -18.111 | -4.214 | 17.031 | 1.00 | 0.00 | |
| ATOM | 341 | C | VAL | A | 23 | -14.528 | -6.032 | 16.613 | 1.00 | 0.00 | C |
| ATOM | 342 | O | VAL | A | 23 | -14.186 | -5.453 | 15.590 | 1.00 | 0.00 | O |
| ATOM | 343 | N | VAL | A | 24 | -13.920 | -7.129 | 17.078 | 1.00 | 0.00 | N |
| ATOM | 344 | HN | VAL | A | 24 | -14.316 | -7.609 | 17.861 | 1.00 | 0.00 | |
| ATOM | 345 | CA | VAL | A | 24 | -12.702 | -7.643 | 16.480 | 1.00 | 0.00 | C |
| ATOM | 346 | HA | VAL | A | 24 | -12.544 | -7.049 | 15.692 | 1.00 | 0.00 | |
| ATOM | 347 | CB | VAL | A | 24 | -11.507 | -7.553 | 17.443 | 1.00 | 0.00 | C |
| ATOM | 348 | HB | VAL | A | 24 | -11.655 | -8.218 | 18.175 | 1.00 | 0.00 | |
| ATOM | 349 | CG1 | VAL | A | 24 | -10.193 | -7.917 | 16.749 | 1.00 | 0.00 | C |
| ATOM | 350 | 1HG1 | VAL | A | 24 | -9.440 | -7.848 | 17.403 | 1.00 | 0.00 | |

```
ATOM    351 2HG1 VAL A  24     -10.248  -8.853  16.402  1.00  0.00
ATOM    352 3HG1 VAL A  24     -10.032  -7.288  15.988  1.00  0.00
ATOM    353  CG2 VAL A  24     -11.424  -6.191  18.101  1.00  0.00           C
ATOM    354 1HG2 VAL A  24     -10.638  -6.167  18.719  1.00  0.00
ATOM    355 2HG2 VAL A  24     -11.317  -5.487  17.398  1.00  0.00
ATOM    356 3HG2 VAL A  24     -12.262  -6.020  18.620  1.00  0.00
ATOM    357  C   VAL A  24     -12.903  -9.089  16.036  1.00  0.00           C
ATOM    358  O   VAL A  24     -13.262  -9.946  16.858  1.00  0.00           O
ATOM    359  N   PHE A  25     -12.549  -9.357  14.766  1.00  0.00           N
ATOM    360  HN  PHE A  25     -12.181  -8.609  14.214  1.00  0.00
ATOM    361  CA  PHE A  25     -12.668 -10.665  14.147  1.00  0.00           C
ATOM    362  HA  PHE A  25     -13.027 -11.269  14.859  1.00  0.00
ATOM    363  CB  PHE A  25     -13.622 -10.591  12.965  1.00  0.00           C
ATOM    364  HB1 PHE A  25     -13.222  -9.982  12.280  1.00  0.00
ATOM    365  HB2 PHE A  25     -13.713 -11.510  12.581  1.00  0.00
ATOM    366  CG  PHE A  25     -15.008 -10.084  13.275  1.00  0.00           C
ATOM    367  CD1 PHE A  25     -16.010 -10.958  13.686  1.00  0.00           C
ATOM    368  HD1 PHE A  25     -15.796 -11.926  13.821  1.00  0.00
ATOM    369  CE1 PHE A  25     -17.301 -10.505  13.911  1.00  0.00           C
ATOM    370  HE1 PHE A  25     -18.013 -11.147  14.197  1.00  0.00
ATOM    371  CZ  PHE A  25     -17.605  -9.170  13.741  1.00  0.00           C
ATOM    372  HZ  PHE A  25     -18.529  -8.837  13.930  1.00  0.00
ATOM    373  CD2 PHE A  25     -15.334  -8.751  13.092  1.00  0.00           C
ATOM    374  HD2 PHE A  25     -14.629  -8.105  12.799  1.00  0.00
ATOM    375  CE2 PHE A  25     -16.632  -8.301  13.311  1.00  0.00           C
ATOM    376  HE2 PHE A  25     -16.858  -7.340  13.155  1.00  0.00
ATOM    377  C   PHE A  25     -11.303 -11.156  13.660  1.00  0.00           C
ATOM    378  O   PHE A  25     -10.438 -10.367  13.353  1.00  0.00           O
ATOM    379  N   ARG A  26     -11.134 -12.480  13.642  1.00  0.00           N
ATOM    380  HN  ARG A  26     -11.828 -13.037  14.098  1.00  0.00
ATOM    381  CA  ARG A  26     -10.018 -13.184  13.011  1.00  0.00           C
ATOM    382  HA  ARG A  26      -9.162 -12.686  13.153  1.00  0.00
ATOM    383  CB  ARG A  26      -9.874 -14.570  13.658  1.00  0.00           C
ATOM    384  HB1 ARG A  26      -9.907 -14.444  14.649  1.00  0.00
ATOM    385  HB2 ARG A  26     -10.656 -15.121  13.366  1.00  0.00
ATOM    386  CG  ARG A  26      -8.616 -15.376  13.349  1.00  0.00           C
ATOM    387  HG1 ARG A  26      -8.592 -15.563  12.367  1.00  0.00
ATOM    388  HG2 ARG A  26      -7.818 -14.829  13.604  1.00  0.00
ATOM    389  CD  ARG A  26      -8.562 -16.710  14.106  1.00  0.00           C
ATOM    390  HD1 ARG A  26      -8.019 -17.358  13.572  1.00  0.00
ATOM    391  HD2 ARG A  26      -8.123 -16.557  14.991  1.00  0.00
ATOM    392  NE  ARG A  26      -9.859 -17.364  14.380  1.00  0.00           N
ATOM    393  HE  ARG A  26     -10.271 -17.886  13.633  1.00  0.00
ATOM    394  CZ  ARG A  26     -10.541 -17.321  15.552  1.00  0.00           C
ATOM    395  NH1 ARG A  26     -10.052 -16.680  16.597  1.00  0.00           N
ATOM    396 1HH1 ARG A  26      -9.168 -16.218  16.531  1.00  0.00
ATOM    397 2HH1 ARG A  26     -10.566 -16.657  17.455  1.00  0.00
ATOM    398  NH2 ARG A  26     -11.723 -17.916  15.663  1.00  0.00           N
ATOM    399 1HH2 ARG A  26     -12.115 -18.399  14.880  1.00  0.00
ATOM    400 2HH2 ARG A  26     -12.221 -17.881  16.529  1.00  0.00
ATOM    401  C   ARG A  26     -10.337 -13.253  11.519  1.00  0.00           C
ATOM    402  O   ARG A  26     -11.440 -13.573  11.153  1.00  0.00           O
ATOM    403  N   GLY A  27      -9.395 -12.837  10.684  1.00  0.00           N
ATOM    404  HN  GLY A  27      -8.519 -12.536  11.060  1.00  0.00
ATOM    405  CA  GLY A  27      -9.585 -12.802   9.250  1.00  0.00           C
ATOM    406  HA1 GLY A  27     -10.365 -13.385   9.023  1.00  0.00
ATOM    407  HA2 GLY A  27      -9.785 -11.859   8.984  1.00  0.00
ATOM    408  C   GLY A  27      -8.346 -13.292   8.535  1.00  0.00           C
ATOM    409  O   GLY A  27      -7.427 -13.802   9.156  1.00  0.00           O
ATOM    410  N   ARG A  28      -8.323 -13.125   7.211  1.00  0.00           N
ATOM    411  HN  ARG A  28      -9.128 -12.743   6.756  1.00  0.00
ATOM    412  CA  ARG A  28      -7.178 -13.477   6.421  1.00  0.00           C
ATOM    413  HA  ARG A  28      -6.383 -13.301   7.001  1.00  0.00
ATOM    414  CB  ARG A  28      -7.234 -14.950   6.041  1.00  0.00           C
ATOM    415  HB1 ARG A  28      -6.432 -15.172   5.486  1.00  0.00
ATOM    416  HB2 ARG A  28      -7.224 -15.500   6.876  1.00  0.00
ATOM    417  CG  ARG A  28      -8.481 -15.291   5.250  1.00  0.00           C
ATOM    418  HG1 ARG A  28      -9.284 -15.097   5.814  1.00  0.00
ATOM    419  HG2 ARG A  28      -8.507 -14.723   4.427  1.00  0.00
ATOM    420  CD  ARG A  28      -8.527 -16.736   4.834  1.00  0.00           C
ATOM    421  HD1 ARG A  28      -7.644 -17.014   4.456  1.00  0.00
```

| | | | | | | | | | | | |
|---|---|---|---|---|---|---|---|---|---|---|---|
| ATOM | 422 | HD2 | ARG | A | 28 | -8.750 | -17.313 | 5.619 | 1.00 | 0.00 | |
| ATOM | 423 | NE | ARG | A | 28 | -9.549 | -16.917 | 3.816 | 1.00 | 0.00 | N |
| ATOM | 424 | HE | ARG | A | 28 | -9.891 | -16.105 | 3.343 | 1.00 | 0.00 | |
| ATOM | 425 | CZ | ARG | A | 28 | -10.048 | -18.083 | 3.482 | 1.00 | 0.00 | C |
| ATOM | 426 | NH1 | ARG | A | 28 | -9.525 | -19.183 | 3.994 | 1.00 | 0.00 | N |
| ATOM | 427 | 1HH1 | ARG | A | 28 | -8.756 | -19.122 | 4.630 | 1.00 | 0.00 | |
| ATOM | 428 | 2HH1 | ARG | A | 28 | -9.898 | -20.077 | 3.746 | 1.00 | 0.00 | |
| ATOM | 429 | NH2 | ARG | A | 28 | -11.029 | -18.146 | 2.604 | 1.00 | 0.00 | N |
| ATOM | 430 | 1HH2 | ARG | A | 28 | -11.389 | -17.307 | 2.196 | 1.00 | 0.00 | |
| ATOM | 431 | 2HH2 | ARG | A | 28 | -11.413 | -19.033 | 2.346 | 1.00 | 0.00 | |
| ATOM | 432 | C | ARG | A | 28 | -7.136 | -12.613 | 5.165 | 1.00 | 0.00 | C |
| ATOM | 433 | O | ARG | A | 28 | -8.130 | -12.019 | 4.800 | 1.00 | 0.00 | O |
| ATOM | 434 | N | HIS | A | 29 | -5.955 | -12.562 | 4.543 | 1.00 | 0.00 | N |
| ATOM | 435 | HN | HIS | A | 29 | -5.148 | -12.898 | 5.029 | 1.00 | 0.00 | |
| ATOM | 436 | CA | HIS | A | 29 | -5.781 | -12.042 | 3.192 | 1.00 | 0.00 | C |
| ATOM | 437 | HA | HIS | A | 29 | -6.184 | -11.127 | 3.176 | 1.00 | 0.00 | |
| ATOM | 438 | CB | HIS | A | 29 | -4.283 | -11.930 | 2.836 | 1.00 | 0.00 | C |
| ATOM | 439 | HB1 | HIS | A | 29 | -3.791 | -11.528 | 3.608 | 1.00 | 0.00 | |
| ATOM | 440 | HB2 | HIS | A | 29 | -3.921 | -12.843 | 2.649 | 1.00 | 0.00 | |
| ATOM | 441 | ND1 | HIS | A | 29 | -3.949 | -11.610 | 0.371 | 1.00 | 0.00 | N |
| ATOM | 442 | CG | HIS | A | 29 | -4.009 | -11.085 | 1.640 | 1.00 | 0.00 | C |
| ATOM | 443 | CE1 | HIS | A | 29 | -3.704 | -10.640 | -0.493 | 1.00 | 0.00 | C |
| ATOM | 444 | HE1 | HIS | A | 29 | -3.616 | -10.746 | -1.483 | 1.00 | 0.00 | |
| ATOM | 445 | NE2 | HIS | A | 29 | -3.595 | -9.495 | 0.183 | 1.00 | 0.00 | N |
| ATOM | 446 | HE2 | HIS | A | 29 | -3.413 | -8.596 | -0.215 | 1.00 | 0.00 | |
| ATOM | 447 | CD2 | HIS | A | 29 | -3.775 | -9.759 | 1.516 | 1.00 | 0.00 | C |
| ATOM | 448 | HD2 | HIS | A | 29 | -3.740 | -9.093 | 2.261 | 1.00 | 0.00 | |
| ATOM | 449 | C | HIS | A | 29 | -6.521 | -12.959 | 2.220 | 1.00 | 0.00 | C |
| ATOM | 450 | O | HIS | A | 29 | -6.614 | -14.161 | 2.430 | 1.00 | 0.00 | O |
| ATOM | 451 | N | ARG | A | 30 | -7.041 | -12.368 | 1.151 | 1.00 | 0.00 | N |
| ATOM | 452 | HN | ARG | A | 30 | -6.828 | -11.406 | 0.978 | 1.00 | 0.00 | |
| ATOM | 453 | CA | ARG | A | 30 | -7.908 | -13.073 | 0.225 | 1.00 | 0.00 | C |
| ATOM | 454 | HA | ARG | A | 30 | -8.484 | -13.688 | 0.763 | 1.00 | 0.00 | |
| ATOM | 455 | CB | ARG | A | 30 | -8.829 | -12.077 | -0.466 | 1.00 | 0.00 | C |
| ATOM | 456 | HB1 | ARG | A | 30 | -8.271 | -11.343 | -0.852 | 1.00 | 0.00 | |
| ATOM | 457 | HB2 | ARG | A | 30 | -9.313 | -12.550 | -1.202 | 1.00 | 0.00 | |
| ATOM | 458 | CG | ARG | A | 30 | -9.858 | -11.463 | 0.482 | 1.00 | 0.00 | C |
| ATOM | 459 | HG1 | ARG | A | 30 | -10.511 | -12.166 | 0.763 | 1.00 | 0.00 | |
| ATOM | 460 | HG2 | ARG | A | 30 | -9.391 | -11.100 | 1.288 | 1.00 | 0.00 | |
| ATOM | 461 | CD | ARG | A | 30 | -10.606 | -10.336 | -0.211 | 1.00 | 0.00 | C |
| ATOM | 462 | HD1 | ARG | A | 30 | -11.170 | -9.832 | 0.443 | 1.00 | 0.00 | |
| ATOM | 463 | HD2 | ARG | A | 30 | -9.966 | -9.707 | -0.653 | 1.00 | 0.00 | |
| ATOM | 464 | NE | ARG | A | 30 | -11.470 | -10.931 | -1.227 | 1.00 | 0.00 | N |
| ATOM | 465 | HE | ARG | A | 30 | -12.112 | -11.638 | -0.931 | 1.00 | 0.00 | |
| ATOM | 466 | CZ | ARG | A | 30 | -11.468 | -10.605 | -2.513 | 1.00 | 0.00 | C |
| ATOM | 467 | NH1 | ARG | A | 30 | -10.989 | -9.424 | -2.885 | 1.00 | 0.00 | N |
| ATOM | 468 | 1HH1 | ARG | A | 30 | -10.634 | -8.790 | -2.198 | 1.00 | 0.00 | |
| ATOM | 469 | 2HH1 | ARG | A | 30 | -10.983 | -9.169 | -3.852 | 1.00 | 0.00 | |
| ATOM | 470 | NH2 | ARG | A | 30 | -11.922 | -11.475 | -3.404 | 1.00 | 0.00 | N |
| ATOM | 471 | 1HH2 | ARG | A | 30 | -12.261 | -12.367 | -3.104 | 1.00 | 0.00 | |
| ATOM | 472 | 2HH2 | ARG | A | 30 | -11.925 | -11.240 | -4.376 | 1.00 | 0.00 | |
| ATOM | 473 | C | ARG | A | 30 | -7.089 | -13.957 | -0.724 | 1.00 | 0.00 | C |
| ATOM | 474 | O | ARG | A | 30 | -7.539 | -15.003 | -1.063 | 1.00 | 0.00 | O |
| ATOM | 475 | N | GLN | A | 31 | -5.879 | -13.558 | -1.084 | 1.00 | 0.00 | N |
| ATOM | 476 | HN | GLN | A | 31 | -5.563 | -12.652 | -0.803 | 1.00 | 0.00 | |
| ATOM | 477 | CA | GLN | A | 31 | -4.974 | -14.419 | -1.895 | 1.00 | 0.00 | C |
| ATOM | 478 | HA | GLN | A | 31 | -5.572 | -15.022 | -2.424 | 1.00 | 0.00 | |
| ATOM | 479 | CB | GLN | A | 31 | -4.131 | -13.587 | -2.856 | 1.00 | 0.00 | C |
| ATOM | 480 | HB1 | GLN | A | 31 | -3.573 | -12.951 | -2.323 | 1.00 | 0.00 | |
| ATOM | 481 | HB2 | GLN | A | 31 | -3.534 | -14.202 | -3.371 | 1.00 | 0.00 | |
| ATOM | 482 | CG | GLN | A | 31 | -4.939 | -12.785 | -3.843 | 1.00 | 0.00 | C |
| ATOM | 483 | HG1 | GLN | A | 31 | -5.435 | -13.411 | -4.445 | 1.00 | 0.00 | |
| ATOM | 484 | HG2 | GLN | A | 31 | -5.592 | -12.214 | -3.344 | 1.00 | 0.00 | |
| ATOM | 485 | CD | GLN | A | 31 | -4.042 | -11.901 | -4.677 | 1.00 | 0.00 | C |
| ATOM | 486 | OE1 | GLN | A | 31 | -3.409 | -12.347 | -5.649 | 1.00 | 0.00 | O |
| ATOM | 487 | NE2 | GLN | A | 31 | -3.984 | -10.625 | -4.294 | 1.00 | 0.00 | N |
| ATOM | 488 | 1HE2 | GLN | A | 31 | -4.513 | -10.316 | -3.504 | 1.00 | 0.00 | |
| ATOM | 489 | 2HE2 | GLN | A | 31 | -3.411 | -9.978 | -4.797 | 1.00 | 0.00 | |
| ATOM | 490 | C | GLN | A | 31 | -4.049 | -15.242 | -0.982 | 1.00 | 0.00 | C |
| ATOM | 491 | O | GLN | A | 31 | -3.988 | -16.446 | -1.087 | 1.00 | 0.00 | O |
| ATOM | 492 | N | LYS | A | 32 | -3.333 | -14.558 | -0.095 | 1.00 | 0.00 | N |

```
ATOM    493  HN  LYS A  32      -3.444 -13.565  -0.063  1.00  0.00
ATOM    494  CA  LYS A  32      -2.399 -15.173   0.829  1.00  0.00           C
ATOM    495  HA  LYS A  32      -1.996 -15.977   0.391  1.00  0.00
ATOM    496  CB  LYS A  32      -1.292 -14.160   1.151  1.00  0.00           C
ATOM    497  HB1 LYS A  32      -0.977 -13.760   0.290  1.00  0.00
ATOM    498  HB2 LYS A  32      -1.684 -13.444   1.728  1.00  0.00
ATOM    499  CG  LYS A  32      -0.077 -14.707   1.869  1.00  0.00           C
ATOM    500  HG1 LYS A  32      -0.354 -15.012   2.780  1.00  0.00
ATOM    501  HG2 LYS A  32       0.278 -15.485   1.350  1.00  0.00
ATOM    502  CD  LYS A  32       1.026 -13.700   2.024  1.00  0.00           C
ATOM    503  HD1 LYS A  32       1.421 -13.516   1.124  1.00  0.00
ATOM    504  HD2 LYS A  32       0.639 -12.856   2.397  1.00  0.00
ATOM    505  CE  LYS A  32       2.117 -14.191   2.948  1.00  0.00           C
ATOM    506  HE1 LYS A  32       2.852 -13.514   2.985  1.00  0.00
ATOM    507  HE2 LYS A  32       1.742 -14.325   3.865  1.00  0.00
ATOM    508  NZ  LYS A  32       2.680 -15.477   2.474  1.00  0.00           N
ATOM    509  HZ1 LYS A  32       3.398 -15.778   3.102  1.00  0.00
ATOM    510  HZ2 LYS A  32       1.956 -16.166   2.440  1.00  0.00
ATOM    511  HZ3 LYS A  32       3.067 -15.355   1.560  1.00  0.00
ATOM    512  C   LYS A  32      -3.172 -15.605   2.083  1.00  0.00           C
ATOM    513  O   LYS A  32      -3.078 -14.985   3.127  1.00  0.00           O
ATOM    514  N   THR A  33      -3.926 -16.693   1.965  1.00  0.00           N
ATOM    515  HN  THR A  33      -3.797 -17.284   1.169  1.00  0.00
ATOM    516  CA  THR A  33      -4.942 -17.059   2.960  1.00  0.00           C
ATOM    517  HA  THR A  33      -5.420 -16.207   3.175  1.00  0.00
ATOM    518  CB  THR A  33      -5.939 -18.068   2.379  1.00  0.00           C
ATOM    519  HB  THR A  33      -6.571 -18.384   3.086  1.00  0.00
ATOM    520  OG1 THR A  33      -5.191 -19.178   1.898  1.00  0.00           O
ATOM    521  HG1 THR A  33      -5.814 -19.857   1.511  1.00  0.00
ATOM    522  CG2 THR A  33      -6.753 -17.502   1.238  1.00  0.00           C
ATOM    523 1HG2 THR A  33      -7.383 -18.201   0.900  1.00  0.00
ATOM    524 2HG2 THR A  33      -7.273 -16.711   1.560  1.00  0.00
ATOM    525 3HG2 THR A  33      -6.141 -17.220   0.500  1.00  0.00
ATOM    526  C   THR A  33      -4.294 -17.605   4.236  1.00  0.00           C
ATOM    527  O   THR A  33      -4.983 -17.749   5.232  1.00  0.00           O
ATOM    528  N   ASP A  34      -2.984 -17.886   4.196  1.00  0.00           N
ATOM    529  HN  ASP A  34      -2.495 -17.786   3.329  1.00  0.00
ATOM    530  CA  ASP A  34      -2.237 -18.338   5.387  1.00  0.00           C
ATOM    531  HA  ASP A  34      -2.834 -18.988   5.858  1.00  0.00
ATOM    532  CB  ASP A  34      -0.956 -19.096   5.003  1.00  0.00           C
ATOM    533  HB1 ASP A  34      -0.402 -19.211   5.827  1.00  0.00
ATOM    534  HB2 ASP A  34      -1.218 -19.994   4.650  1.00  0.00
ATOM    535  CG  ASP A  34      -0.092 -18.407   3.946  1.00  0.00           C
ATOM    536  OD1 ASP A  34      -0.630 -18.149   2.834  1.00  0.00           O
ATOM    537  OD2 ASP A  34       1.115 -18.143   4.228  1.00  0.00           O
ATOM    538  C   ASP A  34      -1.951 -17.139   6.315  1.00  0.00           C
ATOM    539  O   ASP A  34      -1.646 -17.315   7.487  1.00  0.00           O
ATOM    540  N   TRP A  35      -2.062 -15.924   5.774  1.00  0.00           N
ATOM    541  HN  TRP A  35      -2.341 -15.860   4.816  1.00  0.00
ATOM    542  CA  TRP A  35      -1.801 -14.685   6.494  1.00  0.00           C
ATOM    543  HA  TRP A  35      -1.038 -14.860   7.116  1.00  0.00
ATOM    544  CB  TRP A  35      -1.408 -13.596   5.496  1.00  0.00           C
ATOM    545  HB1 TRP A  35      -0.548 -13.862   5.060  1.00  0.00
ATOM    546  HB2 TRP A  35      -2.126 -13.528   4.803  1.00  0.00
ATOM    547  CG  TRP A  35      -1.213 -12.231   6.071  1.00  0.00           C
ATOM    548  CD1 TRP A  35      -1.077 -11.901   7.382  1.00  0.00           C
ATOM    549  HD1 TRP A  35      -1.097 -12.550   8.142  1.00  0.00
ATOM    550  NE1 TRP A  35      -0.910 -10.555   7.508  1.00  0.00           N
ATOM    551  HE1 TRP A  35      -0.808 -10.063   8.373  1.00  0.00
ATOM    552  CE2 TRP A  35      -0.903  -9.982   6.267  1.00  0.00           C
ATOM    553  CD2 TRP A  35      -1.058 -11.011   5.326  1.00  0.00           C
ATOM    554  CE3 TRP A  35      -1.045 -10.684   3.961  1.00  0.00           C
ATOM    555  HE3 TRP A  35      -1.151 -11.397   3.267  1.00  0.00
ATOM    556  CZ3 TRP A  35      -0.884  -9.367   3.591  1.00  0.00           C
ATOM    557  HZ3 TRP A  35      -0.888  -9.125   2.621  1.00  0.00
ATOM    558  CZ2 TRP A  35      -0.674  -8.664   5.881  1.00  0.00           C
ATOM    559  HZ2 TRP A  35      -0.485  -7.954   6.559  1.00  0.00
ATOM    560  CH2 TRP A  35      -0.716  -8.369   4.545  1.00  0.00           C
ATOM    561  HH2 TRP A  35      -0.624  -7.418   4.249  1.00  0.00
ATOM    562  C   TRP A  35      -3.034 -14.282   7.315  1.00  0.00           C
ATOM    563  O   TRP A  35      -4.009 -13.761   6.771  1.00  0.00           O
```

```
ATOM    564  N   GLU A  36      -2.975 -14.529   8.631  1.00  0.00           N
ATOM    565  HN  GLU A  36      -2.142 -14.930   9.012  1.00  0.00
ATOM    566  CA  GLU A  36      -4.078 -14.234   9.516  1.00  0.00           C
ATOM    567  HA  GLU A  36      -4.913 -14.383   8.986  1.00  0.00
ATOM    568  CB  GLU A  36      -4.083 -15.178  10.711  1.00  0.00           C
ATOM    569  HB1 GLU A  36      -3.152 -15.247  11.069  1.00  0.00
ATOM    570  HB2 GLU A  36      -4.684 -14.800  11.415  1.00  0.00
ATOM    571  CG  GLU A  36      -4.570 -16.573  10.357  1.00  0.00           C
ATOM    572  HG1 GLU A  36      -5.406 -16.487   9.816  1.00  0.00
ATOM    573  HG2 GLU A  36      -3.864 -17.024   9.811  1.00  0.00
ATOM    574  CD  GLU A  36      -4.870 -17.445  11.561  1.00  0.00           C
ATOM    575  OE1 GLU A  36      -4.070 -17.394  12.507  1.00  0.00           O
ATOM    576  OE2 GLU A  36      -5.919 -18.141  11.565  1.00  0.00           O
ATOM    577  C   GLU A  36      -3.969 -12.779   9.972  1.00  0.00           C
ATOM    578  O   GLU A  36      -2.860 -12.257  10.169  1.00  0.00           O
ATOM    579  N   VAL A  37      -5.127 -12.134  10.125  1.00  0.00           N
ATOM    580  HN  VAL A  37      -5.978 -12.616   9.919  1.00  0.00
ATOM    581  CA  VAL A  37      -5.193 -10.789  10.570  1.00  0.00           C
ATOM    582  HA  VAL A  37      -4.311 -10.638  11.017  1.00  0.00
ATOM    583  CB  VAL A  37      -5.358  -9.815   9.394  1.00  0.00           C
ATOM    584  HB  VAL A  37      -5.405  -8.893   9.777  1.00  0.00
ATOM    585  CG1 VAL A  37      -4.173  -9.866   8.444  1.00  0.00           C
ATOM    586 1HG1 VAL A  37      -4.318  -9.220   7.695  1.00  0.00
ATOM    587 2HG1 VAL A  37      -3.339  -9.621   8.938  1.00  0.00
ATOM    588 3HG1 VAL A  37      -4.082 -10.791   8.074  1.00  0.00
ATOM    589  CG2 VAL A  37      -6.643 -10.054   8.643  1.00  0.00           C
ATOM    590 1HG2 VAL A  37      -6.714  -9.403   7.888  1.00  0.00
ATOM    591 2HG2 VAL A  37      -6.649 -10.986   8.281  1.00  0.00
ATOM    592 3HG2 VAL A  37      -7.419  -9.932   9.262  1.00  0.00
ATOM    593  C   VAL A  37      -6.345 -10.632  11.556  1.00  0.00           C
ATOM    594  O   VAL A  37      -7.295 -11.399  11.547  1.00  0.00           O
ATOM    595  N   ALA A  38      -6.230  -9.613  12.404  1.00  0.00           N
ATOM    596  HN  ALA A  38      -5.342  -9.159  12.480  1.00  0.00
ATOM    597  CA  ALA A  38      -7.320  -9.126  13.223  1.00  0.00           C
ATOM    598  HA  ALA A  38      -7.967  -9.877  13.353  1.00  0.00
ATOM    599  CB  ALA A  38      -6.804  -8.701  14.564  1.00  0.00           C
ATOM    600  HB1 ALA A  38      -7.563  -8.366  15.122  1.00  0.00
ATOM    601  HB2 ALA A  38      -6.374  -9.481  15.018  1.00  0.00
ATOM    602  HB3 ALA A  38      -6.130  -7.972  14.446  1.00  0.00
ATOM    603  C   ALA A  38      -7.968  -7.952  12.508  1.00  0.00           C
ATOM    604  O   ALA A  38      -7.276  -7.049  12.072  1.00  0.00           O
ATOM    605  N   ILE A  39      -9.292  -7.976  12.408  1.00  0.00           N
ATOM    606  HN  ILE A  39      -9.793  -8.758  12.779  1.00  0.00
ATOM    607  CA  ILE A  39     -10.024  -6.911  11.782  1.00  0.00           C
ATOM    608  HA  ILE A  39      -9.367  -6.220  11.481  1.00  0.00
ATOM    609  CB  ILE A  39     -10.748  -7.469  10.549  1.00  0.00           C
ATOM    610  HB  ILE A  39     -11.339  -8.222  10.838  1.00  0.00
ATOM    611  CG2 ILE A  39     -11.661  -6.432   9.913  1.00  0.00           C
ATOM    612 1HG2 ILE A  39     -12.114  -6.831   9.116  1.00  0.00
ATOM    613 2HG2 ILE A  39     -12.350  -6.142  10.577  1.00  0.00
ATOM    614 3HG2 ILE A  39     -11.119  -5.641   9.628  1.00  0.00
ATOM    615  CG1 ILE A  39      -9.735  -8.014   9.549  1.00  0.00           C
ATOM    616 1HG1 ILE A  39      -9.151  -7.266   9.235  1.00  0.00
ATOM    617 2HG1 ILE A  39      -9.174  -8.709   9.998  1.00  0.00
ATOM    618  CD  ILE A  39     -10.376  -8.635   8.355  1.00  0.00           C
ATOM    619  HD1 ILE A  39      -9.668  -8.974   7.735  1.00  0.00
ATOM    620  HD2 ILE A  39     -10.956  -9.396   8.646  1.00  0.00
ATOM    621  HD3 ILE A  39     -10.933  -7.952   7.883  1.00  0.00
ATOM    622  C   ILE A  39     -10.962  -6.266  12.815  1.00  0.00           C
ATOM    623  O   ILE A  39     -11.895  -6.919  13.300  1.00  0.00           O
ATOM    624  N   LYS A  40     -10.676  -5.005  13.175  1.00  0.00           N
ATOM    625  HN  LYS A  40      -9.849  -4.579  12.810  1.00  0.00
ATOM    626  CA  LYS A  40     -11.526  -4.226  14.082  1.00  0.00           C
ATOM    627  HA  LYS A  40     -11.925  -4.910  14.693  1.00  0.00
ATOM    628  CB  LYS A  40     -10.737  -3.182  14.868  1.00  0.00           C
ATOM    629  HB1 LYS A  40     -10.120  -2.722  14.230  1.00  0.00
ATOM    630  HB2 LYS A  40     -11.390  -2.520  15.236  1.00  0.00
ATOM    631  CG  LYS A  40      -9.899  -3.681  16.026  1.00  0.00           C
ATOM    632  HG1 LYS A  40     -10.464  -4.270  16.604  1.00  0.00
ATOM    633  HG2 LYS A  40      -9.130  -4.208  15.665  1.00  0.00
ATOM    634  CD  LYS A  40      -9.363  -2.531  16.864  1.00  0.00           C
```

| | | | | | | | | | | | |
|---|---|---|---|---|---|---|---|---|---|---|---|
| ATOM | 635 | HD1 | LYS | A | 40 | -8.806 | -1.942 | 16.278 | 1.00 | 0.00 | |
| ATOM | 636 | HD2 | LYS | A | 40 | -10.137 | -2.009 | 17.222 | 1.00 | 0.00 | |
| ATOM | 637 | CE | LYS | A | 40 | -8.512 | -2.989 | 18.031 | 1.00 | 0.00 | C |
| ATOM | 638 | HE1 | LYS | A | 40 | -9.100 | -3.381 | 18.739 | 1.00 | 0.00 | |
| ATOM | 639 | HE2 | LYS | A | 40 | -7.861 | -3.680 | 17.718 | 1.00 | 0.00 | |
| ATOM | 640 | NZ | LYS | A | 40 | -7.754 | -1.862 | 18.621 | 1.00 | 0.00 | N |
| ATOM | 641 | HZ1 | LYS | A | 40 | -7.204 | -2.195 | 19.387 | 1.00 | 0.00 | |
| ATOM | 642 | HZ2 | LYS | A | 40 | -7.154 | -1.464 | 17.927 | 1.00 | 0.00 | |
| ATOM | 643 | HZ3 | LYS | A | 40 | -8.393 | -1.165 | 18.948 | 1.00 | 0.00 | |
| ATOM | 644 | C | LYS | A | 40 | -12.557 | -3.448 | 13.271 | 1.00 | 0.00 | C |
| ATOM | 645 | O | LYS | A | 40 | -12.190 | -2.681 | 12.406 | 1.00 | 0.00 | O |
| ATOM | 646 | N | SER | A | 41 | -13.838 | -3.656 | 13.584 | 1.00 | 0.00 | N |
| ATOM | 647 | HN | SER | A | 41 | -14.059 | -4.406 | 14.208 | 1.00 | 0.00 | |
| ATOM | 648 | CA | SER | A | 41 | -14.923 | -2.850 | 13.065 | 1.00 | 0.00 | C |
| ATOM | 649 | HA | SER | A | 41 | -14.588 | -2.404 | 12.235 | 1.00 | 0.00 | |
| ATOM | 650 | CB | SER | A | 41 | -16.075 | -3.711 | 12.687 | 1.00 | 0.00 | C |
| ATOM | 651 | HB1 | SER | A | 41 | -15.834 | -4.292 | 11.910 | 1.00 | 0.00 | |
| ATOM | 652 | HB2 | SER | A | 41 | -16.348 | -4.284 | 13.460 | 1.00 | 0.00 | |
| ATOM | 653 | OG | SER | A | 41 | -17.182 | -2.911 | 12.316 | 1.00 | 0.00 | O |
| ATOM | 654 | HG1 | SER | A | 41 | -17.950 | -3.500 | 12.064 | 1.00 | 0.00 | |
| ATOM | 655 | C | SER | A | 41 | -15.335 | -1.821 | 14.113 | 1.00 | 0.00 | C |
| ATOM | 656 | O | SER | A | 41 | -15.595 | -2.188 | 15.252 | 1.00 | 0.00 | O |
| ATOM | 657 | N | ILE | A | 42 | -15.385 | -0.552 | 13.707 | 1.00 | 0.00 | N |
| ATOM | 658 | HN | ILE | A | 42 | -15.227 | -0.363 | 12.738 | 1.00 | 0.00 | |
| ATOM | 659 | CA | ILE | A | 42 | -15.654 | 0.571 | 14.584 | 1.00 | 0.00 | C |
| ATOM | 660 | HA | ILE | A | 42 | -15.933 | 0.184 | 15.463 | 1.00 | 0.00 | |
| ATOM | 661 | CB | ILE | A | 42 | -14.388 | 1.420 | 14.773 | 1.00 | 0.00 | C |
| ATOM | 662 | HB | ILE | A | 42 | -14.124 | 1.762 | 13.871 | 1.00 | 0.00 | |
| ATOM | 663 | CG2 | ILE | A | 42 | -14.670 | 2.626 | 15.675 | 1.00 | 0.00 | C |
| ATOM | 664 | 1HG2 | ILE | A | 42 | -13.834 | 3.163 | 15.784 | 1.00 | 0.00 | |
| ATOM | 665 | 2HG2 | ILE | A | 42 | -15.380 | 3.195 | 15.259 | 1.00 | 0.00 | |
| ATOM | 666 | 3HG2 | ILE | A | 42 | -14.980 | 2.307 | 16.571 | 1.00 | 0.00 | |
| ATOM | 667 | CG1 | ILE | A | 42 | -13.196 | 0.602 | 15.277 | 1.00 | 0.00 | C |
| ATOM | 668 | 1HG1 | ILE | A | 42 | -13.370 | 0.340 | 16.226 | 1.00 | 0.00 | |
| ATOM | 669 | 2HG1 | ILE | A | 42 | -13.115 | -0.220 | 14.714 | 1.00 | 0.00 | |
| ATOM | 670 | CD | ILE | A | 42 | -11.868 | 1.364 | 15.219 | 1.00 | 0.00 | C |
| ATOM | 671 | HD1 | ILE | A | 42 | -11.133 | 0.778 | 15.560 | 1.00 | 0.00 | |
| ATOM | 672 | HD2 | ILE | A | 42 | -11.675 | 1.625 | 14.273 | 1.00 | 0.00 | |
| ATOM | 673 | HD3 | ILE | A | 42 | -11.930 | 2.185 | 15.786 | 1.00 | 0.00 | |
| ATOM | 674 | C | ILE | A | 42 | -16.775 | 1.419 | 13.967 | 1.00 | 0.00 | C |
| ATOM | 675 | O | ILE | A | 42 | -16.597 | 2.022 | 12.915 | 1.00 | 0.00 | O |
| ATOM | 676 | N | ASN | A | 43 | -17.927 | 1.493 | 14.636 | 1.00 | 0.00 | N |
| ATOM | 677 | HN | ASN | A | 43 | -18.034 | 0.963 | 15.477 | 1.00 | 0.00 | |
| ATOM | 678 | CA | ASN | A | 43 | -19.037 | 2.325 | 14.176 | 1.00 | 0.00 | C |
| ATOM | 679 | HA | ASN | A | 43 | -19.220 | 2.067 | 13.227 | 1.00 | 0.00 | |
| ATOM | 680 | CB | ASN | A | 43 | -20.294 | 2.040 | 14.995 | 1.00 | 0.00 | C |
| ATOM | 681 | HB1 | ASN | A | 43 | -20.567 | 1.089 | 14.848 | 1.00 | 0.00 | |
| ATOM | 682 | HB2 | ASN | A | 43 | -20.088 | 2.182 | 15.963 | 1.00 | 0.00 | |
| ATOM | 683 | CG | ASN | A | 43 | -21.454 | 2.932 | 14.619 | 1.00 | 0.00 | C |
| ATOM | 684 | OD1 | ASN | A | 43 | -21.708 | 3.932 | 15.280 | 1.00 | 0.00 | O |
| ATOM | 685 | ND2 | ASN | A | 43 | -22.152 | 2.581 | 13.559 | 1.00 | 0.00 | N |
| ATOM | 686 | 1HD2 | ASN | A | 43 | -21.904 | 1.759 | 13.047 | 1.00 | 0.00 | |
| ATOM | 687 | 2HD2 | ASN | A | 43 | -22.930 | 3.137 | 13.266 | 1.00 | 0.00 | |
| ATOM | 688 | C | ASN | A | 43 | -18.630 | 3.809 | 14.246 | 1.00 | 0.00 | C |
| ATOM | 689 | O | ASN | A | 43 | -18.177 | 4.275 | 15.278 | 1.00 | 0.00 | O |
| ATOM | 690 | N | LYS | A | 44 | -18.828 | 4.547 | 13.148 | 1.00 | 0.00 | N |
| ATOM | 691 | HN | LYS | A | 44 | -19.317 | 4.136 | 12.379 | 1.00 | 0.00 | |
| ATOM | 692 | CA | LYS | A | 44 | -18.356 | 5.947 | 13.018 | 1.00 | 0.00 | C |
| ATOM | 693 | HA | LYS | A | 44 | -17.412 | 5.934 | 13.347 | 1.00 | 0.00 | |
| ATOM | 694 | CB | LYS | A | 44 | -18.388 | 6.415 | 11.558 | 1.00 | 0.00 | C |
| ATOM | 695 | HB1 | LYS | A | 44 | -19.224 | 6.067 | 11.133 | 1.00 | 0.00 | |
| ATOM | 696 | HB2 | LYS | A | 44 | -18.400 | 7.415 | 11.547 | 1.00 | 0.00 | |
| ATOM | 697 | CG | LYS | A | 44 | -17.209 | 5.951 | 10.720 | 1.00 | 0.00 | C |
| ATOM | 698 | HG1 | LYS | A | 44 | -16.401 | 6.467 | 11.004 | 1.00 | 0.00 | |
| ATOM | 699 | HG2 | LYS | A | 44 | -17.062 | 4.978 | 10.898 | 1.00 | 0.00 | |
| ATOM | 700 | CD | LYS | A | 44 | -17.378 | 6.132 | 9.230 | 1.00 | 0.00 | C |
| ATOM | 701 | HD1 | LYS | A | 44 | -16.991 | 5.333 | 8.769 | 1.00 | 0.00 | |
| ATOM | 702 | HD2 | LYS | A | 44 | -18.356 | 6.192 | 9.028 | 1.00 | 0.00 | |
| ATOM | 703 | CE | LYS | A | 44 | -16.695 | 7.378 | 8.695 | 1.00 | 0.00 | C |
| ATOM | 704 | HE1 | LYS | A | 44 | -17.121 | 8.189 | 9.096 | 1.00 | 0.00 | |
| ATOM | 705 | HE2 | LYS | A | 44 | -15.725 | 7.356 | 8.939 | 1.00 | 0.00 | |

| | | | | | | | | | | | |
|---|---|---|---|---|---|---|---|---|---|---|---|
| ATOM | 706 | NZ | LYS | A | 44 | -16.807 | 7.469 | 7.217 | 1.00 | 0.00 | N |
| ATOM | 707 | HZ1 | LYS | A | 44 | -16.348 | 8.298 | 6.898 | 1.00 | 0.00 | |
| ATOM | 708 | HZ2 | LYS | A | 44 | -16.376 | 6.669 | 6.800 | 1.00 | 0.00 | |
| ATOM | 709 | HZ3 | LYS | A | 44 | -17.772 | 7.501 | 6.957 | 1.00 | 0.00 | |
| ATOM | 710 | C | LYS | A | 44 | -19.193 | 6.897 | 13.881 | 1.00 | 0.00 | C |
| ATOM | 711 | O | LYS | A | 44 | -18.660 | 7.866 | 14.425 | 1.00 | 0.00 | O |
| ATOM | 712 | N | LYS | A | 45 | -20.498 | 6.625 | 14.005 | 1.00 | 0.00 | N |
| ATOM | 713 | HN | LYS | A | 45 | -20.868 | 5.805 | 13.569 | 1.00 | 0.00 | |
| ATOM | 714 | CA | LYS | A | 45 | -21.400 | 7.501 | 14.765 | 1.00 | 0.00 | C |
| ATOM | 715 | HA | LYS | A | 45 | -21.174 | 8.436 | 14.491 | 1.00 | 0.00 | |
| ATOM | 716 | CB | LYS | A | 45 | -22.865 | 7.245 | 14.397 | 1.00 | 0.00 | C |
| ATOM | 717 | HB1 | LYS | A | 45 | -22.954 | 6.295 | 14.097 | 1.00 | 0.00 | |
| ATOM | 718 | HB2 | LYS | A | 45 | -23.426 | 7.392 | 15.212 | 1.00 | 0.00 | |
| ATOM | 719 | CG | LYS | A | 45 | -23.398 | 8.141 | 13.290 | 1.00 | 0.00 | C |
| ATOM | 720 | HG1 | LYS | A | 45 | -23.705 | 8.999 | 13.703 | 1.00 | 0.00 | |
| ATOM | 721 | HG2 | LYS | A | 45 | -22.651 | 8.327 | 12.652 | 1.00 | 0.00 | |
| ATOM | 722 | CD | LYS | A | 45 | -24.557 | 7.554 | 12.503 | 1.00 | 0.00 | C |
| ATOM | 723 | HD1 | LYS | A | 45 | -24.414 | 6.569 | 12.407 | 1.00 | 0.00 | |
| ATOM | 724 | HD2 | LYS | A | 45 | -25.405 | 7.721 | 13.006 | 1.00 | 0.00 | |
| ATOM | 725 | CE | LYS | A | 45 | -24.687 | 8.161 | 11.121 | 1.00 | 0.00 | C |
| ATOM | 726 | HE1 | LYS | A | 45 | -25.258 | 8.981 | 11.169 | 1.00 | 0.00 | |
| ATOM | 727 | HE2 | LYS | A | 45 | -23.780 | 8.406 | 10.778 | 1.00 | 0.00 | |
| ATOM | 728 | NZ | LYS | A | 45 | -25.308 | 7.211 | 10.170 | 1.00 | 0.00 | N |
| ATOM | 729 | HZ1 | LYS | A | 45 | -25.380 | 7.640 | 9.269 | 1.00 | 0.00 | |
| ATOM | 730 | HZ2 | LYS | A | 45 | -24.743 | 6.389 | 10.104 | 1.00 | 0.00 | |
| ATOM | 731 | HZ3 | LYS | A | 45 | -26.221 | 6.963 | 10.495 | 1.00 | 0.00 | |
| ATOM | 732 | C | LYS | A | 45 | -21.151 | 7.335 | 16.273 | 1.00 | 0.00 | C |
| ATOM | 733 | O | LYS | A | 45 | -21.156 | 8.310 | 17.003 | 1.00 | 0.00 | O |
| ATOM | 734 | N | ASN | A | 46 | -20.940 | 6.100 | 16.738 | 1.00 | 0.00 | N |
| ATOM | 735 | HN | ASN | A | 46 | -20.994 | 5.325 | 16.109 | 1.00 | 0.00 | |
| ATOM | 736 | CA | ASN | A | 46 | -20.634 | 5.859 | 18.137 | 1.00 | 0.00 | C |
| ATOM | 737 | HA | ASN | A | 46 | -21.423 | 6.223 | 18.632 | 1.00 | 0.00 | |
| ATOM | 738 | CB | ASN | A | 46 | -20.491 | 4.377 | 18.474 | 1.00 | 0.00 | C |
| ATOM | 739 | HB1 | ASN | A | 46 | -19.900 | 3.943 | 17.794 | 1.00 | 0.00 | |
| ATOM | 740 | HB2 | ASN | A | 46 | -20.078 | 4.286 | 19.380 | 1.00 | 0.00 | |
| ATOM | 741 | CG | ASN | A | 46 | -21.806 | 3.658 | 18.482 | 1.00 | 0.00 | C |
| ATOM | 742 | OD1 | ASN | A | 46 | -22.839 | 4.278 | 18.680 | 1.00 | 0.00 | O |
| ATOM | 743 | ND2 | ASN | A | 46 | -21.770 | 2.357 | 18.246 | 1.00 | 0.00 | N |
| ATOM | 744 | 1HD2 | ASN | A | 46 | -20.895 | 1.904 | 18.074 | 1.00 | 0.00 | |
| ATOM | 745 | 2HD2 | ASN | A | 46 | -22.618 | 1.826 | 18.239 | 1.00 | 0.00 | |
| ATOM | 746 | C | ASN | A | 46 | -19.339 | 6.578 | 18.489 | 1.00 | 0.00 | C |
| ATOM | 747 | O | ASN | A | 46 | -19.231 | 7.201 | 19.550 | 1.00 | 0.00 | O |
| ATOM | 748 | N | LEU | A | 47 | -18.365 | 6.474 | 17.584 | 1.00 | 0.00 | N |
| ATOM | 749 | HN | LEU | A | 47 | -18.561 | 6.009 | 16.720 | 1.00 | 0.00 | |
| ATOM | 750 | CA | LEU | A | 47 | -17.043 | 7.003 | 17.799 | 1.00 | 0.00 | C |
| ATOM | 751 | HA | LEU | A | 47 | -16.719 | 6.627 | 18.667 | 1.00 | 0.00 | |
| ATOM | 752 | CB | LEU | A | 47 | -16.153 | 6.550 | 16.650 | 1.00 | 0.00 | C |
| ATOM | 753 | HB1 | LEU | A | 47 | -16.147 | 5.550 | 16.637 | 1.00 | 0.00 | |
| ATOM | 754 | HB2 | LEU | A | 47 | -16.549 | 6.892 | 15.798 | 1.00 | 0.00 | |
| ATOM | 755 | CG | LEU | A | 47 | -14.716 | 7.020 | 16.714 | 1.00 | 0.00 | C |
| ATOM | 756 | HG | LEU | A | 47 | -14.688 | 8.016 | 16.793 | 1.00 | 0.00 | |
| ATOM | 757 | CD1 | LEU | A | 47 | -14.026 | 6.448 | 17.935 | 1.00 | 0.00 | C |
| ATOM | 758 | 1HD1 | LEU | A | 47 | -13.079 | 6.767 | 17.963 | 1.00 | 0.00 | |
| ATOM | 759 | 2HD1 | LEU | A | 47 | -14.504 | 6.750 | 18.760 | 1.00 | 0.00 | |
| ATOM | 760 | 3HD1 | LEU | A | 47 | -14.040 | 5.449 | 17.888 | 1.00 | 0.00 | |
| ATOM | 761 | CD2 | LEU | A | 47 | -14.010 | 6.618 | 15.442 | 1.00 | 0.00 | C |
| ATOM | 762 | 1HD2 | LEU | A | 47 | -13.059 | 6.925 | 15.477 | 1.00 | 0.00 | |
| ATOM | 763 | 2HD2 | LEU | A | 47 | -14.035 | 5.623 | 15.347 | 1.00 | 0.00 | |
| ATOM | 764 | 3HD2 | LEU | A | 47 | -14.468 | 7.039 | 14.659 | 1.00 | 0.00 | |
| ATOM | 765 | C | LEU | A | 47 | -17.085 | 8.534 | 17.895 | 1.00 | 0.00 | C |
| ATOM | 766 | O | LEU | A | 47 | -16.308 | 9.124 | 18.644 | 1.00 | 0.00 | O |
| ATOM | 767 | N | SER | A | 48 | -17.986 | 9.160 | 17.131 | 1.00 | 0.00 | N |
| ATOM | 768 | HN | SER | A | 48 | -18.602 | 8.610 | 16.566 | 1.00 | 0.00 | |
| ATOM | 769 | CA | SER | A | 48 | -18.107 | 10.611 | 17.088 | 1.00 | 0.00 | C |
| ATOM | 770 | HA | SER | A | 48 | -17.167 | 10.949 | 17.043 | 1.00 | 0.00 | |
| ATOM | 771 | CB | SER | A | 48 | -18.808 | 11.068 | 15.846 | 1.00 | 0.00 | C |
| ATOM | 772 | HB1 | SER | A | 48 | -18.481 | 11.972 | 15.571 | 1.00 | 0.00 | |
| ATOM | 773 | HB2 | SER | A | 48 | -18.660 | 10.419 | 15.100 | 1.00 | 0.00 | |
| ATOM | 774 | OG | SER | A | 48 | -20.194 | 11.158 | 16.076 | 1.00 | 0.00 | O |
| ATOM | 775 | HG1 | SER | A | 48 | -20.651 | 11.464 | 15.241 | 1.00 | 0.00 | |
| ATOM | 776 | C | SER | A | 48 | -18.813 | 11.143 | 18.348 | 1.00 | 0.00 | C |

```
ATOM    777  O   SER A  48     -18.821  12.330  18.570  1.00  0.00           O
ATOM    778  N   LYS A  49     -19.393  10.254  19.159  1.00  0.00           N
ATOM    779  HN  LYS A  49     -19.409   9.293  18.882  1.00  0.00
ATOM    780  CA  LYS A  49     -20.017  10.630  20.458  1.00  0.00           C
ATOM    781  HA  LYS A  49     -20.206  11.610  20.390  1.00  0.00
ATOM    782  CB  LYS A  49     -21.325   9.864  20.700  1.00  0.00           C
ATOM    783  HB1 LYS A  49     -21.130   8.886  20.623  1.00  0.00
ATOM    784  HB2 LYS A  49     -21.637  10.069  21.628  1.00  0.00
ATOM    785  CG  LYS A  49     -22.469  10.181  19.750  1.00  0.00           C
ATOM    786  HG1 LYS A  49     -22.727  11.139  19.872  1.00  0.00
ATOM    787  HG2 LYS A  49     -22.153  10.038  18.812  1.00  0.00
ATOM    788  CD  LYS A  49     -23.697   9.322  19.977  1.00  0.00           C
ATOM    789  HD1 LYS A  49     -23.414   8.364  20.030  1.00  0.00
ATOM    790  HD2 LYS A  49     -24.123   9.592  20.841  1.00  0.00
ATOM    791  CE  LYS A  49     -24.715   9.471  18.862  1.00  0.00           C
ATOM    792  HE1 LYS A  49     -24.964  10.435  18.767  1.00  0.00
ATOM    793  HE2 LYS A  49     -24.318   9.143  18.005  1.00  0.00
ATOM    794  NZ  LYS A  49     -25.948   8.696  19.132  1.00  0.00           N
ATOM    795  HZ1 LYS A  49     -26.592   8.820  18.377  1.00  0.00
ATOM    796  HZ2 LYS A  49     -25.720   7.726  19.220  1.00  0.00
ATOM    797  HZ3 LYS A  49     -26.365   9.019  19.982  1.00  0.00
ATOM    798  C   LYS A  49     -19.056  10.337  21.614  1.00  0.00           C
ATOM    799  O   LYS A  49     -19.407  10.553  22.757  1.00  0.00           O
ATOM    800  N   SER A  50     -17.842   9.872  21.294  1.00  0.00           N
ATOM    801  HN  SER A  50     -17.624   9.742  20.327  1.00  0.00
ATOM    802  CA  SER A  50     -16.818   9.545  22.285  1.00  0.00           C
ATOM    803  HA  SER A  50     -17.268   9.550  23.178  1.00  0.00
ATOM    804  CB  SER A  50     -16.266   8.170  22.041  1.00  0.00           C
ATOM    805  HB1 SER A  50     -15.813   7.830  22.865  1.00  0.00
ATOM    806  HB2 SER A  50     -17.002   7.545  21.783  1.00  0.00
ATOM    807  OG  SER A  50     -15.305   8.186  20.981  1.00  0.00           O
ATOM    808  HG1 SER A  50     -14.952   7.261  20.837  1.00  0.00
ATOM    809  C   SER A  50     -15.697  10.586  22.233  1.00  0.00           C
ATOM    810  O   SER A  50     -15.669  11.429  21.349  1.00  0.00           O
ATOM    811  N   GLN A  51     -14.725  10.441  23.121  1.00  0.00           N
ATOM    812  HN  GLN A  51     -14.806   9.731  23.821  1.00  0.00
ATOM    813  CA  GLN A  51     -13.573  11.266  23.105  1.00  0.00           C
ATOM    814  HA  GLN A  51     -13.781  12.036  22.502  1.00  0.00
ATOM    815  CB  GLN A  51     -13.320  11.769  24.518  1.00  0.00           C
ATOM    816  HB1 GLN A  51     -13.129  10.985  25.109  1.00  0.00
ATOM    817  HB2 GLN A  51     -12.527  12.378  24.505  1.00  0.00
ATOM    818  CG  GLN A  51     -14.510  12.529  25.083  1.00  0.00           C
ATOM    819  HG1 GLN A  51     -14.741  13.284  24.469  1.00  0.00
ATOM    820  HG2 GLN A  51     -15.291  11.909  25.158  1.00  0.00
ATOM    821  CD  GLN A  51     -14.195  13.088  26.450  1.00  0.00           C
ATOM    822  OE1 GLN A  51     -13.404  12.517  27.212  1.00  0.00           O
ATOM    823  NE2 GLN A  51     -14.796  14.234  26.759  1.00  0.00           N
ATOM    824 1HE2 GLN A  51     -15.411  14.669  26.102  1.00  0.00
ATOM    825 2HE2 GLN A  51     -14.632  14.660  27.649  1.00  0.00
ATOM    826  C   GLN A  51     -12.380  10.498  22.529  1.00  0.00           C
ATOM    827  O   GLN A  51     -11.247  10.969  22.658  1.00  0.00           O
ATOM    828  N   ILE A  52     -12.652   9.399  21.812  1.00  0.00           N
ATOM    829  HN  ILE A  52     -13.607   9.134  21.676  1.00  0.00
ATOM    830  CA  ILE A  52     -11.607   8.580  21.226  1.00  0.00           C
ATOM    831  HA  ILE A  52     -10.903   8.507  21.933  1.00  0.00
ATOM    832  CB  ILE A  52     -12.107   7.157  20.886  1.00  0.00           C
ATOM    833  HB  ILE A  52     -12.858   7.251  20.233  1.00  0.00
ATOM    834  CG2 ILE A  52     -11.001   6.341  20.230  1.00  0.00           C
ATOM    835 1HG2 ILE A  52     -11.343   5.426  20.018  1.00  0.00
ATOM    836 2HG2 ILE A  52     -10.710   6.792  19.386  1.00  0.00
ATOM    837 3HG2 ILE A  52     -10.224   6.270  20.855  1.00  0.00
ATOM    838  CG1 ILE A  52     -12.692   6.422  22.090  1.00  0.00           C
ATOM    839 1HG1 ILE A  52     -13.504   6.915  22.402  1.00  0.00
ATOM    840 2HG1 ILE A  52     -12.951   5.499  21.807  1.00  0.00
ATOM    841  CD  ILE A  52     -11.750   6.301  23.253  1.00  0.00           C
ATOM    842  HD1 ILE A  52     -12.202   5.811  23.998  1.00  0.00
ATOM    843  HD2 ILE A  52     -10.934   5.798  22.970  1.00  0.00
ATOM    844  HD3 ILE A  52     -11.487   7.214  23.565  1.00  0.00
ATOM    845  C   ILE A  52     -11.065   9.288  19.973  1.00  0.00           C
ATOM    846  O   ILE A  52     -11.844   9.715  19.099  1.00  0.00           O
ATOM    847  N   LEU A  53      -9.726   9.353  19.881  1.00  0.00           N
```

| | | | | | | | | | | | |
|---|---|---|---|---|---|---|---|---|---|---|---|
| ATOM | 848 | HN | LEU | A | 53 | -9.191 | 9.008 | 20.652 | 1.00 | 0.00 | |
| ATOM | 849 | CA | LEU | A | 53 | -9.013 | 9.877 | 18.765 | 1.00 | 0.00 | C |
| ATOM | 850 | HA | LEU | A | 53 | -9.676 | 10.365 | 18.197 | 1.00 | 0.00 | |
| ATOM | 851 | CB | LEU | A | 53 | -7.937 | 10.827 | 19.291 | 1.00 | 0.00 | C |
| ATOM | 852 | HB1 | LEU | A | 53 | -7.242 | 10.282 | 19.760 | 1.00 | 0.00 | |
| ATOM | 853 | HB2 | LEU | A | 53 | -7.523 | 11.293 | 18.509 | 1.00 | 0.00 | |
| ATOM | 854 | CG | LEU | A | 53 | -8.438 | 11.889 | 20.262 | 1.00 | 0.00 | C |
| ATOM | 855 | HG | LEU | A | 53 | -8.825 | 11.425 | 21.059 | 1.00 | 0.00 | |
| ATOM | 856 | CD1 | LEU | A | 53 | -7.295 | 12.770 | 20.723 | 1.00 | 0.00 | C |
| ATOM | 857 | 1HD1 | LEU | A | 53 | -7.641 | 13.460 | 21.359 | 1.00 | 0.00 | |
| ATOM | 858 | 2HD1 | LEU | A | 53 | -6.606 | 12.209 | 21.183 | 1.00 | 0.00 | |
| ATOM | 859 | 3HD1 | LEU | A | 53 | -6.883 | 13.222 | 19.932 | 1.00 | 0.00 | |
| ATOM | 860 | CD2 | LEU | A | 53 | -9.525 | 12.738 | 19.623 | 1.00 | 0.00 | C |
| ATOM | 861 | 1HD2 | LEU | A | 53 | -9.838 | 13.427 | 20.277 | 1.00 | 0.00 | |
| ATOM | 862 | 2HD2 | LEU | A | 53 | -9.159 | 13.192 | 18.811 | 1.00 | 0.00 | |
| ATOM | 863 | 3HD2 | LEU | A | 53 | -10.293 | 12.154 | 19.361 | 1.00 | 0.00 | |
| ATOM | 864 | C | LEU | A | 53 | -8.396 | 8.730 | 17.963 | 1.00 | 0.00 | C |
| ATOM | 865 | O | LEU | A | 53 | -7.221 | 8.430 | 18.088 | 1.00 | 0.00 | O |
| ATOM | 866 | N | LEU | A | 54 | -9.192 | 8.156 | 17.067 | 1.00 | 0.00 | N |
| ATOM | 867 | HN | LEU | A | 54 | -10.123 | 8.504 | 16.958 | 1.00 | 0.00 | |
| ATOM | 868 | CA | LEU | A | 54 | -8.766 | 7.052 | 16.247 | 1.00 | 0.00 | C |
| ATOM | 869 | HA | LEU | A | 54 | -8.397 | 6.362 | 16.870 | 1.00 | 0.00 | |
| ATOM | 870 | CB | LEU | A | 54 | -9.976 | 6.495 | 15.481 | 1.00 | 0.00 | C |
| ATOM | 871 | HB1 | LEU | A | 54 | -10.678 | 6.240 | 16.146 | 1.00 | 0.00 | |
| ATOM | 872 | HB2 | LEU | A | 54 | -10.328 | 7.215 | 14.883 | 1.00 | 0.00 | |
| ATOM | 873 | CG | LEU | A | 54 | -9.673 | 5.270 | 14.619 | 1.00 | 0.00 | C |
| ATOM | 874 | HG | LEU | A | 54 | -8.890 | 5.476 | 14.032 | 1.00 | 0.00 | |
| ATOM | 875 | CD1 | LEU | A | 54 | -9.301 | 4.094 | 15.504 | 1.00 | 0.00 | C |
| ATOM | 876 | 1HD1 | LEU | A | 54 | -9.104 | 3.297 | 14.934 | 1.00 | 0.00 | |
| ATOM | 877 | 2HD1 | LEU | A | 54 | -8.492 | 4.326 | 16.044 | 1.00 | 0.00 | |
| ATOM | 878 | 3HD1 | LEU | A | 54 | -10.062 | 3.885 | 16.118 | 1.00 | 0.00 | |
| ATOM | 879 | CD2 | LEU | A | 54 | -10.839 | 4.932 | 13.695 | 1.00 | 0.00 | C |
| ATOM | 880 | 1HD2 | LEU | A | 54 | -10.608 | 4.128 | 13.147 | 1.00 | 0.00 | |
| ATOM | 881 | 2HD2 | LEU | A | 54 | -11.653 | 4.739 | 14.243 | 1.00 | 0.00 | |
| ATOM | 882 | 3HD2 | LEU | A | 54 | -11.020 | 5.707 | 13.090 | 1.00 | 0.00 | |
| ATOM | 883 | C | LEU | A | 54 | -7.673 | 7.519 | 15.278 | 1.00 | 0.00 | C |
| ATOM | 884 | O | LEU | A | 54 | -6.742 | 6.769 | 14.971 | 1.00 | 0.00 | O |
| ATOM | 885 | N | GLY | A | 55 | -7.828 | 8.742 | 14.764 | 1.00 | 0.00 | N |
| ATOM | 886 | HN | GLY | A | 55 | -8.669 | 9.243 | 14.970 | 1.00 | 0.00 | |
| ATOM | 887 | CA | GLY | A | 55 | -6.840 | 9.372 | 13.927 | 1.00 | 0.00 | C |
| ATOM | 888 | HA1 | GLY | A | 55 | -6.840 | 8.898 | 13.047 | 1.00 | 0.00 | |
| ATOM | 889 | HA2 | GLY | A | 55 | -7.117 | 10.324 | 13.793 | 1.00 | 0.00 | |
| ATOM | 890 | C | GLY | A | 55 | -5.454 | 9.316 | 14.548 | 1.00 | 0.00 | C |
| ATOM | 891 | O | GLY | A | 55 | -4.475 | 9.040 | 13.872 | 1.00 | 0.00 | O |
| ATOM | 892 | N | LYS | A | 56 | -5.384 | 9.558 | 15.856 | 1.00 | 0.00 | N |
| ATOM | 893 | HN | LYS | A | 56 | -6.225 | 9.772 | 16.352 | 1.00 | 0.00 | |
| ATOM | 894 | CA | LYS | A | 56 | -4.134 | 9.523 | 16.585 | 1.00 | 0.00 | C |
| ATOM | 895 | HA | LYS | A | 56 | -3.514 | 10.103 | 16.057 | 1.00 | 0.00 | |
| ATOM | 896 | CB | LYS | A | 56 | -4.321 | 10.089 | 17.987 | 1.00 | 0.00 | C |
| ATOM | 897 | HB1 | LYS | A | 56 | -4.707 | 11.008 | 17.907 | 1.00 | 0.00 | |
| ATOM | 898 | HB2 | LYS | A | 56 | -4.961 | 9.501 | 18.481 | 1.00 | 0.00 | |
| ATOM | 899 | CG | LYS | A | 56 | -3.033 | 10.179 | 18.794 | 1.00 | 0.00 | C |
| ATOM | 900 | HG1 | LYS | A | 56 | -2.498 | 9.347 | 18.647 | 1.00 | 0.00 | |
| ATOM | 901 | HG2 | LYS | A | 56 | -2.509 | 10.972 | 18.483 | 1.00 | 0.00 | |
| ATOM | 902 | CD | LYS | A | 56 | -3.318 | 10.327 | 20.280 | 1.00 | 0.00 | C |
| ATOM | 903 | HD1 | LYS | A | 56 | -2.460 | 10.537 | 20.749 | 1.00 | 0.00 | |
| ATOM | 904 | HD2 | LYS | A | 56 | -3.960 | 11.083 | 20.407 | 1.00 | 0.00 | |
| ATOM | 905 | CE | LYS | A | 56 | -3.915 | 9.078 | 20.904 | 1.00 | 0.00 | C |
| ATOM | 906 | HE1 | LYS | A | 56 | -4.757 | 8.834 | 20.422 | 1.00 | 0.00 | |
| ATOM | 907 | HE2 | LYS | A | 56 | -3.261 | 8.325 | 20.835 | 1.00 | 0.00 | |
| ATOM | 908 | NZ | LYS | A | 56 | -4.239 | 9.283 | 22.331 | 1.00 | 0.00 | N |
| ATOM | 909 | HZ1 | LYS | A | 56 | -4.629 | 8.443 | 22.708 | 1.00 | 0.00 | |
| ATOM | 910 | HZ2 | LYS | A | 56 | -3.406 | 9.518 | 22.832 | 1.00 | 0.00 | |
| ATOM | 911 | HZ3 | LYS | A | 56 | -4.902 | 10.027 | 22.419 | 1.00 | 0.00 | |
| ATOM | 912 | C | LYS | A | 56 | -3.583 | 8.094 | 16.666 | 1.00 | 0.00 | C |
| ATOM | 913 | O | LYS | A | 56 | -2.380 | 7.888 | 16.434 | 1.00 | 0.00 | O |
| ATOM | 914 | N | GLU | A | 57 | -4.442 | 7.127 | 17.030 | 1.00 | 0.00 | N |
| ATOM | 915 | HN | GLU | A | 57 | -5.391 | 7.368 | 17.231 | 1.00 | 0.00 | |
| ATOM | 916 | CA | GLU | A | 57 | -4.022 | 5.730 | 17.139 | 1.00 | 0.00 | C |
| ATOM | 917 | HA | GLU | A | 57 | -3.325 | 5.702 | 17.855 | 1.00 | 0.00 | |
| ATOM | 918 | CB | GLU | A | 57 | -5.192 | 4.832 | 17.561 | 1.00 | 0.00 | C |

```
ATOM    919  HB1 GLU A  57      -5.950   4.977  16.925  1.00  0.00
ATOM    920  HB2 GLU A  57      -4.895   3.878  17.513  1.00  0.00
ATOM    921  CG  GLU A  57      -5.697   5.100  18.970  1.00  0.00           C
ATOM    922  HG1 GLU A  57      -5.044   4.704  19.615  1.00  0.00
ATOM    923  HG2 GLU A  57      -5.738   6.090  19.103  1.00  0.00
ATOM    924  CD  GLU A  57      -7.071   4.525  19.275  1.00  0.00           C
ATOM    925  OE1 GLU A  57      -7.334   3.403  18.822  1.00  0.00           O
ATOM    926  OE2 GLU A  57      -7.902   5.231  19.911  1.00  0.00           O
ATOM    927  C   GLU A  57      -3.437   5.272  15.793  1.00  0.00           C
ATOM    928  O   GLU A  57      -2.404   4.607  15.756  1.00  0.00           O
ATOM    929  N   ILE A  58      -4.103   5.659  14.697  1.00  0.00           N
ATOM    930  HN  ILE A  58      -4.909   6.241  14.805  1.00  0.00
ATOM    931  CA  ILE A  58      -3.699   5.265  13.356  1.00  0.00           C
ATOM    932  HA  ILE A  58      -3.597   4.271  13.400  1.00  0.00
ATOM    933  CB  ILE A  58      -4.773   5.602  12.311  1.00  0.00           C
ATOM    934  HB  ILE A  58      -5.022   6.566  12.402  1.00  0.00
ATOM    935  CG2 ILE A  58      -4.208   5.379  10.910  1.00  0.00           C
ATOM    936 1HG2 ILE A  58      -4.907   5.598  10.230  1.00  0.00
ATOM    937 2HG2 ILE A  58      -3.413   5.970  10.773  1.00  0.00
ATOM    938 3HG2 ILE A  58      -3.934   4.423  10.810  1.00  0.00
ATOM    939  CG1 ILE A  58      -6.058   4.791  12.537  1.00  0.00           C
ATOM    940 1HG1 ILE A  58      -5.875   3.832  12.323  1.00  0.00
ATOM    941 2HG1 ILE A  58      -6.326   4.873  13.497  1.00  0.00
ATOM    942  CD  ILE A  58      -7.222   5.230  11.701  1.00  0.00           C
ATOM    943  HD1 ILE A  58      -8.015   4.656  11.906  1.00  0.00
ATOM    944  HD2 ILE A  58      -7.439   6.184  11.907  1.00  0.00
ATOM    945  HD3 ILE A  58      -6.989   5.143  10.733  1.00  0.00
ATOM    946  C   ILE A  58      -2.362   5.913  12.999  1.00  0.00           C
ATOM    947  O   ILE A  58      -1.497   5.266  12.422  1.00  0.00           O
ATOM    948  N   LYS A  59      -2.217   7.194  13.320  1.00  0.00           N
ATOM    949  HN  LYS A  59      -2.964   7.662  13.792  1.00  0.00
ATOM    950  CA  LYS A  59      -1.003   7.940  13.007  1.00  0.00           C
ATOM    951  HA  LYS A  59      -0.859   7.873  12.020  1.00  0.00
ATOM    952  CB  LYS A  59      -1.178   9.412  13.373  1.00  0.00           C
ATOM    953  HB1 LYS A  59      -1.857   9.804  12.753  1.00  0.00
ATOM    954  HB2 LYS A  59      -1.519   9.457  14.312  1.00  0.00
ATOM    955  CG  LYS A  59       0.077  10.271  13.292  1.00  0.00           C
ATOM    956  HG1 LYS A  59       0.770   9.891  13.905  1.00  0.00
ATOM    957  HG2 LYS A  59       0.419  10.257  12.352  1.00  0.00
ATOM    958  CD  LYS A  59      -0.206  11.715  13.697  1.00  0.00           C
ATOM    959  HD1 LYS A  59      -0.795  12.133  13.005  1.00  0.00
ATOM    960  HD2 LYS A  59      -0.676  11.715  14.580  1.00  0.00
ATOM    961  CE  LYS A  59       1.042  12.556  13.826  1.00  0.00           C
ATOM    962  HE1 LYS A  59       0.886  13.285  14.493  1.00  0.00
ATOM    963  HE2 LYS A  59       1.801  11.982  14.131  1.00  0.00
ATOM    964  NZ  LYS A  59       1.409  13.172  12.535  1.00  0.00           N
ATOM    965  HZ1 LYS A  59       2.237  13.721  12.651  1.00  0.00
ATOM    966  HZ2 LYS A  59       1.577  12.454  11.859  1.00  0.00
ATOM    967  HZ3 LYS A  59       0.662  13.757  12.221  1.00  0.00
ATOM    968  C   LYS A  59       0.180   7.310  13.743  1.00  0.00           C
ATOM    969  O   LYS A  59       1.254   7.170  13.191  1.00  0.00           O
ATOM    970  N   ILE A  60      -0.044   6.910  14.993  1.00  0.00           N
ATOM    971  HN  ILE A  60      -0.966   6.991  15.372  1.00  0.00
ATOM    972  CA  ILE A  60       1.016   6.358  15.823  1.00  0.00           C
ATOM    973  HA  ILE A  60       1.818   6.942  15.699  1.00  0.00
ATOM    974  CB  ILE A  60       0.604   6.417  17.296  1.00  0.00           C
ATOM    975  HB  ILE A  60      -0.350   6.129  17.385  1.00  0.00
ATOM    976  CG2 ILE A  60       1.417   5.446  18.116  1.00  0.00           C
ATOM    977 1HG2 ILE A  60       1.134   5.499  19.074  1.00  0.00
ATOM    978 2HG2 ILE A  60       1.269   4.517  17.776  1.00  0.00
ATOM    979 3HG2 ILE A  60       2.387   5.678  18.042  1.00  0.00
ATOM    980  CG1 ILE A  60       0.710   7.849  17.825  1.00  0.00           C
ATOM    981 1HG1 ILE A  60       1.677   8.078  17.933  1.00  0.00
ATOM    982 2HG1 ILE A  60       0.294   8.467  17.157  1.00  0.00
ATOM    983  CD  ILE A  60       0.026   8.069  19.155  1.00  0.00           C
ATOM    984  HD1 ILE A  60       0.138   9.023  19.434  1.00  0.00
ATOM    985  HD2 ILE A  60      -0.948   7.859  19.068  1.00  0.00
ATOM    986  HD3 ILE A  60       0.435   7.470  19.843  1.00  0.00
ATOM    987  C   ILE A  60       1.372   4.934  15.353  1.00  0.00           C
ATOM    988  O   ILE A  60       2.534   4.576  15.291  1.00  0.00           O
ATOM    989  N   LEU A  61       0.364   4.132  15.018  1.00  0.00           N
```

| | | | | | | | | | | | |
|---|---|---|---|---|---|---|---|---|---|---|---|
| ATOM | 990 | HN | LEU | A | 61 | -0.574 | 4.471 | 15.087 | 1.00 | 0.00 | |
| ATOM | 991 | CA | LEU | A | 61 | 0.597 | 2.766 | 14.551 | 1.00 | 0.00 | C |
| ATOM | 992 | HA | LEU | A | 61 | 1.156 | 2.310 | 15.244 | 1.00 | 0.00 | |
| ATOM | 993 | CB | LEU | A | 61 | -0.747 | 2.048 | 14.421 | 1.00 | 0.00 | C |
| ATOM | 994 | HB1 | LEU | A | 61 | -1.396 | 2.681 | 13.999 | 1.00 | 0.00 | |
| ATOM | 995 | HB2 | LEU | A | 61 | -0.618 | 1.256 | 13.824 | 1.00 | 0.00 | |
| ATOM | 996 | CG | LEU | A | 61 | -1.356 | 1.554 | 15.732 | 1.00 | 0.00 | C |
| ATOM | 997 | HG | LEU | A | 61 | -1.397 | 2.338 | 16.352 | 1.00 | 0.00 | |
| ATOM | 998 | CD1 | LEU | A | 61 | -2.770 | 1.054 | 15.521 | 1.00 | 0.00 | C |
| ATOM | 999 | 1HD1 | LEU | A | 61 | -3.145 | 0.737 | 16.392 | 1.00 | 0.00 | |
| ATOM | 1000 | 2HD1 | LEU | A | 61 | -3.337 | 1.797 | 15.165 | 1.00 | 0.00 | |
| ATOM | 1001 | 3HD1 | LEU | A | 61 | -2.762 | 0.298 | 14.867 | 1.00 | 0.00 | |
| ATOM | 1002 | CD2 | LEU | A | 61 | -0.494 | 0.477 | 16.397 | 1.00 | 0.00 | C |
| ATOM | 1003 | 1HD2 | LEU | A | 61 | -0.927 | 0.182 | 17.249 | 1.00 | 0.00 | |
| ATOM | 1004 | 2HD2 | LEU | A | 61 | -0.404 | -0.305 | 15.781 | 1.00 | 0.00 | |
| ATOM | 1005 | 3HD2 | LEU | A | 61 | 0.412 | 0.851 | 16.596 | 1.00 | 0.00 | |
| ATOM | 1006 | C | LEU | A | 61 | 1.367 | 2.776 | 13.213 | 1.00 | 0.00 | C |
| ATOM | 1007 | O | LEU | A | 61 | 2.213 | 1.936 | 12.993 | 1.00 | 0.00 | O |
| ATOM | 1008 | N | LYS | A | 62 | 1.055 | 3.724 | 12.326 | 1.00 | 0.00 | N |
| ATOM | 1009 | HN | LYS | A | 62 | 0.329 | 4.373 | 12.555 | 1.00 | 0.00 | |
| ATOM | 1010 | CA | LYS | A | 62 | 1.734 | 3.857 | 11.022 | 1.00 | 0.00 | C |
| ATOM | 1011 | HA | LYS | A | 62 | 1.642 | 2.971 | 10.568 | 1.00 | 0.00 | |
| ATOM | 1012 | CB | LYS | A | 62 | 1.076 | 4.940 | 10.165 | 1.00 | 0.00 | C |
| ATOM | 1013 | HB1 | LYS | A | 62 | 0.861 | 5.716 | 10.758 | 1.00 | 0.00 | |
| ATOM | 1014 | HB2 | LYS | A | 62 | 1.734 | 5.225 | 9.468 | 1.00 | 0.00 | |
| ATOM | 1015 | CG | LYS | A | 62 | -0.204 | 4.534 | 9.453 | 1.00 | 0.00 | C |
| ATOM | 1016 | HG1 | LYS | A | 62 | -0.019 | 3.708 | 8.920 | 1.00 | 0.00 | |
| ATOM | 1017 | HG2 | LYS | A | 62 | -0.899 | 4.336 | 10.144 | 1.00 | 0.00 | |
| ATOM | 1018 | CD | LYS | A | 62 | -0.753 | 5.594 | 8.514 | 1.00 | 0.00 | C |
| ATOM | 1019 | HD1 | LYS | A | 62 | -1.547 | 5.216 | 8.037 | 1.00 | 0.00 | |
| ATOM | 1020 | HD2 | LYS | A | 62 | -1.036 | 6.384 | 9.057 | 1.00 | 0.00 | |
| ATOM | 1021 | CE | LYS | A | 62 | 0.256 | 6.057 | 7.479 | 1.00 | 0.00 | C |
| ATOM | 1022 | HE1 | LYS | A | 62 | 0.832 | 6.774 | 7.872 | 1.00 | 0.00 | |
| ATOM | 1023 | HE2 | LYS | A | 62 | 0.827 | 5.285 | 7.201 | 1.00 | 0.00 | |
| ATOM | 1024 | NZ | LYS | A | 62 | -0.411 | 6.601 | 6.276 | 1.00 | 0.00 | N |
| ATOM | 1025 | HZ1 | LYS | A | 62 | 0.280 | 6.896 | 5.616 | 1.00 | 0.00 | |
| ATOM | 1026 | HZ2 | LYS | A | 62 | -0.984 | 5.893 | 5.864 | 1.00 | 0.00 | |
| ATOM | 1027 | HZ3 | LYS | A | 62 | -0.980 | 7.382 | 6.535 | 1.00 | 0.00 | |
| ATOM | 1028 | C | LYS | A | 62 | 3.207 | 4.217 | 11.246 | 1.00 | 0.00 | C |
| ATOM | 1029 | O | LYS | A | 62 | 4.079 | 3.744 | 10.560 | 1.00 | 0.00 | O |
| ATOM | 1030 | N | GLU | A | 63 | 3.455 | 5.079 | 12.228 | 1.00 | 0.00 | N |
| ATOM | 1031 | HN | GLU | A | 63 | 2.692 | 5.385 | 12.797 | 1.00 | 0.00 | |
| ATOM | 1032 | CA | GLU | A | 63 | 4.778 | 5.596 | 12.513 | 1.00 | 0.00 | C |
| ATOM | 1033 | HA | GLU | A | 63 | 5.185 | 5.928 | 11.662 | 1.00 | 0.00 | |
| ATOM | 1034 | CB | GLU | A | 63 | 4.638 | 6.773 | 13.483 | 1.00 | 0.00 | C |
| ATOM | 1035 | HB1 | GLU | A | 63 | 3.928 | 7.387 | 13.138 | 1.00 | 0.00 | |
| ATOM | 1036 | HB2 | GLU | A | 63 | 4.366 | 6.418 | 14.378 | 1.00 | 0.00 | |
| ATOM | 1037 | CG | GLU | A | 63 | 5.896 | 7.581 | 13.670 | 1.00 | 0.00 | C |
| ATOM | 1038 | HG1 | GLU | A | 63 | 6.519 | 7.048 | 14.242 | 1.00 | 0.00 | |
| ATOM | 1039 | HG2 | GLU | A | 63 | 6.307 | 7.719 | 12.769 | 1.00 | 0.00 | |
| ATOM | 1040 | CD | GLU | A | 63 | 5.693 | 8.947 | 14.322 | 1.00 | 0.00 | C |
| ATOM | 1041 | OE1 | GLU | A | 63 | 4.549 | 9.426 | 14.368 | 1.00 | 0.00 | O |
| ATOM | 1042 | OE2 | GLU | A | 63 | 6.694 | 9.539 | 14.773 | 1.00 | 0.00 | O |
| ATOM | 1043 | C | GLU | A | 63 | 5.652 | 4.455 | 13.059 | 1.00 | 0.00 | C |
| ATOM | 1044 | O | GLU | A | 63 | 6.831 | 4.434 | 12.857 | 1.00 | 0.00 | O |
| ATOM | 1045 | N | LEU | A | 64 | 5.035 | 3.525 | 13.788 | 1.00 | 0.00 | N |
| ATOM | 1046 | HN | LEU | A | 64 | 4.050 | 3.608 | 13.936 | 1.00 | 0.00 | |
| ATOM | 1047 | CA | LEU | A | 64 | 5.734 | 2.397 | 14.375 | 1.00 | 0.00 | C |
| ATOM | 1048 | HA | LEU | A | 64 | 6.678 | 2.677 | 14.549 | 1.00 | 0.00 | |
| ATOM | 1049 | CB | LEU | A | 64 | 5.058 | 2.032 | 15.695 | 1.00 | 0.00 | C |
| ATOM | 1050 | HB1 | LEU | A | 64 | 4.121 | 1.743 | 15.497 | 1.00 | 0.00 | |
| ATOM | 1051 | HB2 | LEU | A | 64 | 5.562 | 1.272 | 16.105 | 1.00 | 0.00 | |
| ATOM | 1052 | CG | LEU | A | 64 | 5.003 | 3.173 | 16.711 | 1.00 | 0.00 | C |
| ATOM | 1053 | HG | LEU | A | 64 | 4.646 | 3.973 | 16.228 | 1.00 | 0.00 | |
| ATOM | 1054 | CD1 | LEU | A | 64 | 4.053 | 2.873 | 17.857 | 1.00 | 0.00 | C |
| ATOM | 1055 | 1HD1 | LEU | A | 64 | 4.048 | 3.642 | 18.496 | 1.00 | 0.00 | |
| ATOM | 1056 | 2HD1 | LEU | A | 64 | 3.131 | 2.732 | 17.497 | 1.00 | 0.00 | |
| ATOM | 1057 | 3HD1 | LEU | A | 64 | 4.355 | 2.047 | 18.333 | 1.00 | 0.00 | |
| ATOM | 1058 | CD2 | LEU | A | 64 | 6.387 | 3.517 | 17.243 | 1.00 | 0.00 | C |
| ATOM | 1059 | 1HD2 | LEU | A | 64 | 6.314 | 4.265 | 17.903 | 1.00 | 0.00 | |
| ATOM | 1060 | 2HD2 | LEU | A | 64 | 6.781 | 2.714 | 17.691 | 1.00 | 0.00 | |

```
ATOM   1061 3HD2 LEU A  64       6.975   3.798  16.485  1.00  0.00
ATOM   1062  C   LEU A  64       5.724   1.223  13.387  1.00  0.00           C
ATOM   1063  O   LEU A  64       4.734   0.968  12.735  1.00  0.00           O
ATOM   1064  N   GLN A  65       6.873   0.570  13.237  1.00  0.00           N
ATOM   1065  HN  GLN A  65       7.692   0.914  13.697  1.00  0.00
ATOM   1066  CA  GLN A  65       6.970  -0.607  12.439  1.00  0.00           C
ATOM   1067  HA  GLN A  65       6.070  -1.042  12.472  1.00  0.00
ATOM   1068  CB  GLN A  65       7.261  -0.283  10.984  1.00  0.00           C
ATOM   1069  HB1 GLN A  65       6.612   0.410  10.672  1.00  0.00
ATOM   1070  HB2 GLN A  65       8.192   0.076  10.913  1.00  0.00
ATOM   1071  CG  GLN A  65       7.135  -1.518  10.097  1.00  0.00           C
ATOM   1072  HG1 GLN A  65       7.799  -2.197  10.409  1.00  0.00
ATOM   1073  HG2 GLN A  65       6.210  -1.884  10.199  1.00  0.00
ATOM   1074  CD  GLN A  65       7.375  -1.287   8.629  1.00  0.00           C
ATOM   1075  OE1 GLN A  65       8.129  -2.031   8.004  1.00  0.00           O
ATOM   1076  NE2 GLN A  65       6.708  -0.282   8.063  1.00  0.00           N
ATOM   1077 1HE2 GLN A  65       6.084   0.276   8.611  1.00  0.00
ATOM   1078 2HE2 GLN A  65       6.831  -0.085   7.090  1.00  0.00
ATOM   1079  C   GLN A  65       8.043  -1.495  13.049  1.00  0.00           C
ATOM   1080  O   GLN A  65       9.202  -1.329  12.785  1.00  0.00           O
ATOM   1081  N   HIS A  66       7.607  -2.397  13.920  1.00  0.00           N
ATOM   1082  HN  HIS A  66       6.622  -2.556  13.992  1.00  0.00
ATOM   1083  CA  HIS A  66       8.475  -3.142  14.750  1.00  0.00           C
ATOM   1084  HA  HIS A  66       9.299  -3.313  14.209  1.00  0.00
ATOM   1085  CB  HIS A  66       8.845  -2.325  15.993  1.00  0.00           C
ATOM   1086  HB1 HIS A  66       9.101  -1.398  15.719  1.00  0.00
ATOM   1087  HB2 HIS A  66       8.061  -2.285  16.612  1.00  0.00
ATOM   1088  ND1 HIS A  66       9.816  -3.863  17.740  1.00  0.00           N
ATOM   1089  CG  HIS A  66       9.988  -2.921  16.732  1.00  0.00           C
ATOM   1090  CE1 HIS A  66      10.998  -4.238  18.168  1.00  0.00           C
ATOM   1091  HE1 HIS A  66      11.175  -4.905  18.892  1.00  0.00
ATOM   1092  NE2 HIS A  66      11.931  -3.570  17.465  1.00  0.00           N
ATOM   1093  HE2 HIS A  66      12.920  -3.659  17.583  1.00  0.00
ATOM   1094  CD2 HIS A  66      11.316  -2.757  16.572  1.00  0.00           C
ATOM   1095  HD2 HIS A  66      11.764  -2.149  15.917  1.00  0.00
ATOM   1096  C   HIS A  66       7.797  -4.460  15.109  1.00  0.00           C
ATOM   1097  O   HIS A  66       6.609  -4.516  15.256  1.00  0.00           O
ATOM   1098  N   GLU A  67       8.603  -5.499  15.283  1.00  0.00           N
ATOM   1099  HN  GLU A  67       9.590  -5.339  15.271  1.00  0.00
ATOM   1100  CA  GLU A  67       8.125  -6.867  15.491  1.00  0.00           C
ATOM   1101  HA  GLU A  67       7.435  -7.020  14.783  1.00  0.00
ATOM   1102  CB  GLU A  67       9.272  -7.871  15.320  1.00  0.00           C
ATOM   1103  HB1 GLU A  67       8.913  -8.794  15.461  1.00  0.00
ATOM   1104  HB2 GLU A  67       9.630  -7.792  14.390  1.00  0.00
ATOM   1105  CG  GLU A  67      10.420  -7.651  16.294  1.00  0.00           C
ATOM   1106  HG1 GLU A  67      10.541  -6.666  16.416  1.00  0.00
ATOM   1107  HG2 GLU A  67      10.170  -8.067  17.169  1.00  0.00
ATOM   1108  CD  GLU A  67      11.750  -8.235  15.862  1.00  0.00           C
ATOM   1109  OE1 GLU A  67      11.750  -9.429  15.517  1.00  0.00           O
ATOM   1110  OE2 GLU A  67      12.774  -7.481  15.860  1.00  0.00           O
ATOM   1111  C   GLU A  67       7.476  -7.014  16.872  1.00  0.00           C
ATOM   1112  O   GLU A  67       6.787  -7.986  17.107  1.00  0.00           O
ATOM   1113  N   ASN A  68       7.709  -6.055  17.775  1.00  0.00           N
ATOM   1114  HN  ASN A  68       8.257  -5.265  17.502  1.00  0.00
ATOM   1115  CA  ASN A  68       7.194  -6.112  19.152  1.00  0.00           C
ATOM   1116  HA  ASN A  68       6.782  -7.016  19.267  1.00  0.00
ATOM   1117  CB  ASN A  68       8.328  -5.975  20.160  1.00  0.00           C
ATOM   1118  HB1 ASN A  68       8.855  -5.153  19.944  1.00  0.00
ATOM   1119  HB2 ASN A  68       7.937  -5.889  21.076  1.00  0.00
ATOM   1120  CG  ASN A  68       9.263  -7.167  20.143  1.00  0.00           C
ATOM   1121  OD1 ASN A  68      10.465  -7.005  19.943  1.00  0.00           O
ATOM   1122  ND2 ASN A  68       8.721  -8.364  20.356  1.00  0.00           N
ATOM   1123 1HD2 ASN A  68       7.738  -8.446  20.518  1.00  0.00
ATOM   1124 2HD2 ASN A  68       9.297  -9.181  20.354  1.00  0.00
ATOM   1125  C   ASN A  68       6.126  -5.032  19.359  1.00  0.00           C
ATOM   1126  O   ASN A  68       5.755  -4.699  20.503  1.00  0.00           O
ATOM   1127  N   ILE A  69       5.621  -4.492  18.238  1.00  0.00           N
ATOM   1128  HN  ILE A  69       6.018  -4.758  17.359  1.00  0.00
ATOM   1129  CA  ILE A  69       4.522  -3.537  18.244  1.00  0.00           C
ATOM   1130  HA  ILE A  69       4.120  -3.542  19.159  1.00  0.00
ATOM   1131  CB  ILE A  69       5.048  -2.123  17.942  1.00  0.00           C
```

| | | | | | | | | | | | |
|---|---|---|---|---|---|---|---|---|---|---|---|
| ATOM | 1132 | HB | ILE | A | 69 | 5.513 | -2.129 | 17.057 | 1.00 | 0.00 | |
| ATOM | 1133 | CG2 | ILE | A | 69 | 3.884 | -1.146 | 17.836 | 1.00 | 0.00 | C |
| ATOM | 1134 | 1HG2 | ILE | A | 69 | 4.234 | -0.230 | 17.640 | 1.00 | 0.00 | |
| ATOM | 1135 | 2HG2 | ILE | A | 69 | 3.273 | -1.434 | 17.099 | 1.00 | 0.00 | |
| ATOM | 1136 | 3HG2 | ILE | A | 69 | 3.381 | -1.132 | 18.700 | 1.00 | 0.00 | |
| ATOM | 1137 | CG1 | ILE | A | 69 | 6.096 | -1.678 | 18.967 | 1.00 | 0.00 | C |
| ATOM | 1138 | 1HG1 | ILE | A | 69 | 5.703 | -1.759 | 19.883 | 1.00 | 0.00 | |
| ATOM | 1139 | 2HG1 | ILE | A | 69 | 6.892 | -2.278 | 18.892 | 1.00 | 0.00 | |
| ATOM | 1140 | CD | ILE | A | 69 | 6.562 | -0.250 | 18.778 | 1.00 | 0.00 | C |
| ATOM | 1141 | HD1 | ILE | A | 69 | 7.242 | -0.026 | 19.477 | 1.00 | 0.00 | |
| ATOM | 1142 | HD2 | ILE | A | 69 | 6.971 | -0.150 | 17.871 | 1.00 | 0.00 | |
| ATOM | 1143 | HD3 | ILE | A | 69 | 5.781 | 0.369 | 18.862 | 1.00 | 0.00 | |
| ATOM | 1144 | C | ILE | A | 69 | 3.485 | -3.988 | 17.214 | 1.00 | 0.00 | C |
| ATOM | 1145 | O | ILE | A | 69 | 3.796 | -4.189 | 16.035 | 1.00 | 0.00 | O |
| ATOM | 1146 | N | VAL | A | 70 | 2.241 | -4.100 | 17.652 | 1.00 | 0.00 | N |
| ATOM | 1147 | HN | VAL | A | 70 | 2.021 | -3.775 | 18.572 | 1.00 | 0.00 | |
| ATOM | 1148 | CA | VAL | A | 70 | 1.198 | -4.672 | 16.852 | 1.00 | 0.00 | C |
| ATOM | 1149 | HA | VAL | A | 70 | 1.465 | -5.621 | 16.685 | 1.00 | 0.00 | |
| ATOM | 1150 | CB | VAL | A | 70 | -0.150 | -4.702 | 17.595 | 1.00 | 0.00 | C |
| ATOM | 1151 | HB | VAL | A | 70 | 0.007 | -5.090 | 18.503 | 1.00 | 0.00 | |
| ATOM | 1152 | CG1 | VAL | A | 70 | -0.717 | -3.302 | 17.786 | 1.00 | 0.00 | C |
| ATOM | 1153 | 1HG1 | VAL | A | 70 | -1.590 | -3.359 | 18.270 | 1.00 | 0.00 | |
| ATOM | 1154 | 2HG1 | VAL | A | 70 | -0.075 | -2.752 | 18.320 | 1.00 | 0.00 | |
| ATOM | 1155 | 3HG1 | VAL | A | 70 | -0.859 | -2.875 | 16.893 | 1.00 | 0.00 | |
| ATOM | 1156 | CG2 | VAL | A | 70 | -1.149 | -5.594 | 16.888 | 1.00 | 0.00 | C |
| ATOM | 1157 | 1HG2 | VAL | A | 70 | -2.012 | -5.593 | 17.393 | 1.00 | 0.00 | |
| ATOM | 1158 | 2HG2 | VAL | A | 70 | -1.304 | -5.252 | 15.961 | 1.00 | 0.00 | |
| ATOM | 1159 | 3HG2 | VAL | A | 70 | -0.790 | -6.526 | 16.843 | 1.00 | 0.00 | |
| ATOM | 1160 | C | VAL | A | 70 | 1.131 | -3.894 | 15.541 | 1.00 | 0.00 | C |
| ATOM | 1161 | O | VAL | A | 70 | 1.115 | -2.671 | 15.548 | 1.00 | 0.00 | O |
| ATOM | 1162 | N | ALA | A | 71 | 1.164 | -4.643 | 14.416 | 1.00 | 0.00 | N |
| ATOM | 1163 | HN | ALA | A | 71 | 1.176 | -5.638 | 14.511 | 1.00 | 0.00 | |
| ATOM | 1164 | CA | ALA | A | 71 | 1.183 | -4.089 | 13.097 | 1.00 | 0.00 | C |
| ATOM | 1165 | HA | ALA | A | 71 | 1.801 | -3.303 | 13.117 | 1.00 | 0.00 | |
| ATOM | 1166 | CB | ALA | A | 71 | 1.716 | -5.134 | 12.149 | 1.00 | 0.00 | C |
| ATOM | 1167 | HB1 | ALA | A | 71 | 1.736 | -4.763 | 11.221 | 1.00 | 0.00 | |
| ATOM | 1168 | HB2 | ALA | A | 71 | 2.642 | -5.393 | 12.425 | 1.00 | 0.00 | |
| ATOM | 1169 | HB3 | ALA | A | 71 | 1.124 | -5.939 | 12.173 | 1.00 | 0.00 | |
| ATOM | 1170 | C | ALA | A | 71 | -0.218 | -3.619 | 12.697 | 1.00 | 0.00 | C |
| ATOM | 1171 | O | ALA | A | 71 | -1.179 | -4.278 | 12.970 | 1.00 | 0.00 | O |
| ATOM | 1172 | N | LEU | A | 72 | -0.296 | -2.463 | 12.037 | 1.00 | 0.00 | N |
| ATOM | 1173 | HN | LEU | A | 72 | 0.498 | -1.856 | 12.060 | 1.00 | 0.00 | |
| ATOM | 1174 | CA | LEU | A | 72 | -1.463 | -2.031 | 11.287 | 1.00 | 0.00 | C |
| ATOM | 1175 | HA | LEU | A | 72 | -2.234 | -2.612 | 11.547 | 1.00 | 0.00 | |
| ATOM | 1176 | CB | LEU | A | 72 | -1.785 | -0.575 | 11.631 | 1.00 | 0.00 | C |
| ATOM | 1177 | HB1 | LEU | A | 72 | -2.004 | -0.529 | 12.606 | 1.00 | 0.00 | |
| ATOM | 1178 | HB2 | LEU | A | 72 | -0.970 | -0.026 | 11.446 | 1.00 | 0.00 | |
| ATOM | 1179 | CG | LEU | A | 72 | -2.944 | 0.046 | 10.867 | 1.00 | 0.00 | C |
| ATOM | 1180 | HG | LEU | A | 72 | -2.762 | -0.057 | 9.889 | 1.00 | 0.00 | |
| ATOM | 1181 | CD1 | LEU | A | 72 | -4.220 | -0.669 | 11.180 | 1.00 | 0.00 | C |
| ATOM | 1182 | 1HD1 | LEU | A | 72 | -4.971 | -0.249 | 10.670 | 1.00 | 0.00 | |
| ATOM | 1183 | 2HD1 | LEU | A | 72 | -4.136 | -1.630 | 10.918 | 1.00 | 0.00 | |
| ATOM | 1184 | 3HD1 | LEU | A | 72 | -4.406 | -0.605 | 12.161 | 1.00 | 0.00 | |
| ATOM | 1185 | CD2 | LEU | A | 72 | -3.085 | 1.528 | 11.179 | 1.00 | 0.00 | C |
| ATOM | 1186 | 1HD2 | LEU | A | 72 | -3.854 | 1.906 | 10.663 | 1.00 | 0.00 | |
| ATOM | 1187 | 2HD2 | LEU | A | 72 | -3.250 | 1.648 | 12.158 | 1.00 | 0.00 | |
| ATOM | 1188 | 3HD2 | LEU | A | 72 | -2.244 | 2.004 | 10.922 | 1.00 | 0.00 | |
| ATOM | 1189 | C | LEU | A | 72 | -1.132 | -2.169 | 9.796 | 1.00 | 0.00 | C |
| ATOM | 1190 | O | LEU | A | 72 | -0.185 | -1.542 | 9.319 | 1.00 | 0.00 | O |
| ATOM | 1191 | N | TYR | A | 73 | -1.946 | -2.943 | 9.068 | 1.00 | 0.00 | N |
| ATOM | 1192 | HN | TYR | A | 73 | -2.742 | -3.358 | 9.509 | 1.00 | 0.00 | |
| ATOM | 1193 | CA | TYR | A | 73 | -1.710 | -3.201 | 7.651 | 1.00 | 0.00 | C |
| ATOM | 1194 | HA | TYR | A | 73 | -0.737 | -3.012 | 7.516 | 1.00 | 0.00 | |
| ATOM | 1195 | CB | TYR | A | 73 | -2.024 | -4.655 | 7.313 | 1.00 | 0.00 | C |
| ATOM | 1196 | HB1 | TYR | A | 73 | -2.984 | -4.835 | 7.529 | 1.00 | 0.00 | |
| ATOM | 1197 | HB2 | TYR | A | 73 | -1.870 | -4.798 | 6.335 | 1.00 | 0.00 | |
| ATOM | 1198 | CG | TYR | A | 73 | -1.187 | -5.647 | 8.063 | 1.00 | 0.00 | C |
| ATOM | 1199 | CD1 | TYR | A | 73 | 0.193 | -5.600 | 7.990 | 1.00 | 0.00 | C |
| ATOM | 1200 | HD1 | TYR | A | 73 | 0.631 | -4.897 | 7.430 | 1.00 | 0.00 | |
| ATOM | 1201 | CE1 | TYR | A | 73 | 0.980 | -6.508 | 8.677 | 1.00 | 0.00 | C |
| ATOM | 1202 | HE1 | TYR | A | 73 | 1.976 | -6.474 | 8.594 | 1.00 | 0.00 | |

| | | | | | | | | | | | |
|---|---|---|---|---|---|---|---|---|---|---|---|
| ATOM | 1203 | CZ | TYR | A | 73 | 0.380 | -7.465 | 9.480 | 1.00 | 0.00 | C |
| ATOM | 1204 | OH | TYR | A | 73 | 1.146 | -8.348 | 10.188 | 1.00 | 0.00 | O |
| ATOM | 1205 | HH | TYR | A | 73 | 0.552 | -8.962 | 10.708 | 1.00 | 0.00 | |
| ATOM | 1206 | CD2 | TYR | A | 73 | -1.767 | -6.619 | 8.853 | 1.00 | 0.00 | C |
| ATOM | 1207 | HD2 | TYR | A | 73 | -2.764 | -6.667 | 8.917 | 1.00 | 0.00 | |
| ATOM | 1208 | CE2 | TYR | A | 73 | -0.999 | -7.527 | 9.557 | 1.00 | 0.00 | C |
| ATOM | 1209 | HE2 | TYR | A | 73 | -1.439 | -8.226 | 10.120 | 1.00 | 0.00 | |
| ATOM | 1210 | C | TYR | A | 73 | -2.561 | -2.277 | 6.767 | 1.00 | 0.00 | C |
| ATOM | 1211 | O | TYR | A | 73 | -2.141 | -1.926 | 5.692 | 1.00 | 0.00 | O |
| ATOM | 1212 | N | ASP | A | 74 | -3.770 | -1.923 | 7.220 | 1.00 | 0.00 | N |
| ATOM | 1213 | HN | ASP | A | 74 | -4.027 | -2.125 | 8.165 | 1.00 | 0.00 | |
| ATOM | 1214 | CA | ASP | A | 74 | -4.699 | -1.254 | 6.354 | 1.00 | 0.00 | C |
| ATOM | 1215 | HA | ASP | A | 74 | -4.154 | -0.550 | 5.898 | 1.00 | 0.00 | |
| ATOM | 1216 | CB | ASP | A | 74 | -5.282 | -2.224 | 5.329 | 1.00 | 0.00 | C |
| ATOM | 1217 | HB1 | ASP | A | 74 | -4.542 | -2.802 | 4.984 | 1.00 | 0.00 | |
| ATOM | 1218 | HB2 | ASP | A | 74 | -5.965 | -2.794 | 5.786 | 1.00 | 0.00 | |
| ATOM | 1219 | CG | ASP | A | 74 | -5.946 | -1.551 | 4.144 | 1.00 | 0.00 | C |
| ATOM | 1220 | OD1 | ASP | A | 74 | -5.585 | -0.408 | 3.854 | 1.00 | 0.00 | O |
| ATOM | 1221 | OD2 | ASP | A | 74 | -6.781 | -2.205 | 3.486 | 1.00 | 0.00 | O |
| ATOM | 1222 | C | ASP | A | 74 | -5.824 | -0.622 | 7.165 | 1.00 | 0.00 | C |
| ATOM | 1223 | O | ASP | A | 74 | -6.193 | -1.104 | 8.250 | 1.00 | 0.00 | O |
| ATOM | 1224 | N | VAL | A | 75 | -6.356 | 0.459 | 6.591 | 1.00 | 0.00 | N |
| ATOM | 1225 | HN | VAL | A | 75 | -5.924 | 0.808 | 5.760 | 1.00 | 0.00 | |
| ATOM | 1226 | CA | VAL | A | 75 | -7.523 | 1.161 | 7.097 | 1.00 | 0.00 | C |
| ATOM | 1227 | HA | VAL | A | 75 | -7.899 | 0.616 | 7.846 | 1.00 | 0.00 | |
| ATOM | 1228 | CB | VAL | A | 75 | -7.137 | 2.535 | 7.656 | 1.00 | 0.00 | C |
| ATOM | 1229 | HB | VAL | A | 75 | -6.838 | 3.102 | 6.889 | 1.00 | 0.00 | |
| ATOM | 1230 | CG1 | VAL | A | 75 | -8.318 | 3.231 | 8.299 | 1.00 | 0.00 | C |
| ATOM | 1231 | 1HG1 | VAL | A | 75 | -8.029 | 4.121 | 8.651 | 1.00 | 0.00 | |
| ATOM | 1232 | 2HG1 | VAL | A | 75 | -9.040 | 3.360 | 7.619 | 1.00 | 0.00 | |
| ATOM | 1233 | 3HG1 | VAL | A | 75 | -8.665 | 2.672 | 9.052 | 1.00 | 0.00 | |
| ATOM | 1234 | CG2 | VAL | A | 75 | -5.988 | 2.420 | 8.645 | 1.00 | 0.00 | C |
| ATOM | 1235 | 1HG2 | VAL | A | 75 | -5.756 | 3.328 | 8.994 | 1.00 | 0.00 | |
| ATOM | 1236 | 2HG2 | VAL | A | 75 | -6.260 | 1.830 | 9.405 | 1.00 | 0.00 | |
| ATOM | 1237 | 3HG2 | VAL | A | 75 | -5.192 | 2.026 | 8.186 | 1.00 | 0.00 | |
| ATOM | 1238 | C | VAL | A | 75 | -8.525 | 1.289 | 5.945 | 1.00 | 0.00 | C |
| ATOM | 1239 | O | VAL | A | 75 | -8.167 | 1.777 | 4.892 | 1.00 | 0.00 | O |
| ATOM | 1240 | N | GLN | A | 76 | -9.746 | 0.778 | 6.150 | 1.00 | 0.00 | N |
| ATOM | 1241 | HN | GLN | A | 76 | -9.927 | 0.288 | 7.003 | 1.00 | 0.00 | |
| ATOM | 1242 | CA | GLN | A | 76 | -10.814 | 0.910 | 5.180 | 1.00 | 0.00 | C |
| ATOM | 1243 | HA | GLN | A | 76 | -10.416 | 1.406 | 4.408 | 1.00 | 0.00 | |
| ATOM | 1244 | CB | GLN | A | 76 | -11.316 | -0.447 | 4.677 | 1.00 | 0.00 | C |
| ATOM | 1245 | HB1 | GLN | A | 76 | -11.623 | -0.982 | 5.464 | 1.00 | 0.00 | |
| ATOM | 1246 | HB2 | GLN | A | 76 | -12.087 | -0.289 | 4.060 | 1.00 | 0.00 | |
| ATOM | 1247 | CG | GLN | A | 76 | -10.277 | -1.265 | 3.930 | 1.00 | 0.00 | C |
| ATOM | 1248 | HG1 | GLN | A | 76 | -9.462 | -1.348 | 4.503 | 1.00 | 0.00 | |
| ATOM | 1249 | HG2 | GLN | A | 76 | -10.653 | -2.174 | 3.751 | 1.00 | 0.00 | |
| ATOM | 1250 | CD | GLN | A | 76 | -9.873 | -0.648 | 2.612 | 1.00 | 0.00 | C |
| ATOM | 1251 | OE1 | GLN | A | 76 | -10.705 | -0.308 | 1.790 | 1.00 | 0.00 | O |
| ATOM | 1252 | NE2 | GLN | A | 76 | -8.577 | -0.488 | 2.393 | 1.00 | 0.00 | N |
| ATOM | 1253 | 1HE2 | GLN | A | 76 | -7.914 | -0.771 | 3.086 | 1.00 | 0.00 | |
| ATOM | 1254 | 2HE2 | GLN | A | 76 | -8.261 | -0.084 | 1.535 | 1.00 | 0.00 | |
| ATOM | 1255 | C | GLN | A | 76 | -11.961 | 1.675 | 5.839 | 1.00 | 0.00 | C |
| ATOM | 1256 | O | GLN | A | 76 | -12.435 | 1.308 | 6.894 | 1.00 | 0.00 | O |
| ATOM | 1257 | N | GLU | A | 77 | -12.395 | 2.731 | 5.155 | 1.00 | 0.00 | N |
| ATOM | 1258 | HN | GLU | A | 77 | -11.945 | 2.947 | 4.289 | 1.00 | 0.00 | |
| ATOM | 1259 | CA | GLU | A | 77 | -13.464 | 3.573 | 5.583 | 1.00 | 0.00 | C |
| ATOM | 1260 | HA | GLU | A | 77 | -13.656 | 3.370 | 6.543 | 1.00 | 0.00 | |
| ATOM | 1261 | CB | GLU | A | 77 | -13.014 | 5.021 | 5.450 | 1.00 | 0.00 | C |
| ATOM | 1262 | HB1 | GLU | A | 77 | -12.136 | 5.122 | 5.918 | 1.00 | 0.00 | |
| ATOM | 1263 | HB2 | GLU | A | 77 | -12.899 | 5.227 | 4.478 | 1.00 | 0.00 | |
| ATOM | 1264 | CG | GLU | A | 77 | -13.982 | 6.031 | 6.036 | 1.00 | 0.00 | C |
| ATOM | 1265 | HG1 | GLU | A | 77 | -14.646 | 6.273 | 5.328 | 1.00 | 0.00 | |
| ATOM | 1266 | HG2 | GLU | A | 77 | -14.456 | 5.599 | 6.804 | 1.00 | 0.00 | |
| ATOM | 1267 | CD | GLU | A | 77 | -13.331 | 7.312 | 6.537 | 1.00 | 0.00 | C |
| ATOM | 1268 | OE1 | GLU | A | 77 | -12.129 | 7.269 | 6.870 | 1.00 | 0.00 | O |
| ATOM | 1269 | OE2 | GLU | A | 77 | -14.023 | 8.350 | 6.597 | 1.00 | 0.00 | O |
| ATOM | 1270 | C | GLU | A | 77 | -14.699 | 3.268 | 4.722 | 1.00 | 0.00 | C |
| ATOM | 1271 | O | GLU | A | 77 | -14.666 | 3.423 | 3.514 | 1.00 | 0.00 | O |
| ATOM | 1272 | N | LEU | A | 78 | -15.770 | 2.792 | 5.366 | 1.00 | 0.00 | N |
| ATOM | 1273 | HN | LEU | A | 78 | -15.650 | 2.445 | 6.296 | 1.00 | 0.00 | |

```
ATOM   1274  CA  LEU A  78     -17.113   2.753   4.776  1.00  0.00           C
ATOM   1275  HA  LEU A  78     -17.043   2.854   3.784  1.00  0.00
ATOM   1276  CB  LEU A  78     -17.759   1.393   5.068  1.00  0.00           C
ATOM   1277  HB1 LEU A  78     -17.448   1.091   5.969  1.00  0.00
ATOM   1278  HB2 LEU A  78     -18.751   1.519   5.082  1.00  0.00
ATOM   1279  CG  LEU A  78     -17.437   0.282   4.059  1.00  0.00           C
ATOM   1280  HG  LEU A  78     -17.885   0.527   3.199  1.00  0.00
ATOM   1281  CD1 LEU A  78     -15.937   0.162   3.820  1.00  0.00           C
ATOM   1282 1HD1 LEU A  78     -15.762  -0.568   3.160  1.00  0.00
ATOM   1283 2HD1 LEU A  78     -15.586   1.027   3.461  1.00  0.00
ATOM   1284 3HD1 LEU A  78     -15.478  -0.051   4.682  1.00  0.00
ATOM   1285  CD2 LEU A  78     -17.975  -1.059   4.518  1.00  0.00           C
ATOM   1286 1HD2 LEU A  78     -17.749  -1.759   3.840  1.00  0.00
ATOM   1287 2HD2 LEU A  78     -17.563  -1.300   5.397  1.00  0.00
ATOM   1288 3HD2 LEU A  78     -18.968  -1.002   4.620  1.00  0.00
ATOM   1289  C   LEU A  78     -17.913   3.926   5.346  1.00  0.00           C
ATOM   1290  O   LEU A  78     -17.424   4.642   6.224  1.00  0.00           O
ATOM   1291  N   PRO A  79     -19.135   4.225   4.832  1.00  0.00           N
ATOM   1292  CD  PRO A  79     -19.820   3.506   3.745  1.00  0.00           C
ATOM   1293  HD1 PRO A  79     -20.359   2.744   4.105  1.00  0.00
ATOM   1294  HD2 PRO A  79     -19.162   3.162   3.075  1.00  0.00
ATOM   1295  CA  PRO A  79     -19.903   5.378   5.323  1.00  0.00           C
ATOM   1296  HA  PRO A  79     -19.323   6.192   5.324  1.00  0.00
ATOM   1297  CB  PRO A  79     -21.097   5.466   4.356  1.00  0.00           C
ATOM   1298  HB1 PRO A  79     -21.928   5.114   4.787  1.00  0.00
ATOM   1299  HB2 PRO A  79     -21.247   6.411   4.064  1.00  0.00
ATOM   1300  CG  PRO A  79     -20.699   4.593   3.164  1.00  0.00           C
ATOM   1301  HG1 PRO A  79     -21.508   4.193   2.733  1.00  0.00
ATOM   1302  HG2 PRO A  79     -20.191   5.127   2.488  1.00  0.00
ATOM   1303  C   PRO A  79     -20.365   5.197   6.780  1.00  0.00           C
ATOM   1304  O   PRO A  79     -20.465   6.169   7.503  1.00  0.00           O
ATOM   1305  N   ASN A  80     -20.604   3.936   7.177  1.00  0.00           N
ATOM   1306  HN  ASN A  80     -20.461   3.208   6.506  1.00  0.00
ATOM   1307  CA  ASN A  80     -21.062   3.536   8.529  1.00  0.00           C
ATOM   1308  HA  ASN A  80     -21.639   4.302   8.813  1.00  0.00
ATOM   1309  CB  ASN A  80     -21.850   2.223   8.496  1.00  0.00           C
ATOM   1310  HB1 ASN A  80     -21.267   1.505   8.116  1.00  0.00
ATOM   1311  HB2 ASN A  80     -22.113   1.980   9.430  1.00  0.00
ATOM   1312  CG  ASN A  80     -23.106   2.313   7.654  1.00  0.00           C
ATOM   1313  OD1 ASN A  80     -23.861   3.280   7.764  1.00  0.00           O
ATOM   1314  ND2 ASN A  80     -23.331   1.317   6.808  1.00  0.00           N
ATOM   1315 1HD2 ASN A  80     -22.686   0.555   6.749  1.00  0.00
ATOM   1316 2HD2 ASN A  80     -24.146   1.327   6.229  1.00  0.00
ATOM   1317  C   ASN A  80     -19.890   3.331   9.499  1.00  0.00           C
ATOM   1318  O   ASN A  80     -19.996   3.689  10.669  1.00  0.00           O
ATOM   1319  N   SER A  81     -18.806   2.703   9.015  1.00  0.00           N
ATOM   1320  HN  SER A  81     -18.665   2.701   8.025  1.00  0.00
ATOM   1321  CA  SER A  81     -17.805   2.012   9.866  1.00  0.00           C
ATOM   1322  HA  SER A  81     -17.932   2.347  10.800  1.00  0.00
ATOM   1323  CB  SER A  81     -18.052   0.531   9.849  1.00  0.00           C
ATOM   1324  HB1 SER A  81     -17.839   0.149   8.950  1.00  0.00
ATOM   1325  HB2 SER A  81     -17.499   0.074  10.545  1.00  0.00
ATOM   1326  OG  SER A  81     -19.418   0.275  10.128  1.00  0.00           O
ATOM   1327  HG1 SER A  81     -19.581  -0.711  10.116  1.00  0.00
ATOM   1328  C   SER A  81     -16.375   2.323   9.400  1.00  0.00           C
ATOM   1329  O   SER A  81     -16.149   2.740   8.261  1.00  0.00           O
ATOM   1330  N   VAL A  82     -15.416   2.129  10.309  1.00  0.00           N
ATOM   1331  HN  VAL A  82     -15.677   1.997  11.265  1.00  0.00
ATOM   1332  CA  VAL A  82     -14.016   2.103   9.960  1.00  0.00           C
ATOM   1333  HA  VAL A  82     -13.981   2.277   8.976  1.00  0.00
ATOM   1334  CB  VAL A  82     -13.208   3.208  10.658  1.00  0.00           C
ATOM   1335  HB  VAL A  82     -13.384   3.128  11.639  1.00  0.00
ATOM   1336  CG1 VAL A  82     -11.727   3.054  10.409  1.00  0.00           C
ATOM   1337 1HG1 VAL A  82     -11.232   3.787  10.876  1.00  0.00
ATOM   1338 2HG1 VAL A  82     -11.420   2.169  10.760  1.00  0.00
ATOM   1339 3HG1 VAL A  82     -11.547   3.104   9.427  1.00  0.00
ATOM   1340  CG2 VAL A  82     -13.631   4.581  10.215  1.00  0.00           C
ATOM   1341 1HG2 VAL A  82     -13.083   5.269  10.691  1.00  0.00
ATOM   1342 2HG2 VAL A  82     -13.494   4.670   9.228  1.00  0.00
ATOM   1343 3HG2 VAL A  82     -14.598   4.718  10.431  1.00  0.00
ATOM   1344  C   VAL A  82     -13.474   0.722  10.326  1.00  0.00           C
```

```
ATOM   1345  O   VAL A  82     -13.777   0.185  11.369  1.00  0.00           O
ATOM   1346  N   PHE A  83     -12.635   0.181   9.450  1.00  0.00           N
ATOM   1347  HN  PHE A  83     -12.405   0.696   8.624  1.00  0.00
ATOM   1348  CA  PHE A  83     -12.049  -1.104   9.640  1.00  0.00           C
ATOM   1349  HA  PHE A  83     -12.434  -1.465  10.489  1.00  0.00
ATOM   1350  CB  PHE A  83     -12.411  -2.025   8.473  1.00  0.00           C
ATOM   1351  HB1 PHE A  83     -12.247  -1.534   7.618  1.00  0.00
ATOM   1352  HB2 PHE A  83     -11.823  -2.833   8.512  1.00  0.00
ATOM   1353  CG  PHE A  83     -13.834  -2.499   8.458  1.00  0.00           C
ATOM   1354  CD1 PHE A  83     -14.211  -3.625   9.159  1.00  0.00           C
ATOM   1355  HD1 PHE A  83     -13.531  -4.118   9.702  1.00  0.00
ATOM   1356  CE1 PHE A  83     -15.524  -4.086   9.123  1.00  0.00           C
ATOM   1357  HE1 PHE A  83     -15.776  -4.915   9.623  1.00  0.00
ATOM   1358  CZ  PHE A  83     -16.476  -3.403   8.401  1.00  0.00           C
ATOM   1359  HZ  PHE A  83     -17.420  -3.733   8.372  1.00  0.00
ATOM   1360  CD2 PHE A  83     -14.797  -1.819   7.740  1.00  0.00           C
ATOM   1361  HD2 PHE A  83     -14.543  -0.998   7.228  1.00  0.00
ATOM   1362  CE2 PHE A  83     -16.117  -2.261   7.719  1.00  0.00           C
ATOM   1363  HE2 PHE A  83     -16.806  -1.748   7.207  1.00  0.00
ATOM   1364  C   PHE A  83     -10.537  -0.939   9.752  1.00  0.00           C
ATOM   1365  O   PHE A  83      -9.899  -0.327   8.875  1.00  0.00           O
ATOM   1366  N   LEU A  84      -9.978  -1.505  10.821  1.00  0.00           N
ATOM   1367  HN  LEU A  84     -10.574  -1.866  11.538  1.00  0.00
ATOM   1368  CA  LEU A  84      -8.539  -1.621  10.990  1.00  0.00           C
ATOM   1369  HA  LEU A  84      -8.090  -1.036  10.315  1.00  0.00
ATOM   1370  CB  LEU A  84      -8.182  -1.158  12.401  1.00  0.00           C
ATOM   1371  HB1 LEU A  84      -8.960  -1.356  12.997  1.00  0.00
ATOM   1372  HB2 LEU A  84      -7.387  -1.682  12.706  1.00  0.00
ATOM   1373  CG  LEU A  84      -7.849   0.321  12.545  1.00  0.00           C
ATOM   1374  HG  LEU A  84      -6.934   0.468  12.168  1.00  0.00
ATOM   1375  CD1 LEU A  84      -8.791   1.190  11.750  1.00  0.00           C
ATOM   1376 1HD1 LEU A  84      -8.540   2.151  11.870  1.00  0.00
ATOM   1377 2HD1 LEU A  84      -8.730   0.948  10.782  1.00  0.00
ATOM   1378 3HD1 LEU A  84      -9.727   1.047  12.071  1.00  0.00
ATOM   1379  CD2 LEU A  84      -7.847   0.727  14.008  1.00  0.00           C
ATOM   1380 1HD2 LEU A  84      -7.627   1.700  14.084  1.00  0.00
ATOM   1381 2HD2 LEU A  84      -8.750   0.558  14.402  1.00  0.00
ATOM   1382 3HD2 LEU A  84      -7.162   0.191  14.502  1.00  0.00
ATOM   1383  C   LEU A  84      -8.135  -3.082  10.767  1.00  0.00           C
ATOM   1384  O   LEU A  84      -8.675  -3.992  11.420  1.00  0.00           O
ATOM   1385  N   VAL A  85      -7.230  -3.304   9.818  1.00  0.00           N
ATOM   1386  HN  VAL A  85      -6.931  -2.546   9.238  1.00  0.00
ATOM   1387  CA  VAL A  85      -6.669  -4.622   9.609  1.00  0.00           C
ATOM   1388  HA  VAL A  85      -7.308  -5.273  10.019  1.00  0.00
ATOM   1389  CB  VAL A  85      -6.536  -4.960   8.122  1.00  0.00           C
ATOM   1390  HB  VAL A  85      -5.852  -4.342   7.734  1.00  0.00
ATOM   1391  CG1 VAL A  85      -6.038  -6.379   7.917  1.00  0.00           C
ATOM   1392 1HG1 VAL A  85      -5.962  -6.568   6.938  1.00  0.00
ATOM   1393 2HG1 VAL A  85      -5.141  -6.483   8.347  1.00  0.00
ATOM   1394 3HG1 VAL A  85      -6.683  -7.022   8.331  1.00  0.00
ATOM   1395  CG2 VAL A  85      -7.831  -4.719   7.367  1.00  0.00           C
ATOM   1396 1HG2 VAL A  85      -7.703  -4.951   6.403  1.00  0.00
ATOM   1397 2HG2 VAL A  85      -8.554  -5.291   7.754  1.00  0.00
ATOM   1398 3HG2 VAL A  85      -8.089  -3.756   7.447  1.00  0.00
ATOM   1399  C   VAL A  85      -5.313  -4.657  10.292  1.00  0.00           C
ATOM   1400  O   VAL A  85      -4.415  -3.932   9.907  1.00  0.00           O
ATOM   1401  N   MET A  86      -5.180  -5.525  11.291  1.00  0.00           N
ATOM   1402  HN  MET A  86      -5.919  -6.175  11.469  1.00  0.00
ATOM   1403  CA  MET A  86      -4.009  -5.559  12.122  1.00  0.00           C
ATOM   1404  HA  MET A  86      -3.349  -4.965  11.661  1.00  0.00
ATOM   1405  CB  MET A  86      -4.342  -5.042  13.520  1.00  0.00           C
ATOM   1406  HB1 MET A  86      -4.964  -5.682  13.971  1.00  0.00
ATOM   1407  HB2 MET A  86      -3.500  -4.963  14.054  1.00  0.00
ATOM   1408  CG  MET A  86      -4.994  -3.711  13.486  1.00  0.00           C
ATOM   1409  HG1 MET A  86      -4.349  -3.117  13.006  1.00  0.00
ATOM   1410  HG2 MET A  86      -5.812  -3.836  12.925  1.00  0.00
ATOM   1411  SD  MET A  86      -5.367  -3.136  15.131  1.00  0.00           S
ATOM   1412  CE  MET A  86      -3.741  -2.554  15.603  1.00  0.00           C
ATOM   1413  HE1 MET A  86      -3.775  -2.187  16.532  1.00  0.00
ATOM   1414  HE2 MET A  86      -3.092  -3.314  15.569  1.00  0.00
ATOM   1415  HE3 MET A  86      -3.448  -1.836  14.972  1.00  0.00
```

```
ATOM   1416  C   MET A  86      -3.468  -6.976  12.227  1.00  0.00           C
ATOM   1417  O   MET A  86      -4.104  -7.911  11.885  1.00  0.00           O
ATOM   1418  N   GLU A  87      -2.249  -7.063  12.713  1.00  0.00           N
ATOM   1419  HN  GLU A  87      -1.744  -6.214  12.870  1.00  0.00
ATOM   1420  CA  GLU A  87      -1.597  -8.307  13.033  1.00  0.00           C
ATOM   1421  HA  GLU A  87      -1.489  -8.837  12.192  1.00  0.00
ATOM   1422  CB  GLU A  87      -0.236  -7.977  13.633  1.00  0.00           C
ATOM   1423  HB1 GLU A  87       0.299  -7.469  12.958  1.00  0.00
ATOM   1424  HB2 GLU A  87      -0.371  -7.410  14.446  1.00  0.00
ATOM   1425  CG  GLU A  87       0.558  -9.168  14.038  1.00  0.00           C
ATOM   1426  HG1 GLU A  87       0.069  -9.640  14.771  1.00  0.00
ATOM   1427  HG2 GLU A  87       0.642  -9.776  13.248  1.00  0.00
ATOM   1428  CD  GLU A  87       1.936  -8.803  14.524  1.00  0.00           C
ATOM   1429  OE1 GLU A  87       2.094  -7.692  15.088  1.00  0.00           O
ATOM   1430  OE2 GLU A  87       2.864  -9.625  14.291  1.00  0.00           O
ATOM   1431  C   GLU A  87      -2.468  -9.090  14.009  1.00  0.00           C
ATOM   1432  O   GLU A  87      -2.978  -8.524  14.980  1.00  0.00           O
ATOM   1433  N   TYR A  88      -2.680 -10.366  13.705  1.00  0.00           N
ATOM   1434  HN  TYR A  88      -2.351 -10.717  12.828  1.00  0.00
ATOM   1435  CA  TYR A  88      -3.381 -11.273  14.615  1.00  0.00           C
ATOM   1436  HA  TYR A  88      -4.041 -10.704  15.105  1.00  0.00
ATOM   1437  CB  TYR A  88      -4.125 -12.375  13.865  1.00  0.00           C
ATOM   1438  HB1 TYR A  88      -4.814 -11.951  13.278  1.00  0.00
ATOM   1439  HB2 TYR A  88      -3.468 -12.875  13.301  1.00  0.00
ATOM   1440  CG  TYR A  88      -4.823 -13.369  14.757  1.00  0.00           C
ATOM   1441  CD1 TYR A  88      -5.885 -12.962  15.547  1.00  0.00           C
ATOM   1442  HD1 TYR A  88      -6.190 -12.011  15.504  1.00  0.00
ATOM   1443  CE1 TYR A  88      -6.534 -13.842  16.395  1.00  0.00           C
ATOM   1444  HE1 TYR A  88      -7.291 -13.522  16.965  1.00  0.00
ATOM   1445  CZ  TYR A  88      -6.136 -15.162  16.453  1.00  0.00           C
ATOM   1446  OH  TYR A  88      -6.799 -15.979  17.320  1.00  0.00           O
ATOM   1447  HH  TYR A  88      -6.418 -16.902  17.266  1.00  0.00
ATOM   1448  CD2 TYR A  88      -4.431 -14.703  14.825  1.00  0.00           C
ATOM   1449  HD2 TYR A  88      -3.673 -15.022  14.256  1.00  0.00
ATOM   1450  CE2 TYR A  88      -5.073 -15.598  15.669  1.00  0.00           C
ATOM   1451  HE2 TYR A  88      -4.773 -16.551  15.713  1.00  0.00
ATOM   1452  C   TYR A  88      -2.365 -11.917  15.553  1.00  0.00           C
ATOM   1453  O   TYR A  88      -1.407 -12.515  15.092  1.00  0.00           O
ATOM   1454  N   CYS A  89      -2.590 -11.771  16.858  1.00  0.00           N
ATOM   1455  HN  CYS A  89      -3.361 -11.207  17.154  1.00  0.00
ATOM   1456  CA  CYS A  89      -1.763 -12.397  17.859  1.00  0.00           C
ATOM   1457  HA  CYS A  89      -0.921 -12.684  17.401  1.00  0.00
ATOM   1458  CB  CYS A  89      -1.416 -11.408  18.957  1.00  0.00           C
ATOM   1459  HB1 CYS A  89      -2.289 -11.205  19.400  1.00  0.00
ATOM   1460  HB2 CYS A  89      -0.834 -11.922  19.587  1.00  0.00
ATOM   1461  SG  CYS A  89      -0.603  -9.921  18.335  1.00  0.00           S
ATOM   1462  HG1 CYS A  89      -0.399  -9.309  19.099  1.00  0.00
ATOM   1463  C   CYS A  89      -2.522 -13.590  18.436  1.00  0.00           C
ATOM   1464  O   CYS A  89      -3.540 -13.405  19.122  1.00  0.00           O
ATOM   1465  N   ASN A  90      -2.032 -14.798  18.155  1.00  0.00           N
ATOM   1466  HN  ASN A  90      -1.112 -14.876  17.771  1.00  0.00
ATOM   1467  CA  ASN A  90      -2.819 -16.021  18.399  1.00  0.00           C
ATOM   1468  HA  ASN A  90      -3.760 -15.758  18.184  1.00  0.00
ATOM   1469  CB  ASN A  90      -2.383 -17.156  17.473  1.00  0.00           C
ATOM   1470  HB1 ASN A  90      -2.997 -17.934  17.606  1.00  0.00
ATOM   1471  HB2 ASN A  90      -2.444 -16.841  16.526  1.00  0.00
ATOM   1472  CG  ASN A  90      -0.958 -17.612  17.740  1.00  0.00           C
ATOM   1473  OD1 ASN A  90      -0.260 -17.042  18.579  1.00  0.00           O
ATOM   1474  ND2 ASN A  90      -0.501 -18.621  17.022  1.00  0.00           N
ATOM   1475 1HD2 ASN A  90      -1.087 -19.052  16.336  1.00  0.00
ATOM   1476 2HD2 ASN A  90       0.431 -18.954  17.164  1.00  0.00
ATOM   1477  C   ASN A  90      -2.723 -16.478  19.870  1.00  0.00           C
ATOM   1478  O   ASN A  90      -3.271 -17.498  20.208  1.00  0.00           O
ATOM   1479  N   GLY A  91      -2.058 -15.698  20.732  1.00  0.00           N
ATOM   1480  HN  GLY A  91      -1.696 -14.826  20.404  1.00  0.00
ATOM   1481  CA  GLY A  91      -1.833 -16.067  22.150  1.00  0.00           C
ATOM   1482  HA1 GLY A  91      -2.028 -17.044  22.240  1.00  0.00
ATOM   1483  HA2 GLY A  91      -0.868 -15.900  22.354  1.00  0.00
ATOM   1484  C   GLY A  91      -2.690 -15.298  23.154  1.00  0.00           C
ATOM   1485  O   GLY A  91      -2.618 -15.565  24.349  1.00  0.00           O
ATOM   1486  N   GLY A  92      -3.502 -14.350  22.684  1.00  0.00           N
```

```
ATOM   1487  HN  GLY A  92      -3.563 -14.210  21.696  1.00  0.00
ATOM   1488  CA  GLY A  92      -4.312 -13.502  23.576  1.00  0.00           C
ATOM   1489  HA1 GLY A  92      -4.990 -13.022  23.019  1.00  0.00
ATOM   1490  HA2 GLY A  92      -4.778 -14.096  24.232  1.00  0.00
ATOM   1491  C   GLY A  92      -3.446 -12.487  24.323  1.00  0.00           C
ATOM   1492  O   GLY A  92      -2.341 -12.132  23.871  1.00  0.00           O
ATOM   1493  N   ASP A  93      -3.947 -12.015  25.469  1.00  0.00           N
ATOM   1494  HN  ASP A  93      -4.802 -12.397  25.819  1.00  0.00
ATOM   1495  CA  ASP A  93      -3.282 -10.961  26.221  1.00  0.00           C
ATOM   1496  HA  ASP A  93      -2.646 -10.515  25.592  1.00  0.00
ATOM   1497  CB  ASP A  93      -4.275  -9.875  26.667  1.00  0.00           C
ATOM   1498  HB1 ASP A  93      -3.751  -9.131  27.082  1.00  0.00
ATOM   1499  HB2 ASP A  93      -4.746  -9.537  25.852  1.00  0.00
ATOM   1500  CG  ASP A  93      -5.342 -10.302  27.672  1.00  0.00           C
ATOM   1501  OD1 ASP A  93      -5.092 -11.254  28.430  1.00  0.00           O
ATOM   1502  OD2 ASP A  93      -6.403  -9.635  27.710  1.00  0.00           O
ATOM   1503  C   ASP A  93      -2.475 -11.590  27.368  1.00  0.00           C
ATOM   1504  O   ASP A  93      -2.715 -12.735  27.794  1.00  0.00           O
ATOM   1505  N   LEU A  94      -1.491 -10.833  27.839  1.00  0.00           N
ATOM   1506  HN  LEU A  94      -1.344  -9.932  27.430  1.00  0.00
ATOM   1507  CA  LEU A  94      -0.619 -11.253  28.920  1.00  0.00           C
ATOM   1508  HA  LEU A  94      -0.202 -12.119  28.644  1.00  0.00
ATOM   1509  CB  LEU A  94       0.456 -10.185  29.130  1.00  0.00           C
ATOM   1510  HB1 LEU A  94       1.008 -10.136  28.297  1.00  0.00
ATOM   1511  HB2 LEU A  94      -0.005  -9.310  29.277  1.00  0.00
ATOM   1512  CG  LEU A  94       1.403 -10.417  30.304  1.00  0.00           C
ATOM   1513  HG  LEU A  94       0.854 -10.438  31.139  1.00  0.00
ATOM   1514  CD1 LEU A  94       2.112 -11.747  30.184  1.00  0.00           C
ATOM   1515 1HD1 LEU A  94       2.724 -11.870  30.965  1.00  0.00
ATOM   1516 2HD1 LEU A  94       1.437 -12.485  30.170  1.00  0.00
ATOM   1517 3HD1 LEU A  94       2.644 -11.766  29.338  1.00  0.00
ATOM   1518  CD2 LEU A  94       2.430  -9.289  30.398  1.00  0.00           C
ATOM   1519 1HD2 LEU A  94       3.041  -9.458  31.172  1.00  0.00
ATOM   1520 2HD2 LEU A  94       2.964  -9.253  29.553  1.00  0.00
ATOM   1521 3HD2 LEU A  94       1.957  -8.418  30.531  1.00  0.00
ATOM   1522  C   LEU A  94      -1.434 -11.471  30.196  1.00  0.00           C
ATOM   1523  O   LEU A  94      -1.082 -12.323  30.999  1.00  0.00           O
ATOM   1524  N   ALA A  95      -2.521 -10.700  30.355  1.00  0.00           N
ATOM   1525  HN  ALA A  95      -2.740 -10.034  29.642  1.00  0.00
ATOM   1526  CA  ALA A  95      -3.410 -10.788  31.534  1.00  0.00           C
ATOM   1527  HA  ALA A  95      -2.871 -10.519  32.332  1.00  0.00
ATOM   1528  CB  ALA A  95      -4.559  -9.821  31.408  1.00  0.00           C
ATOM   1529  HB1 ALA A  95      -5.147  -9.896  32.213  1.00  0.00
ATOM   1530  HB2 ALA A  95      -4.204  -8.889  31.340  1.00  0.00
ATOM   1531  HB3 ALA A  95      -5.088 -10.037  30.587  1.00  0.00
ATOM   1532  C   ALA A  95      -3.920 -12.227  31.699  1.00  0.00           C
ATOM   1533  O   ALA A  95      -3.856 -12.784  32.788  1.00  0.00           O
ATOM   1534  N   ASP A  96      -4.399 -12.808  30.592  1.00  0.00           N
ATOM   1535  HN  ASP A  96      -4.365 -12.289  29.738  1.00  0.00
ATOM   1536  CA  ASP A  96      -4.966 -14.147  30.555  1.00  0.00           C
ATOM   1537  HA  ASP A  96      -5.564 -14.222  31.353  1.00  0.00
ATOM   1538  CB  ASP A  96      -5.834 -14.340  29.310  1.00  0.00           C
ATOM   1539  HB1 ASP A  96      -5.313 -14.040  28.511  1.00  0.00
ATOM   1540  HB2 ASP A  96      -6.050 -15.312  29.221  1.00  0.00
ATOM   1541  CG  ASP A  96      -7.148 -13.548  29.366  1.00  0.00           C
ATOM   1542  OD1 ASP A  96      -7.778 -13.515  30.435  1.00  0.00           O
ATOM   1543  OD2 ASP A  96      -7.529 -12.951  28.352  1.00  0.00           O
ATOM   1544  C   ASP A  96      -3.846 -15.190  30.670  1.00  0.00           C
ATOM   1545  O   ASP A  96      -4.049 -16.248  31.245  1.00  0.00           O
ATOM   1546  N   TYR A  97      -2.665 -14.881  30.133  1.00  0.00           N
ATOM   1547  HN  TYR A  97      -2.568 -14.018  29.637  1.00  0.00
ATOM   1548  CA  TYR A  97      -1.508 -15.766  30.247  1.00  0.00           C
ATOM   1549  HA  TYR A  97      -1.796 -16.647  29.872  1.00  0.00
ATOM   1550  CB  TYR A  97      -0.333 -15.223  29.436  1.00  0.00           C
ATOM   1551  HB1 TYR A  97      -0.651 -15.033  28.507  1.00  0.00
ATOM   1552  HB2 TYR A  97      -0.021 -14.374  29.862  1.00  0.00
ATOM   1553  CG  TYR A  97       0.850 -16.149  29.336  1.00  0.00           C
ATOM   1554  CD1 TYR A  97       0.803 -17.277  28.535  1.00  0.00           C
ATOM   1555  HD1 TYR A  97      -0.033 -17.480  28.025  1.00  0.00
ATOM   1556  CE1 TYR A  97       1.888 -18.131  28.424  1.00  0.00           C
ATOM   1557  HE1 TYR A  97       1.829 -18.946  27.847  1.00  0.00
```

```
ATOM   1558  CZ  TYR A  97       3.059 -17.851  29.118  1.00  0.00           C
ATOM   1559  OH  TYR A  97       4.134 -18.690  28.989  1.00  0.00           O
ATOM   1560  HH  TYR A  97       4.891 -18.351  29.548  1.00  0.00
ATOM   1561  CD2 TYR A  97       2.027 -15.880  30.017  1.00  0.00           C
ATOM   1562  HD2 TYR A  97       2.082 -15.066  30.596  1.00  0.00
ATOM   1563  CE2 TYR A  97       3.129 -16.713  29.914  1.00  0.00           C
ATOM   1564  HE2 TYR A  97       3.970 -16.497  30.409  1.00  0.00
ATOM   1565  C   TYR A  97      -1.109 -15.916  31.715  1.00  0.00           C
ATOM   1566  O   TYR A  97      -0.782 -17.013  32.149  1.00  0.00           O
ATOM   1567  N   LEU A  98      -1.119 -14.795  32.459  1.00  0.00           N
ATOM   1568  HN  LEU A  98      -1.401 -13.937  32.029  1.00  0.00
ATOM   1569  CA  LEU A  98      -0.729 -14.778  33.893  1.00  0.00           C
ATOM   1570  HA  LEU A  98       0.147 -15.254  33.975  1.00  0.00
ATOM   1571  CB  LEU A  98      -0.570 -13.331  34.369  1.00  0.00           C
ATOM   1572  HB1 LEU A  98      -1.387 -12.825  34.092  1.00  0.00
ATOM   1573  HB2 LEU A  98      -0.506 -13.341  35.367  1.00  0.00
ATOM   1574  CG  LEU A  98       0.651 -12.585  33.827  1.00  0.00           C
ATOM   1575  HG  LEU A  98       0.530 -12.497  32.838  1.00  0.00
ATOM   1576  CD1 LEU A  98       0.763 -11.200  34.428  1.00  0.00           C
ATOM   1577 1HD1 LEU A  98       1.568 -10.739  34.055  1.00  0.00
ATOM   1578 2HD1 LEU A  98      -0.057 -10.672  34.204  1.00  0.00
ATOM   1579 3HD1 LEU A  98       0.851 -11.274  35.421  1.00  0.00
ATOM   1580  CD2 LEU A  98       1.931 -13.348  34.077  1.00  0.00           C
ATOM   1581 1HD2 LEU A  98       2.704 -12.831  33.710  1.00  0.00
ATOM   1582 2HD2 LEU A  98       2.054 -13.479  35.061  1.00  0.00
ATOM   1583 3HD2 LEU A  98       1.882 -14.239  33.626  1.00  0.00
ATOM   1584  C   LEU A  98      -1.784 -15.509  34.731  1.00  0.00           C
ATOM   1585  O   LEU A  98      -1.445 -16.170  35.697  1.00  0.00           O
ATOM   1586  N   GLN A  99      -3.058 -15.329  34.356  1.00  0.00           N
ATOM   1587  HN  GLN A  99      -3.244 -14.673  33.624  1.00  0.00
ATOM   1588  CA  GLN A  99      -4.187 -16.029  34.948  1.00  0.00           C
ATOM   1589  HA  GLN A  99      -4.221 -15.755  35.909  1.00  0.00
ATOM   1590  CB  GLN A  99      -5.480 -15.603  34.244  1.00  0.00           C
ATOM   1591  HB1 GLN A  99      -5.523 -14.604  34.233  1.00  0.00
ATOM   1592  HB2 GLN A  99      -5.460 -15.947  33.305  1.00  0.00
ATOM   1593  CG  GLN A  99      -6.740 -16.126  34.916  1.00  0.00           C
ATOM   1594  HG1 GLN A  99      -7.513 -16.004  34.294  1.00  0.00
ATOM   1595  HG2 GLN A  99      -6.622 -17.099  35.114  1.00  0.00
ATOM   1596  CD  GLN A  99      -7.039 -15.400  36.211  1.00  0.00           C
ATOM   1597  OE1 GLN A  99      -6.673 -14.233  36.376  1.00  0.00           O
ATOM   1598  NE2 GLN A  99      -7.680 -16.094  37.150  1.00  0.00           N
ATOM   1599 1HE2 GLN A  99      -7.938 -17.045  36.980  1.00  0.00
ATOM   1600 2HE2 GLN A  99      -7.903 -15.664  38.025  1.00  0.00
ATOM   1601  C   GLN A  99      -3.974 -17.551  34.853  1.00  0.00           C
ATOM   1602  O   GLN A  99      -4.300 -18.291  35.765  1.00  0.00           O
ATOM   1603  N   ALA A 100      -3.406 -17.993  33.734  1.00  0.00           N
ATOM   1604  HN  ALA A 100      -3.081 -17.320  33.069  1.00  0.00
ATOM   1605  CA  ALA A 100      -3.232 -19.414  33.425  1.00  0.00           C
ATOM   1606  HA  ALA A 100      -4.058 -19.867  33.760  1.00  0.00
ATOM   1607  CB  ALA A 100      -3.128 -19.615  31.930  1.00  0.00           C
ATOM   1608  HB1 ALA A 100      -3.010 -20.588  31.732  1.00  0.00
ATOM   1609  HB2 ALA A 100      -3.963 -19.285  31.490  1.00  0.00
ATOM   1610  HB3 ALA A 100      -2.342 -19.105  31.579  1.00  0.00
ATOM   1611  C   ALA A 100      -1.996 -19.994  34.138  1.00  0.00           C
ATOM   1612  O   ALA A 100      -2.019 -21.155  34.526  1.00  0.00           O
ATOM   1613  N   LYS A 101      -0.922 -19.199  34.260  1.00  0.00           N
ATOM   1614  HN  LYS A 101      -1.024 -18.226  34.053  1.00  0.00
ATOM   1615  CA  LYS A 101       0.402 -19.693  34.685  1.00  0.00           C
ATOM   1616  HA  LYS A 101       0.342 -20.689  34.622  1.00  0.00
ATOM   1617  CB  LYS A 101       1.489 -19.151  33.751  1.00  0.00           C
ATOM   1618  HB1 LYS A 101       1.456 -18.152  33.784  1.00  0.00
ATOM   1619  HB2 LYS A 101       2.375 -19.467  34.091  1.00  0.00
ATOM   1620  CG  LYS A 101       1.377 -19.569  32.293  1.00  0.00           C
ATOM   1621  HG1 LYS A 101       0.424 -19.476  32.004  1.00  0.00
ATOM   1622  HG2 LYS A 101       1.953 -18.967  31.741  1.00  0.00
ATOM   1623  CD  LYS A 101       1.808 -20.990  32.049  1.00  0.00           C
ATOM   1624  HD1 LYS A 101       2.490 -21.235  32.738  1.00  0.00
ATOM   1625  HD2 LYS A 101       1.008 -21.582  32.146  1.00  0.00
ATOM   1626  CE  LYS A 101       2.408 -21.210  30.676  1.00  0.00           C
ATOM   1627  HE1 LYS A 101       2.023 -20.547  30.034  1.00  0.00
ATOM   1628  HE2 LYS A 101       3.400 -21.091  30.723  1.00  0.00
```

| | | | | | | | | | | | |
|---|---|---|---|---|---|---|---|---|---|---|---|
| ATOM | 1629 | NZ | LYS | A | 101 | 2.121 | -22.573 | 30.173 | 1.00 | 0.00 | N |
| ATOM | 1630 | HZ1 | LYS | A | 101 | 2.529 | -22.687 | 29.267 | 1.00 | 0.00 | |
| ATOM | 1631 | HZ2 | LYS | A | 101 | 2.508 | -23.248 | 30.801 | 1.00 | 0.00 | |
| ATOM | 1632 | HZ3 | LYS | A | 101 | 1.132 | -22.704 | 30.112 | 1.00 | 0.00 | |
| ATOM | 1633 | C | LYS | A | 101 | 0.727 | -19.275 | 36.136 | 1.00 | 0.00 | C |
| ATOM | 1634 | O | LYS | A | 101 | 1.587 | -19.885 | 36.768 | 1.00 | 0.00 | O |
| ATOM | 1635 | N | GLY | A | 102 | 0.076 | -18.213 | 36.632 | 1.00 | 0.00 | N |
| ATOM | 1636 | HN | GLY | A | 102 | -0.575 | -17.740 | 36.038 | 1.00 | 0.00 | |
| ATOM | 1637 | CA | GLY | A | 102 | 0.263 | -17.708 | 37.989 | 1.00 | 0.00 | C |
| ATOM | 1638 | HA1 | GLY | A | 102 | -0.591 | -17.276 | 38.278 | 1.00 | 0.00 | |
| ATOM | 1639 | HA2 | GLY | A | 102 | 0.460 | -18.487 | 38.584 | 1.00 | 0.00 | |
| ATOM | 1640 | C | GLY | A | 102 | 1.396 | -16.703 | 38.073 | 1.00 | 0.00 | C |
| ATOM | 1641 | O | GLY | A | 102 | 1.162 | -15.496 | 38.030 | 1.00 | 0.00 | O |
| ATOM | 1642 | N | THR | A | 103 | 2.621 | -17.212 | 38.219 | 1.00 | 0.00 | N |
| ATOM | 1643 | HN | THR | A | 103 | 2.709 | -18.203 | 38.319 | 1.00 | 0.00 | |
| ATOM | 1644 | CA | THR | A | 103 | 3.831 | -16.416 | 38.241 | 1.00 | 0.00 | C |
| ATOM | 1645 | HA | THR | A | 103 | 3.548 | -15.482 | 38.021 | 1.00 | 0.00 | |
| ATOM | 1646 | CB | THR | A | 103 | 4.496 | -16.409 | 39.627 | 1.00 | 0.00 | C |
| ATOM | 1647 | HB | THR | A | 103 | 5.398 | -15.981 | 39.567 | 1.00 | 0.00 | |
| ATOM | 1648 | OG1 | THR | A | 103 | 4.671 | -17.758 | 40.053 | 1.00 | 0.00 | O |
| ATOM | 1649 | HG1 | THR | A | 103 | 5.103 | -17.769 | 40.955 | 1.00 | 0.00 | |
| ATOM | 1650 | CG2 | THR | A | 103 | 3.704 | -15.662 | 40.673 | 1.00 | 0.00 | C |
| ATOM | 1651 | 1HG2 | THR | A | 103 | 4.190 | -15.696 | 41.546 | 1.00 | 0.00 | |
| ATOM | 1652 | 2HG2 | THR | A | 103 | 3.593 | -14.709 | 40.390 | 1.00 | 0.00 | |
| ATOM | 1653 | 3HG2 | THR | A | 103 | 2.804 | -16.086 | 40.776 | 1.00 | 0.00 | |
| ATOM | 1654 | C | THR | A | 103 | 4.786 | -16.980 | 37.196 | 1.00 | 0.00 | C |
| ATOM | 1655 | O | THR | A | 103 | 4.608 | -18.104 | 36.754 | 1.00 | 0.00 | O |
| ATOM | 1656 | N | LEU | A | 104 | 5.811 | -16.197 | 36.841 | 1.00 | 0.00 | N |
| ATOM | 1657 | HN | LEU | A | 104 | 5.921 | -15.319 | 37.306 | 1.00 | 0.00 | |
| ATOM | 1658 | CA | LEU | A | 104 | 6.774 | -16.559 | 35.812 | 1.00 | 0.00 | C |
| ATOM | 1659 | HA | LEU | A | 104 | 6.550 | -17.485 | 35.508 | 1.00 | 0.00 | |
| ATOM | 1660 | CB | LEU | A | 104 | 6.654 | -15.568 | 34.652 | 1.00 | 0.00 | C |
| ATOM | 1661 | HB1 | LEU | A | 104 | 6.994 | -14.682 | 34.968 | 1.00 | 0.00 | |
| ATOM | 1662 | HB2 | LEU | A | 104 | 7.230 | -15.899 | 33.904 | 1.00 | 0.00 | |
| ATOM | 1663 | CG | LEU | A | 104 | 5.246 | -15.358 | 34.100 | 1.00 | 0.00 | C |
| ATOM | 1664 | HG | LEU | A | 104 | 4.672 | -15.003 | 34.838 | 1.00 | 0.00 | |
| ATOM | 1665 | CD1 | LEU | A | 104 | 5.269 | -14.341 | 32.967 | 1.00 | 0.00 | C |
| ATOM | 1666 | 1HD1 | LEU | A | 104 | 4.342 | -14.213 | 32.615 | 1.00 | 0.00 | |
| ATOM | 1667 | 2HD1 | LEU | A | 104 | 5.619 | -13.469 | 33.309 | 1.00 | 0.00 | |
| ATOM | 1668 | 3HD1 | LEU | A | 104 | 5.861 | -14.673 | 32.233 | 1.00 | 0.00 | |
| ATOM | 1669 | CD2 | LEU | A | 104 | 4.632 | -16.670 | 33.632 | 1.00 | 0.00 | C |
| ATOM | 1670 | 1HD2 | LEU | A | 104 | 3.713 | -16.500 | 33.277 | 1.00 | 0.00 | |
| ATOM | 1671 | 2HD2 | LEU | A | 104 | 5.200 | -17.065 | 32.910 | 1.00 | 0.00 | |
| ATOM | 1672 | 3HD2 | LEU | A | 104 | 4.580 | -17.307 | 34.401 | 1.00 | 0.00 | |
| ATOM | 1673 | C | LEU | A | 104 | 8.190 | -16.534 | 36.395 | 1.00 | 0.00 | C |
| ATOM | 1674 | O | LEU | A | 104 | 8.519 | -15.669 | 37.193 | 1.00 | 0.00 | O |
| ATOM | 1675 | N | SER | A | 105 | 9.021 | -17.481 | 35.948 | 1.00 | 0.00 | N |
| ATOM | 1676 | HN | SER | A | 105 | 8.646 | -18.217 | 35.384 | 1.00 | 0.00 | |
| ATOM | 1677 | CA | SER | A | 105 | 10.443 | -17.491 | 36.244 | 1.00 | 0.00 | C |
| ATOM | 1678 | HA | SER | A | 105 | 10.521 | -17.592 | 37.236 | 1.00 | 0.00 | |
| ATOM | 1679 | CB | SER | A | 105 | 11.120 | -18.666 | 35.598 | 1.00 | 0.00 | C |
| ATOM | 1680 | HB1 | SER | A | 105 | 12.086 | -18.692 | 35.855 | 1.00 | 0.00 | |
| ATOM | 1681 | HB2 | SER | A | 105 | 10.677 | -19.517 | 35.879 | 1.00 | 0.00 | |
| ATOM | 1682 | OG | SER | A | 105 | 11.040 | -18.571 | 34.174 | 1.00 | 0.00 | O |
| ATOM | 1683 | HG1 | SER | A | 105 | 11.495 | -19.360 | 33.761 | 1.00 | 0.00 | |
| ATOM | 1684 | C | SER | A | 105 | 11.077 | -16.181 | 35.767 | 1.00 | 0.00 | C |
| ATOM | 1685 | O | SER | A | 105 | 10.603 | -15.551 | 34.828 | 1.00 | 0.00 | O |
| ATOM | 1686 | N | GLU | A | 106 | 12.184 | -15.811 | 36.406 | 1.00 | 0.00 | N |
| ATOM | 1687 | HN | GLU | A | 106 | 12.484 | -16.349 | 37.194 | 1.00 | 0.00 | |
| ATOM | 1688 | CA | GLU | A | 106 | 12.969 | -14.671 | 36.014 | 1.00 | 0.00 | C |
| ATOM | 1689 | HA | GLU | A | 106 | 12.357 | -13.881 | 36.049 | 1.00 | 0.00 | |
| ATOM | 1690 | CB | GLU | A | 106 | 14.112 | -14.440 | 37.000 | 1.00 | 0.00 | C |
| ATOM | 1691 | HB1 | GLU | A | 106 | 14.671 | -15.268 | 37.039 | 1.00 | 0.00 | |
| ATOM | 1692 | HB2 | GLU | A | 106 | 14.668 | -13.678 | 36.669 | 1.00 | 0.00 | |
| ATOM | 1693 | CG | GLU | A | 106 | 13.646 | -14.114 | 38.407 | 1.00 | 0.00 | C |
| ATOM | 1694 | HG1 | GLU | A | 106 | 12.900 | -13.451 | 38.340 | 1.00 | 0.00 | |
| ATOM | 1695 | HG2 | GLU | A | 106 | 13.306 | -14.957 | 38.823 | 1.00 | 0.00 | |
| ATOM | 1696 | CD | GLU | A | 106 | 14.713 | -13.533 | 39.322 | 1.00 | 0.00 | C |
| ATOM | 1697 | OE1 | GLU | A | 106 | 15.688 | -12.916 | 38.794 | 1.00 | 0.00 | O |
| ATOM | 1698 | OE2 | GLU | A | 106 | 14.561 | -13.676 | 40.564 | 1.00 | 0.00 | O |
| ATOM | 1699 | C | GLU | A | 106 | 13.477 | -14.879 | 34.586 | 1.00 | 0.00 | C |

```
ATOM   1700  O   GLU A 106      13.559 -13.924  33.835  1.00  0.00           O
ATOM   1701  N   ASP A 107      13.775 -16.131  34.208  1.00  0.00           N
ATOM   1702  HN  ASP A 107      13.705 -16.869  34.879  1.00  0.00
ATOM   1703  CA  ASP A 107      14.205 -16.456  32.828  1.00  0.00           C
ATOM   1704  HA  ASP A 107      15.055 -15.954  32.668  1.00  0.00
ATOM   1705  CB  ASP A 107      14.525 -17.946  32.652  1.00  0.00           C
ATOM   1706  HB1 ASP A 107      13.816 -18.478  33.115  1.00  0.00
ATOM   1707  HB2 ASP A 107      14.516 -18.157  31.675  1.00  0.00
ATOM   1708  CG  ASP A 107      15.875 -18.371  33.214  1.00  0.00           C
ATOM   1709  OD1 ASP A 107      16.701 -17.483  33.523  1.00  0.00           O
ATOM   1710  OD2 ASP A 107      16.112 -19.596  33.309  1.00  0.00           O
ATOM   1711  C   ASP A 107      13.121 -15.997  31.849  1.00  0.00           C
ATOM   1712  O   ASP A 107      13.408 -15.274  30.921  1.00  0.00           O
ATOM   1713  N   THR A 108      11.876 -16.411  32.104  1.00  0.00           N
ATOM   1714  HN  THR A 108      11.719 -16.982  32.910  1.00  0.00
ATOM   1715  CA  THR A 108      10.732 -16.068  31.259  1.00  0.00           C
ATOM   1716  HA  THR A 108      10.965 -16.422  30.353  1.00  0.00
ATOM   1717  CB  THR A 108       9.447 -16.744  31.757  1.00  0.00           C
ATOM   1718  HB  THR A 108       9.251 -16.464  32.697  1.00  0.00
ATOM   1719  OG1 THR A 108       9.645 -18.159  31.768  1.00  0.00           O
ATOM   1720  HG1 THR A 108       8.813 -18.610  32.091  1.00  0.00
ATOM   1721  CG2 THR A 108       8.241 -16.401  30.913  1.00  0.00           C
ATOM   1722 1HG2 THR A 108       7.435 -16.865  31.279  1.00  0.00
ATOM   1723 2HG2 THR A 108       8.093 -15.412  30.930  1.00  0.00
ATOM   1724 3HG2 THR A 108       8.398 -16.698  29.971  1.00  0.00
ATOM   1725  C   THR A 108      10.538 -14.540  31.217  1.00  0.00           C
ATOM   1726  O   THR A 108      10.254 -13.982  30.161  1.00  0.00           O
ATOM   1727  N   ILE A 109      10.653 -13.882  32.385  1.00  0.00           N
ATOM   1728  HN  ILE A 109      10.931 -14.393  33.198  1.00  0.00
ATOM   1729  CA  ILE A 109      10.389 -12.452  32.513  1.00  0.00           C
ATOM   1730  HA  ILE A 109       9.483 -12.284  32.125  1.00  0.00
ATOM   1731  CB  ILE A 109      10.336 -12.028  33.995  1.00  0.00           C
ATOM   1732  HB  ILE A 109      11.106 -12.460  34.465  1.00  0.00
ATOM   1733  CG2 ILE A 109      10.507 -10.520  34.159  1.00  0.00           C
ATOM   1734 1HG2 ILE A 109      10.467 -10.284  35.130  1.00  0.00
ATOM   1735 2HG2 ILE A 109      11.391 -10.242  33.784  1.00  0.00
ATOM   1736 3HG2 ILE A 109       9.774 -10.046  33.670  1.00  0.00
ATOM   1737  CG1 ILE A 109       9.040 -12.511  34.658  1.00  0.00           C
ATOM   1738 1HG1 ILE A 109       8.285 -11.941  34.334  1.00  0.00
ATOM   1739 2HG1 ILE A 109       8.880 -13.461  34.390  1.00  0.00
ATOM   1740  CD  ILE A 109       9.068 -12.444  36.178  1.00  0.00           C
ATOM   1741  HD1 ILE A 109       8.196 -12.772  36.542  1.00  0.00
ATOM   1742  HD2 ILE A 109       9.811 -13.018  36.523  1.00  0.00
ATOM   1743  HD3 ILE A 109       9.216 -11.498  36.467  1.00  0.00
ATOM   1744  C   ILE A 109      11.431 -11.679  31.696  1.00  0.00           C
ATOM   1745  O   ILE A 109      11.102 -10.675  31.072  1.00  0.00           O
ATOM   1746  N   ARG A 110      12.673 -12.170  31.683  1.00  0.00           N
ATOM   1747  HN  ARG A 110      12.858 -13.005  32.202  1.00  0.00
ATOM   1748  CA  ARG A 110      13.787 -11.545  30.943  1.00  0.00           C
ATOM   1749  HA  ARG A 110      13.860 -10.603  31.270  1.00  0.00
ATOM   1750  CB  ARG A 110      15.101 -12.279  31.232  1.00  0.00           C
ATOM   1751  HB1 ARG A 110      15.194 -12.382  32.222  1.00  0.00
ATOM   1752  HB2 ARG A 110      15.059 -13.182  30.805  1.00  0.00
ATOM   1753  CG  ARG A 110      16.342 -11.567  30.712  1.00  0.00           C
ATOM   1754  HG1 ARG A 110      16.238 -11.412  29.730  1.00  0.00
ATOM   1755  HG2 ARG A 110      16.431 -10.688  31.181  1.00  0.00
ATOM   1756  CD  ARG A 110      17.605 -12.384  30.951  1.00  0.00           C
ATOM   1757  HD1 ARG A 110      17.453 -13.327  30.655  1.00  0.00
ATOM   1758  HD2 ARG A 110      18.362 -11.989  30.430  1.00  0.00
ATOM   1759  NE  ARG A 110      18.003 -12.418  32.358  1.00  0.00           N
ATOM   1760  HE  ARG A 110      18.434 -11.598  32.734  1.00  0.00
ATOM   1761  CZ  ARG A 110      17.836 -13.459  33.182  1.00  0.00           C
ATOM   1762  NH1 ARG A 110      17.267 -14.573  32.751  1.00  0.00           N
ATOM   1763 1HH1 ARG A 110      16.958 -14.642  31.803  1.00  0.00
ATOM   1764 2HH1 ARG A 110      17.146 -15.346  33.374  1.00  0.00
ATOM   1765  NH2 ARG A 110      18.209 -13.362  34.447  1.00  0.00           N
ATOM   1766 1HH2 ARG A 110      18.614 -12.513  34.785  1.00  0.00
ATOM   1767 2HH2 ARG A 110      18.086 -14.138  35.066  1.00  0.00
ATOM   1768  C   ARG A 110      13.481 -11.555  29.439  1.00  0.00           C
ATOM   1769  O   ARG A 110      13.745 -10.580  28.731  1.00  0.00           O
ATOM   1770  N   VAL A 111      12.918 -12.669  28.963  1.00  0.00           N
```

```
ATOM   1771  HN  VAL A 111      12.718 -13.410  29.603  1.00  0.00
ATOM   1772  CA  VAL A 111      12.582 -12.858  27.563  1.00  0.00           C
ATOM   1773  HA  VAL A 111      13.417 -12.673  27.045  1.00  0.00
ATOM   1774  CB  VAL A 111      12.151 -14.313  27.293  1.00  0.00           C
ATOM   1775  HB  VAL A 111      11.512 -14.576  28.015  1.00  0.00
ATOM   1776  CG1 VAL A 111      11.403 -14.462  25.976  1.00  0.00           C
ATOM   1777 1HG1 VAL A 111      11.145 -15.419  25.846  1.00  0.00
ATOM   1778 2HG1 VAL A 111      10.580 -13.895  25.994  1.00  0.00
ATOM   1779 3HG1 VAL A 111      11.993 -14.170  25.223  1.00  0.00
ATOM   1780  CG2 VAL A 111      13.338 -15.261  27.357  1.00  0.00           C
ATOM   1781 1HG2 VAL A 111      13.029 -16.195  27.178  1.00  0.00
ATOM   1782 2HG2 VAL A 111      14.014 -14.996  26.669  1.00  0.00
ATOM   1783 3HG2 VAL A 111      13.752 -15.215  28.266  1.00  0.00
ATOM   1784  C   VAL A 111      11.488 -11.856  27.170  1.00  0.00           C
ATOM   1785  O   VAL A 111      11.602 -11.201  26.145  1.00  0.00           O
ATOM   1786  N   PHE A 112      10.435 -11.766  27.990  1.00  0.00           N
ATOM   1787  HN  PHE A 112      10.413 -12.351  28.801  1.00  0.00
ATOM   1788  CA  PHE A 112       9.319 -10.860  27.761  1.00  0.00           C
ATOM   1789  HA  PHE A 112       8.970 -11.074  26.849  1.00  0.00
ATOM   1790  CB  PHE A 112       8.209 -11.104  28.787  1.00  0.00           C
ATOM   1791  HB1 PHE A 112       8.628 -11.203  29.690  1.00  0.00
ATOM   1792  HB2 PHE A 112       7.599 -10.312  28.788  1.00  0.00
ATOM   1793  CG  PHE A 112       7.369 -12.328  28.536  1.00  0.00           C
ATOM   1794  CD1 PHE A 112       7.756 -13.297  27.615  1.00  0.00           C
ATOM   1795  HD1 PHE A 112       8.609 -13.180  27.107  1.00  0.00
ATOM   1796  CE1 PHE A 112       6.977 -14.422  27.391  1.00  0.00           C
ATOM   1797  HE1 PHE A 112       7.262 -15.100  26.714  1.00  0.00
ATOM   1798  CZ  PHE A 112       5.812 -14.606  28.100  1.00  0.00           C
ATOM   1799  HZ  PHE A 112       5.252 -15.420  27.944  1.00  0.00
ATOM   1800  CD2 PHE A 112       6.201 -12.538  29.248  1.00  0.00           C
ATOM   1801  HD2 PHE A 112       5.914 -11.869  29.933  1.00  0.00
ATOM   1802  CE2 PHE A 112       5.421 -13.666  29.024  1.00  0.00           C
ATOM   1803  HE2 PHE A 112       4.572 -13.795  29.537  1.00  0.00
ATOM   1804  C   PHE A 112       9.809  -9.406  27.803  1.00  0.00           C
ATOM   1805  O   PHE A 112       9.396  -8.567  26.999  1.00  0.00           O
ATOM   1806  N   LEU A 113      10.691  -9.122  28.754  1.00  0.00           N
ATOM   1807  HN  LEU A 113      11.090  -9.870  29.284  1.00  0.00
ATOM   1808  CA  LEU A 113      11.090  -7.767  29.045  1.00  0.00           C
ATOM   1809  HA  LEU A 113      10.269  -7.197  29.010  1.00  0.00
ATOM   1810  CB  LEU A 113      11.701  -7.741  30.442  1.00  0.00           C
ATOM   1811  HB1 LEU A 113      11.052  -8.167  31.072  1.00  0.00
ATOM   1812  HB2 LEU A 113      12.545  -8.277  30.421  1.00  0.00
ATOM   1813  CG  LEU A 113      12.039  -6.367  30.985  1.00  0.00           C
ATOM   1814  HG  LEU A 113      12.664  -5.910  30.353  1.00  0.00
ATOM   1815  CD1 LEU A 113      10.782  -5.514  31.097  1.00  0.00           C
ATOM   1816 1HD1 LEU A 113      11.022  -4.612  31.456  1.00  0.00
ATOM   1817 2HD1 LEU A 113      10.366  -5.413  30.193  1.00  0.00
ATOM   1818 3HD1 LEU A 113      10.133  -5.957  31.715  1.00  0.00
ATOM   1819  CD2 LEU A 113      12.729  -6.508  32.331  1.00  0.00           C
ATOM   1820 1HD2 LEU A 113      12.951  -5.601  32.688  1.00  0.00
ATOM   1821 2HD2 LEU A 113      12.120  -6.978  32.970  1.00  0.00
ATOM   1822 3HD2 LEU A 113      13.570  -7.038  32.221  1.00  0.00
ATOM   1823  C   LEU A 113      12.085  -7.277  27.991  1.00  0.00           C
ATOM   1824  O   LEU A 113      12.113  -6.108  27.653  1.00  0.00           O
ATOM   1825  N   HIS A 114      12.912  -8.188  27.488  1.00  0.00           N
ATOM   1826  HN  HIS A 114      12.915  -9.111  27.873  1.00  0.00
ATOM   1827  CA  HIS A 114      13.806  -7.875  26.398  1.00  0.00           C
ATOM   1828  HA  HIS A 114      14.413  -7.150  26.723  1.00  0.00
ATOM   1829  CB  HIS A 114      14.672  -9.089  26.048  1.00  0.00           C
ATOM   1830  HB1 HIS A 114      15.152  -9.396  26.870  1.00  0.00
ATOM   1831  HB2 HIS A 114      14.084  -9.824  25.711  1.00  0.00
ATOM   1832  ND1 HIS A 114      15.491  -9.128  23.664  1.00  0.00           N
ATOM   1833  CG  HIS A 114      15.693  -8.805  25.001  1.00  0.00           C
ATOM   1834  CE1 HIS A 114      16.552  -8.768  22.965  1.00  0.00           C
ATOM   1835  HE1 HIS A 114      16.684  -8.908  21.984  1.00  0.00
ATOM   1836  NE2 HIS A 114      17.425  -8.180  23.805  1.00  0.00           N
ATOM   1837  HE2 HIS A 114      18.313  -7.798  23.548  1.00  0.00
ATOM   1838  CD2 HIS A 114      16.896  -8.191  25.070  1.00  0.00           C
ATOM   1839  HD2 HIS A 114      17.322  -7.814  25.893  1.00  0.00
ATOM   1840  C   HIS A 114      12.984  -7.364  25.202  1.00  0.00           C
ATOM   1841  O   HIS A 114      13.364  -6.408  24.531  1.00  0.00           O
```

```
ATOM   1842  N   GLN A 115      11.836  -7.994  24.951  1.00  0.00           N
ATOM   1843  HN  GLN A 115      11.541  -8.727  25.564  1.00  0.00
ATOM   1844  CA  GLN A 115      11.003  -7.648  23.816  1.00  0.00           C
ATOM   1845  HA  GLN A 115      11.612  -7.531  23.031  1.00  0.00
ATOM   1846  CB  GLN A 115      10.045  -8.794  23.511  1.00  0.00           C
ATOM   1847  HB1 GLN A 115       9.530  -9.009  24.341  1.00  0.00
ATOM   1848  HB2 GLN A 115       9.414  -8.500  22.794  1.00  0.00
ATOM   1849  CG  GLN A 115      10.760 -10.056  23.042  1.00  0.00           C
ATOM   1850  HG1 GLN A 115      11.267  -9.855  22.204  1.00  0.00
ATOM   1851  HG2 GLN A 115      11.396 -10.357  23.752  1.00  0.00
ATOM   1852  CD  GLN A 115       9.788 -11.170  22.761  1.00  0.00           C
ATOM   1853  OE1 GLN A 115       9.003 -11.102  21.842  1.00  0.00           O
ATOM   1854  NE2 GLN A 115       9.819 -12.200  23.568  1.00  0.00           N
ATOM   1855 1HE2 GLN A 115      10.468 -12.220  24.328  1.00  0.00
ATOM   1856 2HE2 GLN A 115       9.193 -12.967  23.424  1.00  0.00
ATOM   1857  C   GLN A 115      10.276  -6.320  24.084  1.00  0.00           C
ATOM   1858  O   GLN A 115      10.226  -5.461  23.227  1.00  0.00           O
ATOM   1859  N   ILE A 116       9.726  -6.172  25.287  1.00  0.00           N
ATOM   1860  HN  ILE A 116       9.773  -6.934  25.933  1.00  0.00
ATOM   1861  CA  ILE A 116       9.059  -4.947  25.700  1.00  0.00           C
ATOM   1862  HA  ILE A 116       8.276  -4.833  25.089  1.00  0.00
ATOM   1863  CB  ILE A 116       8.541  -5.062  27.143  1.00  0.00           C
ATOM   1864  HB  ILE A 116       9.269  -5.435  27.718  1.00  0.00
ATOM   1865  CG2 ILE A 116       8.179  -3.692  27.679  1.00  0.00           C
ATOM   1866 1HG2 ILE A 116       7.844  -3.779  28.617  1.00  0.00
ATOM   1867 2HG2 ILE A 116       8.988  -3.105  27.667  1.00  0.00
ATOM   1868 3HG2 ILE A 116       7.466  -3.288  27.106  1.00  0.00
ATOM   1869  CG1 ILE A 116       7.359  -6.021  27.239  1.00  0.00           C
ATOM   1870 1HG1 ILE A 116       6.543  -5.565  26.882  1.00  0.00
ATOM   1871 2HG1 ILE A 116       7.555  -6.832  26.688  1.00  0.00
ATOM   1872  CD  ILE A 116       7.065  -6.470  28.636  1.00  0.00           C
ATOM   1873  HD1 ILE A 116       6.283  -7.093  28.628  1.00  0.00
ATOM   1874  HD2 ILE A 116       7.864  -6.942  29.010  1.00  0.00
ATOM   1875  HD3 ILE A 116       6.852  -5.675  29.204  1.00  0.00
ATOM   1876  C   ILE A 116      10.025  -3.769  25.540  1.00  0.00           C
ATOM   1877  O   ILE A 116       9.658  -2.734  25.009  1.00  0.00           O
ATOM   1878  N   ALA A 117      11.259  -3.960  25.997  1.00  0.00           N
ATOM   1879  HN  ALA A 117      11.477  -4.846  26.407  1.00  0.00
ATOM   1880  CA  ALA A 117      12.307  -2.961  25.937  1.00  0.00           C
ATOM   1881  HA  ALA A 117      11.960  -2.164  26.432  1.00  0.00
ATOM   1882  CB  ALA A 117      13.550  -3.493  26.609  1.00  0.00           C
ATOM   1883  HB1 ALA A 117      14.273  -2.803  26.567  1.00  0.00
ATOM   1884  HB2 ALA A 117      13.347  -3.706  27.565  1.00  0.00
ATOM   1885  HB3 ALA A 117      13.853  -4.322  26.140  1.00  0.00
ATOM   1886  C   ALA A 117      12.609  -2.572  24.495  1.00  0.00           C
ATOM   1887  O   ALA A 117      12.846  -1.398  24.187  1.00  0.00           O
ATOM   1888  N   ALA A 118      12.636  -3.567  23.606  1.00  0.00           N
ATOM   1889  HN  ALA A 118      12.482  -4.504  23.919  1.00  0.00
ATOM   1890  CA  ALA A 118      12.884  -3.324  22.191  1.00  0.00           C
ATOM   1891  HA  ALA A 118      13.771  -2.870  22.108  1.00  0.00
ATOM   1892  CB  ALA A 118      12.977  -4.631  21.465  1.00  0.00           C
ATOM   1893  HB1 ALA A 118      13.147  -4.462  20.494  1.00  0.00
ATOM   1894  HB2 ALA A 118      13.728  -5.171  21.846  1.00  0.00
ATOM   1895  HB3 ALA A 118      12.118  -5.133  21.570  1.00  0.00
ATOM   1896  C   ALA A 118      11.782  -2.418  21.619  1.00  0.00           C
ATOM   1897  O   ALA A 118      12.074  -1.507  20.832  1.00  0.00           O
ATOM   1898  N   ALA A 119      10.526  -2.642  22.052  1.00  0.00           N
ATOM   1899  HN  ALA A 119      10.371  -3.406  22.678  1.00  0.00
ATOM   1900  CA  ALA A 119       9.375  -1.827  21.656  1.00  0.00           C
ATOM   1901  HA  ALA A 119       9.358  -1.810  20.656  1.00  0.00
ATOM   1902  CB  ALA A 119       8.094  -2.457  22.152  1.00  0.00           C
ATOM   1903  HB1 ALA A 119       7.316  -1.892  21.876  1.00  0.00
ATOM   1904  HB2 ALA A 119       7.998  -3.371  21.759  1.00  0.00
ATOM   1905  HB3 ALA A 119       8.120  -2.524  23.149  1.00  0.00
ATOM   1906  C   ALA A 119       9.530  -0.407  22.198  1.00  0.00           C
ATOM   1907  O   ALA A 119       9.251   0.561  21.509  1.00  0.00           O
ATOM   1908  N   MET A 120       9.961  -0.305  23.458  1.00  0.00           N
ATOM   1909  HN  MET A 120      10.202  -1.139  23.955  1.00  0.00
ATOM   1910  CA  MET A 120      10.091   0.966  24.126  1.00  0.00           C
ATOM   1911  HA  MET A 120       9.194   1.403  24.057  1.00  0.00
ATOM   1912  CB  MET A 120      10.444   0.763  25.600  1.00  0.00           C
```

```
ATOM   1913  HB1 MET A 120      11.248   0.171  25.652  1.00  0.00
ATOM   1914  HB2 MET A 120      10.661   1.655  25.997  1.00  0.00
ATOM   1915  CG  MET A 120       9.331   0.130  26.428  1.00  0.00           C
ATOM   1916  HG1 MET A 120       9.444  -0.854  26.289  1.00  0.00
ATOM   1917  HG2 MET A 120       9.560   0.361  27.374  1.00  0.00
ATOM   1918  SD  MET A 120       7.675   0.694  25.974  1.00  0.00           S
ATOM   1919  CE  MET A 120       7.833   2.424  26.419  1.00  0.00           C
ATOM   1920  HE1 MET A 120       6.974   2.898  26.224  1.00  0.00
ATOM   1921  HE2 MET A 120       8.042   2.500  27.394  1.00  0.00
ATOM   1922  HE3 MET A 120       8.571   2.839  25.887  1.00  0.00
ATOM   1923  C   MET A 120      11.166   1.808  23.420  1.00  0.00           C
ATOM   1924  O   MET A 120      11.025   3.012  23.286  1.00  0.00           O
ATOM   1925  N   ARG A 121      12.241   1.162  22.966  1.00  0.00           N
ATOM   1926  HN  ARG A 121      12.303   0.173  23.098  1.00  0.00
ATOM   1927  CA  ARG A 121      13.345   1.861  22.273  1.00  0.00           C
ATOM   1928  HA  ARG A 121      13.756   2.511  22.912  1.00  0.00
ATOM   1929  CB  ARG A 121      14.452   0.876  21.872  1.00  0.00           C
ATOM   1930  HB1 ARG A 121      14.464   0.127  22.534  1.00  0.00
ATOM   1931  HB2 ARG A 121      14.240   0.515  20.964  1.00  0.00
ATOM   1932  CG  ARG A 121      15.848   1.496  21.827  1.00  0.00           C
ATOM   1933  HG1 ARG A 121      15.877   2.139  21.061  1.00  0.00
ATOM   1934  HG2 ARG A 121      15.995   1.991  22.683  1.00  0.00
ATOM   1935  CD  ARG A 121      16.995   0.499  21.653  1.00  0.00           C
ATOM   1936  HD1 ARG A 121      17.044   0.229  20.691  1.00  0.00
ATOM   1937  HD2 ARG A 121      17.851   0.943  21.918  1.00  0.00
ATOM   1938  NE  ARG A 121      16.884  -0.740  22.448  1.00  0.00           N
ATOM   1939  HE  ARG A 121      16.074  -1.307  22.299  1.00  0.00
ATOM   1940  CZ  ARG A 121      17.778  -1.173  23.355  1.00  0.00           C
ATOM   1941  NH1 ARG A 121      18.885  -0.479  23.578  1.00  0.00           N
ATOM   1942 1HH1 ARG A 121      19.056   0.366  23.072  1.00  0.00
ATOM   1943 2HH1 ARG A 121      19.550  -0.801  24.252  1.00  0.00
ATOM   1944  NH2 ARG A 121      17.553  -2.293  24.040  1.00  0.00           N
ATOM   1945 1HH2 ARG A 121      16.716  -2.817  23.881  1.00  0.00
ATOM   1946 2HH2 ARG A 121      18.221  -2.610  24.713  1.00  0.00
ATOM   1947  C   ARG A 121      12.766   2.635  21.086  1.00  0.00           C
ATOM   1948  O   ARG A 121      13.099   3.797  20.876  1.00  0.00           O
ATOM   1949  N   ILE A 122      11.833   2.004  20.365  1.00  0.00           N
ATOM   1950  HN  ILE A 122      11.562   1.078  20.627  1.00  0.00
ATOM   1951  CA  ILE A 122      11.197   2.627  19.205  1.00  0.00           C
ATOM   1952  HA  ILE A 122      11.941   2.986  18.642  1.00  0.00
ATOM   1953  CB  ILE A 122      10.407   1.598  18.358  1.00  0.00           C
ATOM   1954  HB  ILE A 122       9.625   1.293  18.901  1.00  0.00
ATOM   1955  CG2 ILE A 122       9.876   2.243  17.091  1.00  0.00           C
ATOM   1956 1HG2 ILE A 122       9.370   1.565  16.558  1.00  0.00
ATOM   1957 2HG2 ILE A 122       9.268   2.999  17.332  1.00  0.00
ATOM   1958 3HG2 ILE A 122      10.641   2.590  16.548  1.00  0.00
ATOM   1959  CG1 ILE A 122      11.220   0.342  18.034  1.00  0.00           C
ATOM   1960 1HG1 ILE A 122      11.459  -0.111  18.893  1.00  0.00
ATOM   1961 2HG1 ILE A 122      10.652  -0.269  17.482  1.00  0.00
ATOM   1962  CD  ILE A 122      12.503   0.609  17.270  1.00  0.00           C
ATOM   1963  HD1 ILE A 122      12.974  -0.256  17.097  1.00  0.00
ATOM   1964  HD2 ILE A 122      12.287   1.050  16.399  1.00  0.00
ATOM   1965  HD3 ILE A 122      13.094   1.209  17.810  1.00  0.00
ATOM   1966  C   ILE A 122      10.297   3.770  19.690  1.00  0.00           C
ATOM   1967  O   ILE A 122      10.352   4.852  19.161  1.00  0.00           O
ATOM   1968  N   LEU A 123       9.459   3.509  20.697  1.00  0.00           N
ATOM   1969  HN  LEU A 123       9.440   2.593  21.097  1.00  0.00
ATOM   1970  CA  LEU A 123       8.571   4.540  21.221  1.00  0.00           C
ATOM   1971  HA  LEU A 123       7.931   4.763  20.486  1.00  0.00
ATOM   1972  CB  LEU A 123       7.798   4.014  22.435  1.00  0.00           C
ATOM   1973  HB1 LEU A 123       8.467   3.693  23.105  1.00  0.00
ATOM   1974  HB2 LEU A 123       7.281   4.778  22.820  1.00  0.00
ATOM   1975  CG  LEU A 123       6.815   2.874  22.186  1.00  0.00           C
ATOM   1976  HG  LEU A 123       7.307   2.011  22.303  1.00  0.00
ATOM   1977  CD1 LEU A 123       5.669   2.953  23.187  1.00  0.00           C
ATOM   1978 1HD1 LEU A 123       5.028   2.204  23.020  1.00  0.00
ATOM   1979 2HD1 LEU A 123       6.032   2.880  24.116  1.00  0.00
ATOM   1980 3HD1 LEU A 123       5.195   3.827  23.083  1.00  0.00
ATOM   1981  CD2 LEU A 123       6.269   2.898  20.768  1.00  0.00           C
ATOM   1982 1HD2 LEU A 123       5.631   2.138  20.644  1.00  0.00
ATOM   1983 2HD2 LEU A 123       5.793   3.763  20.609  1.00  0.00
```

| | | | | | | | | | | | |
|---|---|---|---|---|---|---|---|---|---|---|---|
| ATOM | 1984 | 3HD2 | LEU | A | 123 | 7.024 | 2.808 | 20.119 | 1.00 | 0.00 | |
| ATOM | 1985 | C | LEU | A | 123 | 9.391 | 5.772 | 21.605 | 1.00 | 0.00 | C |
| ATOM | 1986 | O | LEU | A | 123 | 9.054 | 6.882 | 21.226 | 1.00 | 0.00 | O |
| ATOM | 1987 | N | HIS | A | 124 | 10.475 | 5.542 | 22.347 | 1.00 | 0.00 | N |
| ATOM | 1988 | HN | HIS | A | 124 | 10.736 | 4.591 | 22.515 | 1.00 | 0.00 | |
| ATOM | 1989 | CA | HIS | A | 124 | 11.312 | 6.603 | 22.935 | 1.00 | 0.00 | C |
| ATOM | 1990 | HA | HIS | A | 124 | 10.681 | 7.182 | 23.452 | 1.00 | 0.00 | |
| ATOM | 1991 | CB | HIS | A | 124 | 12.352 | 6.019 | 23.882 | 1.00 | 0.00 | C |
| ATOM | 1992 | HB1 | HIS | A | 124 | 11.885 | 5.487 | 24.588 | 1.00 | 0.00 | |
| ATOM | 1993 | HB2 | HIS | A | 124 | 12.963 | 5.421 | 23.364 | 1.00 | 0.00 | |
| ATOM | 1994 | ND1 | HIS | A | 124 | 12.653 | 7.957 | 25.483 | 1.00 | 0.00 | N |
| ATOM | 1995 | CG | HIS | A | 124 | 13.185 | 7.054 | 24.560 | 1.00 | 0.00 | C |
| ATOM | 1996 | CE1 | HIS | A | 124 | 13.634 | 8.716 | 25.950 | 1.00 | 0.00 | C |
| ATOM | 1997 | HE1 | HIS | A | 124 | 13.538 | 9.449 | 26.624 | 1.00 | 0.00 | |
| ATOM | 1998 | NE2 | HIS | A | 124 | 14.786 | 8.329 | 25.358 | 1.00 | 0.00 | N |
| ATOM | 1999 | HE2 | HIS | A | 124 | 15.687 | 8.732 | 25.518 | 1.00 | 0.00 | |
| ATOM | 2000 | CD2 | HIS | A | 124 | 14.515 | 7.295 | 24.503 | 1.00 | 0.00 | C |
| ATOM | 2001 | HD2 | HIS | A | 124 | 15.180 | 6.806 | 23.938 | 1.00 | 0.00 | |
| ATOM | 2002 | C | HIS | A | 124 | 12.001 | 7.429 | 21.845 | 1.00 | 0.00 | C |
| ATOM | 2003 | O | HIS | A | 124 | 12.120 | 8.640 | 21.963 | 1.00 | 0.00 | O |
| ATOM | 2004 | N | SER | A | 125 | 12.454 | 6.763 | 20.786 | 1.00 | 0.00 | N |
| ATOM | 2005 | HN | SER | A | 125 | 12.303 | 5.776 | 20.733 | 1.00 | 0.00 | |
| ATOM | 2006 | CA | SER | A | 125 | 13.154 | 7.420 | 19.716 | 1.00 | 0.00 | C |
| ATOM | 2007 | HA | SER | A | 125 | 13.839 | 7.998 | 20.160 | 1.00 | 0.00 | |
| ATOM | 2008 | CB | SER | A | 125 | 13.880 | 6.425 | 18.876 | 1.00 | 0.00 | C |
| ATOM | 2009 | HB1 | SER | A | 125 | 14.461 | 6.889 | 18.207 | 1.00 | 0.00 | |
| ATOM | 2010 | HB2 | SER | A | 125 | 14.442 | 5.827 | 19.448 | 1.00 | 0.00 | |
| ATOM | 2011 | OG | SER | A | 125 | 12.956 | 5.623 | 18.176 | 1.00 | 0.00 | O |
| ATOM | 2012 | HG1 | SER | A | 125 | 13.449 | 4.958 | 17.615 | 1.00 | 0.00 | |
| ATOM | 2013 | C | SER | A | 125 | 12.194 | 8.283 | 18.881 | 1.00 | 0.00 | C |
| ATOM | 2014 | O | SER | A | 125 | 12.623 | 9.252 | 18.275 | 1.00 | 0.00 | O |
| ATOM | 2015 | N | LYS | A | 126 | 10.902 | 7.941 | 18.872 | 1.00 | 0.00 | N |
| ATOM | 2016 | HN | LYS | A | 126 | 10.625 | 7.113 | 19.360 | 1.00 | 0.00 | |
| ATOM | 2017 | CA | LYS | A | 126 | 9.860 | 8.731 | 18.172 | 1.00 | 0.00 | C |
| ATOM | 2018 | HA | LYS | A | 126 | 10.358 | 9.208 | 17.448 | 1.00 | 0.00 | |
| ATOM | 2019 | CB | LYS | A | 126 | 8.765 | 7.828 | 17.598 | 1.00 | 0.00 | C |
| ATOM | 2020 | HB1 | LYS | A | 126 | 8.241 | 7.457 | 18.365 | 1.00 | 0.00 | |
| ATOM | 2021 | HB2 | LYS | A | 126 | 8.168 | 8.394 | 17.030 | 1.00 | 0.00 | |
| ATOM | 2022 | CG | LYS | A | 126 | 9.228 | 6.654 | 16.748 | 1.00 | 0.00 | C |
| ATOM | 2023 | HG1 | LYS | A | 126 | 10.030 | 6.241 | 17.180 | 1.00 | 0.00 | |
| ATOM | 2024 | HG2 | LYS | A | 126 | 8.490 | 5.980 | 16.702 | 1.00 | 0.00 | |
| ATOM | 2025 | CD | LYS | A | 126 | 9.600 | 7.024 | 15.348 | 1.00 | 0.00 | C |
| ATOM | 2026 | HD1 | LYS | A | 126 | 8.849 | 7.546 | 14.944 | 1.00 | 0.00 | |
| ATOM | 2027 | HD2 | LYS | A | 126 | 10.424 | 7.590 | 15.374 | 1.00 | 0.00 | |
| ATOM | 2028 | CE | LYS | A | 126 | 9.866 | 5.812 | 14.498 | 1.00 | 0.00 | C |
| ATOM | 2029 | HE1 | LYS | A | 126 | 10.616 | 5.281 | 14.893 | 1.00 | 0.00 | |
| ATOM | 2030 | HE2 | LYS | A | 126 | 9.044 | 5.245 | 14.452 | 1.00 | 0.00 | |
| ATOM | 2031 | NZ | LYS | A | 126 | 10.240 | 6.227 | 13.130 | 1.00 | 0.00 | N |
| ATOM | 2032 | HZ1 | LYS | A | 126 | 10.415 | 5.416 | 12.572 | 1.00 | 0.00 | |
| ATOM | 2033 | HZ2 | LYS | A | 126 | 9.493 | 6.755 | 12.726 | 1.00 | 0.00 | |
| ATOM | 2034 | HZ3 | LYS | A | 126 | 11.065 | 6.791 | 13.167 | 1.00 | 0.00 | |
| ATOM | 2035 | C | LYS | A | 126 | 9.168 | 9.710 | 19.129 | 1.00 | 0.00 | C |
| ATOM | 2036 | O | LYS | A | 126 | 8.267 | 10.415 | 18.724 | 1.00 | 0.00 | O |
| ATOM | 2037 | N | GLY | A | 127 | 9.536 | 9.680 | 20.415 | 1.00 | 0.00 | N |
| ATOM | 2038 | HN | GLY | A | 127 | 10.235 | 9.022 | 20.696 | 1.00 | 0.00 | |
| ATOM | 2039 | CA | GLY | A | 127 | 8.963 | 10.566 | 21.431 | 1.00 | 0.00 | C |
| ATOM | 2040 | HA1 | GLY | A | 127 | 9.490 | 10.443 | 22.272 | 1.00 | 0.00 | |
| ATOM | 2041 | HA2 | GLY | A | 127 | 9.067 | 11.506 | 21.107 | 1.00 | 0.00 | |
| ATOM | 2042 | C | GLY | A | 127 | 7.488 | 10.282 | 21.707 | 1.00 | 0.00 | C |
| ATOM | 2043 | O | GLY | A | 127 | 6.728 | 11.205 | 21.964 | 1.00 | 0.00 | O |
| ATOM | 2044 | N | ILE | A | 128 | 7.115 | 8.993 | 21.732 | 1.00 | 0.00 | N |
| ATOM | 2045 | HN | ILE | A | 128 | 7.809 | 8.296 | 21.555 | 1.00 | 0.00 | |
| ATOM | 2046 | CA | ILE | A | 128 | 5.758 | 8.558 | 22.002 | 1.00 | 0.00 | C |
| ATOM | 2047 | HA | ILE | A | 128 | 5.200 | 9.387 | 21.994 | 1.00 | 0.00 | |
| ATOM | 2048 | CB | ILE | A | 128 | 5.262 | 7.595 | 20.914 | 1.00 | 0.00 | C |
| ATOM | 2049 | HB | ILE | A | 128 | 5.920 | 6.847 | 20.826 | 1.00 | 0.00 | |
| ATOM | 2050 | CG2 | ILE | A | 128 | 3.913 | 6.986 | 21.311 | 1.00 | 0.00 | C |
| ATOM | 2051 | 1HG2 | ILE | A | 128 | 3.605 | 6.362 | 20.593 | 1.00 | 0.00 | |
| ATOM | 2052 | 2HG2 | ILE | A | 128 | 4.014 | 6.482 | 22.169 | 1.00 | 0.00 | |
| ATOM | 2053 | 3HG2 | ILE | A | 128 | 3.240 | 7.716 | 21.430 | 1.00 | 0.00 | |
| ATOM | 2054 | CG1 | ILE | A | 128 | 5.202 | 8.280 | 19.551 | 1.00 | 0.00 | C |

```
ATOM   2055 1HG1 ILE A 128       4.460   8.950  19.565  1.00  0.00
ATOM   2056 2HG1 ILE A 128       6.071   8.748  19.392  1.00  0.00
ATOM   2057  CD  ILE A 128       4.961   7.346  18.394  1.00  0.00           C
ATOM   2058  HD1 ILE A 128       4.936   7.870  17.543  1.00  0.00
ATOM   2059  HD2 ILE A 128       5.699   6.673  18.349  1.00  0.00
ATOM   2060  HD3 ILE A 128       4.088   6.875  18.522  1.00  0.00
ATOM   2061  C   ILE A 128       5.709   7.880  23.372  1.00  0.00           C
ATOM   2062  O   ILE A 128       6.517   7.017  23.678  1.00  0.00           O
ATOM   2063  N   ILE A 129       4.724   8.282  24.167  1.00  0.00           N
ATOM   2064  HN  ILE A 129       4.139   9.027  23.846  1.00  0.00
ATOM   2065  CA  ILE A 129       4.445   7.717  25.456  1.00  0.00           C
ATOM   2066  HA  ILE A 129       5.204   7.103  25.671  1.00  0.00
ATOM   2067  CB  ILE A 129       4.362   8.809  26.543  1.00  0.00           C
ATOM   2068  HB  ILE A 129       3.546   9.357  26.361  1.00  0.00
ATOM   2069  CG2 ILE A 129       4.209   8.204  27.920  1.00  0.00           C
ATOM   2070 1HG2 ILE A 129       4.158   8.934  28.601  1.00  0.00
ATOM   2071 2HG2 ILE A 129       3.372   7.658  27.953  1.00  0.00
ATOM   2072 3HG2 ILE A 129       4.996   7.619  28.116  1.00  0.00
ATOM   2073  CG1 ILE A 129       5.557   9.750  26.504  1.00  0.00           C
ATOM   2074 1HG1 ILE A 129       6.353   9.284  26.891  1.00  0.00
ATOM   2075 2HG1 ILE A 129       5.746   9.999  25.554  1.00  0.00
ATOM   2076  CD  ILE A 129       5.326  11.006  27.281  1.00  0.00           C
ATOM   2077  HD1 ILE A 129       6.138  11.587  27.224  1.00  0.00
ATOM   2078  HD2 ILE A 129       4.540  11.493  26.901  1.00  0.00
ATOM   2079  HD3 ILE A 129       5.146  10.778  28.238  1.00  0.00
ATOM   2080  C   ILE A 129       3.124   6.970  25.340  1.00  0.00           C
ATOM   2081  O   ILE A 129       2.142   7.529  24.877  1.00  0.00           O
ATOM   2082  N   HIS A 130       3.102   5.726  25.814  1.00  0.00           N
ATOM   2083  HN  HIS A 130       3.921   5.369  26.263  1.00  0.00
ATOM   2084  CA  HIS A 130       1.941   4.870  25.706  1.00  0.00           C
ATOM   2085  HA  HIS A 130       1.535   5.029  24.806  1.00  0.00
ATOM   2086  CB  HIS A 130       2.395   3.411  25.803  1.00  0.00           C
ATOM   2087  HB1 HIS A 130       3.108   3.244  25.122  1.00  0.00
ATOM   2088  HB2 HIS A 130       2.762   3.243  26.718  1.00  0.00
ATOM   2089  ND1 HIS A 130       0.248   2.267  26.444  1.00  0.00           N
ATOM   2090  CG  HIS A 130       1.303   2.431  25.566  1.00  0.00           C
ATOM   2091  CE1 HIS A 130      -0.550   1.336  25.979  1.00  0.00           C
ATOM   2092  HE1 HIS A 130      -1.393   1.014  26.410  1.00  0.00
ATOM   2093  NE2 HIS A 130      -0.035   0.892  24.821  1.00  0.00           N
ATOM   2094  HE2 HIS A 130      -0.432   0.178  24.244  1.00  0.00
ATOM   2095  CD2 HIS A 130       1.108   1.568  24.562  1.00  0.00           C
ATOM   2096  HD2 HIS A 130       1.700   1.445  23.765  1.00  0.00
ATOM   2097  C   HIS A 130       0.895   5.250  26.764  1.00  0.00           C
ATOM   2098  O   HIS A 130      -0.266   5.437  26.437  1.00  0.00           O
ATOM   2099  N   ARG A 131       1.331   5.296  28.029  1.00  0.00           N
ATOM   2100  HN  ARG A 131       2.251   4.947  28.207  1.00  0.00
ATOM   2101  CA  ARG A 131       0.574   5.816  29.185  1.00  0.00           C
ATOM   2102  HA  ARG A 131       1.282   5.963  29.876  1.00  0.00
ATOM   2103  CB  ARG A 131      -0.120   7.142  28.864  1.00  0.00           C
ATOM   2104  HB1 ARG A 131      -0.782   6.987  28.131  1.00  0.00
ATOM   2105  HB2 ARG A 131      -0.597   7.460  29.683  1.00  0.00
ATOM   2106  CG  ARG A 131       0.838   8.232  28.418  1.00  0.00           C
ATOM   2107  HG1 ARG A 131       1.727   8.081  28.850  1.00  0.00
ATOM   2108  HG2 ARG A 131       0.939   8.190  27.424  1.00  0.00
ATOM   2109  CD  ARG A 131       0.346   9.605  28.795  1.00  0.00           C
ATOM   2110  HD1 ARG A 131       0.335   9.694  29.791  1.00  0.00
ATOM   2111  HD2 ARG A 131       0.954  10.296  28.405  1.00  0.00
ATOM   2112  NE  ARG A 131      -0.995   9.870  28.311  1.00  0.00           N
ATOM   2113  HE  ARG A 131      -1.336   9.310  27.556  1.00  0.00
ATOM   2114  CZ  ARG A 131      -1.797  10.799  28.798  1.00  0.00           C
ATOM   2115  NH1 ARG A 131      -1.388  11.595  29.775  1.00  0.00           N
ATOM   2116 1HH1 ARG A 131      -0.466  11.495  30.149  1.00  0.00
ATOM   2117 2HH1 ARG A 131      -2.001  12.296  30.139  1.00  0.00
ATOM   2118  NH2 ARG A 131      -3.013  10.917  28.302  1.00  0.00           N
ATOM   2119 1HH2 ARG A 131      -3.315  10.308  27.568  1.00  0.00
ATOM   2120 2HH2 ARG A 131      -3.633  11.615  28.659  1.00  0.00
ATOM   2121  C   ARG A 131      -0.456   4.809  29.719  1.00  0.00           C
ATOM   2122  O   ARG A 131      -1.005   5.023  30.787  1.00  0.00           O
ATOM   2123  N   ASP A 132      -0.673   3.699  29.019  1.00  0.00           N
ATOM   2124  HN  ASP A 132      -0.113   3.518  28.210  1.00  0.00
ATOM   2125  CA  ASP A 132      -1.701   2.739  29.395  1.00  0.00           C
```

| | | | | | | | | | | | |
|---|---|---|---|---|---|---|---|---|---|---|---|
| ATOM | 2126 | HA | ASP | A | 132 | -1.902 | 2.885 | 30.364 | 1.00 | 0.00 | |
| ATOM | 2127 | CB | ASP | A | 132 | -2.964 | 3.000 | 28.585 | 1.00 | 0.00 | C |
| ATOM | 2128 | HB1 | ASP | A | 132 | -3.132 | 3.986 | 28.591 | 1.00 | 0.00 | |
| ATOM | 2129 | HB2 | ASP | A | 132 | -2.793 | 2.697 | 27.647 | 1.00 | 0.00 | |
| ATOM | 2130 | CG | ASP | A | 132 | -4.230 | 2.308 | 29.075 | 1.00 | 0.00 | C |
| ATOM | 2131 | OD1 | ASP | A | 132 | -4.342 | 2.046 | 30.321 | 1.00 | 0.00 | O |
| ATOM | 2132 | OD2 | ASP | A | 132 | -5.115 | 2.037 | 28.199 | 1.00 | 0.00 | O |
| ATOM | 2133 | C | ASP | A | 132 | -1.180 | 1.310 | 29.205 | 1.00 | 0.00 | C |
| ATOM | 2134 | O | ASP | A | 132 | -1.948 | 0.418 | 28.900 | 1.00 | 0.00 | O |
| ATOM | 2135 | N | LEU | A | 133 | 0.125 | 1.101 | 29.401 | 1.00 | 0.00 | N |
| ATOM | 2136 | HN | LEU | A | 133 | 0.725 | 1.880 | 29.582 | 1.00 | 0.00 | |
| ATOM | 2137 | CA | LEU | A | 133 | 0.691 | -0.240 | 29.356 | 1.00 | 0.00 | C |
| ATOM | 2138 | HA | LEU | A | 133 | 0.420 | -0.630 | 28.476 | 1.00 | 0.00 | |
| ATOM | 2139 | CB | LEU | A | 133 | 2.213 | -0.201 | 29.457 | 1.00 | 0.00 | C |
| ATOM | 2140 | HB1 | LEU | A | 133 | 2.456 | 0.476 | 30.152 | 1.00 | 0.00 | |
| ATOM | 2141 | HB2 | LEU | A | 133 | 2.523 | -1.106 | 29.749 | 1.00 | 0.00 | |
| ATOM | 2142 | CG | LEU | A | 133 | 2.970 | 0.153 | 28.194 | 1.00 | 0.00 | C |
| ATOM | 2143 | HG | LEU | A | 133 | 2.732 | 1.088 | 27.930 | 1.00 | 0.00 | |
| ATOM | 2144 | CD1 | LEU | A | 133 | 4.469 | 0.058 | 28.462 | 1.00 | 0.00 | C |
| ATOM | 2145 | 1HD1 | LEU | A | 133 | 4.972 | 0.291 | 27.630 | 1.00 | 0.00 | |
| ATOM | 2146 | 2HD1 | LEU | A | 133 | 4.718 | 0.695 | 29.192 | 1.00 | 0.00 | |
| ATOM | 2147 | 3HD1 | LEU | A | 133 | 4.699 | -0.874 | 28.740 | 1.00 | 0.00 | |
| ATOM | 2148 | CD2 | LEU | A | 133 | 2.574 | -0.759 | 27.047 | 1.00 | 0.00 | C |
| ATOM | 2149 | 1HD2 | LEU | A | 133 | 3.087 | -0.505 | 26.227 | 1.00 | 0.00 | |
| ATOM | 2150 | 2HD2 | LEU | A | 133 | 2.780 | -1.707 | 27.289 | 1.00 | 0.00 | |
| ATOM | 2151 | 3HD2 | LEU | A | 133 | 1.594 | -0.666 | 26.870 | 1.00 | 0.00 | |
| ATOM | 2152 | C | LEU | A | 133 | 0.144 | -1.031 | 30.539 | 1.00 | 0.00 | C |
| ATOM | 2153 | O | LEU | A | 133 | 0.164 | -0.569 | 31.664 | 1.00 | 0.00 | O |
| ATOM | 2154 | N | LYS | A | 134 | -0.327 | -2.233 | 30.253 | 1.00 | 0.00 | N |
| ATOM | 2155 | HN | LYS | A | 134 | -0.325 | -2.527 | 29.297 | 1.00 | 0.00 | |
| ATOM | 2156 | CA | LYS | A | 134 | -0.833 | -3.125 | 31.226 | 1.00 | 0.00 | C |
| ATOM | 2157 | HA | LYS | A | 134 | -0.173 | -3.155 | 31.977 | 1.00 | 0.00 | |
| ATOM | 2158 | CB | LYS | A | 134 | -2.162 | -2.599 | 31.775 | 1.00 | 0.00 | C |
| ATOM | 2159 | HB1 | LYS | A | 134 | -2.476 | -3.230 | 32.485 | 1.00 | 0.00 | |
| ATOM | 2160 | HB2 | LYS | A | 134 | -1.997 | -1.699 | 32.178 | 1.00 | 0.00 | |
| ATOM | 2161 | CG | LYS | A | 134 | -3.284 | -2.449 | 30.772 | 1.00 | 0.00 | C |
| ATOM | 2162 | HG1 | LYS | A | 134 | -2.984 | -1.835 | 30.042 | 1.00 | 0.00 | |
| ATOM | 2163 | HG2 | LYS | A | 134 | -3.495 | -3.348 | 30.387 | 1.00 | 0.00 | |
| ATOM | 2164 | CD | LYS | A | 134 | -4.537 | -1.883 | 31.397 | 1.00 | 0.00 | C |
| ATOM | 2165 | HD1 | LYS | A | 134 | -4.766 | -2.432 | 32.200 | 1.00 | 0.00 | |
| ATOM | 2166 | HD2 | LYS | A | 134 | -4.353 | -0.941 | 31.679 | 1.00 | 0.00 | |
| ATOM | 2167 | CE | LYS | A | 134 | -5.725 | -1.882 | 30.459 | 1.00 | 0.00 | C |
| ATOM | 2168 | HE1 | LYS | A | 134 | -5.399 | -1.848 | 29.514 | 1.00 | 0.00 | |
| ATOM | 2169 | HE2 | LYS | A | 134 | -6.256 | -2.718 | 30.598 | 1.00 | 0.00 | |
| ATOM | 2170 | NZ | LYS | A | 134 | -6.615 | -0.717 | 30.691 | 1.00 | 0.00 | N |
| ATOM | 2171 | HZ1 | LYS | A | 134 | -7.385 | -0.753 | 30.054 | 1.00 | 0.00 | |
| ATOM | 2172 | HZ2 | LYS | A | 134 | -6.958 | -0.742 | 31.630 | 1.00 | 0.00 | |
| ATOM | 2173 | HZ3 | LYS | A | 134 | -6.102 | 0.129 | 30.546 | 1.00 | 0.00 | |
| ATOM | 2174 | C | LYS | A | 134 | -0.945 | -4.500 | 30.586 | 1.00 | 0.00 | C |
| ATOM | 2175 | O | LYS | A | 134 | -0.811 | -4.625 | 29.383 | 1.00 | 0.00 | O |
| ATOM | 2176 | N | PRO | A | 135 | -1.133 | -5.580 | 31.368 | 1.00 | 0.00 | N |
| ATOM | 2177 | CD | PRO | A | 135 | -1.237 | -5.574 | 32.836 | 1.00 | 0.00 | C |
| ATOM | 2178 | HD1 | PRO | A | 135 | -2.162 | -5.326 | 33.123 | 1.00 | 0.00 | |
| ATOM | 2179 | HD2 | PRO | A | 135 | -0.581 | -4.933 | 33.234 | 1.00 | 0.00 | |
| ATOM | 2180 | CA | PRO | A | 135 | -1.170 | -6.932 | 30.814 | 1.00 | 0.00 | C |
| ATOM | 2181 | HA | PRO | A | 135 | -0.302 | -7.118 | 30.354 | 1.00 | 0.00 | |
| ATOM | 2182 | CB | PRO | A | 135 | -1.451 | -7.811 | 32.038 | 1.00 | 0.00 | C |
| ATOM | 2183 | HB1 | PRO | A | 135 | -2.432 | -7.967 | 32.152 | 1.00 | 0.00 | |
| ATOM | 2184 | HB2 | PRO | A | 135 | -0.980 | -8.690 | 31.967 | 1.00 | 0.00 | |
| ATOM | 2185 | CG | PRO | A | 135 | -0.905 | -7.010 | 33.193 | 1.00 | 0.00 | C |
| ATOM | 2186 | HG1 | PRO | A | 135 | -1.348 | -7.276 | 34.049 | 1.00 | 0.00 | |
| ATOM | 2187 | HG2 | PRO | A | 135 | 0.083 | -7.138 | 33.276 | 1.00 | 0.00 | |
| ATOM | 2188 | C | PRO | A | 135 | -2.244 | -7.123 | 29.734 | 1.00 | 0.00 | C |
| ATOM | 2189 | O | PRO | A | 135 | -2.032 | -7.896 | 28.806 | 1.00 | 0.00 | O |
| ATOM | 2190 | N | GLN | A | 136 | -3.359 | -6.393 | 29.847 | 1.00 | 0.00 | N |
| ATOM | 2191 | HN | GLN | A | 136 | -3.442 | -5.758 | 30.615 | 1.00 | 0.00 | |
| ATOM | 2192 | CA | GLN | A | 136 | -4.475 | -6.488 | 28.879 | 1.00 | 0.00 | C |
| ATOM | 2193 | HA | GLN | A | 136 | -4.691 | -7.460 | 28.785 | 1.00 | 0.00 | |
| ATOM | 2194 | CB | GLN | A | 136 | -5.721 | -5.777 | 29.417 | 1.00 | 0.00 | C |
| ATOM | 2195 | HB1 | GLN | A | 136 | -5.459 | -4.849 | 29.681 | 1.00 | 0.00 | |
| ATOM | 2196 | HB2 | GLN | A | 136 | -6.397 | -5.736 | 28.681 | 1.00 | 0.00 | |

```
ATOM   2197  CG  GLN A 136      -6.377  -6.448  30.627  1.00  0.00           C
ATOM   2198  HG1 GLN A 136      -7.330  -6.148  30.663  1.00  0.00
ATOM   2199  HG2 GLN A 136      -6.344  -7.437  30.483  1.00  0.00
ATOM   2200  CD  GLN A 136      -5.748  -6.157  31.989  1.00  0.00           C
ATOM   2201  OE1 GLN A 136      -4.798  -5.374  32.150  1.00  0.00           O
ATOM   2202  NE2 GLN A 136      -6.274  -6.834  33.007  1.00  0.00           N
ATOM   2203 1HE2 GLN A 136      -7.024  -7.475  32.846  1.00  0.00
ATOM   2204 2HE2 GLN A 136      -5.919  -6.701  33.932  1.00  0.00
ATOM   2205  C   GLN A 136      -4.038  -5.913  27.507  1.00  0.00           C
ATOM   2206  O   GLN A 136      -4.591  -6.258  26.477  1.00  0.00           O
ATOM   2207  N   ASN A 137      -3.011  -5.056  27.510  1.00  0.00           N
ATOM   2208  HN  ASN A 137      -2.531  -4.904  28.374  1.00  0.00
ATOM   2209  CA  ASN A 137      -2.548  -4.330  26.335  1.00  0.00           C
ATOM   2210  HA  ASN A 137      -3.257  -4.455  25.641  1.00  0.00
ATOM   2211  CB  ASN A 137      -2.417  -2.838  26.642  1.00  0.00           C
ATOM   2212  HB1 ASN A 137      -1.986  -2.726  27.538  1.00  0.00
ATOM   2213  HB2 ASN A 137      -1.846  -2.410  25.942  1.00  0.00
ATOM   2214  CG  ASN A 137      -3.755  -2.137  26.663  1.00  0.00           C
ATOM   2215  OD1 ASN A 137      -4.759  -2.697  26.224  1.00  0.00           O
ATOM   2216  ND2 ASN A 137      -3.784  -0.911  27.165  1.00  0.00           N
ATOM   2217 1HD2 ASN A 137      -2.944  -0.491  27.508  1.00  0.00
ATOM   2218 2HD2 ASN A 137      -4.646  -0.406  27.201  1.00  0.00
ATOM   2219  C   ASN A 137      -1.224  -4.921  25.824  1.00  0.00           C
ATOM   2220  O   ASN A 137      -0.583  -4.346  24.974  1.00  0.00           O
ATOM   2221  N   ILE A 138      -0.835  -6.083  26.342  1.00  0.00           N
ATOM   2222  HN  ILE A 138      -1.375  -6.496  27.075  1.00  0.00
ATOM   2223  CA  ILE A 138       0.353  -6.765  25.869  1.00  0.00           C
ATOM   2224  HA  ILE A 138       0.762  -6.197  25.155  1.00  0.00
ATOM   2225  CB  ILE A 138       1.382  -6.911  26.994  1.00  0.00           C
ATOM   2226  HB  ILE A 138       0.950  -7.376  27.766  1.00  0.00
ATOM   2227  CG2 ILE A 138       2.537  -7.768  26.541  1.00  0.00           C
ATOM   2228 1HG2 ILE A 138       3.200  -7.854  27.285  1.00  0.00
ATOM   2229 2HG2 ILE A 138       2.200  -8.675  26.288  1.00  0.00
ATOM   2230 3HG2 ILE A 138       2.977  -7.343  25.750  1.00  0.00
ATOM   2231  CG1 ILE A 138       1.858  -5.552  27.497  1.00  0.00           C
ATOM   2232 1HG1 ILE A 138       2.465  -5.154  26.809  1.00  0.00
ATOM   2233 2HG1 ILE A 138       1.060  -4.962  27.620  1.00  0.00
ATOM   2234  CD  ILE A 138       2.607  -5.607  28.808  1.00  0.00           C
ATOM   2235  HD1 ILE A 138       2.885  -4.683  29.072  1.00  0.00
ATOM   2236  HD2 ILE A 138       2.013  -5.991  29.515  1.00  0.00
ATOM   2237  HD3 ILE A 138       3.418  -6.183  28.705  1.00  0.00
ATOM   2238  C   ILE A 138      -0.075  -8.117  25.297  1.00  0.00           C
ATOM   2239  O   ILE A 138      -0.567  -8.961  26.010  1.00  0.00           O
ATOM   2240  N   LEU A 139       0.095  -8.276  23.992  1.00  0.00           N
ATOM   2241  HN  LEU A 139       0.607  -7.582  23.486  1.00  0.00
ATOM   2242  CA  LEU A 139      -0.431  -9.412  23.274  1.00  0.00           C
ATOM   2243  HA  LEU A 139      -1.121  -9.833  23.862  1.00  0.00
ATOM   2244  CB  LEU A 139      -1.059  -8.931  21.963  1.00  0.00           C
ATOM   2245  HB1 LEU A 139      -0.368  -8.412  21.460  1.00  0.00
ATOM   2246  HB2 LEU A 139      -1.323  -9.736  21.432  1.00  0.00
ATOM   2247  CG  LEU A 139      -2.290  -8.046  22.113  1.00  0.00           C
ATOM   2248  HG  LEU A 139      -2.034  -7.243  22.652  1.00  0.00
ATOM   2249  CD1 LEU A 139      -2.770  -7.566  20.753  1.00  0.00           C
ATOM   2250 1HD1 LEU A 139      -3.577  -6.987  20.870  1.00  0.00
ATOM   2251 2HD1 LEU A 139      -2.044  -7.040  20.310  1.00  0.00
ATOM   2252 3HD1 LEU A 139      -3.005  -8.355  20.185  1.00  0.00
ATOM   2253  CD2 LEU A 139      -3.405  -8.782  22.857  1.00  0.00           C
ATOM   2254 1HD2 LEU A 139      -4.201  -8.182  22.943  1.00  0.00
ATOM   2255 2HD2 LEU A 139      -3.660  -9.603  22.347  1.00  0.00
ATOM   2256 3HD2 LEU A 139      -3.083  -9.042  23.768  1.00  0.00
ATOM   2257  C   LEU A 139       0.690 -10.407  22.999  1.00  0.00           C
ATOM   2258  O   LEU A 139       1.876 -10.043  22.994  1.00  0.00           O
ATOM   2259  N   LEU A 140       0.276 -11.651  22.745  1.00  0.00           N
ATOM   2260  HN  LEU A 140      -0.703 -11.803  22.609  1.00  0.00
ATOM   2261  CA  LEU A 140       1.166 -12.795  22.655  1.00  0.00           C
ATOM   2262  HA  LEU A 140       2.104 -12.458  22.733  1.00  0.00
ATOM   2263  CB  LEU A 140       0.819 -13.755  23.797  1.00  0.00           C
ATOM   2264  HB1 LEU A 140      -0.153 -13.649  24.007  1.00  0.00
ATOM   2265  HB2 LEU A 140       0.992 -14.688  23.482  1.00  0.00
ATOM   2266  CG  LEU A 140       1.596 -13.550  25.086  1.00  0.00           C
ATOM   2267  HG  LEU A 140       2.558 -13.764  24.915  1.00  0.00
```

```
ATOM   2268  CD1 LEU A 140       1.539 -12.109  25.542  1.00  0.00           C
ATOM   2269 1HD1 LEU A 140       2.058 -12.007  26.390  1.00  0.00
ATOM   2270 2HD1 LEU A 140       1.934 -11.520  24.837  1.00  0.00
ATOM   2271 3HD1 LEU A 140       0.587 -11.847  25.700  1.00  0.00
ATOM   2272  CD2 LEU A 140       1.068 -14.483  26.161  1.00  0.00           C
ATOM   2273 1HD2 LEU A 140       1.583 -14.343  27.006  1.00  0.00
ATOM   2274 2HD2 LEU A 140       0.101 -14.290  26.325  1.00  0.00
ATOM   2275 3HD2 LEU A 140       1.172 -15.431  25.860  1.00  0.00
ATOM   2276  C   LEU A 140       0.986 -13.488  21.305  1.00  0.00           C
ATOM   2277  O   LEU A 140      -0.133 -13.734  20.886  1.00  0.00           O
ATOM   2278  N   SER A 141       2.106 -13.795  20.653  1.00  0.00           N
ATOM   2279  HN  SER A 141       2.981 -13.512  21.046  1.00  0.00
ATOM   2280  CA  SER A 141       2.114 -14.524  19.394  1.00  0.00           C
ATOM   2281  HA  SER A 141       1.189 -14.875  19.248  1.00  0.00
ATOM   2282  CB  SER A 141       2.428 -13.597  18.247  1.00  0.00           C
ATOM   2283  HB1 SER A 141       1.698 -12.923  18.134  1.00  0.00
ATOM   2284  HB2 SER A 141       3.295 -13.126  18.407  1.00  0.00
ATOM   2285  OG  SER A 141       2.542 -14.332  17.032  1.00  0.00           O
ATOM   2286  HG1 SER A 141       2.750 -13.705  16.281  1.00  0.00
ATOM   2287  C   SER A 141       3.119 -15.683  19.489  1.00  0.00           C
ATOM   2288  O   SER A 141       4.265 -15.465  19.867  1.00  0.00           O
ATOM   2289  N   TYR A 142       2.666 -16.901  19.161  1.00  0.00           N
ATOM   2290  HN  TYR A 142       1.721 -16.983  18.845  1.00  0.00
ATOM   2291  CA  TYR A 142       3.489 -18.138  19.240  1.00  0.00           C
ATOM   2292  HA  TYR A 142       4.249 -17.922  19.853  1.00  0.00
ATOM   2293  CB  TYR A 142       2.663 -19.317  19.775  1.00  0.00           C
ATOM   2294  HB1 TYR A 142       1.852 -19.410  19.197  1.00  0.00
ATOM   2295  HB2 TYR A 142       3.223 -20.142  19.700  1.00  0.00
ATOM   2296  CG  TYR A 142       2.180 -19.240  21.204  1.00  0.00           C
ATOM   2297  CD1 TYR A 142       1.045 -18.522  21.554  1.00  0.00           C
ATOM   2298  HD1 TYR A 142       0.534 -18.039  20.843  1.00  0.00
ATOM   2299  CE1 TYR A 142       0.603 -18.459  22.867  1.00  0.00           C
ATOM   2300  HE1 TYR A 142      -0.191 -17.898  23.101  1.00  0.00
ATOM   2301  CZ  TYR A 142       1.262 -19.173  23.854  1.00  0.00           C
ATOM   2302  OH  TYR A 142       0.805 -19.127  25.137  1.00  0.00           O
ATOM   2303  HH  TYR A 142       1.389 -19.693  25.719  1.00  0.00
ATOM   2304  CD2 TYR A 142       2.822 -19.951  22.207  1.00  0.00           C
ATOM   2305  HD2 TYR A 142       3.636 -20.486  21.980  1.00  0.00
ATOM   2306  CE2 TYR A 142       2.365 -19.940  23.519  1.00  0.00           C
ATOM   2307  HE2 TYR A 142       2.830 -20.483  24.218  1.00  0.00
ATOM   2308  C   TYR A 142       3.960 -18.530  17.835  1.00  0.00           C
ATOM   2309  O   TYR A 142       3.173 -18.461  16.901  1.00  0.00           O
ATOM   2310  N   ALA A 143       5.194 -19.033  17.733  1.00  0.00           N
ATOM   2311  HN  ALA A 143       5.738 -19.122  18.567  1.00  0.00
ATOM   2312  CA  ALA A 143       5.802 -19.468  16.448  1.00  0.00           C
ATOM   2313  HA  ALA A 143       5.529 -18.796  15.759  1.00  0.00
ATOM   2314  CB  ALA A 143       7.316 -19.457  16.551  1.00  0.00           C
ATOM   2315  HB1 ALA A 143       7.711 -19.751  15.681  1.00  0.00
ATOM   2316  HB2 ALA A 143       7.629 -18.531  16.763  1.00  0.00
ATOM   2317  HB3 ALA A 143       7.605 -20.081  17.277  1.00  0.00
ATOM   2318  C   ALA A 143       5.278 -20.855  16.025  1.00  0.00           C
ATOM   2319  O   ALA A 143       4.593 -20.967  15.025  1.00  0.00           O
ATOM   2320  N   ASN A 144       5.445 -21.883  16.893  1.00  0.00           N
ATOM   2321  HN  ASN A 144       5.888 -21.714  17.774  1.00  0.00
ATOM   2322  CA  ASN A 144       4.998 -23.209  16.572  1.00  0.00           C
ATOM   2323  HA  ASN A 144       4.893 -23.178  15.578  1.00  0.00
ATOM   2324  CB  ASN A 144       5.987 -24.323  16.987  1.00  0.00           C
ATOM   2325  HB1 ASN A 144       5.610 -25.217  16.746  1.00  0.00
ATOM   2326  HB2 ASN A 144       6.860 -24.192  16.517  1.00  0.00
ATOM   2327  CG  ASN A 144       6.212 -24.261  18.497  1.00  0.00           C
ATOM   2328  OD1 ASN A 144       6.660 -23.239  19.016  1.00  0.00           O
ATOM   2329  ND2 ASN A 144       5.875 -25.367  19.216  1.00  0.00           N
ATOM   2330 1HD2 ASN A 144       5.499 -26.170  18.752  1.00  0.00
ATOM   2331 2HD2 ASN A 144       6.003 -25.377  20.208  1.00  0.00
ATOM   2332  C   ASN A 144       3.715 -23.466  17.290  1.00  0.00           C
ATOM   2333  O   ASN A 144       3.431 -22.907  18.345  1.00  0.00           O
ATOM   2334  N   ARG A 145       2.869 -24.317  16.704  1.00  0.00           N
ATOM   2335  HN  ARG A 145       3.068 -24.695  15.800  1.00  0.00
ATOM   2336  CA  ARG A 145       1.674 -24.675  17.398  1.00  0.00           C
ATOM   2337  HA  ARG A 145       1.751 -24.215  18.283  1.00  0.00
ATOM   2338  CB  ARG A 145       0.365 -24.218  16.725  1.00  0.00           C
```

```
ATOM   2339  HB1 ARG A 145       0.465 -24.307  15.734  1.00  0.00
ATOM   2340  HB2 ARG A 145      -0.380 -24.806  17.039  1.00  0.00
ATOM   2341  CG  ARG A 145       0.004 -22.767  17.046  1.00  0.00           C
ATOM   2342  HG1 ARG A 145       0.675 -22.164  16.615  1.00  0.00
ATOM   2343  HG2 ARG A 145      -0.905 -22.571  16.678  1.00  0.00
ATOM   2344  CD  ARG A 145      -0.004 -22.502  18.554  1.00  0.00           C
ATOM   2345  HD1 ARG A 145      -0.487 -23.235  19.033  1.00  0.00
ATOM   2346  HD2 ARG A 145       0.932 -22.437  18.901  1.00  0.00
ATOM   2347  NE  ARG A 145      -0.706 -21.214  18.796  1.00  0.00           N
ATOM   2348  HE  ARG A 145      -1.055 -20.673  18.031  1.00  0.00
ATOM   2349  CZ  ARG A 145      -0.865 -20.784  20.079  1.00  0.00           C
ATOM   2350  NH1 ARG A 145      -0.344 -21.525  21.099  1.00  0.00           N
ATOM   2351 1HH1 ARG A 145       0.148 -22.373  20.901  1.00  0.00
ATOM   2352 2HH1 ARG A 145      -0.454 -21.220  22.045  1.00  0.00
ATOM   2353  NH2 ARG A 145      -1.523 -19.622  20.355  1.00  0.00           N
ATOM   2354 1HH2 ARG A 145      -1.898 -19.071  19.610  1.00  0.00
ATOM   2355 2HH2 ARG A 145      -1.631 -19.321  21.303  1.00  0.00
ATOM   2356  C   ARG A 145       1.675 -26.162  17.471  1.00  0.00           C
ATOM   2357  O   ARG A 145       2.514 -26.822  16.860  1.00  0.00           O
ATOM   2358  N   ARG A 146       0.763 -26.718  18.301  1.00  0.00           N
ATOM   2359  HN  ARG A 146       0.218 -26.088  18.854  1.00  0.00
ATOM   2360  CA  ARG A 146       0.494 -28.131  18.472  1.00  0.00           C
ATOM   2361  HA  ARG A 146      -0.190 -28.111  19.201  1.00  0.00
ATOM   2362  CB  ARG A 146      -0.050 -28.776  17.171  1.00  0.00           C
ATOM   2363  HB1 ARG A 146      -0.793 -28.205  16.822  1.00  0.00
ATOM   2364  HB2 ARG A 146       0.690 -28.807  16.500  1.00  0.00
ATOM   2365  CG  ARG A 146      -0.590 -30.197  17.338  1.00  0.00           C
ATOM   2366  HG1 ARG A 146       0.054 -30.710  17.906  1.00  0.00
ATOM   2367  HG2 ARG A 146      -1.474 -30.142  17.803  1.00  0.00
ATOM   2368  CD  ARG A 146      -0.787 -30.963  16.029  1.00  0.00           C
ATOM   2369  HD1 ARG A 146      -1.613 -30.654  15.557  1.00  0.00
ATOM   2370  HD2 ARG A 146       0.004 -30.848  15.427  1.00  0.00
ATOM   2371  NE  ARG A 146      -0.929 -32.394  16.404  1.00  0.00           N
ATOM   2372  HE  ARG A 146      -1.823 -32.778  16.634  1.00  0.00
ATOM   2373  CZ  ARG A 146       0.192 -33.169  16.426  1.00  0.00           C
ATOM   2374  NH1 ARG A 146       1.367 -32.674  15.943  1.00  0.00           N
ATOM   2375 1HH1 ARG A 146       1.404 -31.745  15.574  1.00  0.00
ATOM   2376 2HH1 ARG A 146       2.191 -33.240  15.958  1.00  0.00
ATOM   2377  NH2 ARG A 146       0.143 -34.445  16.911  1.00  0.00           N
ATOM   2378 1HH2 ARG A 146      -0.720 -34.817  17.253  1.00  0.00
ATOM   2379 2HH2 ARG A 146       0.970 -35.007  16.924  1.00  0.00
ATOM   2380  C   ARG A 146       1.687 -28.926  18.966  1.00  0.00           C
ATOM   2381  O   ARG A 146       1.715 -30.154  18.841  1.00  0.00           O
ATOM   2382  N   LYS A 147       2.704 -28.258  19.551  1.00  0.00           N
ATOM   2383  HN  LYS A 147       2.635 -27.263  19.617  1.00  0.00
ATOM   2384  CA  LYS A 147       3.881 -28.891  20.086  1.00  0.00           C
ATOM   2385  HA  LYS A 147       3.565 -29.769  20.446  1.00  0.00
ATOM   2386  CB  LYS A 147       5.024 -29.120  19.065  1.00  0.00           C
ATOM   2387  HB1 LYS A 147       5.214 -28.246  18.619  1.00  0.00
ATOM   2388  HB2 LYS A 147       5.832 -29.418  19.573  1.00  0.00
ATOM   2389  CG  LYS A 147       4.775 -30.153  17.958  1.00  0.00           C
ATOM   2390  HG1 LYS A 147       3.944 -29.890  17.468  1.00  0.00
ATOM   2391  HG2 LYS A 147       5.553 -30.130  17.330  1.00  0.00
ATOM   2392  CD  LYS A 147       4.602 -31.594  18.447  1.00  0.00           C
ATOM   2393  HD1 LYS A 147       5.476 -31.916  18.810  1.00  0.00
ATOM   2394  HD2 LYS A 147       3.915 -31.605  19.173  1.00  0.00
ATOM   2395  CE  LYS A 147       4.155 -32.546  17.339  1.00  0.00           C
ATOM   2396  HE1 LYS A 147       3.274 -32.251  16.970  1.00  0.00
ATOM   2397  HE2 LYS A 147       4.835 -32.561  16.606  1.00  0.00
ATOM   2398  NZ  LYS A 147       4.008 -33.915  17.876  1.00  0.00           N
ATOM   2399  HZ1 LYS A 147       3.715 -34.530  17.144  1.00  0.00
ATOM   2400  HZ2 LYS A 147       4.885 -34.226  18.242  1.00  0.00
ATOM   2401  HZ3 LYS A 147       3.324 -33.915  18.605  1.00  0.00
ATOM   2402  C   LYS A 147       4.416 -27.902  21.070  1.00  0.00           C
ATOM   2403  O   LYS A 147       4.138 -26.711  20.950  1.00  0.00           O
ATOM   2404  N   SER A 148       5.190 -28.340  22.077  1.00  0.00           N
ATOM   2405  HN  SER A 148       5.408 -29.308  22.201  1.00  0.00
ATOM   2406  CA  SER A 148       5.681 -27.328  22.960  1.00  0.00           C
ATOM   2407  HA  SER A 148       5.056 -26.568  22.782  1.00  0.00
ATOM   2408  CB  SER A 148       5.679 -27.697  24.446  1.00  0.00           C
ATOM   2409  HB1 SER A 148       4.746 -27.762  24.801  1.00  0.00
```

```
ATOM   2410  HB2 SER A 148       6.157 -28.561  24.605  1.00  0.00
ATOM   2411  OG  SER A 148       6.358 -26.675  25.162  1.00  0.00           O
ATOM   2412  HG1 SER A 148       6.366 -26.900  26.136  1.00  0.00
ATOM   2413  C   SER A 148       7.100 -27.072  22.621  1.00  0.00           C
ATOM   2414  O   SER A 148       7.848 -27.971  22.250  1.00  0.00           O
ATOM   2415  N   SER A 149       7.505 -25.803  22.734  1.00  0.00           N
ATOM   2416  HN  SER A 149       6.858 -25.074  22.957  1.00  0.00
ATOM   2417  CA  SER A 149       8.884 -25.522  22.527  1.00  0.00           C
ATOM   2418  HA  SER A 149       9.315 -26.388  22.273  1.00  0.00
ATOM   2419  CB  SER A 149       9.192 -24.494  21.430  1.00  0.00           C
ATOM   2420  HB1 SER A 149      10.139 -24.573  21.120  1.00  0.00
ATOM   2421  HB2 SER A 149       8.576 -24.606  20.650  1.00  0.00
ATOM   2422  OG  SER A 149       9.012 -23.185  21.939  1.00  0.00           O
ATOM   2423  HG1 SER A 149       9.213 -22.518  21.222  1.00  0.00
ATOM   2424  C   SER A 149       9.274 -24.899  23.819  1.00  0.00           C
ATOM   2425  O   SER A 149       8.403 -24.555  24.616  1.00  0.00           O
ATOM   2426  N   VAL A 150      10.590 -24.729  24.053  1.00  0.00           N
ATOM   2427  HN  VAL A 150      11.242 -24.995  23.343  1.00  0.00
ATOM   2428  CA  VAL A 150      11.097 -24.179  25.285  1.00  0.00           C
ATOM   2429  HA  VAL A 150      10.817 -24.831  25.990  1.00  0.00
ATOM   2430  CB  VAL A 150      12.601 -24.036  25.273  1.00  0.00           C
ATOM   2431  HB  VAL A 150      12.883 -23.444  24.518  1.00  0.00
ATOM   2432  CG1 VAL A 150      13.069 -23.364  26.576  1.00  0.00           C
ATOM   2433 1HG1 VAL A 150      14.065 -23.271  26.564  1.00  0.00
ATOM   2434 2HG1 VAL A 150      12.651 -22.459  26.653  1.00  0.00
ATOM   2435 3HG1 VAL A 150      12.796 -23.925  27.357  1.00  0.00
ATOM   2436  CG2 VAL A 150      13.228 -25.423  25.051  1.00  0.00           C
ATOM   2437 1HG2 VAL A 150      14.225 -25.341  25.041  1.00  0.00
ATOM   2438 2HG2 VAL A 150      12.951 -26.036  25.791  1.00  0.00
ATOM   2439 3HG2 VAL A 150      12.916 -25.794  24.176  1.00  0.00
ATOM   2440  C   VAL A 150      10.512 -22.811  25.491  1.00  0.00           C
ATOM   2441  O   VAL A 150      10.029 -22.478  26.570  1.00  0.00           O
ATOM   2442  N   SER A 151      10.482 -21.989  24.441  1.00  0.00           N
ATOM   2443  HN  SER A 151      10.810 -22.296  23.547  1.00  0.00
ATOM   2444  CA  SER A 151       9.975 -20.663  24.611  1.00  0.00           C
ATOM   2445  HA  SER A 151       9.503 -20.606  25.491  1.00  0.00
ATOM   2446  CB  SER A 151      11.086 -19.603  24.566  1.00  0.00           C
ATOM   2447  HB1 SER A 151      11.769 -19.776  25.276  1.00  0.00
ATOM   2448  HB2 SER A 151      11.531 -19.595  23.670  1.00  0.00
ATOM   2449  OG  SER A 151      10.556 -18.303  24.792  1.00  0.00           O
ATOM   2450  HG1 SER A 151      11.296 -17.631  24.758  1.00  0.00
ATOM   2451  C   SER A 151       9.109 -20.488  23.424  1.00  0.00           C
ATOM   2452  O   SER A 151       8.117 -21.201  23.281  1.00  0.00           O
ATOM   2453  N   GLY A 152       9.452 -19.520  22.552  1.00  0.00           N
ATOM   2454  HN  GLY A 152      10.183 -18.879  22.786  1.00  0.00
ATOM   2455  CA  GLY A 152       8.765 -19.398  21.268  1.00  0.00           C
ATOM   2456  HA1 GLY A 152       9.468 -19.274  20.568  1.00  0.00
ATOM   2457  HA2 GLY A 152       8.284 -20.260  21.108  1.00  0.00
ATOM   2458  C   GLY A 152       7.766 -18.239  21.194  1.00  0.00           C
ATOM   2459  O   GLY A 152       7.207 -17.984  20.121  1.00  0.00           O
ATOM   2460  N   ILE A 153       7.514 -17.559  22.324  1.00  0.00           N
ATOM   2461  HN  ILE A 153       8.041 -17.780  23.145  1.00  0.00
ATOM   2462  CA  ILE A 153       6.508 -16.516  22.402  1.00  0.00           C
ATOM   2463  HA  ILE A 153       5.795 -16.773  21.750  1.00  0.00
ATOM   2464  CB  ILE A 153       5.875 -16.442  23.800  1.00  0.00           C
ATOM   2465  HB  ILE A 153       6.609 -16.413  24.479  1.00  0.00
ATOM   2466  CG2 ILE A 153       5.064 -15.167  23.955  1.00  0.00           C
ATOM   2467 1HG2 ILE A 153       4.661 -15.137  24.870  1.00  0.00
ATOM   2468 2HG2 ILE A 153       5.661 -14.375  23.828  1.00  0.00
ATOM   2469 3HG2 ILE A 153       4.336 -15.149  23.269  1.00  0.00
ATOM   2470  CG1 ILE A 153       5.032 -17.683  24.099  1.00  0.00           C
ATOM   2471 1HG1 ILE A 153       4.167 -17.599  23.604  1.00  0.00
ATOM   2472 2HG1 ILE A 153       5.529 -18.484  23.767  1.00  0.00
ATOM   2473  CD  ILE A 153       4.715 -17.897  25.583  1.00  0.00           C
ATOM   2474  HD1 ILE A 153       4.164 -18.725  25.690  1.00  0.00
ATOM   2475  HD2 ILE A 153       5.568 -17.996  26.095  1.00  0.00
ATOM   2476  HD3 ILE A 153       4.206 -17.110  25.932  1.00  0.00
ATOM   2477  C   ILE A 153       7.148 -15.182  22.009  1.00  0.00           C
ATOM   2478  O   ILE A 153       8.228 -14.847  22.481  1.00  0.00           O
ATOM   2479  N   ARG A 154       6.438 -14.431  21.156  1.00  0.00           N
ATOM   2480  HN  ARG A 154       5.615 -14.825  20.747  1.00  0.00
```

```
ATOM   2481  CA  ARG A 154       6.801 -13.088  20.800  1.00  0.00           C
ATOM   2482  HA  ARG A 154       7.674 -12.899  21.250  1.00  0.00
ATOM   2483  CB  ARG A 154       7.017 -12.977  19.293  1.00  0.00           C
ATOM   2484  HB1 ARG A 154       7.649 -13.697  19.007  1.00  0.00
ATOM   2485  HB2 ARG A 154       6.138 -13.104  18.834  1.00  0.00
ATOM   2486  CG  ARG A 154       7.593 -11.632  18.871  1.00  0.00           C
ATOM   2487  HG1 ARG A 154       6.910 -10.923  19.047  1.00  0.00
ATOM   2488  HG2 ARG A 154       8.411 -11.450  19.417  1.00  0.00
ATOM   2489  CD  ARG A 154       7.976 -11.566  17.413  1.00  0.00           C
ATOM   2490  HD1 ARG A 154       7.477 -12.273  16.912  1.00  0.00
ATOM   2491  HD2 ARG A 154       7.732 -10.666  17.051  1.00  0.00
ATOM   2492  NE  ARG A 154       9.399 -11.770  17.164  1.00  0.00           N
ATOM   2493  HE  ARG A 154      10.040 -11.599  17.913  1.00  0.00
ATOM   2494  CZ  ARG A 154       9.889 -12.170  15.998  1.00  0.00           C
ATOM   2495  NH1 ARG A 154       9.152 -12.031  14.900  1.00  0.00           N
ATOM   2496 1HH1 ARG A 154       8.238 -11.628  14.959  1.00  0.00
ATOM   2497 2HH1 ARG A 154       9.511 -12.329  14.016  1.00  0.00
ATOM   2498  NH2 ARG A 154      11.088 -12.735  15.950  1.00  0.00           N
ATOM   2499 1HH2 ARG A 154      11.617 -12.857  16.790  1.00  0.00
ATOM   2500 2HH2 ARG A 154      11.463 -13.039  15.074  1.00  0.00
ATOM   2501  C   ARG A 154       5.727 -12.120  21.322  1.00  0.00           C
ATOM   2502  O   ARG A 154       4.531 -12.383  21.222  1.00  0.00           O
ATOM   2503  N   ILE A 155       6.198 -11.011  21.902  1.00  0.00           N
ATOM   2504  HN  ILE A 155       7.179 -10.834  21.827  1.00  0.00
ATOM   2505  CA  ILE A 155       5.395 -10.039  22.636  1.00  0.00           C
ATOM   2506  HA  ILE A 155       4.549 -10.492  22.917  1.00  0.00
ATOM   2507  CB  ILE A 155       6.183  -9.557  23.865  1.00  0.00           C
ATOM   2508  HB  ILE A 155       7.102  -9.304  23.562  1.00  0.00
ATOM   2509  CG2 ILE A 155       5.582  -8.297  24.468  1.00  0.00           C
ATOM   2510 1HG2 ILE A 155       6.122  -8.017  25.262  1.00  0.00
ATOM   2511 2HG2 ILE A 155       5.588  -7.565  23.786  1.00  0.00
ATOM   2512 3HG2 ILE A 155       4.641  -8.480  24.752  1.00  0.00
ATOM   2513  CG1 ILE A 155       6.334 -10.676  24.899  1.00  0.00           C
ATOM   2514 1HG1 ILE A 155       6.723 -11.477  24.444  1.00  0.00
ATOM   2515 2HG1 ILE A 155       6.958 -10.363  25.615  1.00  0.00
ATOM   2516  CD  ILE A 155       5.032 -11.096  25.560  1.00  0.00           C
ATOM   2517  HD1 ILE A 155       5.213 -11.827  26.218  1.00  0.00
ATOM   2518  HD2 ILE A 155       4.631 -10.312  26.034  1.00  0.00
ATOM   2519  HD3 ILE A 155       4.395 -11.426  24.863  1.00  0.00
ATOM   2520  C   ILE A 155       5.083  -8.865  21.717  1.00  0.00           C
ATOM   2521  O   ILE A 155       5.960  -8.414  20.947  1.00  0.00           O
ATOM   2522  N   LYS A 156       3.857  -8.338  21.819  1.00  0.00           N
ATOM   2523  HN  LYS A 156       3.206  -8.721  22.474  1.00  0.00
ATOM   2524  CA  LYS A 156       3.461  -7.204  20.979  1.00  0.00           C
ATOM   2525  HA  LYS A 156       4.314  -6.786  20.665  1.00  0.00
ATOM   2526  CB  LYS A 156       2.644  -7.674  19.770  1.00  0.00           C
ATOM   2527  HB1 LYS A 156       1.693  -7.771  20.064  1.00  0.00
ATOM   2528  HB2 LYS A 156       2.705  -6.966  19.067  1.00  0.00
ATOM   2529  CG  LYS A 156       3.076  -9.001  19.136  1.00  0.00           C
ATOM   2530  HG1 LYS A 156       3.640  -9.515  19.783  1.00  0.00
ATOM   2531  HG2 LYS A 156       2.267  -9.539  18.898  1.00  0.00
ATOM   2532  CD  LYS A 156       3.862  -8.808  17.911  1.00  0.00           C
ATOM   2533  HD1 LYS A 156       3.343  -8.199  17.312  1.00  0.00
ATOM   2534  HD2 LYS A 156       4.722  -8.368  18.170  1.00  0.00
ATOM   2535  CE  LYS A 156       4.174 -10.077  17.160  1.00  0.00           C
ATOM   2536  HE1 LYS A 156       4.733 -10.673  17.737  1.00  0.00
ATOM   2537  HE2 LYS A 156       3.319 -10.542  16.930  1.00  0.00
ATOM   2538  NZ  LYS A 156       4.921  -9.784  15.899  1.00  0.00           N
ATOM   2539  HZ1 LYS A 156       5.116 -10.640  15.420  1.00  0.00
ATOM   2540  HZ2 LYS A 156       4.367  -9.193  15.313  1.00  0.00
ATOM   2541  HZ3 LYS A 156       5.780  -9.323  16.120  1.00  0.00
ATOM   2542  C   LYS A 156       2.637  -6.210  21.806  1.00  0.00           C
ATOM   2543  O   LYS A 156       1.529  -6.549  22.248  1.00  0.00           O
ATOM   2544  N   ILE A 157       3.164  -4.980  21.931  1.00  0.00           N
ATOM   2545  HN  ILE A 157       4.080  -4.820  21.564  1.00  0.00
ATOM   2546  CA  ILE A 157       2.472  -3.859  22.575  1.00  0.00           C
ATOM   2547  HA  ILE A 157       2.129  -4.199  23.451  1.00  0.00
ATOM   2548  CB  ILE A 157       3.448  -2.682  22.816  1.00  0.00           C
ATOM   2549  HB  ILE A 157       3.814  -2.403  21.928  1.00  0.00
ATOM   2550  CG2 ILE A 157       2.734  -1.476  23.398  1.00  0.00           C
ATOM   2551 1HG2 ILE A 157       3.392  -0.736  23.541  1.00  0.00
```

```
ATOM   2552 2HG2 ILE A 157       2.023  -1.172  22.764  1.00  0.00
ATOM   2553 3HG2 ILE A 157       2.317  -1.725  24.272  1.00  0.00
ATOM   2554  CG1 ILE A 157       4.645  -3.087  23.671  1.00  0.00           C
ATOM   2555 1HG1 ILE A 157       5.320  -3.540  23.089  1.00  0.00
ATOM   2556 2HG1 ILE A 157       5.049  -2.265  24.073  1.00  0.00
ATOM   2557  CD  ILE A 157       4.286  -4.023  24.785  1.00  0.00           C
ATOM   2558  HD1 ILE A 157       5.108  -4.251  25.307  1.00  0.00
ATOM   2559  HD2 ILE A 157       3.620  -3.584  25.388  1.00  0.00
ATOM   2560  HD3 ILE A 157       3.891  -4.859  24.404  1.00  0.00
ATOM   2561  C   ILE A 157       1.311  -3.412  21.683  1.00  0.00           C
ATOM   2562  O   ILE A 157       1.486  -3.217  20.495  1.00  0.00           O
ATOM   2563  N   ALA A 158       0.162  -3.144  22.306  1.00  0.00           N
ATOM   2564  HN  ALA A 158       0.138  -3.218  23.303  1.00  0.00
ATOM   2565  CA  ALA A 158      -1.053  -2.751  21.620  1.00  0.00           C
ATOM   2566  HA  ALA A 158      -0.799  -2.426  20.709  1.00  0.00
ATOM   2567  CB  ALA A 158      -1.936  -3.955  21.488  1.00  0.00           C
ATOM   2568  HB1 ALA A 158      -2.780  -3.699  21.016  1.00  0.00
ATOM   2569  HB2 ALA A 158      -1.461  -4.661  20.963  1.00  0.00
ATOM   2570  HB3 ALA A 158      -2.157  -4.308  22.397  1.00  0.00
ATOM   2571  C   ALA A 158      -1.754  -1.626  22.389  1.00  0.00           C
ATOM   2572  O   ALA A 158      -1.359  -1.324  23.489  1.00  0.00           O
ATOM   2573  N   ASP A 159      -2.770  -1.020  21.760  1.00  0.00           N
ATOM   2574  HN  ASP A 159      -2.914  -1.257  20.799  1.00  0.00
ATOM   2575  CA  ASP A 159      -3.702  -0.029  22.355  1.00  0.00           C
ATOM   2576  HA  ASP A 159      -4.414   0.095  21.664  1.00  0.00
ATOM   2577  CB  ASP A 159      -4.369  -0.528  23.638  1.00  0.00           C
ATOM   2578  HB1 ASP A 159      -4.684  -1.465  23.487  1.00  0.00
ATOM   2579  HB2 ASP A 159      -3.690  -0.519  24.372  1.00  0.00
ATOM   2580  CG  ASP A 159      -5.565   0.322  24.074  1.00  0.00           C
ATOM   2581  OD1 ASP A 159      -5.744   1.441  23.481  1.00  0.00           O
ATOM   2582  OD2 ASP A 159      -6.330  -0.136  24.983  1.00  0.00           O
ATOM   2583  C   ASP A 159      -2.984   1.290  22.622  1.00  0.00           C
ATOM   2584  O   ASP A 159      -2.600   1.562  23.738  1.00  0.00           O
ATOM   2585  N   PHE A 160      -2.890   2.134  21.600  1.00  0.00           N
ATOM   2586  HN  PHE A 160      -3.252   1.855  20.710  1.00  0.00
ATOM   2587  CA  PHE A 160      -2.287   3.434  21.727  1.00  0.00           C
ATOM   2588  HA  PHE A 160      -1.725   3.433  22.554  1.00  0.00
ATOM   2589  CB  PHE A 160      -1.373   3.674  20.527  1.00  0.00           C
ATOM   2590  HB1 PHE A 160      -1.878   3.467  19.689  1.00  0.00
ATOM   2591  HB2 PHE A 160      -1.094   4.634  20.519  1.00  0.00
ATOM   2592  CG  PHE A 160      -0.131   2.820  20.555  1.00  0.00           C
ATOM   2593  CD1 PHE A 160      -0.156   1.512  20.133  1.00  0.00           C
ATOM   2594  HD1 PHE A 160      -1.007   1.122  19.782  1.00  0.00
ATOM   2595  CE1 PHE A 160       0.983   0.729  20.189  1.00  0.00           C
ATOM   2596  HE1 PHE A 160       0.945  -0.225  19.892  1.00  0.00
ATOM   2597  CZ  PHE A 160       2.164   1.253  20.649  1.00  0.00           C
ATOM   2598  HZ  PHE A 160       2.991   0.692  20.667  1.00  0.00
ATOM   2599  CD2 PHE A 160       1.049   3.311  21.072  1.00  0.00           C
ATOM   2600  HD2 PHE A 160       1.073   4.238  21.447  1.00  0.00
ATOM   2601  CE2 PHE A 160       2.201   2.543  21.081  1.00  0.00           C
ATOM   2602  HE2 PHE A 160       3.061   2.937  21.405  1.00  0.00
ATOM   2603  C   PHE A 160      -3.374   4.495  21.911  1.00  0.00           C
ATOM   2604  O   PHE A 160      -3.138   5.670  21.620  1.00  0.00           O
ATOM   2605  N   GLY A 161      -4.527   4.082  22.460  1.00  0.00           N
ATOM   2606  HN  GLY A 161      -4.605   3.124  22.734  1.00  0.00
ATOM   2607  CA  GLY A 161      -5.666   4.953  22.676  1.00  0.00           C
ATOM   2608  HA1 GLY A 161      -5.947   5.302  21.782  1.00  0.00
ATOM   2609  HA2 GLY A 161      -6.400   4.393  23.061  1.00  0.00
ATOM   2610  C   GLY A 161      -5.357   6.125  23.619  1.00  0.00           C
ATOM   2611  O   GLY A 161      -5.911   7.212  23.455  1.00  0.00           O
ATOM   2612  N   PHE A 162      -4.474   5.910  24.597  1.00  0.00           N
ATOM   2613  HN  PHE A 162      -3.999   5.030  24.631  1.00  0.00
ATOM   2614  CA  PHE A 162      -4.176   6.902  25.612  1.00  0.00           C
ATOM   2615  HA  PHE A 162      -4.896   7.594  25.550  1.00  0.00
ATOM   2616  CB  PHE A 162      -4.212   6.243  27.000  1.00  0.00           C
ATOM   2617  HB1 PHE A 162      -4.828   5.456  26.961  1.00  0.00
ATOM   2618  HB2 PHE A 162      -3.289   5.933  27.227  1.00  0.00
ATOM   2619  CG  PHE A 162      -4.682   7.134  28.122  1.00  0.00           C
ATOM   2620  CD1 PHE A 162      -6.001   7.572  28.175  1.00  0.00           C
ATOM   2621  HD1 PHE A 162      -6.649   7.276  27.473  1.00  0.00
ATOM   2622  CE1 PHE A 162      -6.428   8.417  29.193  1.00  0.00           C
```

```
ATOM   2623  HE1 PHE A 162      -7.376   8.735  29.211  1.00  0.00
ATOM   2624  CZ  PHE A 162      -5.549   8.820  30.181  1.00  0.00           C
ATOM   2625  HZ  PHE A 162      -5.861   9.420  30.918  1.00  0.00
ATOM   2626  CD2 PHE A 162      -3.811   7.536  29.133  1.00  0.00           C
ATOM   2627  HD2 PHE A 162      -2.867   7.207  29.132  1.00  0.00
ATOM   2628  CE2 PHE A 162      -4.241   8.391  30.145  1.00  0.00           C
ATOM   2629  HE2 PHE A 162      -3.598   8.694  30.848  1.00  0.00
ATOM   2630  C   PHE A 162      -2.814   7.565  25.327  1.00  0.00           C
ATOM   2631  O   PHE A 162      -2.358   8.384  26.087  1.00  0.00           O
ATOM   2632  N   ALA A 163      -2.181   7.206  24.205  1.00  0.00           N
ATOM   2633  HN  ALA A 163      -2.680   6.662  23.531  1.00  0.00
ATOM   2634  CA  ALA A 163      -0.789   7.572  23.915  1.00  0.00           C
ATOM   2635  HA  ALA A 163      -0.240   7.384  24.729  1.00  0.00
ATOM   2636  CB  ALA A 163      -0.276   6.721  22.789  1.00  0.00           C
ATOM   2637  HB1 ALA A 163       0.672   6.968  22.589  1.00  0.00
ATOM   2638  HB2 ALA A 163      -0.321   5.757  23.052  1.00  0.00
ATOM   2639  HB3 ALA A 163      -0.838   6.871  21.975  1.00  0.00
ATOM   2640  C   ALA A 163      -0.700   9.059  23.580  1.00  0.00           C
ATOM   2641  O   ALA A 163      -1.696   9.690  23.210  1.00  0.00           O
ATOM   2642  N   ARG A 164       0.499   9.631  23.718  1.00  0.00           N
ATOM   2643  HN  ARG A 164       1.262   9.098  24.084  1.00  0.00
ATOM   2644  CA  ARG A 164       0.706  11.004  23.347  1.00  0.00           C
ATOM   2645  HA  ARG A 164       0.145  11.181  22.538  1.00  0.00
ATOM   2646  CB  ARG A 164       0.214  11.918  24.472  1.00  0.00           C
ATOM   2647  HB1 ARG A 164      -0.251  12.697  24.052  1.00  0.00
ATOM   2648  HB2 ARG A 164      -0.438  11.399  25.025  1.00  0.00
ATOM   2649  CG  ARG A 164       1.273  12.470  25.413  1.00  0.00           C
ATOM   2650  HG1 ARG A 164       1.665  11.717  25.942  1.00  0.00
ATOM   2651  HG2 ARG A 164       1.993  12.909  24.874  1.00  0.00
ATOM   2652  CD  ARG A 164       0.645  13.499  26.368  1.00  0.00           C
ATOM   2653  HD1 ARG A 164       0.253  14.256  25.844  1.00  0.00
ATOM   2654  HD2 ARG A 164      -0.072  13.065  26.914  1.00  0.00
ATOM   2655  NE  ARG A 164       1.630  14.059  27.291  1.00  0.00           N
ATOM   2656  HE  ARG A 164       1.611  13.744  28.240  1.00  0.00
ATOM   2657  CZ  ARG A 164       2.550  14.958  26.956  1.00  0.00           C
ATOM   2658  NH1 ARG A 164       2.395  15.689  25.874  1.00  0.00           N
ATOM   2659 1HH1 ARG A 164       1.584  15.569  25.302  1.00  0.00
ATOM   2660 2HH1 ARG A 164       3.088  16.365  25.624  1.00  0.00
ATOM   2661  NH2 ARG A 164       3.677  15.044  27.635  1.00  0.00           N
ATOM   2662 1HH2 ARG A 164       3.844  14.431  28.407  1.00  0.00
ATOM   2663 2HH2 ARG A 164       4.365  15.723  27.378  1.00  0.00
ATOM   2664  C   ARG A 164       2.168  11.223  22.950  1.00  0.00           C
ATOM   2665  O   ARG A 164       3.024  10.396  23.225  1.00  0.00           O
ATOM   2666  N   TYR A 165       2.389  12.293  22.178  1.00  0.00           N
ATOM   2667  HN  TYR A 165       1.599  12.834  21.888  1.00  0.00
ATOM   2668  CA  TYR A 165       3.691  12.713  21.740  1.00  0.00           C
ATOM   2669  HA  TYR A 165       4.227  11.870  21.689  1.00  0.00
ATOM   2670  CB  TYR A 165       3.615  13.394  20.372  1.00  0.00           C
ATOM   2671  HB1 TYR A 165       2.875  14.067  20.397  1.00  0.00
ATOM   2672  HB2 TYR A 165       4.484  13.860  20.205  1.00  0.00
ATOM   2673  CG  TYR A 165       3.359  12.491  19.201  1.00  0.00           C
ATOM   2674  CD1 TYR A 165       4.394  11.808  18.595  1.00  0.00           C
ATOM   2675  HD1 TYR A 165       5.322  11.906  18.954  1.00  0.00
ATOM   2676  CE1 TYR A 165       4.174  10.990  17.500  1.00  0.00           C
ATOM   2677  HE1 TYR A 165       4.939  10.504  17.077  1.00  0.00
ATOM   2678  CZ  TYR A 165       2.900  10.846  16.993  1.00  0.00           C
ATOM   2679  OH  TYR A 165       2.685  10.040  15.915  1.00  0.00           O
ATOM   2680  HH  TYR A 165       1.713  10.050  15.680  1.00  0.00
ATOM   2681  CD2 TYR A 165       2.093  12.371  18.651  1.00  0.00           C
ATOM   2682  HD2 TYR A 165       1.337  12.894  19.044  1.00  0.00
ATOM   2683  CE2 TYR A 165       1.853  11.540  17.566  1.00  0.00           C
ATOM   2684  HE2 TYR A 165       0.927  11.444  17.201  1.00  0.00
ATOM   2685  C   TYR A 165       4.264  13.717  22.747  1.00  0.00           C
ATOM   2686  O   TYR A 165       3.546  14.631  23.151  1.00  0.00           O
ATOM   2687  N   LEU A 166       5.558  13.568  23.066  1.00  0.00           N
ATOM   2688  HN  LEU A 166       6.028  12.741  22.758  1.00  0.00
ATOM   2689  CA  LEU A 166       6.315  14.542  23.835  1.00  0.00           C
ATOM   2690  HA  LEU A 166       5.736  14.790  24.611  1.00  0.00
ATOM   2691  CB  LEU A 166       7.633  13.913  24.289  1.00  0.00           C
ATOM   2692  HB1 LEU A 166       7.985  13.352  23.540  1.00  0.00
ATOM   2693  HB2 LEU A 166       8.279  14.650  24.486  1.00  0.00
```

| | | | | | | | | | | | |
|---|---|---|---|---|---|---|---|---|---|---|---|
| ATOM | 2694 | CG | LEU | A | 166 | 7.543 | 13.035 | 25.522 | 1.00 | 0.00 | C |
| ATOM | 2695 | HG | LEU | A | 166 | 6.941 | 12.263 | 25.318 | 1.00 | 0.00 | |
| ATOM | 2696 | CD1 | LEU | A | 166 | 8.909 | 12.459 | 25.883 | 1.00 | 0.00 | C |
| ATOM | 2697 | 1HD1 | LEU | A | 166 | 8.824 | 11.885 | 26.697 | 1.00 | 0.00 | |
| ATOM | 2698 | 2HD1 | LEU | A | 166 | 9.250 | 11.908 | 25.121 | 1.00 | 0.00 | |
| ATOM | 2699 | 3HD1 | LEU | A | 166 | 9.548 | 13.206 | 26.067 | 1.00 | 0.00 | |
| ATOM | 2700 | CD2 | LEU | A | 166 | 6.938 | 13.824 | 26.683 | 1.00 | 0.00 | C |
| ATOM | 2701 | 1HD2 | LEU | A | 166 | 6.882 | 13.238 | 27.491 | 1.00 | 0.00 | |
| ATOM | 2702 | 2HD2 | LEU | A | 166 | 7.515 | 14.616 | 26.884 | 1.00 | 0.00 | |
| ATOM | 2703 | 3HD2 | LEU | A | 166 | 6.021 | 14.135 | 26.433 | 1.00 | 0.00 | |
| ATOM | 2704 | C | LEU | A | 166 | 6.626 | 15.771 | 22.981 | 1.00 | 0.00 | C |
| ATOM | 2705 | O | LEU | A | 166 | 7.301 | 15.647 | 21.974 | 1.00 | 0.00 | O |
| ATOM | 2706 | N | HIS | A | 167 | 6.246 | 16.948 | 23.487 | 1.00 | 0.00 | N |
| ATOM | 2707 | HN | HIS | A | 167 | 5.810 | 16.957 | 24.387 | 1.00 | 0.00 | |
| ATOM | 2708 | CA | HIS | A | 167 | 6.431 | 18.213 | 22.802 | 1.00 | 0.00 | C |
| ATOM | 2709 | HA | HIS | A | 167 | 6.363 | 18.076 | 21.814 | 1.00 | 0.00 | |
| ATOM | 2710 | CB | HIS | A | 167 | 5.287 | 19.175 | 23.190 | 1.00 | 0.00 | C |
| ATOM | 2711 | HB1 | HIS | A | 167 | 5.320 | 19.346 | 24.175 | 1.00 | 0.00 | |
| ATOM | 2712 | HB2 | HIS | A | 167 | 5.402 | 20.038 | 22.698 | 1.00 | 0.00 | |
| ATOM | 2713 | ND1 | HIS | A | 167 | 3.590 | 18.191 | 21.588 | 1.00 | 0.00 | N |
| ATOM | 2714 | CG | HIS | A | 167 | 3.928 | 18.640 | 22.864 | 1.00 | 0.00 | C |
| ATOM | 2715 | CE1 | HIS | A | 167 | 2.343 | 17.765 | 21.583 | 1.00 | 0.00 | C |
| ATOM | 2716 | HE1 | HIS | A | 167 | 1.851 | 17.394 | 20.795 | 1.00 | 0.00 | |
| ATOM | 2717 | NE2 | HIS | A | 167 | 1.851 | 17.917 | 22.821 | 1.00 | 0.00 | N |
| ATOM | 2718 | HE2 | HIS | A | 167 | 0.926 | 17.674 | 23.112 | 1.00 | 0.00 | |
| ATOM | 2719 | CD2 | HIS | A | 167 | 2.823 | 18.461 | 23.625 | 1.00 | 0.00 | C |
| ATOM | 2720 | HD2 | HIS | A | 167 | 2.731 | 18.684 | 24.595 | 1.00 | 0.00 | |
| ATOM | 2721 | C | HIS | A | 167 | 7.854 | 18.705 | 23.095 | 1.00 | 0.00 | C |
| ATOM | 2722 | O | HIS | A | 167 | 8.045 | 19.552 | 23.938 | 1.00 | 0.00 | O |
| ATOM | 2723 | N | SER | A | 168 | 8.851 | 18.235 | 22.330 | 1.00 | 0.00 | N |
| ATOM | 2724 | HN | SER | A | 168 | 8.655 | 17.568 | 21.611 | 1.00 | 0.00 | |
| ATOM | 2725 | CA | SER | A | 168 | 10.191 | 18.683 | 22.538 | 1.00 | 0.00 | C |
| ATOM | 2726 | HA | SER | A | 168 | 10.140 | 19.553 | 23.028 | 1.00 | 0.00 | |
| ATOM | 2727 | CB | SER | A | 168 | 11.044 | 17.698 | 23.355 | 1.00 | 0.00 | C |
| ATOM | 2728 | HB1 | SER | A | 168 | 11.963 | 18.064 | 23.504 | 1.00 | 0.00 | |
| ATOM | 2729 | HB2 | SER | A | 168 | 10.615 | 17.506 | 24.238 | 1.00 | 0.00 | |
| ATOM | 2730 | OG | SER | A | 168 | 11.185 | 16.461 | 22.673 | 1.00 | 0.00 | O |
| ATOM | 2731 | HG1 | SER | A | 168 | 11.742 | 15.839 | 23.223 | 1.00 | 0.00 | |
| ATOM | 2732 | C | SER | A | 168 | 10.756 | 18.773 | 21.172 | 1.00 | 0.00 | C |
| ATOM | 2733 | O | SER | A | 168 | 10.035 | 19.099 | 20.229 | 1.00 | 0.00 | O |
| ATOM | 2734 | N | ASN | A | 169 | 12.074 | 18.513 | 21.028 | 1.00 | 0.00 | N |
| ATOM | 2735 | HN | ASN | A | 169 | 12.645 | 18.373 | 21.837 | 1.00 | 0.00 | |
| ATOM | 2736 | CA | ASN | A | 169 | 12.657 | 18.437 | 19.719 | 1.00 | 0.00 | C |
| ATOM | 2737 | HA | ASN | A | 169 | 12.550 | 19.330 | 19.282 | 1.00 | 0.00 | |
| ATOM | 2738 | CB | ASN | A | 169 | 14.154 | 18.082 | 19.764 | 1.00 | 0.00 | C |
| ATOM | 2739 | HB1 | ASN | A | 169 | 14.508 | 17.951 | 18.838 | 1.00 | 0.00 | |
| ATOM | 2740 | HB2 | ASN | A | 169 | 14.670 | 18.808 | 20.218 | 1.00 | 0.00 | |
| ATOM | 2741 | CG | ASN | A | 169 | 14.276 | 16.785 | 20.555 | 1.00 | 0.00 | C |
| ATOM | 2742 | OD1 | ASN | A | 169 | 14.083 | 16.764 | 21.772 | 1.00 | 0.00 | O |
| ATOM | 2743 | ND2 | ASN | A | 169 | 14.532 | 15.662 | 19.834 | 1.00 | 0.00 | N |
| ATOM | 2744 | 1HD2 | ASN | A | 169 | 14.629 | 15.717 | 18.840 | 1.00 | 0.00 | |
| ATOM | 2745 | 2HD2 | ASN | A | 169 | 14.623 | 14.780 | 20.297 | 1.00 | 0.00 | |
| ATOM | 2746 | C | ASN | A | 169 | 11.910 | 17.326 | 19.045 | 1.00 | 0.00 | C |
| ATOM | 2747 | O | ASN | A | 169 | 11.781 | 16.225 | 19.586 | 1.00 | 0.00 | O |
| ATOM | 2748 | N | MET | A | 170 | 11.333 | 17.611 | 17.865 | 1.00 | 0.00 | N |
| ATOM | 2749 | HN | MET | A | 170 | 11.481 | 18.480 | 17.394 | 1.00 | 0.00 | |
| ATOM | 2750 | CA | MET | A | 170 | 10.495 | 16.587 | 17.332 | 1.00 | 0.00 | C |
| ATOM | 2751 | HA | MET | A | 170 | 10.944 | 15.747 | 17.637 | 1.00 | 0.00 | |
| ATOM | 2752 | CB | MET | A | 170 | 9.047 | 16.660 | 17.849 | 1.00 | 0.00 | C |
| ATOM | 2753 | HB1 | MET | A | 170 | 9.049 | 16.959 | 18.803 | 1.00 | 0.00 | |
| ATOM | 2754 | HB2 | MET | A | 170 | 8.529 | 17.313 | 17.296 | 1.00 | 0.00 | |
| ATOM | 2755 | CG | MET | A | 170 | 8.375 | 15.299 | 17.761 | 1.00 | 0.00 | C |
| ATOM | 2756 | HG1 | MET | A | 170 | 8.352 | 15.123 | 16.777 | 1.00 | 0.00 | |
| ATOM | 2757 | HG2 | MET | A | 170 | 9.041 | 14.691 | 18.193 | 1.00 | 0.00 | |
| ATOM | 2758 | SD | MET | A | 170 | 6.727 | 15.127 | 18.511 | 1.00 | 0.00 | S |
| ATOM | 2759 | CE | MET | A | 170 | 7.091 | 13.405 | 18.965 | 1.00 | 0.00 | C |
| ATOM | 2760 | HE1 | MET | A | 170 | 6.301 | 13.010 | 19.434 | 1.00 | 0.00 | |
| ATOM | 2761 | HE2 | MET | A | 170 | 7.286 | 12.877 | 18.139 | 1.00 | 0.00 | |
| ATOM | 2762 | HE3 | MET | A | 170 | 7.886 | 13.381 | 19.572 | 1.00 | 0.00 | |
| ATOM | 2763 | C | MET | A | 170 | 10.460 | 16.698 | 15.843 | 1.00 | 0.00 | C |
| ATOM | 2764 | O | MET | A | 170 | 11.378 | 17.241 | 15.233 | 1.00 | 0.00 | O |

```
ATOM   2765  N   MET A 171       9.362  16.189  15.244  1.00  0.00           N
ATOM   2766  HN  MET A 171       8.615  15.879  15.832  1.00  0.00
ATOM   2767  CA  MET A 171       9.189  16.061  13.823  1.00  0.00           C
ATOM   2768  HA  MET A 171       9.910  15.430  13.535  1.00  0.00
ATOM   2769  CB  MET A 171       7.787  15.517  13.479  1.00  0.00           C
ATOM   2770  HB1 MET A 171       7.106  16.191  13.764  1.00  0.00
ATOM   2771  HB2 MET A 171       7.731  15.385  12.489  1.00  0.00
ATOM   2772  CG  MET A 171       7.465  14.179  14.167  1.00  0.00           C
ATOM   2773  HG1 MET A 171       8.197  13.567  13.870  1.00  0.00
ATOM   2774  HG2 MET A 171       7.568  14.373  15.143  1.00  0.00
ATOM   2775  SD  MET A 171       5.833  13.498  13.791  1.00  0.00           S
ATOM   2776  CE  MET A 171       5.816  12.390  15.224  1.00  0.00           C
ATOM   2777  HE1 MET A 171       4.960  11.873  15.235  1.00  0.00
ATOM   2778  HE2 MET A 171       6.587  11.756  15.165  1.00  0.00
ATOM   2779  HE3 MET A 171       5.891  12.929  16.063  1.00  0.00
ATOM   2780  C   MET A 171       9.358  17.405  13.169  1.00  0.00           C
ATOM   2781  O   MET A 171      10.069  17.521  12.172  1.00  0.00           O
ATOM   2782  N   ALA A 172       8.741  18.460  13.736  1.00  0.00           N
ATOM   2783  HN  ALA A 172       8.155  18.314  14.533  1.00  0.00
ATOM   2784  CA  ALA A 172       8.909  19.792  13.218  1.00  0.00           C
ATOM   2785  HA  ALA A 172       9.872  19.844  12.953  1.00  0.00
ATOM   2786  CB  ALA A 172       8.011  20.113  12.008  1.00  0.00           C
ATOM   2787  HB1 ALA A 172       8.182  21.050  11.705  1.00  0.00
ATOM   2788  HB2 ALA A 172       8.217  19.479  11.263  1.00  0.00
ATOM   2789  HB3 ALA A 172       7.051  20.018  12.271  1.00  0.00
ATOM   2790  C   ALA A 172       8.505  20.723  14.315  1.00  0.00           C
ATOM   2791  O   ALA A 172       7.658  20.377  15.136  1.00  0.00           O
ATOM   2792  N   ALA A 173       9.118  21.925  14.356  1.00  0.00           N
ATOM   2793  HN  ALA A 173       9.832  22.135  13.688  1.00  0.00
ATOM   2794  CA  ALA A 173       8.765  22.908  15.341  1.00  0.00           C
ATOM   2795  HA  ALA A 173       7.767  22.856  15.377  1.00  0.00
ATOM   2796  CB  ALA A 173       9.371  22.665  16.735  1.00  0.00           C
ATOM   2797  HB1 ALA A 173       9.077  23.391  17.357  1.00  0.00
ATOM   2798  HB2 ALA A 173       9.061  21.781  17.084  1.00  0.00
ATOM   2799  HB3 ALA A 173      10.369  22.666  16.669  1.00  0.00
ATOM   2800  C   ALA A 173       9.309  24.212  14.878  1.00  0.00           C
ATOM   2801  O   ALA A 173      10.210  24.266  14.043  1.00  0.00           O
ATOM   2802  N   ASP A 174       8.743  25.309  15.410  1.00  0.00           N
ATOM   2803  HN  ASP A 174       7.973  25.215  16.041  1.00  0.00
ATOM   2804  CA  ASP A 174       9.235  26.608  15.080  1.00  0.00           C
ATOM   2805  HA  ASP A 174       9.780  26.480  14.252  1.00  0.00
ATOM   2806  CB  ASP A 174       8.132  27.648  14.810  1.00  0.00           C
ATOM   2807  HB1 ASP A 174       7.471  27.649  15.560  1.00  0.00
ATOM   2808  HB2 ASP A 174       8.534  28.559  14.719  1.00  0.00
ATOM   2809  CG  ASP A 174       7.432  27.267  13.513  1.00  0.00           C
ATOM   2810  OD1 ASP A 174       7.757  26.183  12.959  1.00  0.00           O
ATOM   2811  OD2 ASP A 174       6.564  28.058  13.057  1.00  0.00           O
ATOM   2812  C   ASP A 174      10.002  27.061  16.278  1.00  0.00           C
ATOM   2813  O   ASP A 174       9.772  26.589  17.394  1.00  0.00           O
ATOM   2814  N   LEU A 175      10.956  27.999  16.089  1.00  0.00           N
ATOM   2815  HN  LEU A 175      11.126  28.369  15.176  1.00  0.00
ATOM   2816  CA  LEU A 175      11.727  28.462  17.217  1.00  0.00           C
ATOM   2817  HA  LEU A 175      12.135  27.630  17.593  1.00  0.00
ATOM   2818  CB  LEU A 175      12.838  29.470  16.851  1.00  0.00           C
ATOM   2819  HB1 LEU A 175      12.420  30.218  16.335  1.00  0.00
ATOM   2820  HB2 LEU A 175      13.224  29.827  17.701  1.00  0.00
ATOM   2821  CG  LEU A 175      13.994  28.893  16.006  1.00  0.00           C
ATOM   2822  HG  LEU A 175      13.618  28.594  15.129  1.00  0.00
ATOM   2823  CD1 LEU A 175      15.047  29.967  15.692  1.00  0.00           C
ATOM   2824 1HD1 LEU A 175      15.781  29.564  15.145  1.00  0.00
ATOM   2825 2HD1 LEU A 175      14.620  30.712  15.180  1.00  0.00
ATOM   2826 3HD1 LEU A 175      15.425  30.323  16.547  1.00  0.00
ATOM   2827  CD2 LEU A 175      14.608  27.649  16.667  1.00  0.00           C
ATOM   2828 1HD2 LEU A 175      15.352  27.301  16.096  1.00  0.00
ATOM   2829 2HD2 LEU A 175      14.966  27.893  17.568  1.00  0.00
ATOM   2830 3HD2 LEU A 175      13.906  26.944  16.767  1.00  0.00
ATOM   2831  C   LEU A 175      10.784  29.148  18.154  1.00  0.00           C
ATOM   2832  O   LEU A 175      10.241  30.209  17.847  1.00  0.00           O
ATOM   2833  N   CYS A 176      10.538  28.513  19.314  1.00  0.00           N
ATOM   2834  HN  CYS A 176      10.996  27.651  19.529  1.00  0.00
ATOM   2835  CA  CYS A 176       9.610  29.092  20.235  1.00  0.00           C
```

| | | | | | | | | | | | |
|---|---|---|---|---|---|---|---|---|---|---|---|
| ATOM | 2836 | HA | CYS | A | 176 | 9.530 | 30.049 | 19.957 | 1.00 | 0.00 | |
| ATOM | 2837 | CB | CYS | A | 176 | 8.237 | 28.390 | 20.197 | 1.00 | 0.00 | C |
| ATOM | 2838 | HB1 | CYS | A | 176 | 7.982 | 28.394 | 19.230 | 1.00 | 0.00 | |
| ATOM | 2839 | HB2 | CYS | A | 176 | 8.428 | 27.452 | 20.487 | 1.00 | 0.00 | |
| ATOM | 2840 | SG | CYS | A | 176 | 6.966 | 29.177 | 21.238 | 1.00 | 0.00 | S |
| ATOM | 2841 | HG1 | CYS | A | 176 | 6.109 | 28.667 | 21.161 | 1.00 | 0.00 | |
| ATOM | 2842 | C | CYS | A | 176 | 10.174 | 28.951 | 21.607 | 1.00 | 0.00 | C |
| ATOM | 2843 | O | CYS | A | 176 | 10.813 | 27.953 | 21.942 | 1.00 | 0.00 | O |
| ATOM | 2844 | N | GLY | A | 177 | 9.943 | 29.984 | 22.435 | 1.00 | 0.00 | N |
| ATOM | 2845 | HN | GLY | A | 177 | 9.485 | 30.793 | 22.068 | 1.00 | 0.00 | |
| ATOM | 2846 | CA | GLY | A | 177 | 10.321 | 29.985 | 23.821 | 1.00 | 0.00 | C |
| ATOM | 2847 | HA1 | GLY | A | 177 | 11.031 | 29.300 | 23.982 | 1.00 | 0.00 | |
| ATOM | 2848 | HA2 | GLY | A | 177 | 10.664 | 30.888 | 24.081 | 1.00 | 0.00 | |
| ATOM | 2849 | C | GLY | A | 177 | 9.062 | 29.648 | 24.551 | 1.00 | 0.00 | C |
| ATOM | 2850 | O | GLY | A | 177 | 8.315 | 28.758 | 24.143 | 1.00 | 0.00 | O |
| ATOM | 2851 | N | SER | A | 178 | 8.779 | 30.347 | 25.667 | 1.00 | 0.00 | N |
| ATOM | 2852 | HN | SER | A | 178 | 9.424 | 31.025 | 26.018 | 1.00 | 0.00 | |
| ATOM | 2853 | CA | SER | A | 178 | 7.532 | 30.103 | 26.347 | 1.00 | 0.00 | C |
| ATOM | 2854 | HA | SER | A | 178 | 7.308 | 29.160 | 26.102 | 1.00 | 0.00 | |
| ATOM | 2855 | CB | SER | A | 178 | 7.601 | 30.257 | 27.874 | 1.00 | 0.00 | C |
| ATOM | 2856 | HB1 | SER | A | 178 | 8.080 | 29.484 | 28.289 | 1.00 | 0.00 | |
| ATOM | 2857 | HB2 | SER | A | 178 | 8.061 | 31.108 | 28.126 | 1.00 | 0.00 | |
| ATOM | 2858 | OG | SER | A | 178 | 6.287 | 30.296 | 28.418 | 1.00 | 0.00 | O |
| ATOM | 2859 | HG1 | SER | A | 178 | 6.340 | 30.396 | 29.412 | 1.00 | 0.00 | |
| ATOM | 2860 | C | SER | A | 178 | 6.555 | 31.130 | 25.870 | 1.00 | 0.00 | C |
| ATOM | 2861 | O | SER | A | 178 | 6.912 | 32.294 | 25.706 | 1.00 | 0.00 | O |
| ATOM | 2862 | N | PRO | A | 179 | 5.353 | 30.704 | 25.556 | 1.00 | 0.00 | N |
| ATOM | 2863 | CD | PRO | A | 179 | 5.156 | 29.396 | 24.957 | 1.00 | 0.00 | C |
| ATOM | 2864 | HD1 | PRO | A | 179 | 5.328 | 28.684 | 25.638 | 1.00 | 0.00 | |
| ATOM | 2865 | HD2 | PRO | A | 179 | 5.783 | 29.279 | 24.187 | 1.00 | 0.00 | |
| ATOM | 2866 | CA | PRO | A | 179 | 4.337 | 31.634 | 25.163 | 1.00 | 0.00 | C |
| ATOM | 2867 | HA | PRO | A | 179 | 4.743 | 32.410 | 24.680 | 1.00 | 0.00 | |
| ATOM | 2868 | CB | PRO | A | 179 | 3.396 | 30.879 | 24.224 | 1.00 | 0.00 | C |
| ATOM | 2869 | HB1 | PRO | A | 179 | 2.442 | 31.094 | 24.433 | 1.00 | 0.00 | |
| ATOM | 2870 | HB2 | PRO | A | 179 | 3.589 | 31.110 | 23.270 | 1.00 | 0.00 | |
| ATOM | 2871 | CG | PRO | A | 179 | 3.696 | 29.398 | 24.499 | 1.00 | 0.00 | C |
| ATOM | 2872 | HG1 | PRO | A | 179 | 3.099 | 29.040 | 25.217 | 1.00 | 0.00 | |
| ATOM | 2873 | HG2 | PRO | A | 179 | 3.584 | 28.851 | 23.669 | 1.00 | 0.00 | |
| ATOM | 2874 | C | PRO | A | 179 | 3.701 | 32.247 | 26.377 | 1.00 | 0.00 | C |
| ATOM | 2875 | O | PRO | A | 179 | 3.287 | 31.523 | 27.285 | 1.00 | 0.00 | O |
| ATOM | 2876 | N | MET | A | 180 | 3.533 | 33.585 | 26.348 | 1.00 | 0.00 | N |
| ATOM | 2877 | HN | MET | A | 180 | 3.752 | 34.045 | 25.488 | 1.00 | 0.00 | |
| ATOM | 2878 | CA | MET | A | 180 | 3.061 | 34.439 | 27.449 | 1.00 | 0.00 | C |
| ATOM | 2879 | HA | MET | A | 180 | 3.661 | 34.230 | 28.221 | 1.00 | 0.00 | |
| ATOM | 2880 | CB | MET | A | 180 | 3.152 | 35.920 | 27.052 | 1.00 | 0.00 | C |
| ATOM | 2881 | HB1 | MET | A | 180 | 2.531 | 36.078 | 26.284 | 1.00 | 0.00 | |
| ATOM | 2882 | HB2 | MET | A | 180 | 2.863 | 36.473 | 27.833 | 1.00 | 0.00 | |
| ATOM | 2883 | CG | MET | A | 180 | 4.539 | 36.389 | 26.636 | 1.00 | 0.00 | C |
| ATOM | 2884 | HG1 | MET | A | 180 | 4.869 | 36.937 | 27.405 | 1.00 | 0.00 | |
| ATOM | 2885 | HG2 | MET | A | 180 | 5.084 | 35.554 | 26.564 | 1.00 | 0.00 | |
| ATOM | 2886 | SD | MET | A | 180 | 4.487 | 37.331 | 25.074 | 1.00 | 0.00 | S |
| ATOM | 2887 | CE | MET | A | 180 | 4.341 | 35.975 | 23.906 | 1.00 | 0.00 | C |
| ATOM | 2888 | HE1 | MET | A | 180 | 4.300 | 36.339 | 22.976 | 1.00 | 0.00 | |
| ATOM | 2889 | HE2 | MET | A | 180 | 5.135 | 35.373 | 23.993 | 1.00 | 0.00 | |
| ATOM | 2890 | HE3 | MET | A | 180 | 3.507 | 35.458 | 24.099 | 1.00 | 0.00 | |
| ATOM | 2891 | C | MET | A | 180 | 1.592 | 34.128 | 27.780 | 1.00 | 0.00 | C |
| ATOM | 2892 | O | MET | A | 180 | 1.142 | 34.341 | 28.913 | 1.00 | 0.00 | O |
| ATOM | 2893 | N | TYR | A | 181 | 0.868 | 33.610 | 26.782 | 1.00 | 0.00 | N |
| ATOM | 2894 | HN | TYR | A | 181 | 1.351 | 33.321 | 25.955 | 1.00 | 0.00 | |
| ATOM | 2895 | CA | TYR | A | 181 | -0.546 | 33.443 | 26.820 | 1.00 | 0.00 | C |
| ATOM | 2896 | HA | TYR | A | 181 | -0.881 | 34.156 | 27.436 | 1.00 | 0.00 | |
| ATOM | 2897 | CB | TYR | A | 181 | -1.093 | 33.619 | 25.416 | 1.00 | 0.00 | C |
| ATOM | 2898 | HB1 | TYR | A | 181 | -0.729 | 32.888 | 24.838 | 1.00 | 0.00 | |
| ATOM | 2899 | HB2 | TYR | A | 181 | -2.090 | 33.551 | 25.452 | 1.00 | 0.00 | |
| ATOM | 2900 | CG | TYR | A | 181 | -0.736 | 34.936 | 24.786 | 1.00 | 0.00 | C |
| ATOM | 2901 | CD1 | TYR | A | 181 | -1.446 | 36.075 | 25.114 | 1.00 | 0.00 | C |
| ATOM | 2902 | HD1 | TYR | A | 181 | -2.166 | 36.023 | 25.806 | 1.00 | 0.00 | |
| ATOM | 2903 | CE1 | TYR | A | 181 | -1.177 | 37.287 | 24.501 | 1.00 | 0.00 | C |
| ATOM | 2904 | HE1 | TYR | A | 181 | -1.724 | 38.094 | 24.723 | 1.00 | 0.00 | |
| ATOM | 2905 | CZ | TYR | A | 181 | -0.147 | 37.377 | 23.579 | 1.00 | 0.00 | C |
| ATOM | 2906 | OH | TYR | A | 181 | 0.119 | 38.582 | 22.999 | 1.00 | 0.00 | O |

```
ATOM   2907  HH  TYR A 181       0.884  38.484  22.362  1.00  0.00
ATOM   2908  CD2 TYR A 181       0.280  35.041  23.842  1.00  0.00           C
ATOM   2909  HD2 TYR A 181       0.796  34.224  23.584  1.00  0.00
ATOM   2910  CE2 TYR A 181       0.590  36.251  23.255  1.00  0.00           C
ATOM   2911  HE2 TYR A 181       1.343  36.314  22.600  1.00  0.00
ATOM   2912  C   TYR A 181      -0.922  32.056  27.360  1.00  0.00           C
ATOM   2913  O   TYR A 181      -2.084  31.792  27.646  1.00  0.00           O
ATOM   2914  N   MET A 182       0.072  31.183  27.513  1.00  0.00           N
ATOM   2915  HN  MET A 182       1.014  31.486  27.369  1.00  0.00
ATOM   2916  CA  MET A 182      -0.171  29.844  27.872  1.00  0.00           C
ATOM   2917  HA  MET A 182      -1.070  29.639  27.484  1.00  0.00
ATOM   2918  CB  MET A 182       0.900  28.943  27.269  1.00  0.00           C
ATOM   2919  HB1 MET A 182       1.125  29.281  26.355  1.00  0.00
ATOM   2920  HB2 MET A 182       1.714  28.982  27.849  1.00  0.00
ATOM   2921  CG  MET A 182       0.458  27.493  27.152  1.00  0.00           C
ATOM   2922  HG1 MET A 182       0.396  27.157  28.092  1.00  0.00
ATOM   2923  HG2 MET A 182      -0.456  27.525  26.748  1.00  0.00
ATOM   2924  SD  MET A 182       1.612  26.540  26.159  1.00  0.00           S
ATOM   2925  CE  MET A 182       1.362  27.253  24.529  1.00  0.00           C
ATOM   2926  HE1 MET A 182       1.961  26.798  23.870  1.00  0.00
ATOM   2927  HE2 MET A 182       0.409  27.129  24.254  1.00  0.00
ATOM   2928  HE3 MET A 182       1.577  28.229  24.556  1.00  0.00
ATOM   2929  C   MET A 182      -0.172  29.706  29.401  1.00  0.00           C
ATOM   2930  O   MET A 182       0.637  30.330  30.114  1.00  0.00           O
ATOM   2931  N   ALA A 183      -1.098  28.886  29.903  1.00  0.00           N
ATOM   2932  HN  ALA A 183      -1.660  28.356  29.268  1.00  0.00
ATOM   2933  CA  ALA A 183      -1.326  28.728  31.325  1.00  0.00           C
ATOM   2934  HA  ALA A 183      -1.525  29.634  31.698  1.00  0.00
ATOM   2935  CB  ALA A 183      -2.523  27.844  31.561  1.00  0.00           C
ATOM   2936  HB1 ALA A 183      -2.672  27.741  32.544  1.00  0.00
ATOM   2937  HB2 ALA A 183      -3.331  28.259  31.143  1.00  0.00
ATOM   2938  HB3 ALA A 183      -2.360  26.946  31.153  1.00  0.00
ATOM   2939  C   ALA A 183      -0.073  28.159  31.978  1.00  0.00           C
ATOM   2940  O   ALA A 183       0.548  27.247  31.441  1.00  0.00           O
ATOM   2941  N   PRO A 184       0.333  28.676  33.154  1.00  0.00           N
ATOM   2942  CD  PRO A 184      -0.342  29.750  33.893  1.00  0.00           C
ATOM   2943  HD1 PRO A 184      -1.136  29.393  34.385  1.00  0.00
ATOM   2944  HD2 PRO A 184      -0.634  30.476  33.270  1.00  0.00
ATOM   2945  CA  PRO A 184       1.551  28.219  33.810  1.00  0.00           C
ATOM   2946  HA  PRO A 184       2.327  28.395  33.204  1.00  0.00
ATOM   2947  CB  PRO A 184       1.562  29.008  35.125  1.00  0.00           C
ATOM   2948  HB1 PRO A 184       1.159  28.472  35.867  1.00  0.00
ATOM   2949  HB2 PRO A 184       2.493  29.274  35.374  1.00  0.00
ATOM   2950  CG  PRO A 184       0.721  30.233  34.841  1.00  0.00           C
ATOM   2951  HG1 PRO A 184       0.309  30.584  35.682  1.00  0.00
ATOM   2952  HG2 PRO A 184       1.272  30.951  34.415  1.00  0.00
ATOM   2953  C   PRO A 184       1.567  26.703  34.082  1.00  0.00           C
ATOM   2954  O   PRO A 184       2.624  26.075  33.983  1.00  0.00           O
ATOM   2955  N   GLU A 185       0.411  26.126  34.432  1.00  0.00           N
ATOM   2956  HN  GLU A 185      -0.416  26.682  34.511  1.00  0.00
ATOM   2957  CA  GLU A 185       0.347  24.691  34.699  1.00  0.00           C
ATOM   2958  HA  GLU A 185       0.997  24.499  35.434  1.00  0.00
ATOM   2959  CB  GLU A 185      -1.032  24.287  35.228  1.00  0.00           C
ATOM   2960  HB1 GLU A 185      -1.009  23.311  35.447  1.00  0.00
ATOM   2961  HB2 GLU A 185      -1.213  24.811  36.060  1.00  0.00
ATOM   2962  CG  GLU A 185      -2.188  24.521  34.272  1.00  0.00           C
ATOM   2963  HG1 GLU A 185      -1.808  24.664  33.358  1.00  0.00
ATOM   2964  HG2 GLU A 185      -2.758  23.700  34.271  1.00  0.00
ATOM   2965  CD  GLU A 185      -3.077  25.705  34.597  1.00  0.00           C
ATOM   2966  OE1 GLU A 185      -2.517  26.767  34.974  1.00  0.00           O
ATOM   2967  OE2 GLU A 185      -4.338  25.578  34.448  1.00  0.00           O
ATOM   2968  C   GLU A 185       0.771  23.923  33.430  1.00  0.00           C
ATOM   2969  O   GLU A 185       1.441  22.922  33.510  1.00  0.00           O
ATOM   2970  N   VAL A 186       0.434  24.461  32.253  1.00  0.00           N
ATOM   2971  HN  VAL A 186      -0.041  25.341  32.248  1.00  0.00
ATOM   2972  CA  VAL A 186       0.729  23.819  30.975  1.00  0.00           C
ATOM   2973  HA  VAL A 186       0.496  22.855  31.104  1.00  0.00
ATOM   2974  CB  VAL A 186      -0.141  24.380  29.837  1.00  0.00           C
ATOM   2975  HB  VAL A 186       0.010  25.368  29.797  1.00  0.00
ATOM   2976  CG1 VAL A 186       0.269  23.783  28.492  1.00  0.00           C
ATOM   2977 1HG1 VAL A 186      -0.309  24.162  27.769  1.00  0.00
```

```
ATOM   2978 2HG1 VAL A 186       1.226  24.007  28.306  1.00  0.00
ATOM   2979 3HG1 VAL A 186       0.159  22.789  28.521  1.00  0.00
ATOM   2980  CG2 VAL A 186      -1.630  24.175  30.099  1.00  0.00           C
ATOM   2981 1HG2 VAL A 186      -2.157  24.553  29.338  1.00  0.00
ATOM   2982 2HG2 VAL A 186      -1.822  23.197  30.185  1.00  0.00
ATOM   2983 3HG2 VAL A 186      -1.887  24.641  30.946  1.00  0.00
ATOM   2984  C   VAL A 186       2.221  23.966  30.655  1.00  0.00           C
ATOM   2985  O   VAL A 186       2.854  23.026  30.285  1.00  0.00           O
ATOM   2986  N   ILE A 187       2.756  25.170  30.784  1.00  0.00           N
ATOM   2987  HN  ILE A 187       2.160  25.942  31.005  1.00  0.00
ATOM   2988  CA  ILE A 187       4.202  25.413  30.613  1.00  0.00           C
ATOM   2989  HA  ILE A 187       4.397  25.256  29.645  1.00  0.00
ATOM   2990  CB  ILE A 187       4.577  26.863  30.980  1.00  0.00           C
ATOM   2991  HB  ILE A 187       4.336  27.009  31.939  1.00  0.00
ATOM   2992  CG2 ILE A 187       6.073  27.073  30.840  1.00  0.00           C
ATOM   2993 1HG2 ILE A 187       6.301  28.016  31.081  1.00  0.00
ATOM   2994 2HG2 ILE A 187       6.556  26.447  31.452  1.00  0.00
ATOM   2995 3HG2 ILE A 187       6.348  26.894  29.895  1.00  0.00
ATOM   2996  CG1 ILE A 187       3.785  27.903  30.177  1.00  0.00           C
ATOM   2997 1HG1 ILE A 187       2.812  27.679  30.237  1.00  0.00
ATOM   2998 2HG1 ILE A 187       3.945  28.805  30.578  1.00  0.00
ATOM   2999  CD  ILE A 187       4.155  27.973  28.715  1.00  0.00           C
ATOM   3000  HD1 ILE A 187       3.598  28.670  28.263  1.00  0.00
ATOM   3001  HD2 ILE A 187       5.122  28.211  28.627  1.00  0.00
ATOM   3002  HD3 ILE A 187       3.990  27.085  28.286  1.00  0.00
ATOM   3003  C   ILE A 187       4.988  24.435  31.489  1.00  0.00           C
ATOM   3004  O   ILE A 187       5.882  23.764  31.009  1.00  0.00           O
ATOM   3005  N   MET A 188       4.641  24.397  32.780  1.00  0.00           N
ATOM   3006  HN  MET A 188       3.803  24.870  33.054  1.00  0.00
ATOM   3007  CA  MET A 188       5.407  23.709  33.816  1.00  0.00           C
ATOM   3008  HA  MET A 188       6.354  23.974  33.635  1.00  0.00
ATOM   3009  CB  MET A 188       4.971  24.156  35.221  1.00  0.00           C
ATOM   3010  HB1 MET A 188       3.972  24.127  35.267  1.00  0.00
ATOM   3011  HB2 MET A 188       5.354  23.518  35.889  1.00  0.00
ATOM   3012  CG  MET A 188       5.423  25.564  35.597  1.00  0.00           C
ATOM   3013  HG1 MET A 188       6.401  25.556  35.389  1.00  0.00
ATOM   3014  HG2 MET A 188       4.947  26.144  34.936  1.00  0.00
ATOM   3015  SD  MET A 188       5.074  26.060  37.361  1.00  0.00           S
ATOM   3016  CE  MET A 188       3.283  26.014  37.425  1.00  0.00           C
ATOM   3017  HE1 MET A 188       2.976  26.267  38.342  1.00  0.00
ATOM   3018  HE2 MET A 188       2.909  26.659  36.758  1.00  0.00
ATOM   3019  HE3 MET A 188       2.965  25.090  37.210  1.00  0.00
ATOM   3020  C   MET A 188       5.228  22.187  33.711  1.00  0.00           C
ATOM   3021  O   MET A 188       6.038  21.455  34.224  1.00  0.00           O
ATOM   3022  N   SER A 189       4.146  21.720  33.078  1.00  0.00           N
ATOM   3023  HN  SER A 189       3.550  22.362  32.595  1.00  0.00
ATOM   3024  CA  SER A 189       3.811  20.294  33.073  1.00  0.00           C
ATOM   3025  HA  SER A 189       4.194  19.936  33.925  1.00  0.00
ATOM   3026  CB  SER A 189       2.328  20.088  33.089  1.00  0.00           C
ATOM   3027  HB1 SER A 189       2.111  19.143  33.334  1.00  0.00
ATOM   3028  HB2 SER A 189       1.898  20.710  33.744  1.00  0.00
ATOM   3029  OG  SER A 189       1.780  20.349  31.805  1.00  0.00           O
ATOM   3030  HG1 SER A 189       0.790  20.208  31.830  1.00  0.00
ATOM   3031  C   SER A 189       4.424  19.575  31.861  1.00  0.00           C
ATOM   3032  O   SER A 189       4.175  18.390  31.661  1.00  0.00           O
ATOM   3033  N   GLN A 190       5.179  20.288  31.030  1.00  0.00           N
ATOM   3034  HN  GLN A 190       5.303  21.267  31.194  1.00  0.00
ATOM   3035  CA  GLN A 190       5.828  19.664  29.884  1.00  0.00           C
ATOM   3036  HA  GLN A 190       5.113  19.287  29.296  1.00  0.00
ATOM   3037  CB  GLN A 190       6.594  20.717  29.080  1.00  0.00           C
ATOM   3038  HB1 GLN A 190       6.191  21.614  29.263  1.00  0.00
ATOM   3039  HB2 GLN A 190       7.549  20.714  29.377  1.00  0.00
ATOM   3040  CG  GLN A 190       6.557  20.470  27.577  1.00  0.00           C
ATOM   3041  HG1 GLN A 190       7.170  19.712  27.355  1.00  0.00
ATOM   3042  HG2 GLN A 190       5.624  20.234  27.307  1.00  0.00
ATOM   3043  CD  GLN A 190       6.990  21.682  26.797  1.00  0.00           C
ATOM   3044  OE1 GLN A 190       6.180  22.358  26.168  1.00  0.00           O
ATOM   3045  NE2 GLN A 190       8.281  21.973  26.853  1.00  0.00           N
ATOM   3046 1HE2 GLN A 190       8.900  21.400  27.390  1.00  0.00
ATOM   3047 2HE2 GLN A 190       8.637  22.766  26.358  1.00  0.00
ATOM   3048  C   GLN A 190       6.736  18.528  30.385  1.00  0.00           C
```

```
ATOM   3049  O   GLN A 190       7.449  18.712  31.351  1.00  0.00           O
ATOM   3050  N   HIS A 191       6.647  17.348  29.752  1.00  0.00           N
ATOM   3051  HN  HIS A 191       5.951  17.249  29.041  1.00  0.00
ATOM   3052  CA  HIS A 191       7.517  16.180  30.042  1.00  0.00           C
ATOM   3053  HA  HIS A 191       7.317  15.529  29.310  1.00  0.00
ATOM   3054  CB  HIS A 191       9.001  16.590  30.002  1.00  0.00           C
ATOM   3055  HB1 HIS A 191       9.119  17.434  30.525  1.00  0.00
ATOM   3056  HB2 HIS A 191       9.551  15.863  30.414  1.00  0.00
ATOM   3057  ND1 HIS A 191      10.889  16.792  28.350  1.00  0.00           N
ATOM   3058  CG  HIS A 191       9.533  16.833  28.630  1.00  0.00           C
ATOM   3059  CE1 HIS A 191      11.082  17.037  27.068  1.00  0.00           C
ATOM   3060  HE1 HIS A 191      11.967  17.082  26.605  1.00  0.00
ATOM   3061  NE2 HIS A 191       9.886  17.214  26.495  1.00  0.00           N
ATOM   3062  HE2 HIS A 191       9.728  17.405  25.526  1.00  0.00
ATOM   3063  CD2 HIS A 191       8.910  17.091  27.455  1.00  0.00           C
ATOM   3064  HD2 HIS A 191       7.924  17.176  27.313  1.00  0.00
ATOM   3065  C   HIS A 191       7.181  15.509  31.391  1.00  0.00           C
ATOM   3066  O   HIS A 191       7.926  14.637  31.837  1.00  0.00           O
ATOM   3067  N   TYR A 192       6.064  15.876  32.029  1.00  0.00           N
ATOM   3068  HN  TYR A 192       5.438  16.516  31.584  1.00  0.00
ATOM   3069  CA  TYR A 192       5.728  15.360  33.379  1.00  0.00           C
ATOM   3070  HA  TYR A 192       6.515  15.582  33.955  1.00  0.00
ATOM   3071  CB  TYR A 192       4.498  16.058  33.981  1.00  0.00           C
ATOM   3072  HB1 TYR A 192       4.546  15.984  34.977  1.00  0.00
ATOM   3073  HB2 TYR A 192       4.519  17.022  33.717  1.00  0.00
ATOM   3074  CG  TYR A 192       3.171  15.496  33.546  1.00  0.00           C
ATOM   3075  CD1 TYR A 192       2.621  15.856  32.335  1.00  0.00           C
ATOM   3076  HD1 TYR A 192       3.102  16.518  31.760  1.00  0.00
ATOM   3077  CE1 TYR A 192       1.422  15.327  31.893  1.00  0.00           C
ATOM   3078  HE1 TYR A 192       1.052  15.598  31.004  1.00  0.00
ATOM   3079  CZ  TYR A 192       0.738  14.425  32.681  1.00  0.00           C
ATOM   3080  OH  TYR A 192      -0.444  13.941  32.200  1.00  0.00           O
ATOM   3081  HH  TYR A 192      -0.836  13.302  32.861  1.00  0.00
ATOM   3082  CD2 TYR A 192       2.486  14.575  34.317  1.00  0.00           C
ATOM   3083  HD2 TYR A 192       2.870  14.289  35.195  1.00  0.00
ATOM   3084  CE2 TYR A 192       1.279  14.040  33.899  1.00  0.00           C
ATOM   3085  HE2 TYR A 192       0.799  13.378  34.474  1.00  0.00
ATOM   3086  C   TYR A 192       5.525  13.837  33.324  1.00  0.00           C
ATOM   3087  O   TYR A 192       5.868  13.123  34.277  1.00  0.00           O
ATOM   3088  N   ASP A 193       4.980  13.347  32.201  1.00  0.00           N
ATOM   3089  HN  ASP A 193       4.814  13.971  31.437  1.00  0.00
ATOM   3090  CA  ASP A 193       4.619  11.945  32.046  1.00  0.00           C
ATOM   3091  HA  ASP A 193       4.654  11.554  32.966  1.00  0.00
ATOM   3092  CB  ASP A 193       3.187  11.800  31.516  1.00  0.00           C
ATOM   3093  HB1 ASP A 193       2.988  10.825  31.421  1.00  0.00
ATOM   3094  HB2 ASP A 193       2.565  12.203  32.187  1.00  0.00
ATOM   3095  CG  ASP A 193       2.912  12.462  30.173  1.00  0.00           C
ATOM   3096  OD1 ASP A 193       3.809  13.191  29.658  1.00  0.00           O
ATOM   3097  OD2 ASP A 193       1.790  12.258  29.651  1.00  0.00           O
ATOM   3098  C   ASP A 193       5.642  11.226  31.155  1.00  0.00           C
ATOM   3099  O   ASP A 193       5.375  10.139  30.671  1.00  0.00           O
ATOM   3100  N   ALA A 194       6.835  11.805  31.006  1.00  0.00           N
ATOM   3101  HN  ALA A 194       7.036  12.625  31.541  1.00  0.00
ATOM   3102  CA  ALA A 194       7.869  11.293  30.091  1.00  0.00           C
ATOM   3103  HA  ALA A 194       7.446  11.254  29.186  1.00  0.00
ATOM   3104  CB  ALA A 194       9.041  12.243  30.041  1.00  0.00           C
ATOM   3105  HB1 ALA A 194       9.735  11.884  29.417  1.00  0.00
ATOM   3106  HB2 ALA A 194       8.732  13.137  29.717  1.00  0.00
ATOM   3107  HB3 ALA A 194       9.433  12.338  30.956  1.00  0.00
ATOM   3108  C   ALA A 194       8.327   9.886  30.502  1.00  0.00           C
ATOM   3109  O   ALA A 194       8.771   9.119  29.642  1.00  0.00           O
ATOM   3110  N   LYS A 195       8.203   9.544  31.792  1.00  0.00           N
ATOM   3111  HN  LYS A 195       7.761  10.182  32.423  1.00  0.00
ATOM   3112  CA  LYS A 195       8.695   8.266  32.303  1.00  0.00           C
ATOM   3113  HA  LYS A 195       9.205   7.857  31.546  1.00  0.00
ATOM   3114  CB  LYS A 195       9.613   8.505  33.499  1.00  0.00           C
ATOM   3115  HB1 LYS A 195       9.213   9.224  34.068  1.00  0.00
ATOM   3116  HB2 LYS A 195       9.674   7.658  34.027  1.00  0.00
ATOM   3117  CG  LYS A 195      11.019   8.929  33.119  1.00  0.00           C
ATOM   3118  HG1 LYS A 195      11.499   8.144  32.728  1.00  0.00
ATOM   3119  HG2 LYS A 195      10.963   9.659  32.438  1.00  0.00
```

```
ATOM   3120  CD  LYS A 195      11.807   9.434  34.302  1.00  0.00           C
ATOM   3121  HD1 LYS A 195      11.450  10.332  34.560  1.00  0.00
ATOM   3122  HD2 LYS A 195      11.682   8.795  35.061  1.00  0.00
ATOM   3123  CE  LYS A 195      13.280   9.570  34.038  1.00  0.00           C
ATOM   3124  HE1 LYS A 195      13.655   8.686  33.760  1.00  0.00
ATOM   3125  HE2 LYS A 195      13.431  10.240  33.311  1.00  0.00
ATOM   3126  NZ  LYS A 195      13.975  10.023  35.261  1.00  0.00           N
ATOM   3127  HZ1 LYS A 195      14.954  10.110  35.075  1.00  0.00
ATOM   3128  HZ2 LYS A 195      13.609  10.910  35.543  1.00  0.00
ATOM   3129  HZ3 LYS A 195      13.833   9.355  35.992  1.00  0.00
ATOM   3130  C   LYS A 195       7.542   7.332  32.696  1.00  0.00           C
ATOM   3131  O   LYS A 195       7.779   6.323  33.349  1.00  0.00           O
ATOM   3132  N   ALA A 196       6.322   7.649  32.273  1.00  0.00           N
ATOM   3133  HN  ALA A 196       6.211   8.395  31.616  1.00  0.00
ATOM   3134  CA  ALA A 196       5.136   6.931  32.747  1.00  0.00           C
ATOM   3135  HA  ALA A 196       5.107   7.072  33.737  1.00  0.00
ATOM   3136  CB  ALA A 196       3.887   7.502  32.121  1.00  0.00           C
ATOM   3137  HB1 ALA A 196       3.088   7.001  32.455  1.00  0.00
ATOM   3138  HB2 ALA A 196       3.802   8.467  32.367  1.00  0.00
ATOM   3139  HB3 ALA A 196       3.943   7.415  31.126  1.00  0.00
ATOM   3140  C   ALA A 196       5.240   5.440  32.432  1.00  0.00           C
ATOM   3141  O   ALA A 196       4.888   4.601  33.255  1.00  0.00           O
ATOM   3142  N   ASP A 197       5.718   5.124  31.222  1.00  0.00           N
ATOM   3143  HN  ASP A 197       6.077   5.850  30.636  1.00  0.00
ATOM   3144  CA  ASP A 197       5.729   3.766  30.741  1.00  0.00           C
ATOM   3145  HA  ASP A 197       4.793   3.432  30.852  1.00  0.00
ATOM   3146  CB  ASP A 197       6.080   3.708  29.256  1.00  0.00           C
ATOM   3147  HB1 ASP A 197       6.927   4.218  29.106  1.00  0.00
ATOM   3148  HB2 ASP A 197       6.215   2.752  28.995  1.00  0.00
ATOM   3149  CG  ASP A 197       5.011   4.295  28.366  1.00  0.00           C
ATOM   3150  OD1 ASP A 197       3.831   4.313  28.797  1.00  0.00           O
ATOM   3151  OD2 ASP A 197       5.371   4.757  27.251  1.00  0.00           O
ATOM   3152  C   ASP A 197       6.685   2.928  31.591  1.00  0.00           C
ATOM   3153  O   ASP A 197       6.464   1.749  31.769  1.00  0.00           O
ATOM   3154  N   LEU A 198       7.734   3.554  32.127  1.00  0.00           N
ATOM   3155  HN  LEU A 198       7.855   4.532  31.959  1.00  0.00
ATOM   3156  CA  LEU A 198       8.708   2.851  32.951  1.00  0.00           C
ATOM   3157  HA  LEU A 198       9.009   2.047  32.438  1.00  0.00
ATOM   3158  CB  LEU A 198       9.914   3.755  33.197  1.00  0.00           C
ATOM   3159  HB1 LEU A 198       9.575   4.653  33.478  1.00  0.00
ATOM   3160  HB2 LEU A 198      10.454   3.356  33.938  1.00  0.00
ATOM   3161  CG  LEU A 198      10.833   3.949  31.988  1.00  0.00           C
ATOM   3162  HG  LEU A 198      10.250   4.230  31.226  1.00  0.00
ATOM   3163  CD1 LEU A 198      11.868   5.042  32.234  1.00  0.00           C
ATOM   3164 1HD1 LEU A 198      12.448   5.140  31.425  1.00  0.00
ATOM   3165 2HD1 LEU A 198      11.402   5.908  32.416  1.00  0.00
ATOM   3166 3HD1 LEU A 198      12.433   4.795  33.022  1.00  0.00
ATOM   3167  CD2 LEU A 198      11.544   2.652  31.614  1.00  0.00           C
ATOM   3168 1HD2 LEU A 198      12.135   2.812  30.823  1.00  0.00
ATOM   3169 2HD2 LEU A 198      12.098   2.341  32.386  1.00  0.00
ATOM   3170 3HD2 LEU A 198      10.865   1.953  31.387  1.00  0.00
ATOM   3171  C   LEU A 198       8.046   2.391  34.261  1.00  0.00           C
ATOM   3172  O   LEU A 198       8.300   1.290  34.728  1.00  0.00           O
ATOM   3173  N   TRP A 199       7.161   3.210  34.830  1.00  0.00           N
ATOM   3174  HN  TRP A 199       7.024   4.127  34.455  1.00  0.00
ATOM   3175  CA  TRP A 199       6.397   2.788  35.982  1.00  0.00           C
ATOM   3176  HA  TRP A 199       7.079   2.544  36.672  1.00  0.00
ATOM   3177  CB  TRP A 199       5.529   3.922  36.528  1.00  0.00           C
ATOM   3178  HB1 TRP A 199       6.122   4.632  36.908  1.00  0.00
ATOM   3179  HB2 TRP A 199       4.986   4.305  35.781  1.00  0.00
ATOM   3180  CG  TRP A 199       4.589   3.478  37.604  1.00  0.00           C
ATOM   3181  CD1 TRP A 199       3.355   2.931  37.431  1.00  0.00           C
ATOM   3182  HD1 TRP A 199       2.920   2.771  36.545  1.00  0.00
ATOM   3183  NE1 TRP A 199       2.794   2.631  38.645  1.00  0.00           N
ATOM   3184  HE1 TRP A 199       1.884   2.239  38.782  1.00  0.00
ATOM   3185  CE2 TRP A 199       3.679   2.954  39.641  1.00  0.00           C
ATOM   3186  CD2 TRP A 199       4.834   3.481  39.026  1.00  0.00           C
ATOM   3187  CE3 TRP A 199       5.909   3.885  39.827  1.00  0.00           C
ATOM   3188  HE3 TRP A 199       6.739   4.262  39.416  1.00  0.00
ATOM   3189  CZ3 TRP A 199       5.799   3.752  41.192  1.00  0.00           C
ATOM   3190  HZ3 TRP A 199       6.563   4.027  41.776  1.00  0.00
```

```
ATOM   3191  CZ2 TRP A 199       3.543   2.863  41.023  1.00  0.00           C
ATOM   3192  HZ2 TRP A 199       2.696   2.542  41.446  1.00  0.00
ATOM   3193  CH2 TRP A 199       4.630   3.236  41.780  1.00  0.00           C
ATOM   3194  HH2 TRP A 199       4.588   3.138  42.774  1.00  0.00
ATOM   3195  C   TRP A 199       5.542   1.575  35.617  1.00  0.00           C
ATOM   3196  O   TRP A 199       5.508   0.588  36.336  1.00  0.00           O
ATOM   3197  N   SER A 200       4.832   1.676  34.499  1.00  0.00           N
ATOM   3198  HN  SER A 200       4.956   2.481  33.919  1.00  0.00
ATOM   3199  CA  SER A 200       3.882   0.656  34.090  1.00  0.00           C
ATOM   3200  HA  SER A 200       3.194   0.600  34.814  1.00  0.00
ATOM   3201  CB  SER A 200       3.180   1.055  32.833  1.00  0.00           C
ATOM   3202  HB1 SER A 200       3.839   1.210  32.097  1.00  0.00
ATOM   3203  HB2 SER A 200       2.530   0.348  32.554  1.00  0.00
ATOM   3204  OG  SER A 200       2.465   2.251  33.042  1.00  0.00           O
ATOM   3205  HG1 SER A 200       1.998   2.512  32.197  1.00  0.00
ATOM   3206  C   SER A 200       4.595  -0.685  33.927  1.00  0.00           C
ATOM   3207  O   SER A 200       4.131  -1.698  34.429  1.00  0.00           O
ATOM   3208  N   ILE A 201       5.721  -0.663  33.208  1.00  0.00           N
ATOM   3209  HN  ILE A 201       6.031   0.212  32.837  1.00  0.00
ATOM   3210  CA  ILE A 201       6.512  -1.842  32.940  1.00  0.00           C
ATOM   3211  HA  ILE A 201       5.936  -2.482  32.430  1.00  0.00
ATOM   3212  CB  ILE A 201       7.723  -1.480  32.050  1.00  0.00           C
ATOM   3213  HB  ILE A 201       8.173  -0.673  32.434  1.00  0.00
ATOM   3214  CG2 ILE A 201       8.773  -2.588  32.046  1.00  0.00           C
ATOM   3215 1HG2 ILE A 201       9.539  -2.319  31.462  1.00  0.00
ATOM   3216 2HG2 ILE A 201       9.102  -2.739  32.978  1.00  0.00
ATOM   3217 3HG2 ILE A 201       8.366  -3.432  31.695  1.00  0.00
ATOM   3218  CG1 ILE A 201       7.262  -1.126  30.633  1.00  0.00           C
ATOM   3219 1HG1 ILE A 201       7.010  -1.974  30.166  1.00  0.00
ATOM   3220 2HG1 ILE A 201       6.460  -0.534  30.705  1.00  0.00
ATOM   3221  CD  ILE A 201       8.305  -0.408  29.779  1.00  0.00           C
ATOM   3222  HD1 ILE A 201       7.922  -0.216  28.876  1.00  0.00
ATOM   3223  HD2 ILE A 201       8.563   0.451  30.221  1.00  0.00
ATOM   3224  HD3 ILE A 201       9.113  -0.989  29.682  1.00  0.00
ATOM   3225  C   ILE A 201       6.911  -2.473  34.287  1.00  0.00           C
ATOM   3226  O   ILE A 201       6.906  -3.695  34.446  1.00  0.00           O
ATOM   3227  N   GLY A 202       7.214  -1.611  35.262  1.00  0.00           N
ATOM   3228  HN  GLY A 202       7.234  -0.635  35.045  1.00  0.00
ATOM   3229  CA  GLY A 202       7.515  -2.023  36.618  1.00  0.00           C
ATOM   3230  HA1 GLY A 202       8.348  -2.576  36.590  1.00  0.00
ATOM   3231  HA2 GLY A 202       7.680  -1.197  37.157  1.00  0.00
ATOM   3232  C   GLY A 202       6.387  -2.826  37.242  1.00  0.00           C
ATOM   3233  O   GLY A 202       6.630  -3.892  37.809  1.00  0.00           O
ATOM   3234  N   THR A 203       5.161  -2.302  37.162  1.00  0.00           N
ATOM   3235  HN  THR A 203       5.041  -1.439  36.672  1.00  0.00
ATOM   3236  CA  THR A 203       3.968  -2.945  37.769  1.00  0.00           C
ATOM   3237  HA  THR A 203       4.180  -3.107  38.733  1.00  0.00
ATOM   3238  CB  THR A 203       2.746  -2.018  37.727  1.00  0.00           C
ATOM   3239  HB  THR A 203       1.960  -2.482  38.136  1.00  0.00
ATOM   3240  OG1 THR A 203       2.416  -1.743  36.369  1.00  0.00           O
ATOM   3241  HG1 THR A 203       1.620  -1.139  36.335  1.00  0.00
ATOM   3242  CG2 THR A 203       2.967  -0.709  38.463  1.00  0.00           C
ATOM   3243 1HG2 THR A 203       2.141  -0.149  38.402  1.00  0.00
ATOM   3244 2HG2 THR A 203       3.172  -0.897  39.424  1.00  0.00
ATOM   3245 3HG2 THR A 203       3.734  -0.219  38.048  1.00  0.00
ATOM   3246  C   THR A 203       3.701  -4.289  37.071  1.00  0.00           C
ATOM   3247  O   THR A 203       3.305  -5.258  37.694  1.00  0.00           O
ATOM   3248  N   VAL A 204       3.941  -4.338  35.764  1.00  0.00           N
ATOM   3249  HN  VAL A 204       4.325  -3.534  35.311  1.00  0.00
ATOM   3250  CA  VAL A 204       3.663  -5.522  34.978  1.00  0.00           C
ATOM   3251  HA  VAL A 204       2.741  -5.815  35.231  1.00  0.00
ATOM   3252  CB  VAL A 204       3.664  -5.207  33.472  1.00  0.00           C
ATOM   3253  HB  VAL A 204       4.469  -4.649  33.272  1.00  0.00
ATOM   3254  CG1 VAL A 204       3.778  -6.460  32.616  1.00  0.00           C
ATOM   3255 1HG1 VAL A 204       3.774  -6.205  31.649  1.00  0.00
ATOM   3256 2HG1 VAL A 204       4.631  -6.935  32.832  1.00  0.00
ATOM   3257 3HG1 VAL A 204       3.003  -7.064  32.804  1.00  0.00
ATOM   3258  CG2 VAL A 204       2.422  -4.421  33.103  1.00  0.00           C
ATOM   3259 1HG2 VAL A 204       2.433  -4.222  32.123  1.00  0.00
ATOM   3260 2HG2 VAL A 204       1.609  -4.959  33.325  1.00  0.00
ATOM   3261 3HG2 VAL A 204       2.405  -3.564  33.617  1.00  0.00
```

```
ATOM   3262  C   VAL A 204       4.672  -6.612  35.345  1.00  0.00           C
ATOM   3263  O   VAL A 204       4.281  -7.745  35.577  1.00  0.00           O
ATOM   3264  N   ILE A 205       5.960  -6.249  35.391  1.00  0.00           N
ATOM   3265  HN  ILE A 205       6.210  -5.318  35.125  1.00  0.00
ATOM   3266  CA  ILE A 205       7.006  -7.163  35.814  1.00  0.00           C
ATOM   3267  HA  ILE A 205       6.991  -7.917  35.157  1.00  0.00
ATOM   3268  CB  ILE A 205       8.392  -6.507  35.775  1.00  0.00           C
ATOM   3269  HB  ILE A 205       8.341  -5.662  36.308  1.00  0.00
ATOM   3270  CG2 ILE A 205       9.433  -7.409  36.411  1.00  0.00           C
ATOM   3271 1HG2 ILE A 205      10.327  -6.963  36.375  1.00  0.00
ATOM   3272 2HG2 ILE A 205       9.186  -7.583  37.364  1.00  0.00
ATOM   3273 3HG2 ILE A 205       9.472  -8.275  35.913  1.00  0.00
ATOM   3274  CG1 ILE A 205       8.808  -6.118  34.363  1.00  0.00           C
ATOM   3275 1HG1 ILE A 205       9.146  -6.930  33.888  1.00  0.00
ATOM   3276 2HG1 ILE A 205       8.013  -5.755  33.878  1.00  0.00
ATOM   3277  CD  ILE A 205       9.891  -5.074  34.351  1.00  0.00           C
ATOM   3278  HD1 ILE A 205      10.130  -4.852  33.406  1.00  0.00
ATOM   3279  HD2 ILE A 205       9.565  -4.250  34.815  1.00  0.00
ATOM   3280  HD3 ILE A 205      10.698  -5.426  34.825  1.00  0.00
ATOM   3281  C   ILE A 205       6.688  -7.636  37.232  1.00  0.00           C
ATOM   3282  O   ILE A 205       6.820  -8.805  37.529  1.00  0.00           O
ATOM   3283  N   TYR A 206       6.311  -6.701  38.108  1.00  0.00           N
ATOM   3284  HN  TYR A 206       6.231  -5.751  37.806  1.00  0.00
ATOM   3285  CA  TYR A 206       6.016  -7.035  39.486  1.00  0.00           C
ATOM   3286  HA  TYR A 206       6.886  -7.366  39.851  1.00  0.00
ATOM   3287  CB  TYR A 206       5.570  -5.818  40.299  1.00  0.00           C
ATOM   3288  HB1 TYR A 206       6.265  -5.104  40.215  1.00  0.00
ATOM   3289  HB2 TYR A 206       4.703  -5.485  39.928  1.00  0.00
ATOM   3290  CG  TYR A 206       5.371  -6.105  41.766  1.00  0.00           C
ATOM   3291  CD1 TYR A 206       4.177  -6.626  42.256  1.00  0.00           C
ATOM   3292  HD1 TYR A 206       3.425  -6.802  41.621  1.00  0.00
ATOM   3293  CE1 TYR A 206       4.006  -6.906  43.602  1.00  0.00           C
ATOM   3294  HE1 TYR A 206       3.136  -7.271  43.934  1.00  0.00
ATOM   3295  CZ  TYR A 206       5.053  -6.675  44.488  1.00  0.00           C
ATOM   3296  OH  TYR A 206       4.968  -6.945  45.828  1.00  0.00           O
ATOM   3297  HH  TYR A 206       5.831  -6.704  46.272  1.00  0.00
ATOM   3298  CD2 TYR A 206       6.407  -5.915  42.658  1.00  0.00           C
ATOM   3299  HD2 TYR A 206       7.284  -5.570  42.323  1.00  0.00
ATOM   3300  CE2 TYR A 206       6.258  -6.192  44.006  1.00  0.00           C
ATOM   3301  HE2 TYR A 206       7.022  -6.043  44.633  1.00  0.00
ATOM   3302  C   TYR A 206       4.928  -8.115  39.532  1.00  0.00           C
ATOM   3303  O   TYR A 206       5.075  -9.099  40.238  1.00  0.00           O
ATOM   3304  N   GLN A 207       3.857  -7.927  38.754  1.00  0.00           N
ATOM   3305  HN  GLN A 207       3.811  -7.124  38.160  1.00  0.00
ATOM   3306  CA  GLN A 207       2.768  -8.871  38.764  1.00  0.00           C
ATOM   3307  HA  GLN A 207       2.498  -8.907  39.726  1.00  0.00
ATOM   3308  CB  GLN A 207       1.584  -8.400  37.933  1.00  0.00           C
ATOM   3309  HB1 GLN A 207       1.202  -7.572  38.344  1.00  0.00
ATOM   3310  HB2 GLN A 207       1.894  -8.199  37.004  1.00  0.00
ATOM   3311  CG  GLN A 207       0.508  -9.474  37.880  1.00  0.00           C
ATOM   3312  HG1 GLN A 207       0.713 -10.096  37.125  1.00  0.00
ATOM   3313  HG2 GLN A 207       0.519  -9.981  38.742  1.00  0.00
ATOM   3314  CD  GLN A 207      -0.875  -8.944  37.676  1.00  0.00           C
ATOM   3315  OE1 GLN A 207      -1.073  -7.771  37.410  1.00  0.00           O
ATOM   3316  NE2 GLN A 207      -1.847  -9.805  37.904  1.00  0.00           N
ATOM   3317 1HE2 GLN A 207      -1.631 -10.738  38.192  1.00  0.00
ATOM   3318 2HE2 GLN A 207      -2.800  -9.525  37.789  1.00  0.00
ATOM   3319  C   GLN A 207       3.245 -10.235  38.246  1.00  0.00           C
ATOM   3320  O   GLN A 207       2.849 -11.259  38.775  1.00  0.00           O
ATOM   3321  N   CYS A 208       4.076 -10.221  37.196  1.00  0.00           N
ATOM   3322  HN  CYS A 208       4.337  -9.338  36.806  1.00  0.00
ATOM   3323  CA  CYS A 208       4.618 -11.435  36.595  1.00  0.00           C
ATOM   3324  HA  CYS A 208       3.819 -11.913  36.231  1.00  0.00
ATOM   3325  CB  CYS A 208       5.603 -11.122  35.472  1.00  0.00           C
ATOM   3326  HB1 CYS A 208       6.234 -10.460  35.878  1.00  0.00
ATOM   3327  HB2 CYS A 208       6.081 -11.987  35.318  1.00  0.00
ATOM   3328  SG  CYS A 208       4.844 -10.489  33.952  1.00  0.00           S
ATOM   3329  HG1 CYS A 208       5.556 -10.311  33.273  1.00  0.00
ATOM   3330  C   CYS A 208       5.353 -12.258  37.654  1.00  0.00           C
ATOM   3331  O   CYS A 208       5.280 -13.488  37.660  1.00  0.00           O
ATOM   3332  N   LEU A 209       6.089 -11.557  38.521  1.00  0.00           N
```

```
ATOM   3333  HN  LEU A 209       5.990 -10.562  38.528  1.00  0.00
ATOM   3334  CA  LEU A 209       7.027 -12.161  39.455  1.00  0.00           C
ATOM   3335  HA  LEU A 209       7.460 -12.934  38.991  1.00  0.00
ATOM   3336  CB  LEU A 209       8.064 -11.103  39.848  1.00  0.00           C
ATOM   3337  HB1 LEU A 209       8.509 -10.784  39.011  1.00  0.00
ATOM   3338  HB2 LEU A 209       7.578 -10.342  40.278  1.00  0.00
ATOM   3339  CG  LEU A 209       9.158 -11.560  40.812  1.00  0.00           C
ATOM   3340  HG  LEU A 209       8.718 -11.928  41.631  1.00  0.00
ATOM   3341  CD1 LEU A 209      10.016 -12.650  40.180  1.00  0.00           C
ATOM   3342 1HD1 LEU A 209      10.724 -12.934  40.827  1.00  0.00
ATOM   3343 2HD1 LEU A 209       9.441 -13.435  39.950  1.00  0.00
ATOM   3344 3HD1 LEU A 209      10.446 -12.296  39.349  1.00  0.00
ATOM   3345  CD2 LEU A 209      10.036 -10.391  41.214  1.00  0.00           C
ATOM   3346 1HD2 LEU A 209      10.745 -10.708  41.844  1.00  0.00
ATOM   3347 2HD2 LEU A 209      10.464 -10.000  40.399  1.00  0.00
ATOM   3348 3HD2 LEU A 209       9.478  -9.694  41.664  1.00  0.00
ATOM   3349  C   LEU A 209       6.280 -12.665  40.691  1.00  0.00           C
ATOM   3350  O   LEU A 209       6.476 -13.798  41.107  1.00  0.00           O
ATOM   3351  N   VAL A 210       5.414 -11.805  41.232  1.00  0.00           N
ATOM   3352  HN  VAL A 210       5.165 -10.996  40.700  1.00  0.00
ATOM   3353  CA  VAL A 210       4.810 -11.968  42.542  1.00  0.00           C
ATOM   3354  HA  VAL A 210       5.412 -12.596  43.034  1.00  0.00
ATOM   3355  CB  VAL A 210       4.746 -10.620  43.279  1.00  0.00           C
ATOM   3356  HB  VAL A 210       4.285  -9.963  42.682  1.00  0.00
ATOM   3357  CG1 VAL A 210       3.929 -10.706  44.552  1.00  0.00           C
ATOM   3358 1HG1 VAL A 210       3.913  -9.811  44.999  1.00  0.00
ATOM   3359 2HG1 VAL A 210       2.995 -10.985  44.330  1.00  0.00
ATOM   3360 3HG1 VAL A 210       4.340 -11.379  45.167  1.00  0.00
ATOM   3361  CG2 VAL A 210       6.127 -10.081  43.557  1.00  0.00           C
ATOM   3362 1HG2 VAL A 210       6.053  -9.206  44.036  1.00  0.00
ATOM   3363 2HG2 VAL A 210       6.630 -10.731  44.126  1.00  0.00
ATOM   3364 3HG2 VAL A 210       6.613  -9.948  42.693  1.00  0.00
ATOM   3365  C   VAL A 210       3.406 -12.578  42.409  1.00  0.00           C
ATOM   3366  O   VAL A 210       3.028 -13.391  43.216  1.00  0.00           O
ATOM   3367  N   GLY A 211       2.629 -12.123  41.423  1.00  0.00           N
ATOM   3368  HN  GLY A 211       2.946 -11.351  40.873  1.00  0.00
ATOM   3369  CA  GLY A 211       1.325 -12.722  41.123  1.00  0.00           C
ATOM   3370  HA1 GLY A 211       1.299 -12.927  40.145  1.00  0.00
ATOM   3371  HA2 GLY A 211       1.251 -13.573  41.644  1.00  0.00
ATOM   3372  C   GLY A 211       0.165 -11.809  41.479  1.00  0.00           C
ATOM   3373  O   GLY A 211      -0.993 -12.204  41.393  1.00  0.00           O
ATOM   3374  N   LYS A 212       0.480 -10.584  41.889  1.00  0.00           N
ATOM   3375  HN  LYS A 212       1.425 -10.388  42.150  1.00  0.00
ATOM   3376  CA  LYS A 212      -0.500  -9.531  41.969  1.00  0.00           C
ATOM   3377  HA  LYS A 212      -1.187  -9.717  41.267  1.00  0.00
ATOM   3378  CB  LYS A 212      -1.193  -9.524  43.335  1.00  0.00           C
ATOM   3379  HB1 LYS A 212      -2.006  -8.946  43.263  1.00  0.00
ATOM   3380  HB2 LYS A 212      -1.468 -10.462  43.544  1.00  0.00
ATOM   3381  CG  LYS A 212      -0.373  -9.016  44.510  1.00  0.00           C
ATOM   3382  HG1 LYS A 212       0.555  -9.381  44.434  1.00  0.00
ATOM   3383  HG2 LYS A 212      -0.342  -8.017  44.471  1.00  0.00
ATOM   3384  CD  LYS A 212      -0.983  -9.453  45.857  1.00  0.00           C
ATOM   3385  HD1 LYS A 212      -1.976  -9.504  45.753  1.00  0.00
ATOM   3386  HD2 LYS A 212      -0.625 -10.357  46.088  1.00  0.00
ATOM   3387  CE  LYS A 212      -0.687  -8.526  47.022  1.00  0.00           C
ATOM   3388  HE1 LYS A 212      -0.976  -7.597  46.791  1.00  0.00
ATOM   3389  HE2 LYS A 212      -1.185  -8.838  47.831  1.00  0.00
ATOM   3390  NZ  LYS A 212       0.759  -8.502  47.341  1.00  0.00           N
ATOM   3391  HZ1 LYS A 212       0.920  -7.884  48.110  1.00  0.00
ATOM   3392  HZ2 LYS A 212       1.062  -9.423  47.584  1.00  0.00
ATOM   3393  HZ3 LYS A 212       1.271  -8.182  46.544  1.00  0.00
ATOM   3394  C   LYS A 212       0.204  -8.216  41.696  1.00  0.00           C
ATOM   3395  O   LYS A 212       1.437  -8.145  41.760  1.00  0.00           O
ATOM   3396  N   PRO A 213      -0.539  -7.143  41.343  1.00  0.00           N
ATOM   3397  CD  PRO A 213      -1.986  -7.129  41.101  1.00  0.00           C
ATOM   3398  HD1 PRO A 213      -2.485  -7.031  41.962  1.00  0.00
ATOM   3399  HD2 PRO A 213      -2.274  -7.970  40.643  1.00  0.00
ATOM   3400  CA  PRO A 213       0.071  -5.829  41.159  1.00  0.00           C
ATOM   3401  HA  PRO A 213       0.852  -5.896  40.539  1.00  0.00
ATOM   3402  CB  PRO A 213      -1.067  -4.941  40.618  1.00  0.00           C
ATOM   3403  HB1 PRO A 213      -1.399  -4.318  41.326  1.00  0.00
```

```
ATOM   3404  HB2 PRO A 213      -0.759  -4.412  39.827  1.00  0.00
ATOM   3405  CG  PRO A 213      -2.166  -5.919  40.211  1.00  0.00           C
ATOM   3406  HG1 PRO A 213      -3.069  -5.514  40.356  1.00  0.00
ATOM   3407  HG2 PRO A 213      -2.068  -6.178  39.250  1.00  0.00
ATOM   3408  C   PRO A 213       0.601  -5.299  42.483  1.00  0.00           C
ATOM   3409  O   PRO A 213       0.140  -5.692  43.546  1.00  0.00           O
ATOM   3410  N   PRO A 214       1.590  -4.390  42.457  1.00  0.00           N
ATOM   3411  CD  PRO A 214       2.204  -3.829  41.237  1.00  0.00           C
ATOM   3412  HD1 PRO A 214       1.498  -3.529  40.596  1.00  0.00
ATOM   3413  HD2 PRO A 214       2.789  -4.509  40.795  1.00  0.00
ATOM   3414  CA  PRO A 214       2.154  -3.852  43.694  1.00  0.00           C
ATOM   3415  HA  PRO A 214       2.383  -4.617  44.296  1.00  0.00
ATOM   3416  CB  PRO A 214       3.355  -3.020  43.201  1.00  0.00           C
ATOM   3417  HB1 PRO A 214       3.466  -2.193  43.752  1.00  0.00
ATOM   3418  HB2 PRO A 214       4.198  -3.557  43.232  1.00  0.00
ATOM   3419  CG  PRO A 214       3.013  -2.656  41.766  1.00  0.00           C
ATOM   3420  HG1 PRO A 214       2.472  -1.816  41.737  1.00  0.00
ATOM   3421  HG2 PRO A 214       3.847  -2.534  41.227  1.00  0.00
ATOM   3422  C   PRO A 214       1.165  -2.971  44.499  1.00  0.00           C
ATOM   3423  O   PRO A 214       1.291  -2.857  45.719  1.00  0.00           O
ATOM   3424  N   PHE A 215       0.202  -2.348  43.814  1.00  0.00           N
ATOM   3425  HN  PHE A 215       0.081  -2.571  42.847  1.00  0.00
ATOM   3426  CA  PHE A 215      -0.676  -1.361  44.415  1.00  0.00           C
ATOM   3427  HA  PHE A 215      -0.680  -1.537  45.399  1.00  0.00
ATOM   3428  CB  PHE A 215      -0.130   0.049  44.161  1.00  0.00           C
ATOM   3429  HB1 PHE A 215      -0.176   0.236  43.180  1.00  0.00
ATOM   3430  HB2 PHE A 215      -0.703   0.705  44.652  1.00  0.00
ATOM   3431  CG  PHE A 215       1.297   0.260  44.604  1.00  0.00           C
ATOM   3432  CD1 PHE A 215       1.638   0.258  45.950  1.00  0.00           C
ATOM   3433  HD1 PHE A 215       0.926   0.115  46.638  1.00  0.00
ATOM   3434  CE1 PHE A 215       2.951   0.452  46.356  1.00  0.00           C
ATOM   3435  HE1 PHE A 215       3.182   0.425  47.328  1.00  0.00
ATOM   3436  CZ  PHE A 215       3.934   0.681  45.422  1.00  0.00           C
ATOM   3437  HZ  PHE A 215       4.876   0.847  45.714  1.00  0.00
ATOM   3438  CD2 PHE A 215       2.299   0.486  43.673  1.00  0.00           C
ATOM   3439  HD2 PHE A 215       2.073   0.509  42.699  1.00  0.00
ATOM   3440  CE2 PHE A 215       3.613   0.681  44.078  1.00  0.00           C
ATOM   3441  HE2 PHE A 215       4.329   0.822  43.395  1.00  0.00
ATOM   3442  C   PHE A 215      -2.095  -1.532  43.864  1.00  0.00           C
ATOM   3443  O   PHE A 215      -2.345  -1.268  42.695  1.00  0.00           O
ATOM   3444  N   GLN A 216      -3.006  -1.987  44.727  1.00  0.00           N
ATOM   3445  HN  GLN A 216      -2.705  -2.198  45.657  1.00  0.00
ATOM   3446  CA  GLN A 216      -4.420  -2.197  44.400  1.00  0.00           C
ATOM   3447  HA  GLN A 216      -4.512  -2.106  43.408  1.00  0.00
ATOM   3448  CB  GLN A 216      -4.862  -3.600  44.825  1.00  0.00           C
ATOM   3449  HB1 GLN A 216      -4.960  -3.615  45.820  1.00  0.00
ATOM   3450  HB2 GLN A 216      -5.745  -3.798  44.400  1.00  0.00
ATOM   3451  CG  GLN A 216      -3.886  -4.695  44.426  1.00  0.00           C
ATOM   3452  HG1 GLN A 216      -3.652  -4.582  43.460  1.00  0.00
ATOM   3453  HG2 GLN A 216      -3.059  -4.608  44.982  1.00  0.00
ATOM   3454  CD  GLN A 216      -4.452  -6.081  44.622  1.00  0.00           C
ATOM   3455  OE1 GLN A 216      -5.497  -6.444  44.074  1.00  0.00           O
ATOM   3456  NE2 GLN A 216      -3.731  -6.886  45.386  1.00  0.00           N
ATOM   3457 1HE2 GLN A 216      -2.875  -6.560  45.788  1.00  0.00
ATOM   3458 2HE2 GLN A 216      -4.042  -7.820  45.562  1.00  0.00
ATOM   3459  C   GLN A 216      -5.255  -1.139  45.125  1.00  0.00           C
ATOM   3460  O   GLN A 216      -4.816  -0.571  46.134  1.00  0.00           O
ATOM   3461  N   ALA A 217      -6.469  -0.898  44.626  1.00  0.00           N
ATOM   3462  HN  ALA A 217      -6.742  -1.339  43.771  1.00  0.00
ATOM   3463  CA  ALA A 217      -7.398  -0.014  45.294  1.00  0.00           C
ATOM   3464  HA  ALA A 217      -7.258  -0.140  46.276  1.00  0.00
ATOM   3465  CB  ALA A 217      -7.091   1.414  44.942  1.00  0.00           C
ATOM   3466  HB1 ALA A 217      -7.737   2.018  45.408  1.00  0.00
ATOM   3467  HB2 ALA A 217      -6.160   1.636  45.231  1.00  0.00
ATOM   3468  HB3 ALA A 217      -7.172   1.538  43.953  1.00  0.00
ATOM   3469  C   ALA A 217      -8.832  -0.380  44.921  1.00  0.00           C
ATOM   3470  O   ALA A 217      -9.063  -1.120  43.964  1.00  0.00           O
ATOM   3471  N   ASN A 218      -9.778   0.186  45.686  1.00  0.00           N
ATOM   3472  HN  ASN A 218      -9.502   0.897  46.333  1.00  0.00
ATOM   3473  CA  ASN A 218     -11.197  -0.184  45.623  1.00  0.00           C
ATOM   3474  HA  ASN A 218     -11.203  -1.176  45.495  1.00  0.00
```

```
ATOM   3475  CB  ASN A 218     -11.933   0.144  46.928  1.00  0.00           C
ATOM   3476  HB1 ASN A 218     -11.959   1.138  47.039  1.00  0.00
ATOM   3477  HB2 ASN A 218     -12.867  -0.209  46.865  1.00  0.00
ATOM   3478  CG  ASN A 218     -11.286  -0.452  48.164  1.00  0.00           C
ATOM   3479  OD1 ASN A 218     -10.584  -1.461  48.089  1.00  0.00           O
ATOM   3480  ND2 ASN A 218     -11.519   0.168  49.310  1.00  0.00           N
ATOM   3481 1HD2 ASN A 218     -12.096   0.985  49.328  1.00  0.00
ATOM   3482 2HD2 ASN A 218     -11.118  -0.180  50.157  1.00  0.00
ATOM   3483  C   ASN A 218     -11.871   0.527  44.437  1.00  0.00           C
ATOM   3484  O   ASN A 218     -12.927   0.113  43.990  1.00  0.00           O
ATOM   3485  N   SER A 219     -11.249   1.603  43.940  1.00  0.00           N
ATOM   3486  HN  SER A 219     -10.364   1.863  44.326  1.00  0.00
ATOM   3487  CA  SER A 219     -11.801   2.412  42.865  1.00  0.00           C
ATOM   3488  HA  SER A 219     -12.172   1.775  42.189  1.00  0.00
ATOM   3489  CB  SER A 219     -12.933   3.265  43.375  1.00  0.00           C
ATOM   3490  HB1 SER A 219     -13.471   3.621  42.611  1.00  0.00
ATOM   3491  HB2 SER A 219     -13.525   2.731  43.979  1.00  0.00
ATOM   3492  OG  SER A 219     -12.444   4.374  44.116  1.00  0.00           O
ATOM   3493  HG1 SER A 219     -13.214   4.922  44.442  1.00  0.00
ATOM   3494  C   SER A 219     -10.700   3.264  42.230  1.00  0.00           C
ATOM   3495  O   SER A 219      -9.584   3.341  42.754  1.00  0.00           O
ATOM   3496  N   PRO A 220     -10.974   3.907  41.065  1.00  0.00           N
ATOM   3497  CD  PRO A 220     -12.132   3.638  40.191  1.00  0.00           C
ATOM   3498  HD1 PRO A 220     -12.949   4.098  40.538  1.00  0.00
ATOM   3499  HD2 PRO A 220     -12.304   2.655  40.128  1.00  0.00
ATOM   3500  CA  PRO A 220     -10.050   4.886  40.482  1.00  0.00           C
ATOM   3501  HA  PRO A 220      -9.176   4.427  40.323  1.00  0.00
ATOM   3502  CB  PRO A 220     -10.791   5.370  39.226  1.00  0.00           C
ATOM   3503  HB1 PRO A 220     -11.330   6.189  39.421  1.00  0.00
ATOM   3504  HB2 PRO A 220     -10.150   5.562  38.483  1.00  0.00
ATOM   3505  CG  PRO A 220     -11.698   4.210  38.860  1.00  0.00           C
ATOM   3506  HG1 PRO A 220     -12.489   4.529  38.338  1.00  0.00
ATOM   3507  HG2 PRO A 220     -11.201   3.525  38.327  1.00  0.00
ATOM   3508  C   PRO A 220      -9.715   6.080  41.397  1.00  0.00           C
ATOM   3509  O   PRO A 220      -8.565   6.508  41.437  1.00  0.00           O
ATOM   3510  N   GLN A 221     -10.716   6.608  42.118  1.00  0.00           N
ATOM   3511  HN  GLN A 221     -11.613   6.166  42.095  1.00  0.00
ATOM   3512  CA  GLN A 221     -10.539   7.815  42.941  1.00  0.00           C
ATOM   3513  HA  GLN A 221      -9.973   8.438  42.402  1.00  0.00
ATOM   3514  CB  GLN A 221     -11.882   8.502  43.201  1.00  0.00           C
ATOM   3515  HB1 GLN A 221     -12.594   7.802  43.255  1.00  0.00
ATOM   3516  HB2 GLN A 221     -11.828   8.990  44.072  1.00  0.00
ATOM   3517  CG  GLN A 221     -12.262   9.500  42.105  1.00  0.00           C
ATOM   3518  HG1 GLN A 221     -13.085   9.991  42.390  1.00  0.00
ATOM   3519  HG2 GLN A 221     -11.511  10.149  41.988  1.00  0.00
ATOM   3520  CD  GLN A 221     -12.538   8.847  40.767  1.00  0.00           C
ATOM   3521  OE1 GLN A 221     -13.025   7.720  40.696  1.00  0.00           O
ATOM   3522  NE2 GLN A 221     -12.236   9.558  39.688  1.00  0.00           N
ATOM   3523 1HE2 GLN A 221     -11.848  10.474  39.786  1.00  0.00
ATOM   3524 2HE2 GLN A 221     -12.397   9.177  38.777  1.00  0.00
ATOM   3525  C   GLN A 221      -9.799   7.476  44.247  1.00  0.00           C
ATOM   3526  O   GLN A 221      -9.163   8.359  44.842  1.00  0.00           O
ATOM   3527  N   ASP A 222      -9.871   6.206  44.673  1.00  0.00           N
ATOM   3528  HN  ASP A 222     -10.433   5.564  44.152  1.00  0.00
ATOM   3529  CA  ASP A 222      -9.161   5.701  45.878  1.00  0.00           C
ATOM   3530  HA  ASP A 222      -9.351   6.360  46.605  1.00  0.00
ATOM   3531  CB  ASP A 222      -9.693   4.328  46.321  1.00  0.00           C
ATOM   3532  HB1 ASP A 222     -10.088   3.875  45.522  1.00  0.00
ATOM   3533  HB2 ASP A 222      -8.922   3.791  46.662  1.00  0.00
ATOM   3534  CG  ASP A 222     -10.758   4.358  47.413  1.00  0.00           C
ATOM   3535  OD1 ASP A 222     -11.281   5.449  47.703  1.00  0.00           O
ATOM   3536  OD2 ASP A 222     -11.048   3.285  47.973  1.00  0.00           O
ATOM   3537  C   ASP A 222      -7.649   5.634  45.609  1.00  0.00           C
ATOM   3538  O   ASP A 222      -6.839   5.972  46.489  1.00  0.00           O
ATOM   3539  N   LEU A 223      -7.282   5.189  44.397  1.00  0.00           N
ATOM   3540  HN  LEU A 223      -7.996   4.930  43.746  1.00  0.00
ATOM   3541  CA  LEU A 223      -5.872   5.065  43.983  1.00  0.00           C
ATOM   3542  HA  LEU A 223      -5.372   4.644  44.740  1.00  0.00
ATOM   3543  CB  LEU A 223      -5.792   4.162  42.746  1.00  0.00           C
ATOM   3544  HB1 LEU A 223      -6.236   3.294  42.969  1.00  0.00
ATOM   3545  HB2 LEU A 223      -6.291   4.610  42.004  1.00  0.00
```

| | | | | | | | | | | | |
|---|---|---|---|---|---|---|---|---|---|---|---|
| ATOM | 3546 | CG | LEU | A | 223 | -4.378 | 3.853 | 42.243 | 1.00 | 0.00 | C |
| ATOM | 3547 | HG | LEU | A | 223 | -3.998 | 4.710 | 41.897 | 1.00 | 0.00 | |
| ATOM | 3548 | CD1 | LEU | A | 223 | -3.460 | 3.354 | 43.358 | 1.00 | 0.00 | C |
| ATOM | 3549 | 1HD1 | LEU | A | 223 | -2.552 | 3.166 | 42.983 | 1.00 | 0.00 | |
| ATOM | 3550 | 2HD1 | LEU | A | 223 | -3.389 | 4.053 | 44.069 | 1.00 | 0.00 | |
| ATOM | 3551 | 3HD1 | LEU | A | 223 | -3.838 | 2.516 | 43.751 | 1.00 | 0.00 | |
| ATOM | 3552 | CD2 | LEU | A | 223 | -4.418 | 2.835 | 41.113 | 1.00 | 0.00 | C |
| ATOM | 3553 | 1HD2 | LEU | A | 223 | -3.487 | 2.648 | 40.800 | 1.00 | 0.00 | |
| ATOM | 3554 | 2HD2 | LEU | A | 223 | -4.836 | 1.988 | 41.442 | 1.00 | 0.00 | |
| ATOM | 3555 | 3HD2 | LEU | A | 223 | -4.958 | 3.200 | 40.354 | 1.00 | 0.00 | |
| ATOM | 3556 | C | LEU | A | 223 | -5.271 | 6.458 | 43.701 | 1.00 | 0.00 | C |
| ATOM | 3557 | O | LEU | A | 223 | -4.068 | 6.668 | 43.885 | 1.00 | 0.00 | O |
| ATOM | 3558 | N | ARG | A | 224 | -6.116 | 7.391 | 43.239 | 1.00 | 0.00 | N |
| ATOM | 3559 | HN | ARG | A | 224 | -7.063 | 7.124 | 43.062 | 1.00 | 0.00 | |
| ATOM | 3560 | CA | ARG | A | 224 | -5.726 | 8.785 | 42.979 | 1.00 | 0.00 | C |
| ATOM | 3561 | HA | ARG | A | 224 | -4.988 | 8.753 | 42.305 | 1.00 | 0.00 | |
| ATOM | 3562 | CB | ARG | A | 224 | -6.906 | 9.575 | 42.398 | 1.00 | 0.00 | C |
| ATOM | 3563 | HB1 | ARG | A | 224 | -7.253 | 9.081 | 41.601 | 1.00 | 0.00 | |
| ATOM | 3564 | HB2 | ARG | A | 224 | -7.621 | 9.629 | 43.095 | 1.00 | 0.00 | |
| ATOM | 3565 | CG | ARG | A | 224 | -6.568 | 10.994 | 41.960 | 1.00 | 0.00 | C |
| ATOM | 3566 | HG1 | ARG | A | 224 | -5.757 | 11.301 | 42.458 | 1.00 | 0.00 | |
| ATOM | 3567 | HG2 | ARG | A | 224 | -6.376 | 10.992 | 40.979 | 1.00 | 0.00 | |
| ATOM | 3568 | CD | ARG | A | 224 | -7.707 | 11.965 | 42.234 | 1.00 | 0.00 | C |
| ATOM | 3569 | HD1 | ARG | A | 224 | -8.584 | 11.486 | 42.199 | 1.00 | 0.00 | |
| ATOM | 3570 | HD2 | ARG | A | 224 | -7.593 | 12.385 | 43.134 | 1.00 | 0.00 | |
| ATOM | 3571 | NE | ARG | A | 224 | -7.728 | 13.023 | 41.240 | 1.00 | 0.00 | N |
| ATOM | 3572 | HE | ARG | A | 224 | -8.533 | 13.092 | 40.650 | 1.00 | 0.00 | |
| ATOM | 3573 | CZ | ARG | A | 224 | -6.750 | 13.906 | 41.060 | 1.00 | 0.00 | C |
| ATOM | 3574 | NH1 | ARG | A | 224 | -5.726 | 13.933 | 41.895 | 1.00 | 0.00 | N |
| ATOM | 3575 | 1HH1 | ARG | A | 224 | -5.688 | 13.290 | 42.660 | 1.00 | 0.00 | |
| ATOM | 3576 | 2HH1 | ARG | A | 224 | -4.989 | 14.596 | 41.762 | 1.00 | 0.00 | |
| ATOM | 3577 | NH2 | ARG | A | 224 | -6.786 | 14.740 | 40.031 | 1.00 | 0.00 | N |
| ATOM | 3578 | 1HH2 | ARG | A | 224 | -7.550 | 14.707 | 39.387 | 1.00 | 0.00 | |
| ATOM | 3579 | 2HH2 | ARG | A | 224 | -6.049 | 15.403 | 39.899 | 1.00 | 0.00 | |
| ATOM | 3580 | C | ARG | A | 224 | -5.220 | 9.435 | 44.272 | 1.00 | 0.00 | C |
| ATOM | 3581 | O | ARG | A | 224 | -4.134 | 10.007 | 44.285 | 1.00 | 0.00 | O |
| ATOM | 3582 | N | MET | A | 225 | -6.007 | 9.318 | 45.350 | 1.00 | 0.00 | N |
| ATOM | 3583 | HN | MET | A | 225 | -6.822 | 8.742 | 45.284 | 1.00 | 0.00 | |
| ATOM | 3584 | CA | MET | A | 225 | -5.730 | 9.998 | 46.632 | 1.00 | 0.00 | C |
| ATOM | 3585 | HA | MET | A | 225 | -5.418 | 10.916 | 46.386 | 1.00 | 0.00 | |
| ATOM | 3586 | CB | MET | A | 225 | -7.002 | 10.113 | 47.481 | 1.00 | 0.00 | C |
| ATOM | 3587 | HB1 | MET | A | 225 | -7.393 | 9.200 | 47.601 | 1.00 | 0.00 | |
| ATOM | 3588 | HB2 | MET | A | 225 | -6.760 | 10.492 | 48.374 | 1.00 | 0.00 | |
| ATOM | 3589 | CG | MET | A | 225 | -8.055 | 11.016 | 46.834 | 1.00 | 0.00 | C |
| ATOM | 3590 | HG1 | MET | A | 225 | -7.602 | 11.903 | 46.744 | 1.00 | 0.00 | |
| ATOM | 3591 | HG2 | MET | A | 225 | -8.199 | 10.621 | 45.926 | 1.00 | 0.00 | |
| ATOM | 3592 | SD | MET | A | 225 | -9.599 | 11.137 | 47.771 | 1.00 | 0.00 | S |
| ATOM | 3593 | CE | MET | A | 225 | -10.649 | 11.991 | 46.589 | 1.00 | 0.00 | C |
| ATOM | 3594 | HE1 | MET | A | 225 | -11.556 | 12.127 | 46.987 | 1.00 | 0.00 | |
| ATOM | 3595 | HE2 | MET | A | 225 | -10.246 | 12.879 | 46.366 | 1.00 | 0.00 | |
| ATOM | 3596 | HE3 | MET | A | 225 | -10.729 | 11.442 | 45.757 | 1.00 | 0.00 | |
| ATOM | 3597 | C | MET | A | 225 | -4.623 | 9.262 | 47.413 | 1.00 | 0.00 | C |
| ATOM | 3598 | O | MET | A | 225 | -3.890 | 9.894 | 48.187 | 1.00 | 0.00 | O |
| ATOM | 3599 | N | PHE | A | 226 | -4.489 | 7.944 | 47.190 | 1.00 | 0.00 | N |
| ATOM | 3600 | HN | PHE | A | 226 | -5.191 | 7.473 | 46.655 | 1.00 | 0.00 | |
| ATOM | 3601 | CA | PHE | A | 226 | -3.346 | 7.163 | 47.704 | 1.00 | 0.00 | C |
| ATOM | 3602 | HA | PHE | A | 226 | -3.350 | 7.224 | 48.702 | 1.00 | 0.00 | |
| ATOM | 3603 | CB | PHE | A | 226 | -3.511 | 5.689 | 47.324 | 1.00 | 0.00 | C |
| ATOM | 3604 | HB1 | PHE | A | 226 | -4.396 | 5.379 | 47.672 | 1.00 | 0.00 | |
| ATOM | 3605 | HB2 | PHE | A | 226 | -3.505 | 5.626 | 46.326 | 1.00 | 0.00 | |
| ATOM | 3606 | CG | PHE | A | 226 | -2.466 | 4.729 | 47.839 | 1.00 | 0.00 | C |
| ATOM | 3607 | CD1 | PHE | A | 226 | -2.611 | 4.103 | 49.070 | 1.00 | 0.00 | C |
| ATOM | 3608 | HD1 | PHE | A | 226 | -3.395 | 4.327 | 49.649 | 1.00 | 0.00 | |
| ATOM | 3609 | CE1 | PHE | A | 226 | -1.679 | 3.168 | 49.506 | 1.00 | 0.00 | C |
| ATOM | 3610 | HE1 | PHE | A | 226 | -1.794 | 2.730 | 50.398 | 1.00 | 0.00 | |
| ATOM | 3611 | CZ | PHE | A | 226 | -0.596 | 2.840 | 48.715 | 1.00 | 0.00 | C |
| ATOM | 3612 | HZ | PHE | A | 226 | 0.076 | 2.171 | 49.033 | 1.00 | 0.00 | |
| ATOM | 3613 | CD2 | PHE | A | 226 | -1.380 | 4.378 | 47.052 | 1.00 | 0.00 | C |
| ATOM | 3614 | HD2 | PHE | A | 226 | -1.265 | 4.807 | 46.156 | 1.00 | 0.00 | |
| ATOM | 3615 | CE2 | PHE | A | 226 | -0.448 | 3.441 | 47.485 | 1.00 | 0.00 | C |
| ATOM | 3616 | HE2 | PHE | A | 226 | 0.330 | 3.204 | 46.903 | 1.00 | 0.00 | |

```
ATOM   3617  C   PHE A 226      -2.033   7.769  47.175  1.00  0.00           C
ATOM   3618  O   PHE A 226      -1.147   8.138  47.947  1.00  0.00           O
ATOM   3619  N   TYR A 227      -1.928   7.892  45.849  1.00  0.00           N
ATOM   3620  HN  TYR A 227      -2.704   7.620  45.280  1.00  0.00
ATOM   3621  CA  TYR A 227      -0.720   8.410  45.189  1.00  0.00           C
ATOM   3622  HA  TYR A 227       0.046   7.887  45.563  1.00  0.00
ATOM   3623  CB  TYR A 227      -0.794   8.189  43.673  1.00  0.00           C
ATOM   3624  HB1 TYR A 227      -1.705   8.461  43.363  1.00  0.00
ATOM   3625  HB2 TYR A 227      -0.107   8.772  43.240  1.00  0.00
ATOM   3626  CG  TYR A 227      -0.554   6.770  43.199  1.00  0.00           C
ATOM   3627  CD1 TYR A 227       0.373   5.930  43.806  1.00  0.00           C
ATOM   3628  HD1 TYR A 227       0.886   6.261  44.598  1.00  0.00
ATOM   3629  CE1 TYR A 227       0.605   4.643  43.340  1.00  0.00           C
ATOM   3630  HE1 TYR A 227       1.267   4.054  43.803  1.00  0.00
ATOM   3631  CZ  TYR A 227      -0.082   4.176  42.233  1.00  0.00           C
ATOM   3632  OH  TYR A 227       0.120   2.922  41.736  1.00  0.00           O
ATOM   3633  HH  TYR A 227      -0.469   2.780  40.940  1.00  0.00
ATOM   3634  CD2 TYR A 227      -1.209   6.288  42.078  1.00  0.00           C
ATOM   3635  HD2 TYR A 227      -1.853   6.882  41.597  1.00  0.00
ATOM   3636  CE2 TYR A 227      -0.994   5.003  41.606  1.00  0.00           C
ATOM   3637  HE2 TYR A 227      -1.500   4.673  40.809  1.00  0.00
ATOM   3638  C   TYR A 227      -0.535   9.898  45.510  1.00  0.00           C
ATOM   3639  O   TYR A 227       0.576  10.363  45.674  1.00  0.00           O
ATOM   3640  N   GLU A 228      -1.645  10.632  45.556  1.00  0.00           N
ATOM   3641  HN  GLU A 228      -2.520  10.163  45.435  1.00  0.00
ATOM   3642  CA  GLU A 228      -1.670  12.095  45.774  1.00  0.00           C
ATOM   3643  HA  GLU A 228      -1.114  12.524  45.062  1.00  0.00
ATOM   3644  CB  GLU A 228      -3.112  12.591  45.665  1.00  0.00           C
ATOM   3645  HB1 GLU A 228      -3.450  12.362  44.752  1.00  0.00
ATOM   3646  HB2 GLU A 228      -3.657  12.111  46.353  1.00  0.00
ATOM   3647  CG  GLU A 228      -3.319  14.078  45.871  1.00  0.00           C
ATOM   3648  HG1 GLU A 228      -2.787  14.364  46.668  1.00  0.00
ATOM   3649  HG2 GLU A 228      -2.988  14.559  45.059  1.00  0.00
ATOM   3650  CD  GLU A 228      -4.779  14.453  46.099  1.00  0.00           C
ATOM   3651  OE1 GLU A 228      -5.668  13.624  45.741  1.00  0.00           O
ATOM   3652  OE2 GLU A 228      -5.036  15.562  46.660  1.00  0.00           O
ATOM   3653  C   GLU A 228      -1.058  12.440  47.133  1.00  0.00           C
ATOM   3654  O   GLU A 228      -0.339  13.438  47.239  1.00  0.00           O
ATOM   3655  N   LYS A 229      -1.303  11.583  48.142  1.00  0.00           N
ATOM   3656  HN  LYS A 229      -1.841  10.762  47.953  1.00  0.00
ATOM   3657  CA  LYS A 229      -0.809  11.806  49.521  1.00  0.00           C
ATOM   3658  HA  LYS A 229      -0.607  12.785  49.520  1.00  0.00
ATOM   3659  CB  LYS A 229      -1.884  11.452  50.556  1.00  0.00           C
ATOM   3660  HB1 LYS A 229      -1.854  10.463  50.699  1.00  0.00
ATOM   3661  HB2 LYS A 229      -1.650  11.919  51.409  1.00  0.00
ATOM   3662  CG  LYS A 229      -3.324  11.821  50.212  1.00  0.00           C
ATOM   3663  HG1 LYS A 229      -3.316  12.332  49.353  1.00  0.00
ATOM   3664  HG2 LYS A 229      -3.840  10.973  50.091  1.00  0.00
ATOM   3665  CD  LYS A 229      -4.043  12.667  51.259  1.00  0.00           C
ATOM   3666  HD1 LYS A 229      -3.759  12.367  52.170  1.00  0.00
ATOM   3667  HD2 LYS A 229      -3.792  13.626  51.130  1.00  0.00
ATOM   3668  CE  LYS A 229      -5.547  12.544  51.157  1.00  0.00           C
ATOM   3669  HE1 LYS A 229      -5.807  12.428  50.198  1.00  0.00
ATOM   3670  HE2 LYS A 229      -5.852  11.750  51.683  1.00  0.00
ATOM   3671  NZ  LYS A 229      -6.227  13.749  51.686  1.00  0.00           N
ATOM   3672  HZ1 LYS A 229      -7.217  13.636  51.604  1.00  0.00
ATOM   3673  HZ2 LYS A 229      -5.984  13.874  52.648  1.00  0.00
ATOM   3674  HZ3 LYS A 229      -5.939  14.551  51.163  1.00  0.00
ATOM   3675  C   LYS A 229       0.475  11.004  49.867  1.00  0.00           C
ATOM   3676  O   LYS A 229       0.833  10.923  51.023  1.00  0.00           O
ATOM   3677  N   ASN A 230       1.173  10.469  48.861  1.00  0.00           N
ATOM   3678  HN  ASN A 230       0.860  10.671  47.933  1.00  0.00
ATOM   3679  CA  ASN A 230       2.328   9.633  48.989  1.00  0.00           C
ATOM   3680  HA  ASN A 230       2.389   9.176  48.102  1.00  0.00
ATOM   3681  CB  ASN A 230       3.587  10.459  49.222  1.00  0.00           C
ATOM   3682  HB1 ASN A 230       3.431  11.388  48.885  1.00  0.00
ATOM   3683  HB2 ASN A 230       3.777  10.488  50.203  1.00  0.00
ATOM   3684  CG  ASN A 230       4.796   9.876  48.506  1.00  0.00           C
ATOM   3685  OD1 ASN A 230       4.912   8.659  48.317  1.00  0.00           O
ATOM   3686  ND2 ASN A 230       5.694  10.737  48.068  1.00  0.00           N
ATOM   3687 1HD2 ASN A 230       5.560  11.717  48.214  1.00  0.00
```

```
ATOM   3688 2HD2 ASN A 230       6.509  10.410  47.590  1.00  0.00
ATOM   3689  C   ASN A 230       2.141   8.592  50.090  1.00  0.00           C
ATOM   3690  O   ASN A 230       3.113   8.193  50.738  1.00  0.00           O
ATOM   3691  N   ARG A 231       0.903   8.121  50.259  1.00  0.00           N
ATOM   3692  HN  ARG A 231       0.139   8.603  49.831  1.00  0.00
ATOM   3693  CA  ARG A 231       0.629   6.919  51.053  1.00  0.00           C
ATOM   3694  HA  ARG A 231       1.140   6.992  51.910  1.00  0.00
ATOM   3695  CB  ARG A 231      -0.861   6.831  51.410  1.00  0.00           C
ATOM   3696  HB1 ARG A 231      -1.384   6.747  50.562  1.00  0.00
ATOM   3697  HB2 ARG A 231      -1.003   6.016  51.971  1.00  0.00
ATOM   3698  CG  ARG A 231      -1.395   8.034  52.179  1.00  0.00           C
ATOM   3699  HG1 ARG A 231      -0.954   8.064  53.076  1.00  0.00
ATOM   3700  HG2 ARG A 231      -1.173   8.866  51.670  1.00  0.00
ATOM   3701  CD  ARG A 231      -2.888   7.991  52.387  1.00  0.00           C
ATOM   3702  HD1 ARG A 231      -3.357   7.988  51.504  1.00  0.00
ATOM   3703  HD2 ARG A 231      -3.139   7.170  52.900  1.00  0.00
ATOM   3704  NE  ARG A 231      -3.324   9.159  53.140  1.00  0.00           N
ATOM   3705  HE  ARG A 231      -2.619   9.765  53.509  1.00  0.00
ATOM   3706  CZ  ARG A 231      -4.594   9.476  53.371  1.00  0.00           C
ATOM   3707  NH1 ARG A 231      -5.566   8.663  52.984  1.00  0.00           N
ATOM   3708 1HH1 ARG A 231      -5.345   7.807  52.516  1.00  0.00
ATOM   3709 2HH1 ARG A 231      -6.520   8.905  53.160  1.00  0.00
ATOM   3710  NH2 ARG A 231      -4.891  10.613  53.976  1.00  0.00           N
ATOM   3711 1HH2 ARG A 231      -4.160  11.234  54.260  1.00  0.00
ATOM   3712 2HH2 ARG A 231      -5.846  10.852  54.150  1.00  0.00
ATOM   3713  C   ARG A 231       1.107   5.700  50.252  1.00  0.00           C
ATOM   3714  O   ARG A 231       0.442   4.664  50.222  1.00  0.00           O
ATOM   3715  N   SER A 232       2.278   5.829  49.611  1.00  0.00           N
ATOM   3716  HN  SER A 232       2.836   6.635  49.810  1.00  0.00
ATOM   3717  CA  SER A 232       2.791   4.860  48.636  1.00  0.00           C
ATOM   3718  HA  SER A 232       2.279   4.007  48.733  1.00  0.00
ATOM   3719  CB  SER A 232       2.545   5.369  47.234  1.00  0.00           C
ATOM   3720  HB1 SER A 232       1.564   5.371  47.042  1.00  0.00
ATOM   3721  HB2 SER A 232       2.904   6.298  47.144  1.00  0.00
ATOM   3722  OG  SER A 232       3.171   4.574  46.227  1.00  0.00           O
ATOM   3723  HG1 SER A 232       2.972   4.960  45.326  1.00  0.00
ATOM   3724  C   SER A 232       4.278   4.594  48.925  1.00  0.00           C
ATOM   3725  O   SER A 232       5.114   5.493  48.832  1.00  0.00           O
ATOM   3726  N   LEU A 233       4.561   3.352  49.335  1.00  0.00           N
ATOM   3727  HN  LEU A 233       3.804   2.702  49.407  1.00  0.00
ATOM   3728  CA  LEU A 233       5.896   2.880  49.684  1.00  0.00           C
ATOM   3729  HA  LEU A 233       6.512   3.666  49.646  1.00  0.00
ATOM   3730  CB  LEU A 233       5.872   2.287  51.101  1.00  0.00           C
ATOM   3731  HB1 LEU A 233       5.104   1.649  51.159  1.00  0.00
ATOM   3732  HB2 LEU A 233       6.729   1.792  51.247  1.00  0.00
ATOM   3733  CG  LEU A 233       5.722   3.299  52.229  1.00  0.00           C
ATOM   3734  HG  LEU A 233       5.004   3.947  51.976  1.00  0.00
ATOM   3735  CD1 LEU A 233       5.300   2.600  53.508  1.00  0.00           C
ATOM   3736 1HD1 LEU A 233       5.205   3.274  54.240  1.00  0.00
ATOM   3737 2HD1 LEU A 233       4.424   2.140  53.363  1.00  0.00
ATOM   3738 3HD1 LEU A 233       5.993   1.926  53.763  1.00  0.00
ATOM   3739  CD2 LEU A 233       7.014   4.090  52.437  1.00  0.00           C
ATOM   3740 1HD2 LEU A 233       6.887   4.745  53.182  1.00  0.00
ATOM   3741 2HD2 LEU A 233       7.756   3.461  52.669  1.00  0.00
ATOM   3742 3HD2 LEU A 233       7.243   4.581  51.596  1.00  0.00
ATOM   3743  C   LEU A 233       6.323   1.812  48.674  1.00  0.00           C
ATOM   3744  O   LEU A 233       5.500   1.024  48.211  1.00  0.00           O
ATOM   3745  N   MET A 234       7.623   1.745  48.393  1.00  0.00           N
ATOM   3746  HN  MET A 234       8.201   2.533  48.607  1.00  0.00
ATOM   3747  CA  MET A 234       8.238   0.574  47.788  1.00  0.00           C
ATOM   3748  HA  MET A 234       8.185   0.710  46.799  1.00  0.00
ATOM   3749  CB  MET A 234       9.703   0.444  48.221  1.00  0.00           C
ATOM   3750  HB1 MET A 234      10.015   1.332  48.559  1.00  0.00
ATOM   3751  HB2 MET A 234       9.759  -0.231  48.957  1.00  0.00
ATOM   3752  CG  MET A 234      10.626   0.001  47.075  1.00  0.00           C
ATOM   3753  HG1 MET A 234      11.491  -0.222  47.525  1.00  0.00
ATOM   3754  HG2 MET A 234      10.206  -0.837  46.727  1.00  0.00
ATOM   3755  SD  MET A 234      10.827   1.281  45.764  1.00  0.00           S
ATOM   3756  CE  MET A 234      12.037   2.357  46.540  1.00  0.00           C
ATOM   3757  HE1 MET A 234      12.248   3.118  45.927  1.00  0.00
ATOM   3758  HE2 MET A 234      11.665   2.713  47.397  1.00  0.00
```

```
ATOM   3759  HE3 MET A 234      12.871   1.839  46.728  1.00  0.00
ATOM   3760  C   MET A 234       7.466  -0.669  48.217  1.00  0.00           C
ATOM   3761  O   MET A 234       7.297  -0.917  49.402  1.00  0.00           O
ATOM   3762  N   PRO A 235       6.953  -1.486  47.270  1.00  0.00           N
ATOM   3763  CD  PRO A 235       7.048  -1.282  45.811  1.00  0.00           C
ATOM   3764  HD1 PRO A 235       8.003  -1.177  45.535  1.00  0.00
ATOM   3765  HD2 PRO A 235       6.531  -0.471  45.538  1.00  0.00
ATOM   3766  CA  PRO A 235       6.245  -2.717  47.621  1.00  0.00           C
ATOM   3767  HA  PRO A 235       5.679  -2.577  48.433  1.00  0.00
ATOM   3768  CB  PRO A 235       5.488  -3.019  46.320  1.00  0.00           C
ATOM   3769  HB1 PRO A 235       5.307  -3.998  46.229  1.00  0.00
ATOM   3770  HB2 PRO A 235       4.625  -2.515  46.282  1.00  0.00
ATOM   3771  CG  PRO A 235       6.442  -2.542  45.231  1.00  0.00           C
ATOM   3772  HG1 PRO A 235       7.150  -3.225  45.050  1.00  0.00
ATOM   3773  HG2 PRO A 235       5.948  -2.341  44.385  1.00  0.00
ATOM   3774  C   PRO A 235       7.201  -3.870  47.972  1.00  0.00           C
ATOM   3775  O   PRO A 235       8.340  -3.848  47.568  1.00  0.00           O
ATOM   3776  N   SER A 236       6.643  -4.930  48.544  1.00  0.00           N
ATOM   3777  HN  SER A 236       5.661  -4.891  48.728  1.00  0.00
ATOM   3778  CA  SER A 236       7.343  -6.132  48.920  1.00  0.00           C
ATOM   3779  HA  SER A 236       8.073  -5.816  49.526  1.00  0.00
ATOM   3780  CB  SER A 236       6.418  -7.044  49.665  1.00  0.00           C
ATOM   3781  HB1 SER A 236       5.554  -7.144  49.172  1.00  0.00
ATOM   3782  HB2 SER A 236       6.838  -7.943  49.787  1.00  0.00
ATOM   3783  OG  SER A 236       6.140  -6.501  50.950  1.00  0.00           O
ATOM   3784  HG1 SER A 236       5.522  -7.113  51.444  1.00  0.00
ATOM   3785  C   SER A 236       7.957  -6.859  47.712  1.00  0.00           C
ATOM   3786  O   SER A 236       7.245  -7.499  46.921  1.00  0.00           O
ATOM   3787  N   ILE A 237       9.295  -6.819  47.643  1.00  0.00           N
ATOM   3788  HN  ILE A 237       9.775  -6.168  48.231  1.00  0.00
ATOM   3789  CA  ILE A 237      10.099  -7.668  46.762  1.00  0.00           C
ATOM   3790  HA  ILE A 237       9.542  -7.829  45.947  1.00  0.00
ATOM   3791  CB  ILE A 237      11.413  -6.962  46.356  1.00  0.00           C
ATOM   3792  HB  ILE A 237      12.001  -6.919  47.163  1.00  0.00
ATOM   3793  CG2 ILE A 237      12.162  -7.774  45.285  1.00  0.00           C
ATOM   3794 1HG2 ILE A 237      13.007  -7.300  45.038  1.00  0.00
ATOM   3795 2HG2 ILE A 237      12.382  -8.680  45.647  1.00  0.00
ATOM   3796 3HG2 ILE A 237      11.584  -7.870  44.475  1.00  0.00
ATOM   3797  CG1 ILE A 237      11.193  -5.502  45.944  1.00  0.00           C
ATOM   3798 1HG1 ILE A 237      10.596  -5.060  46.614  1.00  0.00
ATOM   3799 2HG1 ILE A 237      12.076  -5.032  45.923  1.00  0.00
ATOM   3800  CD  ILE A 237      10.564  -5.345  44.604  1.00  0.00           C
ATOM   3801  HD1 ILE A 237      10.451  -4.372  44.401  1.00  0.00
ATOM   3802  HD2 ILE A 237      11.149  -5.765  43.910  1.00  0.00
ATOM   3803  HD3 ILE A 237       9.670  -5.792  44.601  1.00  0.00
ATOM   3804  C   ILE A 237      10.406  -8.987  47.481  1.00  0.00           C
ATOM   3805  O   ILE A 237      11.157  -9.006  48.453  1.00  0.00           O
ATOM   3806  N   PRO A 238       9.899 -10.137  46.992  1.00  0.00           N
ATOM   3807  CD  PRO A 238       8.978 -10.257  45.862  1.00  0.00           C
ATOM   3808  HD1 PRO A 238       9.437  -9.994  45.013  1.00  0.00
ATOM   3809  HD2 PRO A 238       8.180  -9.672  46.004  1.00  0.00
ATOM   3810  CA  PRO A 238      10.276 -11.438  47.530  1.00  0.00           C
ATOM   3811  HA  PRO A 238       9.884 -11.520  48.446  1.00  0.00
ATOM   3812  CB  PRO A 238       9.746 -12.445  46.492  1.00  0.00           C
ATOM   3813  HB1 PRO A 238      10.443 -12.665  45.809  1.00  0.00
ATOM   3814  HB2 PRO A 238       9.431 -13.287  46.930  1.00  0.00
ATOM   3815  CG  PRO A 238       8.595 -11.723  45.853  1.00  0.00           C
ATOM   3816  HG1 PRO A 238       8.461 -12.046  44.916  1.00  0.00
ATOM   3817  HG2 PRO A 238       7.758 -11.871  46.380  1.00  0.00
ATOM   3818  C   PRO A 238      11.798 -11.603  47.669  1.00  0.00           C
ATOM   3819  O   PRO A 238      12.568 -11.160  46.812  1.00  0.00           O
ATOM   3820  N   ARG A 239      12.192 -12.306  48.732  1.00  0.00           N
ATOM   3821  HN  ARG A 239      11.492 -12.737  49.302  1.00  0.00
ATOM   3822  CA  ARG A 239      13.577 -12.476  49.102  1.00  0.00           C
ATOM   3823  HA  ARG A 239      14.002 -11.574  49.027  1.00  0.00
ATOM   3824  CB  ARG A 239      13.665 -12.957  50.551  1.00  0.00           C
ATOM   3825  HB1 ARG A 239      13.107 -13.782  50.645  1.00  0.00
ATOM   3826  HB2 ARG A 239      14.620 -13.175  50.754  1.00  0.00
ATOM   3827  CG  ARG A 239      13.182 -11.938  51.573  1.00  0.00           C
ATOM   3828  HG1 ARG A 239      12.270 -11.626  51.308  1.00  0.00
ATOM   3829  HG2 ARG A 239      13.136 -12.380  52.469  1.00  0.00
```

```
ATOM   3830  CD  ARG A 239      14.095 -10.730  51.677  1.00  0.00           C
ATOM   3831  HD1 ARG A 239      14.846 -10.935  52.304  1.00  0.00
ATOM   3832  HD2 ARG A 239      14.465 -10.515  50.773  1.00  0.00
ATOM   3833  NE  ARG A 239      13.431  -9.535  52.170  1.00  0.00           N
ATOM   3834  HE  ARG A 239      12.549  -9.647  52.627  1.00  0.00
ATOM   3835  CZ  ARG A 239      13.918  -8.300  52.054  1.00  0.00           C
ATOM   3836  NH1 ARG A 239      15.013  -8.074  51.344  1.00  0.00           N
ATOM   3837 1HH1 ARG A 239      15.480  -8.833  50.890  1.00  0.00
ATOM   3838 2HH1 ARG A 239      15.372  -7.144  51.262  1.00  0.00
ATOM   3839  NH2 ARG A 239      13.306  -7.293  52.649  1.00  0.00           N
ATOM   3840 1HH2 ARG A 239      12.478  -7.458  53.186  1.00  0.00
ATOM   3841 2HH2 ARG A 239      13.670  -6.366  52.563  1.00  0.00
ATOM   3842  C   ARG A 239      14.275 -13.436  48.129  1.00  0.00           C
ATOM   3843  O   ARG A 239      15.489 -13.478  48.110  1.00  0.00           O
ATOM   3844  N   GLU A 240      13.504 -14.170  47.307  1.00  0.00           N
ATOM   3845  HN  GLU A 240      12.512 -14.099  47.416  1.00  0.00
ATOM   3846  CA  GLU A 240      14.018 -15.082  46.246  1.00  0.00           C
ATOM   3847  HA  GLU A 240      14.684 -15.676  46.697  1.00  0.00
ATOM   3848  CB  GLU A 240      12.878 -15.919  45.651  1.00  0.00           C
ATOM   3849  HB1 GLU A 240      12.159 -15.299  45.337  1.00  0.00
ATOM   3850  HB2 GLU A 240      13.236 -16.436  44.873  1.00  0.00
ATOM   3851  CG  GLU A 240      12.257 -16.905  46.627  1.00  0.00           C
ATOM   3852  HG1 GLU A 240      11.767 -17.595  46.094  1.00  0.00
ATOM   3853  HG2 GLU A 240      13.000 -17.341  47.135  1.00  0.00
ATOM   3854  CD  GLU A 240      11.281 -16.311  47.638  1.00  0.00           C
ATOM   3855  OE1 GLU A 240      10.647 -15.297  47.300  1.00  0.00           O
ATOM   3856  OE2 GLU A 240      11.176 -16.856  48.773  1.00  0.00           O
ATOM   3857  C   GLU A 240      14.685 -14.277  45.112  1.00  0.00           C
ATOM   3858  O   GLU A 240      15.484 -14.794  44.357  1.00  0.00           O
ATOM   3859  N   THR A 241      14.282 -13.012  44.991  1.00  0.00           N
ATOM   3860  HN  THR A 241      13.815 -12.591  45.769  1.00  0.00
ATOM   3861  CA  THR A 241      14.482 -12.200  43.787  1.00  0.00           C
ATOM   3862  HA  THR A 241      14.130 -12.737  43.020  1.00  0.00
ATOM   3863  CB  THR A 241      13.687 -10.892  43.896  1.00  0.00           C
ATOM   3864  HB  THR A 241      14.007 -10.347  44.671  1.00  0.00
ATOM   3865  OG1 THR A 241      12.306 -11.206  44.146  1.00  0.00           O
ATOM   3866  HG1 THR A 241      11.779 -10.359  44.219  1.00  0.00
ATOM   3867  CG2 THR A 241      13.817 -10.047  42.650  1.00  0.00           C
ATOM   3868 1HG2 THR A 241      13.286  -9.207  42.760  1.00  0.00
ATOM   3869 2HG2 THR A 241      14.779  -9.817  42.502  1.00  0.00
ATOM   3870 3HG2 THR A 241      13.472 -10.558  41.863  1.00  0.00
ATOM   3871  C   THR A 241      15.977 -11.933  43.564  1.00  0.00           C
ATOM   3872  O   THR A 241      16.664 -11.502  44.456  1.00  0.00           O
ATOM   3873  N   SER A 242      16.447 -12.157  42.339  1.00  0.00           N
ATOM   3874  HN  SER A 242      15.816 -12.497  41.642  1.00  0.00
ATOM   3875  CA  SER A 242      17.857 -11.927  41.958  1.00  0.00           C
ATOM   3876  HA  SER A 242      18.411 -12.491  42.570  1.00  0.00
ATOM   3877  CB  SER A 242      18.147 -12.385  40.559  1.00  0.00           C
ATOM   3878  HB1 SER A 242      19.132 -12.497  40.430  1.00  0.00
ATOM   3879  HB2 SER A 242      17.687 -13.254  40.379  1.00  0.00
ATOM   3880  OG  SER A 242      17.683 -11.435  39.603  1.00  0.00           O
ATOM   3881  HG1 SER A 242      17.888 -11.763  38.681  1.00  0.00
ATOM   3882  C   SER A 242      18.203 -10.452  42.121  1.00  0.00           C
ATOM   3883  O   SER A 242      17.342  -9.579  41.988  1.00  0.00           O
ATOM   3884  N   PRO A 243      19.478 -10.126  42.424  1.00  0.00           N
ATOM   3885  CD  PRO A 243      20.602 -11.065  42.579  1.00  0.00           C
ATOM   3886  HD1 PRO A 243      20.975 -11.320  41.687  1.00  0.00
ATOM   3887  HD2 PRO A 243      20.310 -11.888  43.066  1.00  0.00
ATOM   3888  CA  PRO A 243      19.870  -8.744  42.694  1.00  0.00           C
ATOM   3889  HA  PRO A 243      19.360  -8.407  43.486  1.00  0.00
ATOM   3890  CB  PRO A 243      21.389  -8.840  42.903  1.00  0.00           C
ATOM   3891  HB1 PRO A 243      21.882  -8.688  42.046  1.00  0.00
ATOM   3892  HB2 PRO A 243      21.702  -8.183  43.589  1.00  0.00
ATOM   3893  CG  PRO A 243      21.594 -10.256  43.391  1.00  0.00           C
ATOM   3894  HG1 PRO A 243      22.531 -10.560  43.217  1.00  0.00
ATOM   3895  HG2 PRO A 243      21.397 -10.327  44.369  1.00  0.00
ATOM   3896  C   PRO A 243      19.517  -7.777  41.553  1.00  0.00           C
ATOM   3897  O   PRO A 243      19.065  -6.656  41.803  1.00  0.00           O
ATOM   3898  N   TYR A 244      19.704  -8.229  40.307  1.00  0.00           N
ATOM   3899  HN  TYR A 244      20.000  -9.175  40.174  1.00  0.00
ATOM   3900  CA  TYR A 244      19.491  -7.389  39.142  1.00  0.00           C
```

```
ATOM   3901  HA  TYR A 244      19.962  -6.525  39.321  1.00  0.00
ATOM   3902  CB  TYR A 244      20.124  -8.047  37.919  1.00  0.00           C
ATOM   3903  HB1 TYR A 244      19.893  -9.020  37.933  1.00  0.00
ATOM   3904  HB2 TYR A 244      19.735  -7.624  37.101  1.00  0.00
ATOM   3905  CG  TYR A 244      21.626  -7.934  37.823  1.00  0.00           C
ATOM   3906  CD1 TYR A 244      22.319  -6.861  38.370  1.00  0.00           C
ATOM   3907  HD1 TYR A 244      21.812  -6.139  38.841  1.00  0.00
ATOM   3908  CE1 TYR A 244      23.703  -6.770  38.276  1.00  0.00           C
ATOM   3909  HE1 TYR A 244      24.182  -5.994  38.686  1.00  0.00
ATOM   3910  CZ  TYR A 244      24.422  -7.759  37.611  1.00  0.00           C
ATOM   3911  OH  TYR A 244      25.775  -7.643  37.466  1.00  0.00           O
ATOM   3912  HH  TYR A 244      26.126  -8.436  36.968  1.00  0.00
ATOM   3913  CD2 TYR A 244      22.363  -8.909  37.162  1.00  0.00           C
ATOM   3914  HD2 TYR A 244      21.885  -9.687  36.754  1.00  0.00
ATOM   3915  CE2 TYR A 244      23.747  -8.832  37.052  1.00  0.00           C
ATOM   3916  HE2 TYR A 244      24.253  -9.548  36.571  1.00  0.00
ATOM   3917  C   TYR A 244      17.992  -7.089  38.952  1.00  0.00           C
ATOM   3918  O   TYR A 244      17.623  -5.951  38.699  1.00  0.00           O
ATOM   3919  N   LEU A 245      17.131  -8.102  39.075  1.00  0.00           N
ATOM   3920  HN  LEU A 245      17.476  -9.016  39.287  1.00  0.00
ATOM   3921  CA  LEU A 245      15.691  -7.899  38.905  1.00  0.00           C
ATOM   3922  HA  LEU A 245      15.564  -7.481  38.005  1.00  0.00
ATOM   3923  CB  LEU A 245      14.952  -9.243  38.948  1.00  0.00           C
ATOM   3924  HB1 LEU A 245      15.367  -9.848  38.268  1.00  0.00
ATOM   3925  HB2 LEU A 245      15.072  -9.635  39.860  1.00  0.00
ATOM   3926  CG  LEU A 245      13.456  -9.176  38.662  1.00  0.00           C
ATOM   3927  HG  LEU A 245      13.011  -8.705  39.424  1.00  0.00
ATOM   3928  CD1 LEU A 245      13.180  -8.376  37.403  1.00  0.00           C
ATOM   3929 1HD1 LEU A 245      12.194  -8.346  37.236  1.00  0.00
ATOM   3930 2HD1 LEU A 245      13.527  -7.445  37.517  1.00  0.00
ATOM   3931 3HD1 LEU A 245      13.636  -8.809  36.626  1.00  0.00
ATOM   3932  CD2 LEU A 245      12.866 -10.575  38.557  1.00  0.00           C
ATOM   3933 1HD2 LEU A 245      11.886 -10.510  38.370  1.00  0.00
ATOM   3934 2HD2 LEU A 245      13.315 -11.071  37.814  1.00  0.00
ATOM   3935 3HD2 LEU A 245      13.009 -11.063  39.418  1.00  0.00
ATOM   3936  C   LEU A 245      15.175  -6.960  40.002  1.00  0.00           C
ATOM   3937  O   LEU A 245      14.313  -6.107  39.749  1.00  0.00           O
ATOM   3938  N   ALA A 246      15.704  -7.124  41.218  1.00  0.00           N
ATOM   3939  HN  ALA A 246      16.410  -7.820  41.351  1.00  0.00
ATOM   3940  CA  ALA A 246      15.282  -6.320  42.352  1.00  0.00           C
ATOM   3941  HA  ALA A 246      14.286  -6.383  42.409  1.00  0.00
ATOM   3942  CB  ALA A 246      15.848  -6.880  43.633  1.00  0.00           C
ATOM   3943  HB1 ALA A 246      15.549  -6.317  44.404  1.00  0.00
ATOM   3944  HB2 ALA A 246      15.523  -7.817  43.759  1.00  0.00
ATOM   3945  HB3 ALA A 246      16.847  -6.877  43.585  1.00  0.00
ATOM   3946  C   ALA A 246      15.701  -4.867  42.125  1.00  0.00           C
ATOM   3947  O   ALA A 246      14.931  -3.972  42.350  1.00  0.00           O
ATOM   3948  N   ASN A 247      16.933  -4.658  41.656  1.00  0.00           N
ATOM   3949  HN  ASN A 247      17.530  -5.444  41.498  1.00  0.00
ATOM   3950  CA  ASN A 247      17.440  -3.309  41.364  1.00  0.00           C
ATOM   3951  HA  ASN A 247      17.414  -2.803  42.226  1.00  0.00
ATOM   3952  CB  ASN A 247      18.901  -3.327  40.897  1.00  0.00           C
ATOM   3953  HB1 ASN A 247      19.473  -3.644  41.653  1.00  0.00
ATOM   3954  HB2 ASN A 247      18.982  -3.962  40.129  1.00  0.00
ATOM   3955  CG  ASN A 247      19.414  -1.962  40.447  1.00  0.00           C
ATOM   3956  OD1 ASN A 247      19.199  -1.562  39.306  1.00  0.00           O
ATOM   3957  ND2 ASN A 247      20.102  -1.236  41.324  1.00  0.00           N
ATOM   3958 1HD2 ASN A 247      20.268  -1.589  42.245  1.00  0.00
ATOM   3959 2HD2 ASN A 247      20.453  -0.337  41.063  1.00  0.00
ATOM   3960  C   ASN A 247      16.522  -2.656  40.323  1.00  0.00           C
ATOM   3961  O   ASN A 247      16.150  -1.502  40.454  1.00  0.00           O
ATOM   3962  N   LEU A 248      16.152  -3.434  39.297  1.00  0.00           N
ATOM   3963  HN  LEU A 248      16.469  -4.382  39.279  1.00  0.00
ATOM   3964  CA  LEU A 248      15.319  -2.968  38.220  1.00  0.00           C
ATOM   3965  HA  LEU A 248      15.791  -2.177  37.832  1.00  0.00
ATOM   3966  CB  LEU A 248      15.189  -4.081  37.182  1.00  0.00           C
ATOM   3967  HB1 LEU A 248      16.106  -4.307  36.852  1.00  0.00
ATOM   3968  HB2 LEU A 248      14.790  -4.880  37.632  1.00  0.00
ATOM   3969  CG  LEU A 248      14.324  -3.739  35.974  1.00  0.00           C
ATOM   3970  HG  LEU A 248      13.376  -3.618  36.268  1.00  0.00
ATOM   3971  CD1 LEU A 248      14.783  -2.428  35.324  1.00  0.00           C
```

```
ATOM   3972 1HD1 LEU A 248      14.202  -2.225  34.536  1.00  0.00
ATOM   3973 2HD1 LEU A 248      14.714  -1.684  35.989  1.00  0.00
ATOM   3974 3HD1 LEU A 248      15.732  -2.520  35.023  1.00  0.00
ATOM   3975  CD2 LEU A 248      14.368  -4.882  34.992  1.00  0.00           C
ATOM   3976 1HD2 LEU A 248      13.802  -4.661  34.198  1.00  0.00
ATOM   3977 2HD2 LEU A 248      15.312  -5.032  34.697  1.00  0.00
ATOM   3978 3HD2 LEU A 248      14.020  -5.711  35.430  1.00  0.00
ATOM   3979  C   LEU A 248      13.938  -2.525  38.738  1.00  0.00           C
ATOM   3980  O   LEU A 248      13.484  -1.441  38.421  1.00  0.00           O
ATOM   3981  N   LEU A 249      13.279  -3.383  39.522  1.00  0.00           N
ATOM   3982  HN  LEU A 249      13.711  -4.255  39.752  1.00  0.00
ATOM   3983  CA  LEU A 249      11.955  -3.094  40.053  1.00  0.00           C
ATOM   3984  HA  LEU A 249      11.359  -2.922  39.269  1.00  0.00
ATOM   3985  CB  LEU A 249      11.441  -4.302  40.839  1.00  0.00           C
ATOM   3986  HB1 LEU A 249      12.172  -4.612  41.447  1.00  0.00
ATOM   3987  HB2 LEU A 249      10.656  -4.007  41.385  1.00  0.00
ATOM   3988  CG  LEU A 249      11.003  -5.488  39.985  1.00  0.00           C
ATOM   3989  HG  LEU A 249      11.679  -5.618  39.259  1.00  0.00
ATOM   3990  CD1 LEU A 249      10.961  -6.769  40.805  1.00  0.00           C
ATOM   3991 1HD1 LEU A 249      10.672  -7.528  40.222  1.00  0.00
ATOM   3992 2HD1 LEU A 249      11.871  -6.959  41.173  1.00  0.00
ATOM   3993 3HD1 LEU A 249      10.313  -6.661  41.559  1.00  0.00
ATOM   3994  CD2 LEU A 249       9.647  -5.209  39.342  1.00  0.00           C
ATOM   3995 1HD2 LEU A 249       9.374  -5.994  38.786  1.00  0.00
ATOM   3996 2HD2 LEU A 249       8.965  -5.053  40.057  1.00  0.00
ATOM   3997 3HD2 LEU A 249       9.714  -4.397  38.762  1.00  0.00
ATOM   3998  C   LEU A 249      12.010  -1.846  40.936  1.00  0.00           C
ATOM   3999  O   LEU A 249      11.089  -1.038  40.886  1.00  0.00           O
ATOM   4000  N   LEU A 250      13.072  -1.698  41.742  1.00  0.00           N
ATOM   4001  HN  LEU A 250      13.819  -2.360  41.687  1.00  0.00
ATOM   4002  CA  LEU A 250      13.170  -0.588  42.706  1.00  0.00           C
ATOM   4003  HA  LEU A 250      12.334  -0.622  43.254  1.00  0.00
ATOM   4004  CB  LEU A 250      14.381  -0.750  43.634  1.00  0.00           C
ATOM   4005  HB1 LEU A 250      15.171  -0.981  43.066  1.00  0.00
ATOM   4006  HB2 LEU A 250      14.539   0.129  44.084  1.00  0.00
ATOM   4007  CG  LEU A 250      14.272  -1.814  44.726  1.00  0.00           C
ATOM   4008  HG  LEU A 250      14.036  -2.681  44.286  1.00  0.00
ATOM   4009  CD1 LEU A 250      15.609  -1.991  45.464  1.00  0.00           C
ATOM   4010 1HD1 LEU A 250      15.509  -2.691  46.171  1.00  0.00
ATOM   4011 2HD1 LEU A 250      16.314  -2.272  44.813  1.00  0.00
ATOM   4012 3HD1 LEU A 250      15.873  -1.125  45.888  1.00  0.00
ATOM   4013  CD2 LEU A 250      13.159  -1.476  45.702  1.00  0.00           C
ATOM   4014 1HD2 LEU A 250      13.106  -2.184  46.406  1.00  0.00
ATOM   4015 2HD2 LEU A 250      13.349  -0.593  46.131  1.00  0.00
ATOM   4016 3HD2 LEU A 250      12.289  -1.428  45.212  1.00  0.00
ATOM   4017  C   LEU A 250      13.279   0.733  41.941  1.00  0.00           C
ATOM   4018  O   LEU A 250      12.757   1.747  42.386  1.00  0.00           O
ATOM   4019  N   GLY A 251      13.983   0.700  40.805  1.00  0.00           N
ATOM   4020  HN  GLY A 251      14.326  -0.182  40.482  1.00  0.00
ATOM   4021  CA  GLY A 251      14.273   1.886  40.019  1.00  0.00           C
ATOM   4022  HA1 GLY A 251      14.457   2.635  40.655  1.00  0.00
ATOM   4023  HA2 GLY A 251      15.093   1.699  39.477  1.00  0.00
ATOM   4024  C   GLY A 251      13.128   2.268  39.098  1.00  0.00           C
ATOM   4025  O   GLY A 251      13.019   3.426  38.705  1.00  0.00           O
ATOM   4026  N   LEU A 252      12.310   1.280  38.714  1.00  0.00           N
ATOM   4027  HN  LEU A 252      12.524   0.343  38.991  1.00  0.00
ATOM   4028  CA  LEU A 252      11.120   1.516  37.908  1.00  0.00           C
ATOM   4029  HA  LEU A 252      11.354   2.223  37.241  1.00  0.00
ATOM   4030  CB  LEU A 252      10.731   0.232  37.179  1.00  0.00           C
ATOM   4031  HB1 LEU A 252      10.700  -0.507  37.852  1.00  0.00
ATOM   4032  HB2 LEU A 252       9.821   0.364  36.787  1.00  0.00
ATOM   4033  CG  LEU A 252      11.664  -0.200  36.055  1.00  0.00           C
ATOM   4034  HG  LEU A 252      12.593  -0.242  36.422  1.00  0.00
ATOM   4035  CD1 LEU A 252      11.270  -1.573  35.551  1.00  0.00           C
ATOM   4036 1HD1 LEU A 252      11.888  -1.847  34.814  1.00  0.00
ATOM   4037 2HD1 LEU A 252      11.330  -2.233  36.300  1.00  0.00
ATOM   4038 3HD1 LEU A 252      10.332  -1.545  35.206  1.00  0.00
ATOM   4039  CD2 LEU A 252      11.671   0.800  34.914  1.00  0.00           C
ATOM   4040 1HD2 LEU A 252      12.293   0.484  34.197  1.00  0.00
ATOM   4041 2HD2 LEU A 252      10.747   0.886  34.540  1.00  0.00
ATOM   4042 3HD2 LEU A 252      11.978   1.690  35.252  1.00  0.00
```

```
ATOM   4043  C   LEU A 252       9.976   1.999  38.810  1.00  0.00           C
ATOM   4044  O   LEU A 252       9.239   2.904  38.432  1.00  0.00           O
ATOM   4045  N   LEU A 253       9.844   1.395  39.993  1.00  0.00           N
ATOM   4046  HN  LEU A 253      10.547   0.748  40.289  1.00  0.00
ATOM   4047  CA  LEU A 253       8.692   1.654  40.876  1.00  0.00           C
ATOM   4048  HA  LEU A 253       7.931   1.944  40.296  1.00  0.00
ATOM   4049  CB  LEU A 253       8.308   0.369  41.616  1.00  0.00           C
ATOM   4050  HB1 LEU A 253       9.131   0.000  42.048  1.00  0.00
ATOM   4051  HB2 LEU A 253       7.639   0.604  42.321  1.00  0.00
ATOM   4052  CG  LEU A 253       7.706  -0.727  40.749  1.00  0.00           C
ATOM   4053  HG  LEU A 253       8.364  -0.954  40.031  1.00  0.00
ATOM   4054  CD1 LEU A 253       7.450  -1.989  41.567  1.00  0.00           C
ATOM   4055 1HD1 LEU A 253       7.055  -2.694  40.978  1.00  0.00
ATOM   4056 2HD1 LEU A 253       8.313  -2.319  41.949  1.00  0.00
ATOM   4057 3HD1 LEU A 253       6.814  -1.781  42.310  1.00  0.00
ATOM   4058  CD2 LEU A 253       6.418  -0.236  40.092  1.00  0.00           C
ATOM   4059 1HD2 LEU A 253       6.034  -0.965  39.526  1.00  0.00
ATOM   4060 2HD2 LEU A 253       5.759   0.019  40.800  1.00  0.00
ATOM   4061 3HD2 LEU A 253       6.618   0.560  39.520  1.00  0.00
ATOM   4062  C   LEU A 253       9.032   2.776  41.863  1.00  0.00           C
ATOM   4063  O   LEU A 253       8.900   2.604  43.069  1.00  0.00           O
ATOM   4064  N   GLN A 254       9.433   3.931  41.311  1.00  0.00           N
ATOM   4065  HN  GLN A 254       9.604   3.937  40.326  1.00  0.00
ATOM   4066  CA  GLN A 254       9.637   5.172  42.035  1.00  0.00           C
ATOM   4067  HA  GLN A 254       9.701   4.931  43.003  1.00  0.00
ATOM   4068  CB  GLN A 254      10.927   5.842  41.571  1.00  0.00           C
ATOM   4069  HB1 GLN A 254      10.895   5.940  40.576  1.00  0.00
ATOM   4070  HB2 GLN A 254      10.985   6.747  41.993  1.00  0.00
ATOM   4071  CG  GLN A 254      12.180   5.063  41.935  1.00  0.00           C
ATOM   4072  HG1 GLN A 254      12.038   4.101  41.702  1.00  0.00
ATOM   4073  HG2 GLN A 254      12.947   5.424  41.404  1.00  0.00
ATOM   4074  CD  GLN A 254      12.529   5.154  43.404  1.00  0.00           C
ATOM   4075  OE1 GLN A 254      12.230   6.138  44.068  1.00  0.00           O
ATOM   4076  NE2 GLN A 254      13.169   4.119  43.918  1.00  0.00           N
ATOM   4077 1HE2 GLN A 254      13.395   3.334  43.341  1.00  0.00
ATOM   4078 2HE2 GLN A 254      13.427   4.120  44.884  1.00  0.00
ATOM   4079  C   GLN A 254       8.455   6.103  41.777  1.00  0.00           C
ATOM   4080  O   GLN A 254       8.200   6.472  40.629  1.00  0.00           O
ATOM   4081  N   ARG A 255       7.772   6.502  42.856  1.00  0.00           N
ATOM   4082  HN  ARG A 255       8.110   6.228  43.756  1.00  0.00
ATOM   4083  CA  ARG A 255       6.561   7.315  42.795  1.00  0.00           C
ATOM   4084  HA  ARG A 255       5.893   6.802  42.256  1.00  0.00
ATOM   4085  CB  ARG A 255       5.986   7.529  44.201  1.00  0.00           C
ATOM   4086  HB1 ARG A 255       5.910   6.638  44.648  1.00  0.00
ATOM   4087  HB2 ARG A 255       6.620   8.107  44.715  1.00  0.00
ATOM   4088  CG  ARG A 255       4.611   8.192  44.230  1.00  0.00           C
ATOM   4089  HG1 ARG A 255       4.086   7.870  43.442  1.00  0.00
ATOM   4090  HG2 ARG A 255       4.146   7.923  45.073  1.00  0.00
ATOM   4091  CD  ARG A 255       4.671   9.712  44.178  1.00  0.00           C
ATOM   4092  HD1 ARG A 255       5.577  10.016  44.473  1.00  0.00
ATOM   4093  HD2 ARG A 255       4.507  10.016  43.240  1.00  0.00
ATOM   4094  NE  ARG A 255       3.694  10.382  45.026  1.00  0.00           N
ATOM   4095  HE  ARG A 255       2.966   9.824  45.425  1.00  0.00
ATOM   4096  CZ  ARG A 255       3.700  11.688  45.309  1.00  0.00           C
ATOM   4097  NH1 ARG A 255       4.685  12.466  44.888  1.00  0.00           N
ATOM   4098 1HH1 ARG A 255       5.435  12.078  44.352  1.00  0.00
ATOM   4099 2HH1 ARG A 255       4.679  13.442  45.106  1.00  0.00
ATOM   4100  NH2 ARG A 255       2.710  12.214  46.011  1.00  0.00           N
ATOM   4101 1HH2 ARG A 255       1.959  11.635  46.329  1.00  0.00
ATOM   4102 2HH2 ARG A 255       2.712  13.191  46.224  1.00  0.00
ATOM   4103  C   ARG A 255       6.867   8.638  42.088  1.00  0.00           C
ATOM   4104  O   ARG A 255       6.031   9.160  41.376  1.00  0.00           O
ATOM   4105  N   ASN A 256       8.071   9.166  42.283  1.00  0.00           N
ATOM   4106  HN  ASN A 256       8.732   8.670  42.846  1.00  0.00
ATOM   4107  CA  ASN A 256       8.458  10.444  41.702  1.00  0.00           C
ATOM   4108  HA  ASN A 256       7.612  10.903  41.430  1.00  0.00
ATOM   4109  CB  ASN A 256       9.151  11.326  42.740  1.00  0.00           C
ATOM   4110  HB1 ASN A 256       8.638  11.272  43.596  1.00  0.00
ATOM   4111  HB2 ASN A 256      10.078  10.979  42.884  1.00  0.00
ATOM   4112  CG  ASN A 256       9.252  12.783  42.344  1.00  0.00           C
ATOM   4113  OD1 ASN A 256       8.947  13.161  41.211  1.00  0.00           O
```

```
ATOM   4114  ND2 ASN A 256       9.680  13.610  43.279  1.00  0.00           N
ATOM   4115 1HD2 ASN A 256       9.915  13.261  44.186  1.00  0.00
ATOM   4116 2HD2 ASN A 256       9.769  14.586  43.081  1.00  0.00
ATOM   4117  C   ASN A 256       9.329  10.201  40.461  1.00  0.00           C
ATOM   4118  O   ASN A 256      10.352   9.512  40.526  1.00  0.00           O
ATOM   4119  N   GLN A 257       8.899  10.789  39.335  1.00  0.00           N
ATOM   4120  HN  GLN A 257       8.088  11.371  39.388  1.00  0.00
ATOM   4121  CA  GLN A 257       9.551  10.628  38.027  1.00  0.00           C
ATOM   4122  HA  GLN A 257       9.604   9.648  37.834  1.00  0.00
ATOM   4123  CB  GLN A 257       8.702  11.276  36.920  1.00  0.00           C
ATOM   4124  HB1 GLN A 257       9.194  11.177  36.055  1.00  0.00
ATOM   4125  HB2 GLN A 257       7.834  10.782  36.868  1.00  0.00
ATOM   4126  CG  GLN A 257       8.385  12.763  37.111  1.00  0.00           C
ATOM   4127  HG1 GLN A 257       9.066  13.155  37.729  1.00  0.00
ATOM   4128  HG2 GLN A 257       8.443  13.216  36.221  1.00  0.00
ATOM   4129  CD  GLN A 257       7.003  13.033  37.693  1.00  0.00           C
ATOM   4130  OE1 GLN A 257       6.613  12.476  38.719  1.00  0.00           O
ATOM   4131  NE2 GLN A 257       6.244  13.917  37.054  1.00  0.00           N
ATOM   4132 1HE2 GLN A 257       6.588  14.370  36.231  1.00  0.00
ATOM   4133 2HE2 GLN A 257       5.329  14.130  37.396  1.00  0.00
ATOM   4134  C   GLN A 257      10.978  11.194  38.097  1.00  0.00           C
ATOM   4135  O   GLN A 257      11.902  10.715  37.431  1.00  0.00           O
ATOM   4136  N   LYS A 258      11.138  12.215  38.935  1.00  0.00           N
ATOM   4137  HN  LYS A 258      10.316  12.617  39.337  1.00  0.00
ATOM   4138  CA  LYS A 258      12.421  12.789  39.309  1.00  0.00           C
ATOM   4139  HA  LYS A 258      12.819  13.239  38.510  1.00  0.00
ATOM   4140  CB  LYS A 258      12.150  13.833  40.403  1.00  0.00           C
ATOM   4141  HB1 LYS A 258      11.522  14.516  40.030  1.00  0.00
ATOM   4142  HB2 LYS A 258      11.716  13.367  41.174  1.00  0.00
ATOM   4143  CG  LYS A 258      13.364  14.574  40.941  1.00  0.00           C
ATOM   4144  HG1 LYS A 258      13.883  13.948  41.524  1.00  0.00
ATOM   4145  HG2 LYS A 258      13.931  14.855  40.167  1.00  0.00
ATOM   4146  CD  LYS A 258      13.004  15.814  41.757  1.00  0.00           C
ATOM   4147  HD1 LYS A 258      12.695  16.531  41.132  1.00  0.00
ATOM   4148  HD2 LYS A 258      12.264  15.580  42.387  1.00  0.00
ATOM   4149  CE  LYS A 258      14.164  16.355  42.566  1.00  0.00           C
ATOM   4150  HE1 LYS A 258      14.996  16.321  42.012  1.00  0.00
ATOM   4151  HE2 LYS A 258      13.973  17.302  42.822  1.00  0.00
ATOM   4152  NZ  LYS A 258      14.396  15.573  43.806  1.00  0.00           N
ATOM   4153  HZ1 LYS A 258      15.168  15.964  44.307  1.00  0.00
ATOM   4154  HZ2 LYS A 258      13.577  15.605  44.378  1.00  0.00
ATOM   4155  HZ3 LYS A 258      14.600  14.623  43.568  1.00  0.00
ATOM   4156  C   LYS A 258      13.407  11.676  39.730  1.00  0.00           C
ATOM   4157  O   LYS A 258      14.569  11.710  39.349  1.00  0.00           O
ATOM   4158  N   ASP A 259      12.930  10.677  40.482  1.00  0.00           N
ATOM   4159  HN  ASP A 259      11.942  10.622  40.625  1.00  0.00
ATOM   4160  CA  ASP A 259      13.796   9.639  41.120  1.00  0.00           C
ATOM   4161  HA  ASP A 259      14.711  10.042  41.109  1.00  0.00
ATOM   4162  CB  ASP A 259      13.336   9.350  42.548  1.00  0.00           C
ATOM   4163  HB1 ASP A 259      12.366   9.107  42.521  1.00  0.00
ATOM   4164  HB2 ASP A 259      13.864   8.576  42.896  1.00  0.00
ATOM   4165  CG  ASP A 259      13.504  10.508  43.518  1.00  0.00           C
ATOM   4166  OD1 ASP A 259      14.545  11.215  43.448  1.00  0.00           O
ATOM   4167  OD2 ASP A 259      12.617  10.668  44.368  1.00  0.00           O
ATOM   4168  C   ASP A 259      13.786   8.302  40.346  1.00  0.00           C
ATOM   4169  O   ASP A 259      14.480   7.373  40.712  1.00  0.00           O
ATOM   4170  N   ARG A 260      12.978   8.206  39.289  1.00  0.00           N
ATOM   4171  HN  ARG A 260      12.502   9.028  38.976  1.00  0.00
ATOM   4172  CA  ARG A 260      12.758   6.965  38.574  1.00  0.00           C
ATOM   4173  HA  ARG A 260      12.787   6.229  39.250  1.00  0.00
ATOM   4174  CB  ARG A 260      11.381   7.005  37.908  1.00  0.00           C
ATOM   4175  HB1 ARG A 260      10.713   7.304  38.589  1.00  0.00
ATOM   4176  HB2 ARG A 260      11.414   7.668  37.160  1.00  0.00
ATOM   4177  CG  ARG A 260      10.908   5.676  37.338  1.00  0.00           C
ATOM   4178  HG1 ARG A 260      11.539   5.385  36.618  1.00  0.00
ATOM   4179  HG2 ARG A 260      10.896   4.992  38.068  1.00  0.00
ATOM   4180  CD  ARG A 260       9.509   5.805  36.753  1.00  0.00           C
ATOM   4181  HD1 ARG A 260       9.528   6.426  35.969  1.00  0.00
ATOM   4182  HD2 ARG A 260       9.184   4.906  36.458  1.00  0.00
ATOM   4183  NE  ARG A 260       8.546   6.330  37.707  1.00  0.00           N
ATOM   4184  HE  ARG A 260       8.683   6.105  38.672  1.00  0.00
```

```
ATOM   4185  CZ  ARG A 260       7.496   7.086  37.397  1.00  0.00           C
ATOM   4186  NH1 ARG A 260       7.202   7.364  36.138  1.00  0.00           N
ATOM   4187 1HH1 ARG A 260       7.773   7.004  35.400  1.00  0.00
ATOM   4188 2HH1 ARG A 260       6.409   7.934  35.924  1.00  0.00
ATOM   4189  NH2 ARG A 260       6.725   7.550  38.358  1.00  0.00           N
ATOM   4190 1HH2 ARG A 260       6.931   7.334  39.312  1.00  0.00
ATOM   4191 2HH2 ARG A 260       5.934   8.119  38.134  1.00  0.00
ATOM   4192  C   ARG A 260      13.873   6.756  37.540  1.00  0.00           C
ATOM   4193  O   ARG A 260      14.279   7.666  36.852  1.00  0.00           O
ATOM   4194  N   MET A 261      14.349   5.521  37.449  1.00  0.00           N
ATOM   4195  HN  MET A 261      14.046   4.849  38.125  1.00  0.00
ATOM   4196  CA  MET A 261      15.285   5.086  36.426  1.00  0.00           C
ATOM   4197  HA  MET A 261      16.194   5.406  36.692  1.00  0.00
ATOM   4198  CB  MET A 261      15.244   3.562  36.327  1.00  0.00           C
ATOM   4199  HB1 MET A 261      15.457   3.181  37.227  1.00  0.00
ATOM   4200  HB2 MET A 261      14.320   3.288  36.059  1.00  0.00
ATOM   4201  CG  MET A 261      16.216   2.971  35.327  1.00  0.00           C
ATOM   4202  HG1 MET A 261      16.017   3.423  34.457  1.00  0.00
ATOM   4203  HG2 MET A 261      17.126   3.242  35.642  1.00  0.00
ATOM   4204  SD  MET A 261      16.017   1.189  35.241  1.00  0.00           S
ATOM   4205  CE  MET A 261      16.356   0.743  36.941  1.00  0.00           C
ATOM   4206  HE1 MET A 261      16.280  -0.248  37.047  1.00  0.00
ATOM   4207  HE2 MET A 261      17.281   1.034  37.184  1.00  0.00
ATOM   4208  HE3 MET A 261      15.696   1.194  37.542  1.00  0.00
ATOM   4209  C   MET A 261      14.865   5.683  35.093  1.00  0.00           C
ATOM   4210  O   MET A 261      13.687   5.640  34.767  1.00  0.00           O
ATOM   4211  N   ASP A 262      15.813   6.243  34.337  1.00  0.00           N
ATOM   4212  HN  ASP A 262      16.750   6.294  34.683  1.00  0.00
ATOM   4213  CA  ASP A 262      15.503   6.792  32.991  1.00  0.00           C
ATOM   4214  HA  ASP A 262      14.529   7.020  32.977  1.00  0.00
ATOM   4215  CB  ASP A 262      16.250   8.104  32.733  1.00  0.00           C
ATOM   4216  HB1 ASP A 262      16.008   8.434  31.821  1.00  0.00
ATOM   4217  HB2 ASP A 262      15.962   8.774  33.417  1.00  0.00
ATOM   4218  CG  ASP A 262      17.761   7.975  32.804  1.00  0.00           C
ATOM   4219  OD1 ASP A 262      18.274   6.931  32.332  1.00  0.00           O
ATOM   4220  OD2 ASP A 262      18.416   8.925  33.341  1.00  0.00           O
ATOM   4221  C   ASP A 262      15.769   5.719  31.922  1.00  0.00           C
ATOM   4222  O   ASP A 262      16.310   4.655  32.208  1.00  0.00           O
ATOM   4223  N   PHE A 263      15.354   6.014  30.691  1.00  0.00           N
ATOM   4224  HN  PHE A 263      15.135   6.965  30.475  1.00  0.00
ATOM   4225  CA  PHE A 263      15.206   5.010  29.646  1.00  0.00           C
ATOM   4226  HA  PHE A 263      14.570   4.332  30.013  1.00  0.00
ATOM   4227  CB  PHE A 263      14.600   5.626  28.389  1.00  0.00           C
ATOM   4228  HB1 PHE A 263      15.031   6.514  28.229  1.00  0.00
ATOM   4229  HB2 PHE A 263      14.786   5.020  27.615  1.00  0.00
ATOM   4230  CG  PHE A 263      13.111   5.838  28.469  1.00  0.00           C
ATOM   4231  CD1 PHE A 263      12.230   4.790  28.237  1.00  0.00           C
ATOM   4232  HD1 PHE A 263      12.588   3.882  28.019  1.00  0.00
ATOM   4233  CE1 PHE A 263      10.855   4.989  28.305  1.00  0.00           C
ATOM   4234  HE1 PHE A 263      10.232   4.228  28.124  1.00  0.00
ATOM   4235  CZ  PHE A 263      10.348   6.230  28.621  1.00  0.00           C
ATOM   4236  HZ  PHE A 263       9.359   6.371  28.674  1.00  0.00
ATOM   4237  CD2 PHE A 263      12.584   7.082  28.799  1.00  0.00           C
ATOM   4238  HD2 PHE A 263      13.201   7.845  28.992  1.00  0.00
ATOM   4239  CE2 PHE A 263      11.211   7.275  28.863  1.00  0.00           C
ATOM   4240  HE2 PHE A 263      10.846   8.179  29.086  1.00  0.00
ATOM   4241  C   PHE A 263      16.541   4.325  29.352  1.00  0.00           C
ATOM   4242  O   PHE A 263      16.600   3.117  29.200  1.00  0.00           O
ATOM   4243  N   GLU A 264      17.615   5.113  29.285  1.00  0.00           N
ATOM   4244  HN  GLU A 264      17.532   6.075  29.545  1.00  0.00
ATOM   4245  CA  GLU A 264      18.912   4.603  28.840  1.00  0.00           C
ATOM   4246  HA  GLU A 264      18.782   4.105  27.983  1.00  0.00
ATOM   4247  CB  GLU A 264      19.844   5.782  28.544  1.00  0.00           C
ATOM   4248  HB1 GLU A 264      19.936   6.335  29.372  1.00  0.00
ATOM   4249  HB2 GLU A 264      20.740   5.425  28.281  1.00  0.00
ATOM   4250  CG  GLU A 264      19.320   6.673  27.415  1.00  0.00           C
ATOM   4251  HG1 GLU A 264      18.417   7.007  27.684  1.00  0.00
ATOM   4252  HG2 GLU A 264      19.947   7.446  27.321  1.00  0.00
ATOM   4253  CD  GLU A 264      19.186   6.003  26.040  1.00  0.00           C
ATOM   4254  OE1 GLU A 264      19.805   4.918  25.834  1.00  0.00           O
ATOM   4255  OE2 GLU A 264      18.467   6.566  25.158  1.00  0.00           O
```

```
ATOM   4256  C   GLU A 264      19.457   3.601  29.874  1.00  0.00           C
ATOM   4257  O   GLU A 264      20.071   2.601  29.504  1.00  0.00           O
ATOM   4258  N   ALA A 265      19.190   3.852  31.159  1.00  0.00           N
ATOM   4259  HN  ALA A 265      18.738   4.712  31.397  1.00  0.00
ATOM   4260  CA  ALA A 265      19.531   2.923  32.226  1.00  0.00           C
ATOM   4261  HA  ALA A 265      20.514   2.752  32.161  1.00  0.00
ATOM   4262  CB  ALA A 265      19.235   3.547  33.567  1.00  0.00           C
ATOM   4263  HB1 ALA A 265      19.472   2.903  34.294  1.00  0.00
ATOM   4264  HB2 ALA A 265      19.775   4.382  33.673  1.00  0.00
ATOM   4265  HB3 ALA A 265      18.262   3.770  33.623  1.00  0.00
ATOM   4266  C   ALA A 265      18.751   1.616  32.045  1.00  0.00           C
ATOM   4267  O   ALA A 265      19.303   0.536  32.223  1.00  0.00           O
ATOM   4268  N   PHE A 266      17.460   1.746  31.700  1.00  0.00           N
ATOM   4269  HN  PHE A 266      17.109   2.668  31.537  1.00  0.00
ATOM   4270  CA  PHE A 266      16.532   0.622  31.547  1.00  0.00           C
ATOM   4271  HA  PHE A 266      16.622   0.076  32.380  1.00  0.00
ATOM   4272  CB  PHE A 266      15.087   1.127  31.442  1.00  0.00           C
ATOM   4273  HB1 PHE A 266      14.781   1.406  32.352  1.00  0.00
ATOM   4274  HB2 PHE A 266      15.070   1.916  30.828  1.00  0.00
ATOM   4275  CG  PHE A 266      14.101   0.113  30.916  1.00  0.00           C
ATOM   4276  CD1 PHE A 266      13.781  -1.024  31.641  1.00  0.00           C
ATOM   4277  HD1 PHE A 266      14.175  -1.154  32.551  1.00  0.00
ATOM   4278  CE1 PHE A 266      12.925  -1.984  31.126  1.00  0.00           C
ATOM   4279  HE1 PHE A 266      12.745  -2.822  31.641  1.00  0.00
ATOM   4280  CZ  PHE A 266      12.328  -1.783  29.914  1.00  0.00           C
ATOM   4281  HZ  PHE A 266      11.671  -2.449  29.561  1.00  0.00
ATOM   4282  CD2 PHE A 266      13.526   0.275  29.675  1.00  0.00           C
ATOM   4283  HD2 PHE A 266      13.756   1.077  29.124  1.00  0.00
ATOM   4284  CE2 PHE A 266      12.634  -0.661  29.183  1.00  0.00           C
ATOM   4285  HE2 PHE A 266      12.209  -0.520  28.289  1.00  0.00
ATOM   4286  C   PHE A 266      16.924  -0.245  30.337  1.00  0.00           C
ATOM   4287  O   PHE A 266      17.027  -1.457  30.473  1.00  0.00           O
ATOM   4288  N   PHE A 267      17.131   0.374  29.167  1.00  0.00           N
ATOM   4289  HN  PHE A 267      17.075   1.372  29.137  1.00  0.00
ATOM   4290  CA  PHE A 267      17.442  -0.363  27.906  1.00  0.00           C
ATOM   4291  HA  PHE A 267      16.644  -0.947  27.755  1.00  0.00
ATOM   4292  CB  PHE A 267      17.624   0.568  26.700  1.00  0.00           C
ATOM   4293  HB1 PHE A 267      18.346   1.226  26.913  1.00  0.00
ATOM   4294  HB2 PHE A 267      17.897   0.017  25.912  1.00  0.00
ATOM   4295  CG  PHE A 267      16.403   1.349  26.304  1.00  0.00           C
ATOM   4296  CD1 PHE A 267      15.140   0.774  26.346  1.00  0.00           C
ATOM   4297  HD1 PHE A 267      15.041  -0.191  26.591  1.00  0.00
ATOM   4298  CE1 PHE A 267      14.020   1.526  26.052  1.00  0.00           C
ATOM   4299  HE1 PHE A 267      13.112   1.113  26.117  1.00  0.00
ATOM   4300  CZ  PHE A 267      14.148   2.840  25.671  1.00  0.00           C
ATOM   4301  HZ  PHE A 267      13.332   3.378  25.458  1.00  0.00
ATOM   4302  CD2 PHE A 267      16.512   2.681  25.924  1.00  0.00           C
ATOM   4303  HD2 PHE A 267      17.413   3.114  25.897  1.00  0.00
ATOM   4304  CE2 PHE A 267      15.388   3.418  25.581  1.00  0.00           C
ATOM   4305  HE2 PHE A 267      15.481   4.364  25.270  1.00  0.00
ATOM   4306  C   PHE A 267      18.715  -1.194  28.067  1.00  0.00           C
ATOM   4307  O   PHE A 267      18.836  -2.257  27.479  1.00  0.00           O
ATOM   4308  N   SER A 268      19.655  -0.675  28.855  1.00  0.00           N
ATOM   4309  HN  SER A 268      19.437   0.146  29.383  1.00  0.00
ATOM   4310  CA  SER A 268      20.976  -1.246  28.980  1.00  0.00           C
ATOM   4311  HA  SER A 268      21.044  -1.952  28.275  1.00  0.00
ATOM   4312  CB  SER A 268      22.024  -0.190  28.717  1.00  0.00           C
ATOM   4313  HB1 SER A 268      22.925  -0.611  28.609  1.00  0.00
ATOM   4314  HB2 SER A 268      21.798   0.330  27.893  1.00  0.00
ATOM   4315  OG  SER A 268      22.093   0.722  29.807  1.00  0.00           O
ATOM   4316  HG1 SER A 268      22.790   1.414  29.619  1.00  0.00
ATOM   4317  C   SER A 268      21.153  -1.872  30.368  1.00  0.00           C
ATOM   4318  O   SER A 268      22.283  -2.040  30.834  1.00  0.00           O
ATOM   4319  N   HIS A 269      20.036  -2.226  31.019  1.00  0.00           N
ATOM   4320  HN  HIS A 269      19.159  -2.173  30.541  1.00  0.00
ATOM   4321  CA  HIS A 269      20.061  -2.688  32.409  1.00  0.00           C
ATOM   4322  HA  HIS A 269      20.638  -2.038  32.903  1.00  0.00
ATOM   4323  CB  HIS A 269      18.666  -2.671  33.058  1.00  0.00           C
ATOM   4324  HB1 HIS A 269      18.224  -1.794  32.870  1.00  0.00
ATOM   4325  HB2 HIS A 269      18.114  -3.411  32.675  1.00  0.00
ATOM   4326  ND1 HIS A 269      18.899  -4.097  35.125  1.00  0.00           N
```

```
ATOM   4327  CG  HIS A 269      18.716  -2.853  34.538  1.00  0.00           C
ATOM   4328  CE1 HIS A 269      18.947  -3.951  36.434  1.00  0.00           C
ATOM   4329  HE1 HIS A 269      19.071  -4.687  37.100  1.00  0.00
ATOM   4330  NE2 HIS A 269      18.802  -2.648  36.723  1.00  0.00           N
ATOM   4331  HE2 HIS A 269      18.792  -2.251  37.641  1.00  0.00
ATOM   4332  CD2 HIS A 269      18.670  -1.956  35.554  1.00  0.00           C
ATOM   4333  HD2 HIS A 269      18.559  -0.966  35.463  1.00  0.00
ATOM   4334  C   HIS A 269      20.680  -4.070  32.464  1.00  0.00           C
ATOM   4335  O   HIS A 269      20.353  -4.925  31.650  1.00  0.00           O
ATOM   4336  N   PRO A 270      21.595  -4.318  33.426  1.00  0.00           N
ATOM   4337  CD  PRO A 270      22.114  -3.317  34.372  1.00  0.00           C
ATOM   4338  HD1 PRO A 270      21.373  -2.968  34.946  1.00  0.00
ATOM   4339  HD2 PRO A 270      22.536  -2.559  33.875  1.00  0.00
ATOM   4340  CA  PRO A 270      22.231  -5.623  33.586  1.00  0.00           C
ATOM   4341  HA  PRO A 270      22.875  -5.742  32.831  1.00  0.00
ATOM   4342  CB  PRO A 270      22.803  -5.519  34.998  1.00  0.00           C
ATOM   4343  HB1 PRO A 270      22.128  -5.812  35.675  1.00  0.00
ATOM   4344  HB2 PRO A 270      23.628  -6.077  35.087  1.00  0.00
ATOM   4345  CG  PRO A 270      23.138  -4.067  35.197  1.00  0.00           C
ATOM   4346  HG1 PRO A 270      23.064  -3.820  36.163  1.00  0.00
ATOM   4347  HG2 PRO A 270      24.064  -3.874  34.873  1.00  0.00
ATOM   4348  C   PRO A 270      21.301  -6.848  33.499  1.00  0.00           C
ATOM   4349  O   PRO A 270      21.692  -7.880  32.928  1.00  0.00           O
ATOM   4350  N   PHE A 271      20.084  -6.727  34.047  1.00  0.00           N
ATOM   4351  HN  PHE A 271      19.818  -5.847  34.440  1.00  0.00
ATOM   4352  CA  PHE A 271      19.133  -7.834  34.091  1.00  0.00           C
ATOM   4353  HA  PHE A 271      19.611  -8.546  34.605  1.00  0.00
ATOM   4354  CB  PHE A 271      17.851  -7.413  34.807  1.00  0.00           C
ATOM   4355  HB1 PHE A 271      18.067  -7.198  35.759  1.00  0.00
ATOM   4356  HB2 PHE A 271      17.480  -6.600  34.358  1.00  0.00
ATOM   4357  CG  PHE A 271      16.793  -8.477  34.796  1.00  0.00           C
ATOM   4358  CD1 PHE A 271      16.944  -9.636  35.550  1.00  0.00           C
ATOM   4359  HD1 PHE A 271      17.755  -9.748  36.124  1.00  0.00
ATOM   4360  CE1 PHE A 271      15.988 -10.636  35.517  1.00  0.00           C
ATOM   4361  HE1 PHE A 271      16.105 -11.461  36.070  1.00  0.00
ATOM   4362  CZ  PHE A 271      14.881 -10.492  34.724  1.00  0.00           C
ATOM   4363  HZ  PHE A 271      14.180 -11.205  34.707  1.00  0.00
ATOM   4364  CD2 PHE A 271      15.693  -8.364  33.970  1.00  0.00           C
ATOM   4365  HD2 PHE A 271      15.589  -7.559  33.386  1.00  0.00
ATOM   4366  CE2 PHE A 271      14.734  -9.359  33.947  1.00  0.00           C
ATOM   4367  HE2 PHE A 271      13.928  -9.258  33.364  1.00  0.00
ATOM   4368  C   PHE A 271      18.775  -8.336  32.678  1.00  0.00           C
ATOM   4369  O   PHE A 271      18.373  -9.480  32.525  1.00  0.00           O
ATOM   4370  N   LEU A 272      18.898  -7.466  31.668  1.00  0.00           N
ATOM   4371  HN  LEU A 272      19.361  -6.596  31.836  1.00  0.00
ATOM   4372  CA  LEU A 272      18.381  -7.741  30.329  1.00  0.00           C
ATOM   4373  HA  LEU A 272      17.591  -8.340  30.463  1.00  0.00
ATOM   4374  CB  LEU A 272      17.983  -6.422  29.666  1.00  0.00           C
ATOM   4375  HB1 LEU A 272      18.682  -5.743  29.890  1.00  0.00
ATOM   4376  HB2 LEU A 272      17.966  -6.568  28.677  1.00  0.00
ATOM   4377  CG  LEU A 272      16.635  -5.846  30.069  1.00  0.00           C
ATOM   4378  HG  LEU A 272      16.618  -5.711  31.060  1.00  0.00
ATOM   4379  CD1 LEU A 272      16.428  -4.504  29.400  1.00  0.00           C
ATOM   4380 1HD1 LEU A 272      15.540  -4.131  29.669  1.00  0.00
ATOM   4381 2HD1 LEU A 272      17.153  -3.876  29.682  1.00  0.00
ATOM   4382 3HD1 LEU A 272      16.454  -4.619  28.407  1.00  0.00
ATOM   4383  CD2 LEU A 272      15.519  -6.806  29.716  1.00  0.00           C
ATOM   4384 1HD2 LEU A 272      14.641  -6.411  29.988  1.00  0.00
ATOM   4385 2HD2 LEU A 272      15.520  -6.970  28.729  1.00  0.00
ATOM   4386 3HD2 LEU A 272      15.659  -7.671  30.198  1.00  0.00
ATOM   4387  C   LEU A 272      19.426  -8.452  29.456  1.00  0.00           C
ATOM   4388  O   LEU A 272      19.105  -8.808  28.317  1.00  0.00           O
ATOM   4389  N   GLU A 273      20.666  -8.622  29.946  1.00  0.00           N
ATOM   4390  HN  GLU A 273      20.831  -8.464  30.919  1.00  0.00
ATOM   4391  CA  GLU A 273      21.785  -9.035  29.087  1.00  0.00           C
ATOM   4392  HA  GLU A 273      21.366  -9.467  28.288  1.00  0.00
ATOM   4393  CB  GLU A 273      22.610  -7.818  28.649  1.00  0.00           C
ATOM   4394  HB1 GLU A 273      23.410  -8.139  28.141  1.00  0.00
ATOM   4395  HB2 GLU A 273      22.045  -7.245  28.054  1.00  0.00
ATOM   4396  CG  GLU A 273      23.095  -6.965  29.805  1.00  0.00           C
ATOM   4397  HG1 GLU A 273      22.319  -6.791  30.411  1.00  0.00
```

```
ATOM   4398  HG2 GLU A 273      23.795  -7.480  30.299  1.00  0.00
ATOM   4399  CD  GLU A 273      23.688  -5.627  29.397  1.00  0.00           C
ATOM   4400  OE1 GLU A 273      23.451  -5.197  28.244  1.00  0.00           O
ATOM   4401  OE2 GLU A 273      24.408  -5.030  30.221  1.00  0.00           O
ATOM   4402  C   GLU A 273      22.676 -10.049  29.804  1.00  0.00           C
ATOM   4403  O   GLU A 273      23.847  -9.755  30.071  1.00  0.00           O
ATOM   4404  N   GLN A 274      22.161 -11.258  30.057  1.00  0.00           N
ATOM   4405  HN  GLN A 274      21.225 -11.470  29.776  1.00  0.00
ATOM   4406  CA  GLN A 274      22.975 -12.283  30.752  1.00  0.00           C
ATOM   4407  HA  GLN A 274      23.907 -12.204  30.399  1.00  0.00
ATOM   4408  CB  GLN A 274      22.973 -12.001  32.254  1.00  0.00           C
ATOM   4409  HB1 GLN A 274      23.414 -12.767  32.721  1.00  0.00
ATOM   4410  HB2 GLN A 274      23.495 -11.164  32.420  1.00  0.00
ATOM   4411  CG  GLN A 274      21.583 -11.820  32.844  1.00  0.00           C
ATOM   4412  HG1 GLN A 274      21.081 -11.146  32.303  1.00  0.00
ATOM   4413  HG2 GLN A 274      21.099 -12.694  32.820  1.00  0.00
ATOM   4414  CD  GLN A 274      21.657 -11.342  34.272  1.00  0.00           C
ATOM   4415  OE1 GLN A 274      20.967 -11.841  35.159  1.00  0.00           O
ATOM   4416  NE2 GLN A 274      22.523 -10.373  34.508  1.00  0.00           N
ATOM   4417 1HE2 GLN A 274      23.077 -10.005  33.761  1.00  0.00
ATOM   4418 2HE2 GLN A 274      22.624 -10.007  35.433  1.00  0.00
ATOM   4419  C   GLN A 274      22.480 -13.701  30.441  1.00  0.00           C
ATOM   4420  O   GLN A 274      21.413 -13.883  29.860  1.00  0.00           O
ATOM   4421  N   GLY A 275      23.288 -14.696  30.845  1.00  0.00           N
ATOM   4422  HN  GLY A 275      24.130 -14.442  31.321  1.00  0.00
ATOM   4423  CA  GLY A 275      23.027 -16.147  30.639  1.00  0.00           C
ATOM   4424  HA1 GLY A 275      23.663 -16.481  29.944  1.00  0.00
ATOM   4425  HA2 GLY A 275      23.201 -16.619  31.503  1.00  0.00
ATOM   4426  C   GLY A 275      21.601 -16.440  30.190  1.00  0.00           C
ATOM   4427  OT1 GLY A 275      20.671 -16.461  31.001  1.00  0.00           O
ATOM   4428  OT2 GLY A 275      21.279 -17.641  29.989  1.00  0.00           O
ATOM   4429  HT2 GLY A 275      20.322 -17.686  29.701  1.00  0.00
TER
ATOM      1  C   LIG     1      10.409  -8.574   0.191  1.00  0.00      LIG
ATOM      2  C1  LIG     1       9.283  -8.184  -0.519  1.00  0.00      LIG
ATOM      3  C2  LIG     1       9.197  -6.879  -1.014  1.00  0.00      LIG
ATOM      4  C3  LIG     1      10.230  -5.950  -0.790  1.00  0.00      LIG
ATOM      5  C4  LIG     1      11.357  -6.374  -0.065  1.00  0.00      LIG
ATOM      6  C5  LIG     1      11.448  -7.683   0.420  1.00  0.00      LIG
ATOM      7  C6  LIG     1      10.138  -4.569  -1.305  1.00  0.00      LIG
ATOM      8  C7  LIG     1       8.979  -3.901  -1.452  1.00  0.00      LIG
ATOM      9  N   LIG     1       8.955  -2.589  -1.909  1.00  0.00      LIG
ATOM     10  N1  LIG     1       7.859  -1.812  -2.097  1.00  0.00      LIG
ATOM     11  C8  LIG     1       8.355  -0.640  -2.496  1.00  0.00      LIG
ATOM     12  C9  LIG     1       9.760  -0.637  -2.583  1.00  0.00      LIG
ATOM     13  C10 LIG     1      10.139  -1.912  -2.215  1.00  0.00      LIG
ATOM     14  N2  LIG     1      11.358  -2.527  -2.110  1.00  0.00      LIG
ATOM     15  C11 LIG     1      11.343  -3.759  -1.696  1.00  0.00      LIG
ATOM     16  C12 LIG     1      10.589   0.512  -2.960  1.00  0.00      LIG
ATOM     17  C13 LIG     1      10.108   1.756  -3.315  1.00  0.00      LIG
ATOM     18  C14 LIG     1      12.576   1.722  -3.237  1.00  0.00      LIG
ATOM     19  C15 LIG     1      12.018   0.486  -2.974  1.00  0.00      LIG
ATOM     20  F   LIG     1      10.494  -9.824   0.661  1.00  0.00      LIG
ATOM     21  S   LIG     1      11.362   2.877  -3.615  1.00  0.00      LIG
ATOM     22  C16 LIG     1      13.982   2.088  -3.222  1.00  0.00      LIG
ATOM     23  O   LIG     1      14.904   1.284  -3.403  1.00  0.00      LIG
ATOM     24  N3  LIG     1      14.231   3.406  -2.925  1.00  0.00      LIG
ATOM     25  C17 LIG     1      15.586   3.914  -2.679  1.00  0.00      LIG
ATOM     26  C18 LIG     1      15.552   5.392  -3.029  1.00  0.00      LIG
ATOM     27  C19 LIG     1      16.789   6.257  -3.030  1.00  0.00      LIG
ATOM     28  C20 LIG     1      16.154   5.874  -4.333  1.00  0.00      LIG
ATOM     29  C21 LIG     1      16.133   3.609  -1.252  1.00  0.00      LIG
ATOM     30  F1  LIG     1      15.240   2.920  -0.501  1.00  0.00      LIG
ATOM     31  F2  LIG     1      17.270   2.875  -1.306  1.00  0.00      LIG
ATOM     32  F3  LIG     1      16.424   4.738  -0.560  1.00  0.00      LIG
ATOM     33  H   LIG     1       8.480  -8.894  -0.687  1.00  0.00      LIG
ATOM     34  H1  LIG     1       8.309  -6.606  -1.581  1.00  0.00      LIG
ATOM     35  H2  LIG     1      12.179  -5.695   0.151  1.00  0.00      LIG
ATOM     36  H3  LIG     1      12.320  -8.008   0.978  1.00  0.00      LIG
ATOM     37  H4  LIG     1       8.006  -4.309  -1.216  1.00  0.00      LIG
ATOM     38  H5  LIG     1       7.660   0.165  -2.699  1.00  0.00      LIG
```

```
ATOM     39  H6  LIG     1       12.313  -4.278  -1.631  1.00  0.00       LIG
ATOM     40  H7  LIG     1        9.081   2.079  -3.403  1.00  0.00       LIG
ATOM     41  H8  LIG     1       12.628  -0.381  -2.740  1.00  0.00       LIG
ATOM     42  H9  LIG     1       13.446   3.925  -2.538  1.00  0.00       LIG
ATOM     43  H10 LIG     1       16.248   3.397  -3.386  1.00  0.00       LIG
ATOM     44  H11 LIG     1       14.638   5.876  -2.696  1.00  0.00       LIG
ATOM     45  H12 LIG     1       17.745   5.813  -2.777  1.00  0.00       LIG
ATOM     46  H13 LIG     1       16.682   7.286  -2.702  1.00  0.00       LIG
ATOM     47  H14 LIG     1       16.711   5.189  -4.964  1.00  0.00       LIG
ATOM     48  H15 LIG     1       15.626   6.643  -4.888  1.00  0.00       LIG
ENDMDL
```
